# Identification of pseudotetraivprolide from *Pseudomonas entomophila* give novel insights into the biosynthesis of detoxin/rimosamide-like anti-antibiotics

**DOI:** 10.1101/2025.06.17.660182

**Authors:** Edna Bode, Julia Büllesbach, Kevin Bauer, Yan-Ni Shi, Simon Reiners, Ziheng Cui, Petra Happel, Yi-Ming Shi, Uli Kazmaier, Martin Grininger, Helge B. Bode

## Abstract

Novel variants of known natural product (NP) classes might guide our understanding of biosynthesis, mode of action and potential application as drugs for all members of such NP classes. Here we describe a novel member of the widespread detoxine/rimosamide-like (DRL) natural products named pseudotetraivprolide from *Pseudomonas* strains, which has the characteristic DRL-activity of protecting *Bacillus cereus* against the antibiotic blasticidin S. The generation of several deletion and complemented mutants, heterologous expression experiments, identification and structure elucidation of several derivatives, chemical synthesis of main derivatives, enzymatic characterization of individual biochemical steps and detailed homology modelling of enzyme complexes were performed. This allowed us to show the primary metabolism-derived malonyl CoA-ACP transacylase (AT) FabD acting as *trans*-AT in the biosynthesis, suggest an order for all late-stage modifications and provide a function for the three conserved hypothetical proteins PipDFG acting as last-step acetylation complex thereby stabilizing the final product.

## Introduction

Bacteria carry biosynthesis gene clusters (BGCs) responsible for the production of bioactive natural products (NP). While we are able to speed up identification of NPs and their BGCs based on progress in mass spectrometry and bioinformatics tools like antiSMASH^[1,2]^, respectively, we still do not understand well, how and for which ecological function they were evolved. That they were developed for specific ecological functions is indicative since related BBCs producing structurally similar NPs often share a set of core genes but differ in the presence of accessory genes involved in modifying the core structure. Examples are the biosynthesis of glycopeptide antibiotics^[3]^, pyrrolizidine alkaloids^[4]^ or detoxin/rimosamide^[5]^ NP families among several other NP classes. In the light of applying NPs as clinically used drugs, the knowledge of such novel NP derivatives increases the chances of either finding suitable drugs directly or diversifying them by using the accessory genes or their suggested modifications for compound optimization. Furthermore, although we are able to rather quickly identify promising BGCs of interest involved in the biosynthesis of desired NP families from thousands of available genomes using a set of great tools^[6–11]^, we still have to access both the BGC and the NP to confirm their structure, biosynthesis and function. Here, heterologous expression of complete BGCs following multiple different methods,^[12– 15]^ activation of BGC expression using variation of the culture conditions^[16]^ including its modern elicitor-based variants,^[17,18]^ or BGC activation using promoter exchange approaches^[19–21]^ have been applied successfully.

In our aim to identify novel NPs from entomopathogenic bacteria we discovered a BGC in *Pseudomonas entomophila* L48 with similarity to BGCs of the detoxin/rimosamide family of NPs. Detoxin/rimosamide-like (DRL) NPs are named after the first described members of this diverse NP class^[22,23]^. They are produced by hybrids of non-ribosomal peptide synthetase (NRPS) and polyketide synthase (PKS) found in a variety of Gram-positive *Streptomyces*^[5,23,24]^ and Gram-negative species like *Pseudomonas, Chitinimonas*^[25,26]^ and *Pseudovibrio*^[27]^. They have been described as anti-antibiotics protecting *Bacillus cereus* against the nucleoside antibiotic blasticidin S^[28,29]^, most likely by blocking peptide ABC-importers or oligopeptide transporters^[30,31]^.

We activated the respective BGC using the easyPACId promoter exchange approach^[20]^ that enabled us to analyze the biosynthesis and structure of the produced NPs. Here we show the structure of all main derivatives, the chemical synthesis of two main derivatives and the detailed biosynthesis of a new DRL from *P. entomophila* named pseudotetraivprolide. We elucidated several unusual but conserved DRL features concerning their biosynthesis, including novel insights in the activity of the *trans*-AT polyketide synthase (PKS) module requiring the primary metabolism malonyl CoA-ACP transacylase FabD. We also elucidated the function and interaction of three hypothetical proteins, present in several DRL producing BGCs involved in late-stage acetylation and stabilization of the final product.

## Results and Discussion

### Identification of a new type of detoxin/rimosamide BGC

*P. entomophila* L48^[32]^ is the producer of several known NPs: The lipodepsipeptide entolysin^[33]^, the iron siderophores pseudomonine^[34]^ and pyoverdine^[34]^, pyreudione^[35]^ and the pseudomonas virulence factor (pvf)^[36]^ are all derived from NRPS-encoding BGCs, while the bioactive oxazole-containing labradorins^[37]^ are produced NRPS-independent.

Additionally, a BGC with similarity to the detoxin/rimosamide-like (DRL) NP family is present in the genome, which seems to represent a new member of this widespread NP family. It is composed of eight genes (PSEEN_RS12600-RS12635) *pipA*-*H* (Fig. 1a, Table S1.1).

**Figure 1.**
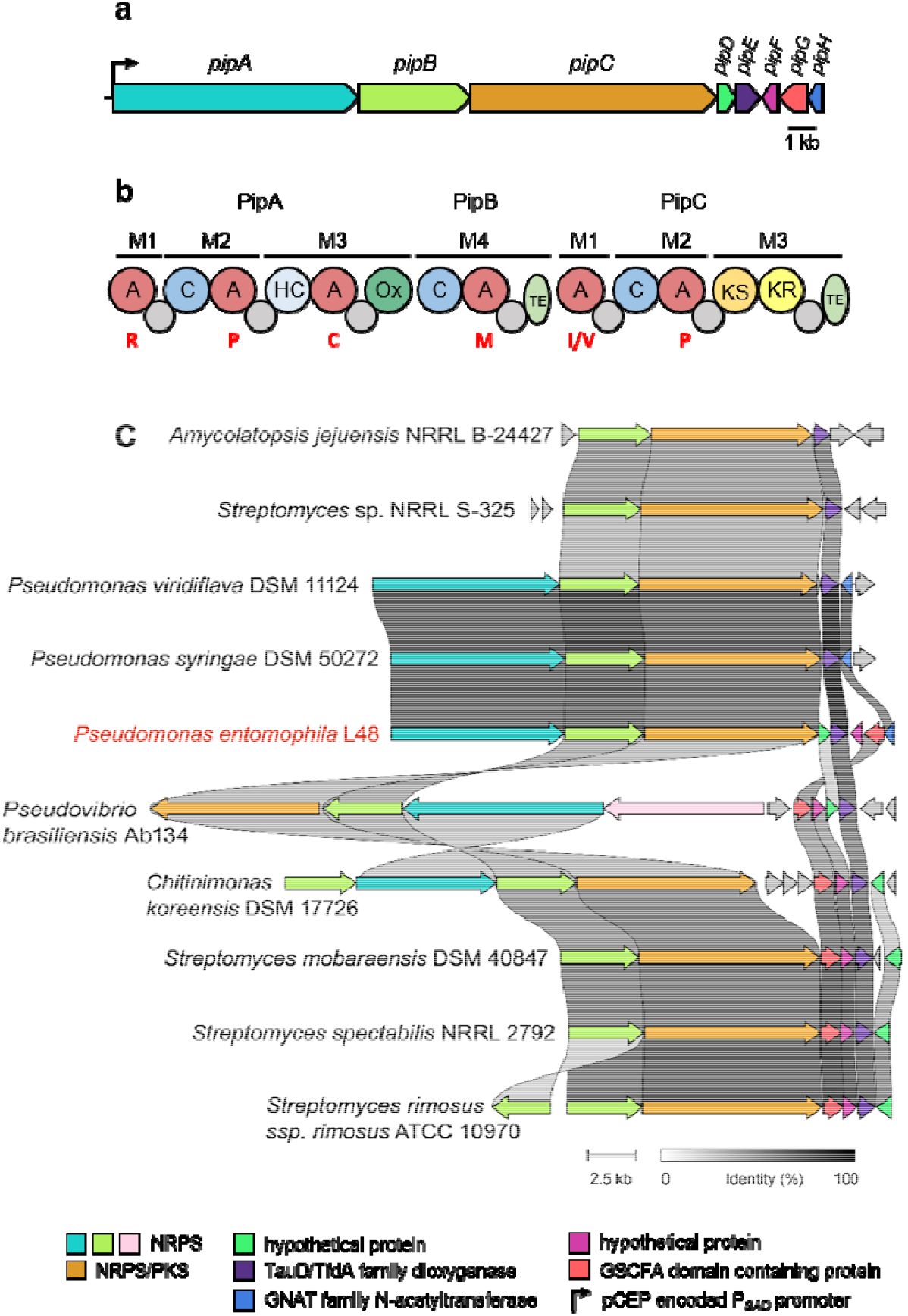
Biosynthetic gene cluster of pseudotetraivprolide (**a**) highlighting domain organization of the NRPSs PipAB and the NRPS/PKS hybrid PipC (**b**). Predicted domains: Adenylation domain (A) with substrate specificities in one-letter amino acid code (red), Thiolation domain (grey circle), Condensation domains (blue), Heterocyclisation domain (HC) (light blue), Oxidation domain (green), Thioesterase domain (TE-domains) (olive), Ketosynthase domain (KS) (orange) and Ketoreductase domain (KR) (yellow). Alignment of DRL BGCs compared to the pseudotetraivprolide BGC of *P. entomophila* L48 (red) using CAGECAT analysis (**c**).

The two non-ribosomal peptide synthetases (NRPS) PipAB are involved in producing a tetrapeptide^[19]^. The signature NRPS/PKS hybrid PipC, characteristic for all DRL BGCs is responsible for the elongated dipeptide unit (Fig. 1b). The distribution of the *pipA*-*H* BGC was analysed via CompArative GEne Cluster Analysis Toolbox CAGECAT^[38]^. The cluster is highly abundant in *Pseudomonas* as depicted in Fig 1c. Characteristic for the *pip* BGC in the family of Pseudomonadaceae is the combination of two NRPS PipAB and one PKS/NRPS PipC. Besides the highly conserved dioxygenase PipE, present in all DRL BGCs, the *pip* BGC encodes four additional genes with similarity to a N-acetyltransferase (PipH), a GSCFA-domain containing protein (PipG), a member of the SGNH/GDSL hydrolase family, named after the highly N-terminal conserved motif with still unknown enzymatic activity, and two hypothetical proteins PipD and PipF with unknown function. PipD, PipF and PipG always occur together and are only associated with the respective DRL BGC. Whereas *pipG* and *pipF* are always oriented next to each other, localization and orientation of *pipD* might be different (Fig. 1c). Notably, the gene *pipH* is found only in BGCs of *P. entomophila, P. viridiflava, P. syringae* and other *Pseudomonas* strains, which share the same NRPS NRPS/PKS gene composition and do not have a Cstarter condensation domain in PipA.

### Activation of the *pip* BGC

Activation of the BGC in *P. entomophila* via the easyPACId approach^[19,20]^ in order to investigate the respective NP leads to a wealth of different derivatives (Fig. 2, Fig. S2.1-S2.15, Table S1.8). Besides the previously described pseudotetratide (**3b**)^[19]^, additional derivatives of the two separate units **3a**-**3c** and **1a**-**1c**, called ivprolide, and as well as the full-length product pseudotetraivprolide (PIP, **4a**-**4e**) were identified (Fig. 2, Fig. 3), while only minute amounts of **3b** and **4e** were detected in the wild type (Fig. 2b, Fig. S2.4 & S2.10). Products **3** and **4** differ in the modification of their N-terminus, which can be acetylated, and the C-terminal proline ring that can be hydroxylated and further acetylated, respectively. The structures of all derivatives were elucidated based on detailed MS-MS analysis (Fig. S2.1-S2.15) as well as NMR analysis for **1a** (Fig. S2.17-S2.21), **4c** (Fig. S2.22-S2.28, Table S2.1) and **4e** (Fig. S2.29-S2.32 Table S2.2; all in *Supplementary Information-2: Compound identification and structure elucidation*). In addition to the main compounds containing isoleucine, corresponding valine-containing ivprolide (**2a**) and pseudotetraivprolide (**5a**-**5e**) were found as minor derivatives (Fig. S2.1, S2.11-S2.15).

**Figure 2.**
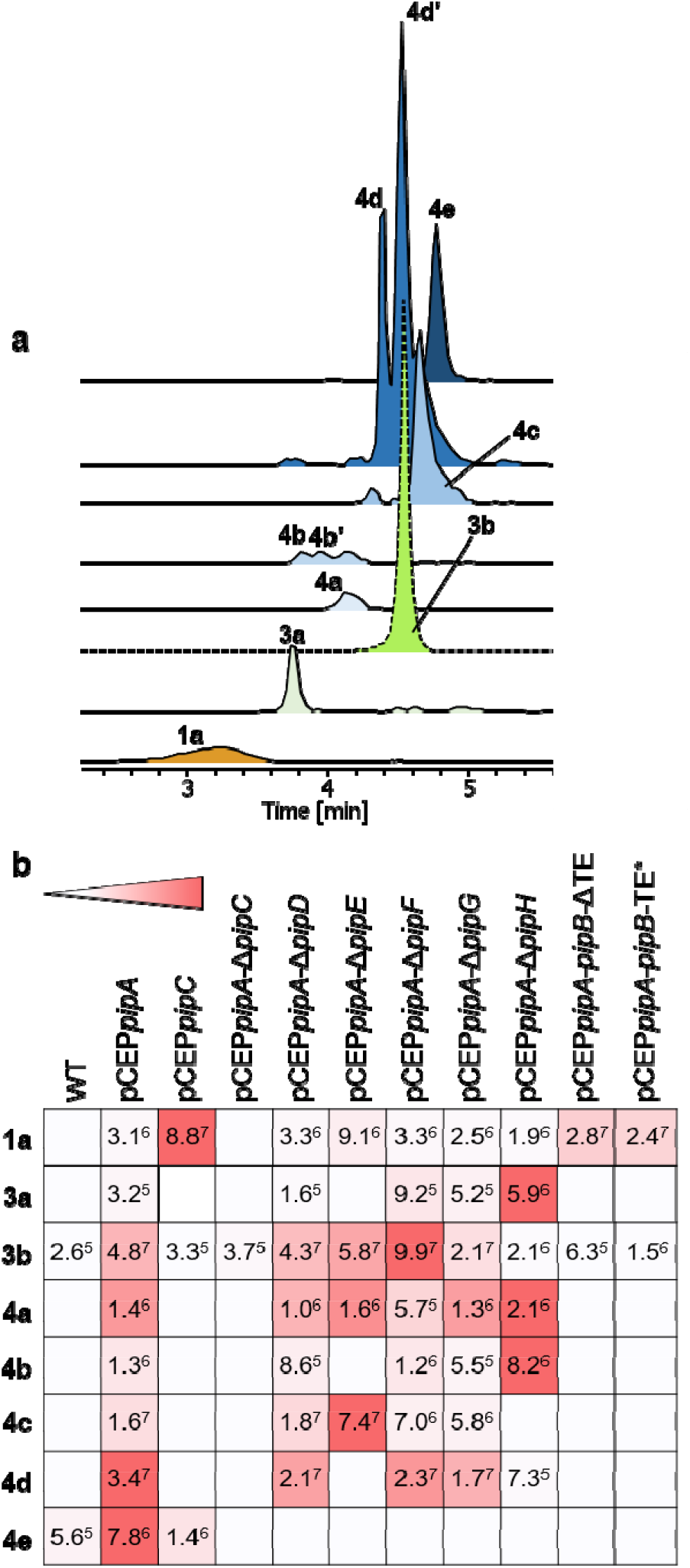
EICs of pseudotetraivprolide derivatives detected in culture extracts of induced pCEP*pipA* mutant, dashed line indicates a 10-fold reduced signal (**a**). Heat map showing the production of pseudotetraivprolide derivatives comparing selected mutants (**b**). Dark red (highest production) to white (no production). Signal intensities from the HPLC/MS analysis are abbreviated as 3.1^6^ for 3.1 × 10^6^.

**Figure 3.**
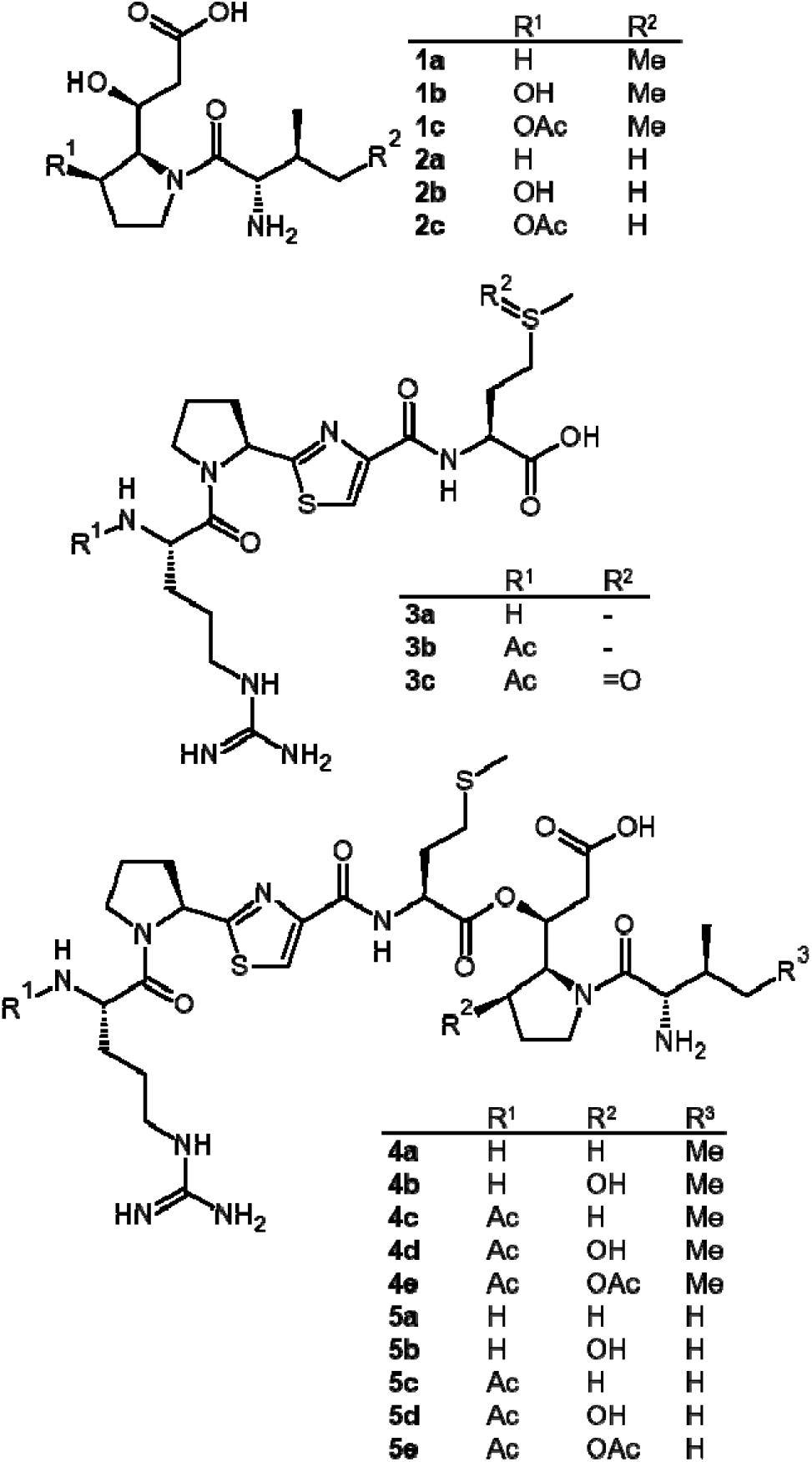
Overview of identified ivprolide (**1 & 2**), pseudotetratide (**3**) and pseudotetraivprolide (**4 & 5**) derivatives from top to bottom.

For full-length PIP derivatives **4b** and **4d** carrying a hydroxy group at the C-terminal proline two isobaric peaks were consistently identified, suggesting two different ester variants most likely formed autocatalytically (Fig. S2.7, S2.9, S2.12 & S2.14). Furthermore, methionine sulfoxide derivatives **3c, 4f, 4g, 5f** and **5g** were also identified under some experimental conditions (Fig. S2.5, S2.7, S2.9, S2.12 & S2.14).

For functional analysis of PIP, we constructed a *pipA* promoter exchange mutant in a low-background WT strain named ΔPELP4 having deletions of BGCs responsible for pseudomonine, entolysin, labradorin, pyreudione and an additional unknown BGC4 (Fig. S1.1a). Such extracts indeed protect *Bacillus cereus* against the antibiotic blasticidin S as previously described for several DRL derivatives^[28]^ (Fig. S1.1b & c). In contrast to pseudovibramide^[27]^ and chitinimide^[26]^, no difference in swarming behaviour was evident for any of the mutants generated compared to the wt strain or PIP producing mutants (data not shown), probably since *P. entomophila* uses entolysin for this purpose^[33]^.

### Chemical synthesis of Pseudotetraivprolides

In order to confirm the overall structure of the full-length derivatives including all stereocenters of the incorporated amino acids, the main product **4c** was synthesized (see *Supplementary Information SI-3: Chemical Synthesis* for details) indeed confirming the structure of the natural product. Briefly, starting from Boc-protected proline **6** the β-hydroxyester **7** (Fig. 4) was prepared according to Greck *et al.*^[39]^ 1,1’-Carbonyldiimidazole (CDI) activation and coupling with potassium methyl malonate (KMM) provided the correspondig β-ketoester which was subjected to an asymmetric Noyori hydrogenation.^[40]^ Standard protecting group manipulations and peptide couplings provided the right-hand side tripeptide **8**. For the left part of the molecule, **6** was converted into thiazole fragment **9** following a procedure from Deng and Taunton’s synthesis of ceratospongamide.^[41]^ After Boc-deprotection, the thiazole building block **9** was coupled with Boc-Orn(Troc)-OH in a HATU mediated peptide coupling with 83% yield. Since, the direct couplings with acetylated ornithine and arginine resulted in strong epimerization at the α-position,^[42,43]^ we used a two-step protocol. After removal of the Boc-carbamate, acetylation with acetic anhydride gave the desired dipeptide **10** in 94% yield. In a last step, the Troc-carbamate was removed to introduce the Di-Boc-guanidine unit. ^[44]^ Deprotection of the two peptide fragments **8** and **11** and coupling with HBTU provided protected pseudotetraivprolid **4c** in acceptable yield. In the last step, a final deprotection was carried out with a cleavage cocktail containing Et3SiH as a scavenger to suppress side reactions induced by the *tert-*butyl cations.^[45]^ Pseudotetraivprolide **5c** was obtained in analogous manner.

**Figure 4.**
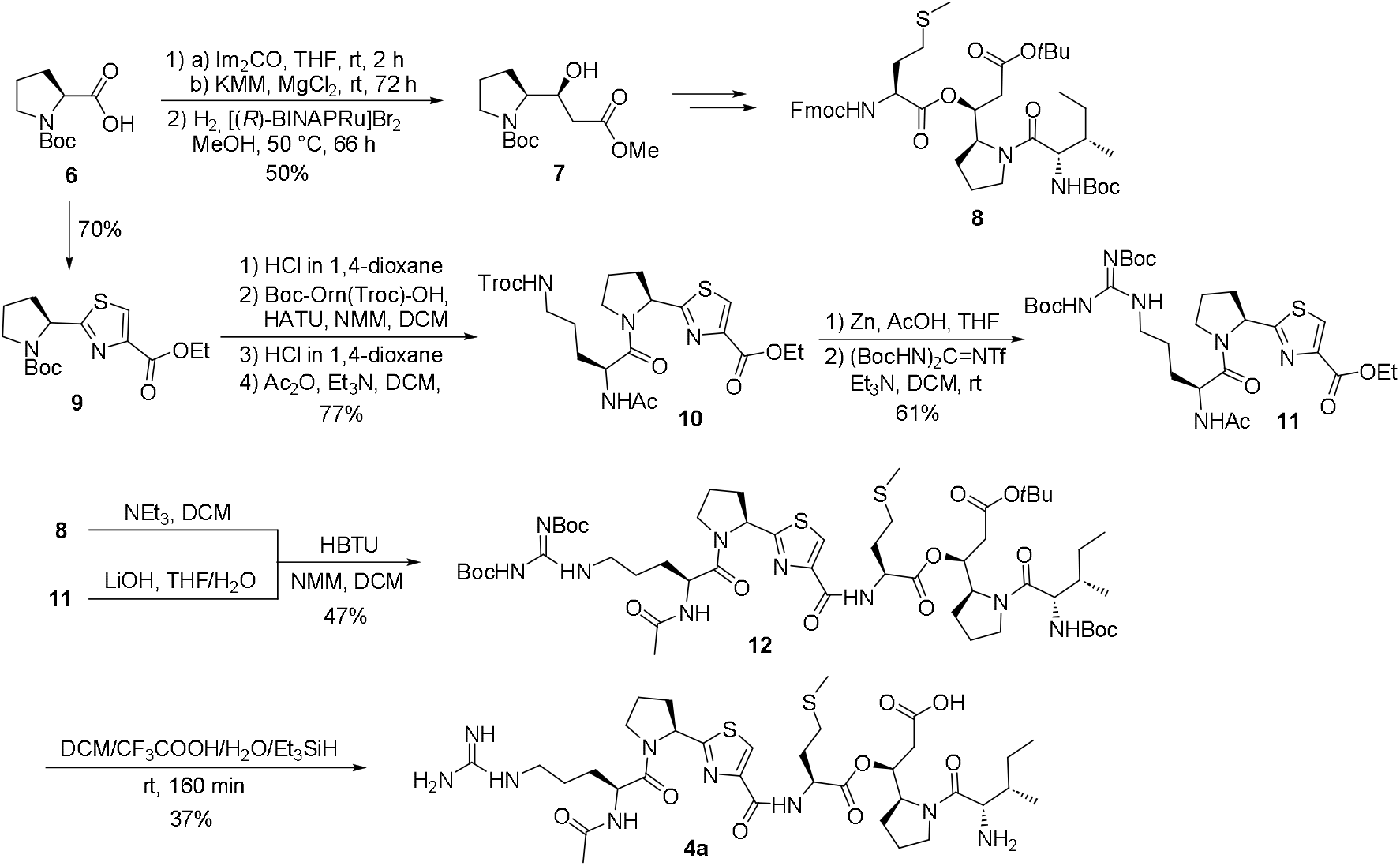
Synthesis of pseudotetraivprolide **4a**. Im (imidazolyl), KMM (potassium methyl malonate), BINAP (2,2’-bis(diphenylphosphino)-1,1’-binaphthyl), HATU (1-[Bis(dimethylamino)methylene]-1H-1,2,3-triazolo[4,5-b]pyridinium 3-oxide hexafluorophosphate), HBTU (3-[Bis(dimethylamino)methyliumyl]-3H-benzotriazol-1-oxide hexafluorophos-phate).

### Functional analysis of *pip* deletions

In order to elucidate the function of all Pip enzymes, the corresponding genes were deleted in-frame in order to avoid polar effects on downstream genes following a detailed HPLC/MS analysis of the produced derivatives (Fig. 2b, Fig. S1.3). All deletions were analysed under the control of the PBAD promoter.

In extracts of deletion mutant Δ*pipC*, only traces of the tetrapeptide **3a** and **3b** were detectable (Fig. S2.3 & S2.4), indicating that their release from the enzyme might be dependent on the presence of PipC. Promoter exchange in front of *pipC* led to overproduction of **1a** and **2a** (Fig. S2.1). A minimal amount of **3b** and **4e** were also produced (Fig. 2b, Fig. S2.4 & S2.10) due to the activity of the natural *pipA* promoter. In the Δ*pipE* mutant an accumulation of all non-hydroxylated compounds **3b, 4a** and **4c** was detected (Fig. S2.4, S2.6 & S2.8). Deletion of either *pipD, pipF* or *pipG* led to accumulation of all precursors that were not *O*-acetylated with **3b** and **4d** as main products (Fig. S2.4 & S2.9). Upon deletion of the N-acetyltransferase encoding gene *pipH* only compounds without an N-terminal acetylation were produced like mainly **3a** and also **4a** and **4b** (Fig. S2.3, S2.6 & S2.7). Traces of **3b** and **4d** in this mutant indicate that there might be another acetyltransferase encoded in the genome catalyzing this reaction. All deletions were successfully complemented with a plasmid-encoded copy of the respective gene (Fig. S1.4) except for *pipG*, which could however be complemented by *pipFG* and *pipFGH* (Fig. S1.5) suggesting that PipF and PipG might build a catalytic complex to enable the final acetylation of **4d**. Deletion of all accessory genes *pipDEFGH* led to accumulation of **3a** and **4a** in the respective mutant, while complementation with plasmid-encoded *pipDEFGH* let to a strong overproduction of **4e** (Fig. S1.6) enabling its preparative isolation and even leading to new derivatives **4h** and **5h** that carry an additional acetyl group at the amine group of Ile or Val, respectively (Fig. S2.16).

### Core structure biosynthesis

PipB and PipC carry a thioesterase (TE) domain and it has been postulated early on that the PipB-TE is responsible for ester bond formation.^[23]^ Indeed, this was shown recently for chitinimide through detailed in vivo and in vitro experiments^[46]^. Similarly, either deletion of the TE domain or exchange of the essential Ser-2856 residue against Ala in PipC resulted in a complete loss of all full-length products while showing accumulation of **1a** and **3b** (Fig. 2b, Fig. S1.7), indicating that **1a** can be released from the PipC-TE by hydrolysis. However, since no hydroxylated or *O*-acetylated products **1b** and **1c** can be detected in the *pipB* TE mutants or when the inducible promoter was inserted in front of *pipC* (Fig. 2b), we postulate that hydroxylation and acetylation predominantly takes place on the full-length products **4**. Heterologous expression of *pipC* or *pipCDE* in *E. coli* showing exclusive production of **1a** (Fig. S1.8) confirmed this. In fact, this timing might be different to that observed in the pseudovibramide^[27]^ and chitinimide^[26]^ biosynthesis, where analogs of **1c** have been found even when parts of the NRPS homologous to PipA were deleted. For the biosynthesis in *P. entomophila* we attribute the presence of **1b** and **1c** in some mutants to hydrolysis of the fully modified full-length products **4b, 4d** and **4e**, since we could also show an increasing amount of both products as well as methyl esters of **3b**, when MeOH was used for compound extraction (data not shown) as also observed for chitinimide^[26]^.

Interestingly production of **1a** from heterologous expression of *pipC* in *E. coli* did not require an acyltransferase (AT), despite the fact that no PipC homolog from any DRL BGCs carries an AT domain. Structural modelling of PipC confirmed the presence of an AT docking domain (ATd) and absence of a classical AT domain (Fig. 5, Fig. S1.9-S1.12) suggesting that the Pip pathway as all other DRL pathways is indeed of the *trans*-AT PKS/NRPS type. Since no typical *trans-*AT is encoded in the pip BGC or anywhere in the *P. entomophila* genome, as confirmed by BLAST search, the primary metabolism malonyl-CoA:ACP-transacylase FabD (PSEEN_RS07520) might act as PipC AT. Indeed, enzyme kinetic analysis of the transacylation reaction confirmed that both FabD from *E. coli* and *P. entomophila* could catalyze the AT-mediated transacylation of the PipC-acyl carrier protein (ACP) (Fig. 5a).^[47,48]^

**Figure 5.**
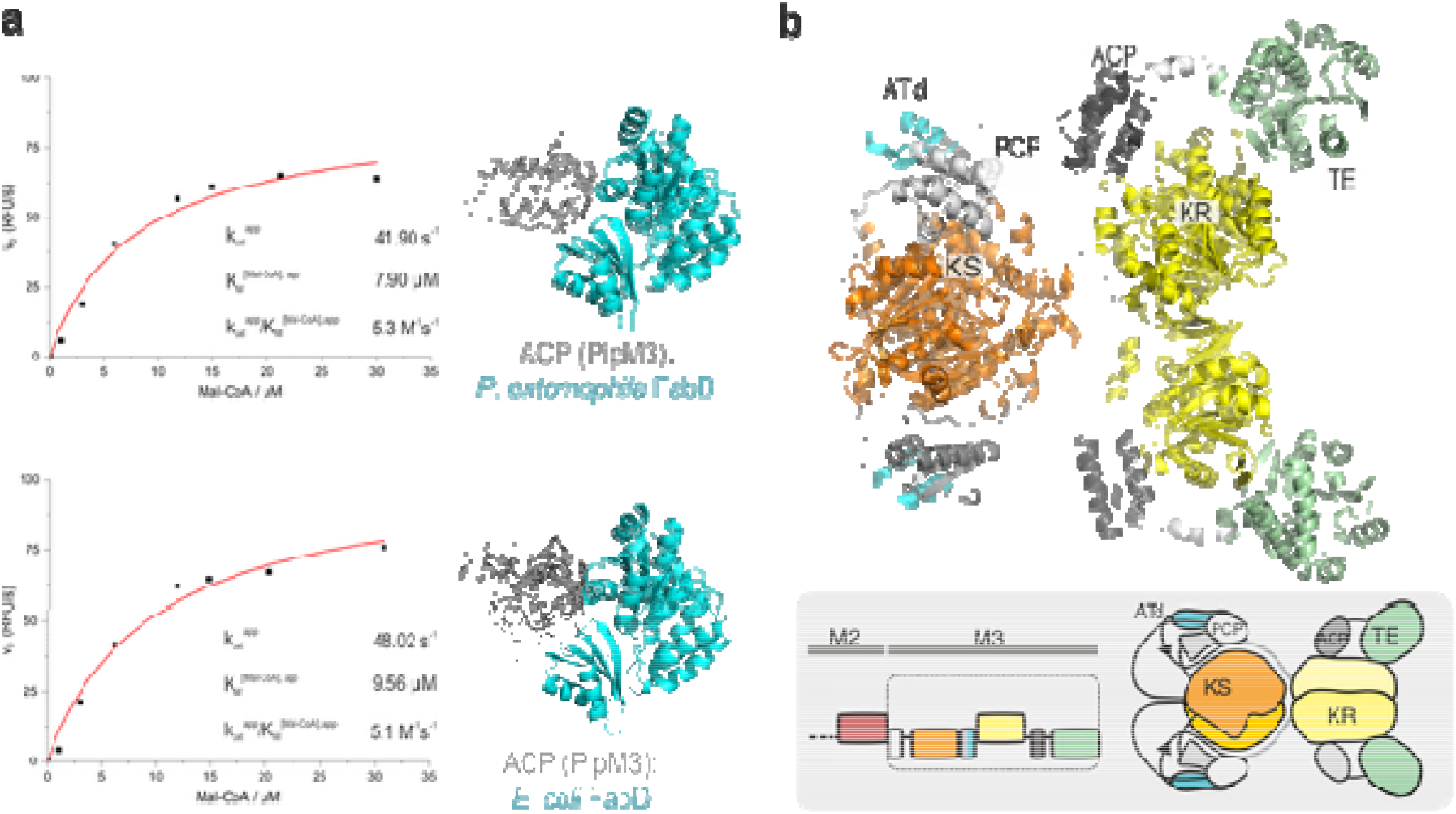
ACP-FabD interaction and structural modeling of PipC/Module-3. (**a**) Michaelis-Menten fits of transacylation titration curves of *P. entomophila* FabD (top panel) and *E. coli* FabD (bottom panel) with malonyl-CoA and ACP in fixed concentration, respectively. Measurements were performed on one biological sample in technical replication. Shown are the two predicted complexes, each represented by five structural models as ranked by AlphaFold based on internal confidence scores (ACP in grey in ribbon representation, FabDs in cyan). The higher positional variability of *P. entomophila* ACP in complex with *E. coli* FabD indicates uncertainty of the prediction. (**b**) PipC/Module-3, including the upstream PCP, modeled by AlphaFold.^[53]^ The N-terminal PCP is positioned at the KS domain, while the internal ACP interacts with the KR domain.^[54,55]^ Although PKS TE domains are typically dimeric, the TE domain is modeled here in its monomeric form interacting with the KR. The inset shows PipC/Module-3 in domain architecture with the upstream C-domain not included in the AlphaFold model in orange. Domain colors as introduced in Fig. 1.

### Biosynthesis of pseudotetraivprolide

Tetratide **3a** is generated from the PipAB NRPSs following standard NRPS biochemistry. It might be released from PipB by premature hydrolysis but normally it is connected to the PipC-bound ivprolide **1a** catalyzed by the PipB-TE (Fig. 6). Similar ester-forming TE domains are also found in other peptide NPs like the Gq protein inhibitor FR900359 from *Chromobacterium vaccini0i* ^[49]^and salinamide from *Streptomyces* CNB-091 ^[50]^. Release of the PipC-bound ester intermediate by the PipC-TE results in the first full-length product pseudotetraivprolide **4a**. Acetylation of the N-terminus by PipH to give **4c** is required also for PipE activity (Fig. S1.13) introducing the hydroxy group at the C-terminal proline ring leading to **4d**. At this stage, transesterification between the two hydroxy groups might lead to two possible variants (Fig. S1.14) having almost identical MS^2^ data (Fig. S2.7). **4d** is finally acetylated by the PipDFG complex to form **4e**.

**Figure 6.**
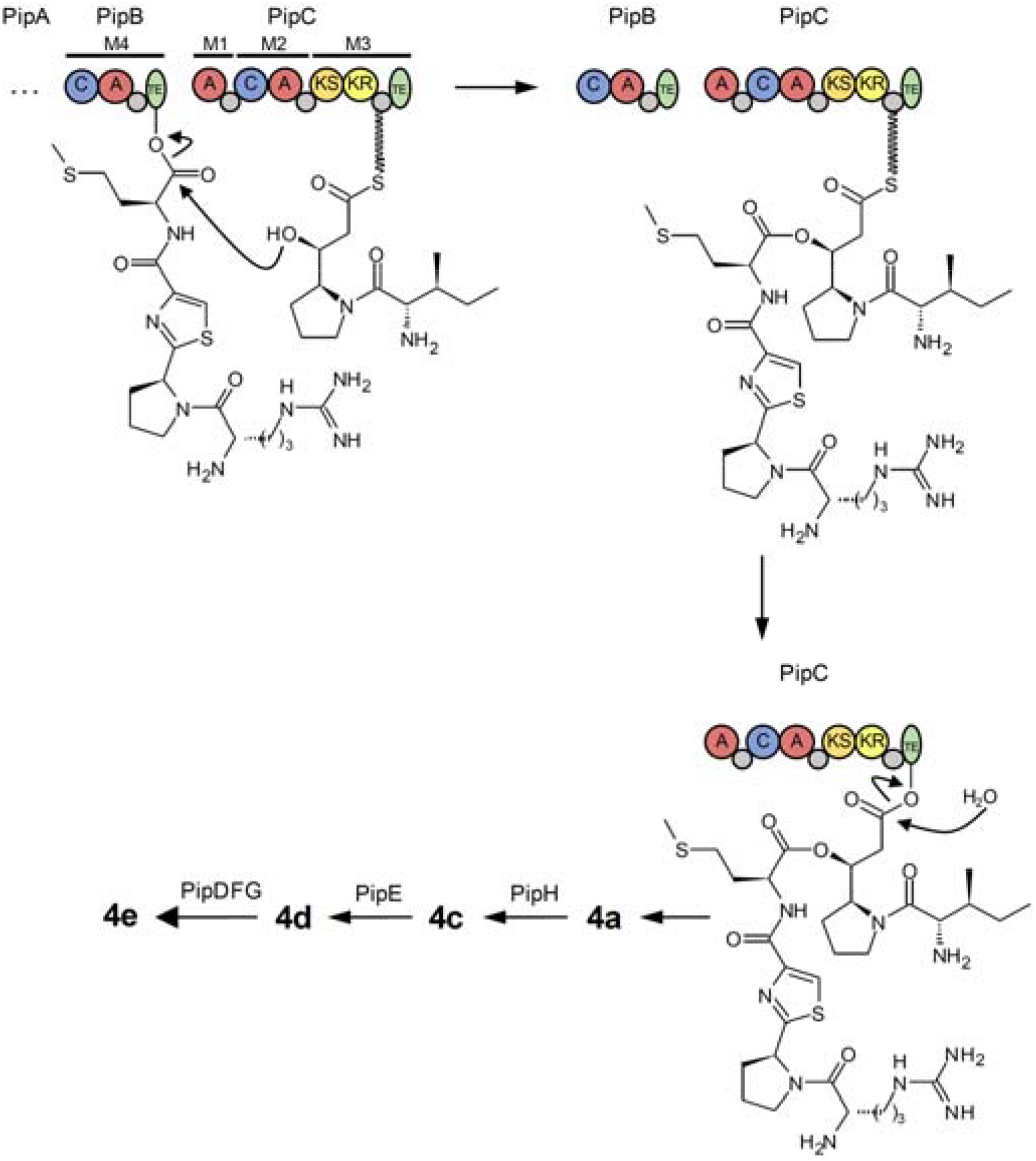
Proposed timing of the late-stage biosynthesis of pseudotetraivprolide focusing on the TEs from PipB and PipC, as well as PipDEFGH. For domain explanations of PipB and PipC see Fig. 1.

Complementation experiments of Δ*pipDEFGH* with plasmid encoded individual genes *pipD*-*pipH* showed that only PipH and not PipE acts on the core structure **4a** (Fig. S1.6), suggesting the N-terminal acetylation being a prerequisite for all further modifications. Strikingly, PipH is only found in strains having a free N-terminus and not in DRL BGCs that already have an acylated or carboxylated N-terminus as in detoxin or chitinimide, respectively (Fig. 1c), suggesting that a free N-terminus as in **4a** might prevent further modifications by PipEDFG.

A co-cultivation experiment with Δ*pipC*-pCEP*pipA* (only producing small amounts of **3b** but having functional PipDEFGH) and Δ*pipE*-pCEP*pipA* (mainly accumulating **4c** but not able to hydroxylate and acetylate it further) resulted in production of **4e**, suggesting that free **4c** can be secreted from the Δ*pipE* cell and enter the PipDEFG-containing cell for further modification (Fig. S1.15).

In closely related Pip BGCs from *Pseudomonas viridiflava* DSM 11124 and *Pseudomonas syringae* DSM 50272 *pipDFG* is missing (Fig. 1c, Fig S1.16). Also notably is a truncated Cstarter domain in PipA and a MbtH-encoding gene in *P. viridiflava*. As expected, extracts of respective promoter exchange mutants showed a similar compound profile compared *P. entomophila* except that **4e** was not detected and **4d**/**4d**□ were the final products in both strains (Fig. S1.17). Expression of *pipDFGH* from *P. entomophila* in *P. viridiflava* resulted in high production of **4e** (Fig. S1.18), raising the question about a not yet identified acetylation mechanism or a function of non-acetylated DRL products in this strain.

Since Δ*pipD*, Δ*pipF* and Δ*pipG* all showed a very similar production profile with accumulation of **4d** as final product (Fig. S1.5 & S1.6), all three enzymes were suggested to be involved in the terminal acetylation. Since no obvious homology to acetyltransferases were observed for any of the three proteins from BLAST-P search, we performed CLEAN^[51]^ (Table S4.1) and structure-based FoldSeek^[52]^ searches (Table S42 & S4.3) suggesting that PipF and PipG might indeed be hydrolases and PipD shows some similarity to an endopeptidase (see *Supplementary Information SI-4: Prediction of structure and function of PipD, PipF and PipG*). For PipF and PipG active sites for such hydrolytic activities could be suggested based on structural alignment with known enzyme active sites (Fig. 7a & b, Fig. S4.1 & S4.2) and a trimeric complex of PipDFG could also be proposed based on an Alphafold model, which additionally seems to be very stable as indicative from MD simulations over 100 ns (Fig. 7c, Fig. S4.3). The surface accessible catalytic triad in PipF might be Ser-18, His-211 and Asp-208 while in PipG the most likely surface accessible catalytic triad might be Cys-47, His-51 and Glu-296, while Ser-46 or Thr-238 as other candidate residues might not be equally surface accessible (Fig. S4.2). In order to test the role of these amino acids, Ser18Ala, His211Ala and Asp208Ala variants of PipG and Thr238Ala, Cys47Ala and His51Ala variants of PipF were generated and compared to the parent variants of both enzymes in vivo, confirming the importance of Ser-118 and His-211 in PipF and an unexpected role for Cys-47 in PipG (Fig. 7d). The large distance between the identified important residues in PipF and PipG suggests that PipDFG might either occur at a higher oligomerization state (Fig. S1.19) or that Cys-47 has another function, which cannot be clarified without a X-ray structure of the complex ideally with **4d** as the substrate for the acetylation.

**Figure 7.**
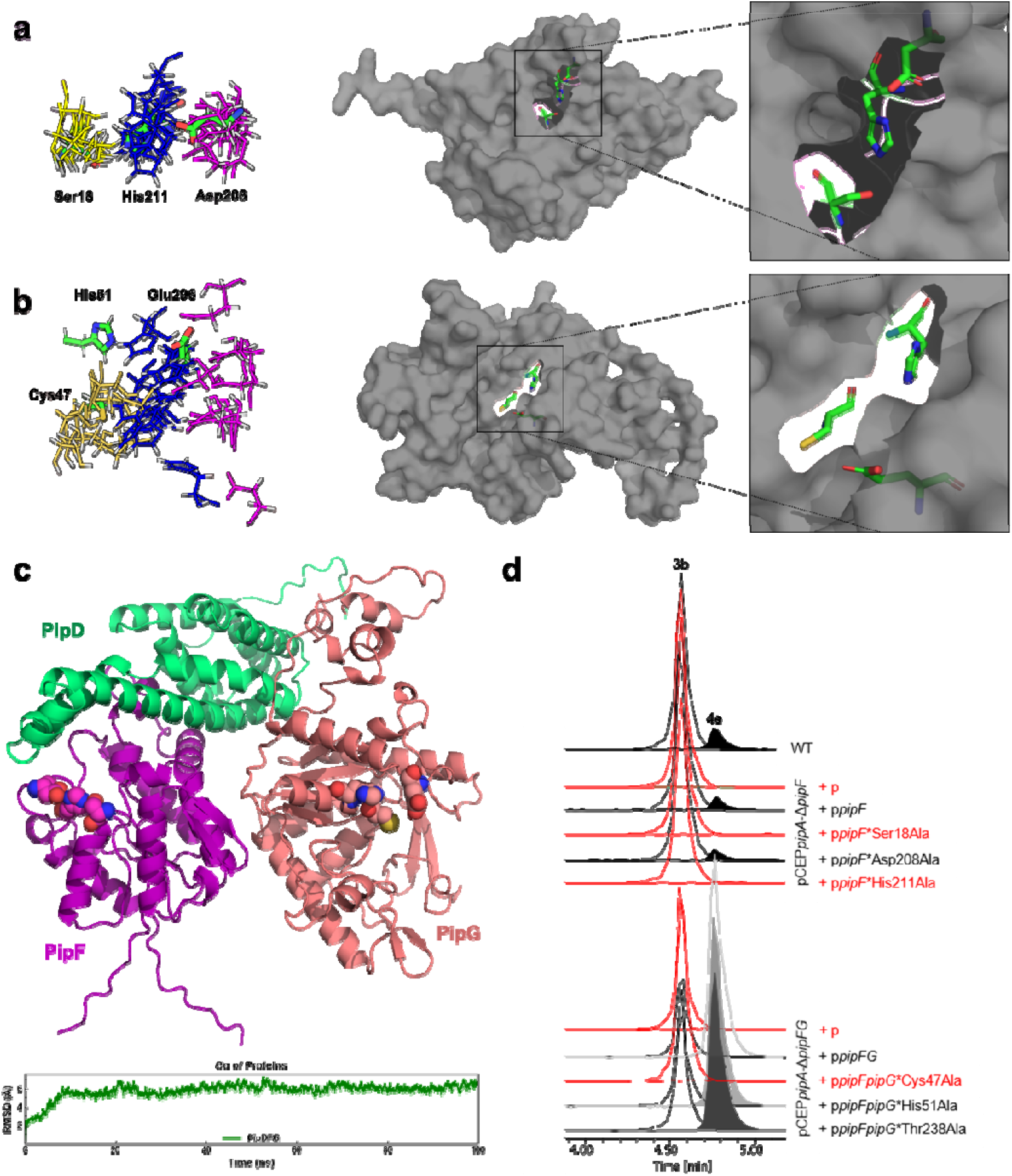
Proposed active sites of PipF and PipG and structure of the PipDFG complex. Proposed active sites of PipF (**a**) and PipG (**b**) are aligned with the active sites of enzymes identified in the FoldSeek results from the PDB100 dataset (left). The yellow, blue, and purple lines represent the residues corresponding to the nucleophilic, basic, and acidic components of the catalytic triad, respectively. Green sticks indicate the catalytic triad of PipF and PipG, respectively. Position and surface exposure of the catalytic triads in PipF and PipG is shown at the right. (**c**) Top: Trimeric complexes of PipD, PipF and PipG as modelled by AlphaFold. The catalytic triads of PipF and PipG are shown as spheres. Bottom: RMSD of Cα in trimeric complexes. (**d**) Production of **4e** in variants of PipF and PipG with specific amino acid exchanges confirming the importance of Ser18 and His211 in PipF and Cys47 in PipG, respectively. Lack of **4e** production is shown in red.

## Conclusions

We have solved the structure and biosynthesis of the pseudotetraivprolide, novel members of the widespread DRL family of NPs found in *Pseudomonas*. Through mutant construction and complementation, heterologous expression and in vitro experiments, we could suggest functions for all enzymes involved in the biosynthesis of this widespread NP class for the first time.

Although especially the identified timing of the individual biosynthetic steps might differ between organisms^[24,26,27]^, our work offers the possibility for a detailed comparison of DRL pathways in different organisms, which might point to different evolutionary adaptations. We could show for the first time that DRL pathways are dependent on FabD, indicating an unexpected link between primary and secondary metabolism. First evidence for a PipDFG complex responsible for the final-step acetylation as provided in our work requires additional structural and biochemical analysis in order to understand the function of all three proteins in this rather simple enzymatic modification, which might led to O-(**4e**) but also N-acetylation (**4h**, Fig. 2.16). While we could also confirm the anti-antibiotic activity specific for pseudotetraivprolide (Fig. S1.1), an unsolved ecological question is why DRL producers actually produce them to protect other bacteria like *B. cereus* against blasticidin S and not themselves against their own or foreign antibiotics as resistance mechanism.

## Author contributions

E.B., J.B. and P.H. constructed all strains, which were analyzed by E.B.. K.B. and U.K. planned and conducted the chemical synthesis. Y.N.S. isolated **1a** and **4c** and elucidated the structure also of **4e** by NMR together with Y.M.S.. S.R., Z.C. and M.G. performed the enzymatic analysis of the transacylation reaction, the structural modeling of PipC and the bioinformatics characterization of PipDFG. E.B. and H.B.B. wrote the paper with input from all authors.

## Acknowledgements

Work in the Bode lab was supported by an ERC Advanced Grant (835108) and the Max Planck Society. We are grateful for Prof. Dr. Graumann and his group providing us with *B. cereus* and access to his S2 laboratory, and to Christian Schelhas for initial protein modeling work.

## Data availability statement

All data and materials can be found within the manuscript, supporting information or can be requested from the corresponding author. There are three files of Supplementary Information: SI-1 (material & methods, Supplementary Tables S1.1-S1.8, Supplementary Figures S1.1-S1.19), SI-2 (compound identification & structure elucidation of all identified derivatives and NMR data of **1a, 4c** and **4e**; Supplementary Figures S2.1-S2.32), SI-3 (chemical synthesis of **4c & 5c**) and SI-4 (PipDFG structure/function prediction; Supplementary Tables S4.1-S4.3, Supplementary Figures S4.1-S4.3 and supplementary notes for structure/function prediction).

