## Supplementary Information SI-1 for "Identification of pseudotetraivprolide from *Pseudomonas entomophila* give novel insights into the biosynthesis of detoxin/rimosamide-like anti-antibiotics"

**Supporting Information SI-1: Microbiology**

**Materials & Methods**

**Cultivation conditions**

*E. coli* S17-1 λpir and *E. coli* ST18 cells, listed in Table S1 were cultivated on LB agar plates or in LB medium (10 g/L tryptone, 5 g/L yeast extract, 5 g/L NaCl, for solid plates 1,5 % (*w/v*) agar-agar was added to the medium) shaking at 200 rpm at 37°. For *E. coli* ST18 cells the LB media were supplemented with aminolaevulinic acid (ALA) 50 µg/mL. If needed, kanamycin was added to the media in a final concentration of 50 µg/mL, chloramphenicol, 34 µg/mL, gentamycin in a final concentration of 10 µg/mL.

*P. entomophila* was grown on LB agar plates or LB medium at 28-30 °C. To prevent the cells from swarming, the agar-agar concentration in solid media was increased up to 2.5 -3 % (*w/v*). When needed, antibiotics were added to the medium in appropriate concentrations: kanamycin 50 µg/mL, gentamicin 75 µg/mL (*P. entomophila*), 5 µg/mL (*P. viridiflava*) and 2 µg/mL (*P. syringae*).

**Production cultures**

For small-scale production cultures of *P. entomophila* wt and mutants, XPP medium was used as described[1]. In general, the respective cells were first cultivated overnight in 5 ml LB medium at 28 °C, shaking at 200 rpm. For analytical samples 5 ml of XPP medium were inoculated with 50 µl of the overnight culture. For induction of the $P_{BAD}$ promoter L-arabinose was added to final concentration of 0.2 % (*w/v*). For *P. syringae* and *P. viridiflava* L-arabinose was added to a final concentration of 2 % (*w/v*). The cultures were grown for 48 to 72 h at 28 °C shaking at 200 rpm.

**DNA isolation**

According to the manufacturers protocols genomic DNA was purified using either Monarch® Genomic DNA Purification Kit (NEB) or Gentra Puregene Yeast/Bact. Kit B (Qiagen).

Plasmid DNA was purified with Monarch® Plasmid Miniprep or with the Invisorb® Plasmid Spin Mini Two Kit (Stratec).

**Polymerase chain reaction (PCR)**

For generation of DNA fragments for the promoter exchange approach or complementation assays in all *Pseudomonas* strains Q5® High Fidelity DNA polymerase (New England Biolabs) was used according the manufacturer instructions. 1 µL of gDNA was added to 25 µL PCR reaction mix. The addition of 0.5 µL $MgCl_2$ [50 mM] and 3% DMSO enhanced the yield of the desired fragment. PCR was performed with Lab Cycler Gradient (Sensoquest GmbH) or peqSTAR 96X Universal (VWR Peqlab) thermocyclers. Promoter exchange mutants were verified by colony PCR using Phire® Green Hot Start II Polymerase (Thermo Scientific). In a first step cells were lysed in 25 QuickExtract™ DNA Extraction Solution (Lucigen). 1 µL of the lysate was added to the PCR reaction mix, 4% DMSO and 0.5 µL and $MgCl_2$ [50 mM] were added to increase the final yield. In all PCR cycles the initial denaturation at 98 °C was performed for two minutes and further conducted as described in the users guidelines. *E. coli* S17-1_pCEP promoter exchange constructs were verified by colony PCR using BioMix™ Red (meridian BIOSCIENCE™) following the manufacturers instructions. In a first step, cell material was resuspended in 25-30 µL 0.02 M NaOH, and lysed at 99 °C for several minutes. An aliquot of 1 µL of the lysate was applied in the PCR reaction mix. All primers are listed in Tables S1.4-S1.9.

**Preparation of extracts**

If not described differently, the compounds of respective cultures were extracted by mixing 100 ml of the respective 48 or 72 h culture with 400 ml acetonitrile. The bacterial debris was pelleted via centrifugation for 30 min at full speed. An aliquot of the supernatant was transferred into a HPLC vial, and further subjected to LC-MS analysis.

**Heat map**

Strains for heat map analysis were cultivated in triplicates. From a 5 mL LB over-night culture 5 mL XPP medium was inoculated at 28 °C, shaking at 200 rpm 72 h. EIC were determined for all derivatives and mean values were calculated.

**Assembly of pCEP and pEB17 constructs and conjugation**

Construction of plasmids for performing promoter exchanges or gene deletions were performed as described[1,2] based on plasmids listed in Table S1.5. The used vectors pCEP and pEB17 (Table S1.3) can only be propagated in *E. coli* ST18 and *E. coli* S17-1λpir cells and not in *Pseudomonas* cells.

Construction of pCEP_*xyz* (listed in Table S1.5) for promoter exchange: Briefly, the first 300-800 bp of the gene of interest were amplified via PCR with corresponding oligos containing overlapping regions to the vector backbone listed in Table 1.5 using the Q5 Polymerase from NEB following the manufacturers protocol for DNA with high GC content. The protocol was further optimized by the addition of up to 3 % DMSO and 2 % $MgCl_2$ [50 mM] in 25 µl PCR reaction mixture. The resulting fragment was separated in an agarose gel. The fragment was cut out from the gel and extracted using Qiagen MinElute® Gel Extraction Kit or Wizard SV Gel and PCR Clean-Up System (Promega) following the manufacturers instruction. The purified fragment was assembled with PstI and BglII linearized pCEP-km vector backbone by NEBuilder® Hifi DNA Assembly Mix (New England Biolabs). The assembled pCEP-*xyz* construct was transformed either in *E. coli* ST18 or into *E. coli* S17-1 λpir respectively.

The construction of pEB17-Δ*xyz* plasmids for deletion followed the described procedure[1]. Briefly, 800-1500 bp upstream and downstream of the target gene(s) with overlapping regions were amplified with oligos listed in Table 1.5. The pEB17 vector backbone was generated via restriction with enzymes PstI and BglII. The PCR fragments with overlapping 25-30 bp to each other and the vector backbone were subsequently ligated by NEBuilder® HiFi DNA Assembly Master Mix following the manufacturers protocol. They were further transformed into *E. coli* ST18 via electroporation.

*E. coli* clones carrying the ligated pCEP-*xyz* or pEB17-Δ-*xyz* construct were verified by colony PCR using oligo VpCEP-fw and VpCEP-rv for pCEP and oligo VpDS132-fw and pCEP-rv for pEB17 constructs, respectively. All primers used for verification are listed in Table S1.4.

The pCEP-gene xyz or pEB17-Δgene-xyz plasmids (all vectors used in this study are listet in Table 1.5) were transformed into *P. entomophila* via conjugation adapted from[3]. The respective *Pseudomonas* strain was cultivated in 5 mL LB medium, and the selected *E. coli* mutant was cultivated in 5 ml LB medium over night with antibiotics and ALA added for ST18 cells. The next day, the *E. coli* donor cells were washed twice in LB medium to eliminate ALA or antibiotics. The *E. coli* donor cells and the *Pseudomonas* acceptor cells were adjusted to an $OD_{600}$ of 6.0. 1 ml donor and 100 µl acceptor cells were mixed and pelleted by centrifugation for 1 min at 21,000 *g*. The cells were dissolved in 50 µl LB medium which were further spotted in one drop on an LB agar plate. The inoculated plate was incubated at 30 °C for 5-6 hours to enable mating of the cells and transfer of the plasmid via rolling circle mechanism. The cell plaque was scraped from the plate using a sterile inoculation loop and dissolved in 500 µl LB medium. Different dilutions from 1:20 – 1:100 were generated. 50 µl of the diluted cell suspension was plated on LB medium containing kanamycin as selective antibiotic and additional ampicillin in a final concentration of 100 µg/ml if *E. coli* S17-1 λpir cells were used. The inoculated plates were incubated for 24 to 48 hours at 30 °C. The obtained colonies for promoter exchange mutants were verified by colony PCR using VpCEP-fw and an individual verification primer respectively (Table S1.4).

**Generation of deletion strains**

Single colonies of the generated insertion mutants of *Pseudomonas* pEB17-Δ*xyz* were streaked onto no salt LB agar (NSLB) containing 15 % sucrose[1,4]. The addition of 3 % agar-agar (w/v) was necessary to prevent cells from swarming. The plates were incubated at 18 °C for 24-72 h to enable deletion of the vector backbone via a second homologous recombination step and the activity of SacB, which is encoded on the pEB17 vector. The obtained colonies were plated on LB agar and in parallel on LB agar containing kanamycin. Clones growing on LB agar only were verified by colony PCR using individual verification primers listed in Table S1.5.

**Anti-antibiotic activity**

*Preparation of extracts*

EB7001 (WT), EB862 (ΔPELP4_pCEP*pipA*), EB849 (ΔPELP4), listed in Table S1.2, were cultivated for 24 h at 28 °C, shaking at 200 rpm in 5 mL LB with 50 µg/mL kanamycin, if needed. To set up a production culture, 100 mL XPPM, containing 2 % XAD adsorber resin, were inoculated with 1 mL from this pre-culture. Before inoculation, the cells were washed twice to eliminate kanamycin. EB862 was induced with 0.2 % L-Arabinose. All cultures were incubated for 72 h at 28 °C, shaking at 200 rpm. The XAD resin was then harvested and extracted with 300 mL methanol. The solvent was evaporated to obtain an oily extract. Its concentration was adjusted with methanol to 20 mg/mL. Cellulose discs were prepared with 10 µL blasticidine-S [1 mg/mL] dissolved in water and the addition of different concentrations of extract from 862, 7001 and 849.

*Bioactivity assay*

The screen for biologic activity was performed as described previously [5]. *Bacillus cereus* ATTC14579 was cultivated in 10 mL LB medium 24 h at 30 °C, shaking at 200 rpm. Subsequently 30 µL of the *Bacillus* culture were spread on LB agar plates. Prepared discs were put onto the fresh inoculated plates. The cells were incubated for 24 h at 30 °C.

**Cloning of pSEVA constructs**

Complementation of deleted genes was carried out by cloning the respective gene into free replicating conjugatable plasmid pSEVA621-Gm [6]. The plasmid backbone was generated via PCR with primers AR-900-fw and AR-970-rv (plasmid and primers are listed in table 1.3 and 1.4.). Respective genes were amplified by PCR as described above. The DNA fragments were assembled with PCR generated pSEVA backbone by NEBuilder® HiFi DNA Assembly Master Mix. The assembled plasmid was transformed into *E. coli* ST18 cells via heat shock or electroporation. The constructs were verified by colony PCR with primers V_pCEP_fw and VPEB-796-pSEVA-rv (listed in Table SX). For complementation of the respective deletion mutant the plasmid was transformed via conjugation. For each deletion mutant an empty plasmid control was generated, by transforming empty pSEVA621-GM into the respective strain. All primers used for generating the pSEVA constructs and their respective strains are listed in Table 1.8,

**Cloning of pACYC and pCOLA constructs and heterologous expression**

For cloning of *pipC*, *pipCD* and *fabD* were amplified with primers listed in Table 1.6. The amplified genes were ligated into PCR generated pACYC_ara or pCOLA_ara backbone respectively, via NEBuilder® Hifi DNA Assembly Mix (New England Biolabs). Assembled constructs were transformed into *E. coli* DH10B::*mtaA* via electroporation. All generated mutants are listed in Table 1.6. For heterologous expression, cells were cultivated overnight in 5 mL LB medium which the required antibiotics. The next day, 5 mL LB medium with antibiotics, were inoculated with 50 µL from the respective overnight culture. The production cultures were induced with 0.2%

L-arabinose (*w/v*) following cultivation for 24 h at 22 °C and shaking at 200 rpm. Culture extraction was conducted as described above.

**Cross feeding experiment via co-cultivation**

*P. entomophila*-pCEP*pipA*, *P. entomophila* Δ*pipC*-pCEP*pipA* and *P. entomophila* Δ*pipE*-CEP*pipA* were cultivated in 5 mL LB medium with 50 µg/mL kanamycin at 28 °C for 24 h (shaking at 200 rpm). For co-cultivations 5 mL of fresh LB medium were inoculated with 50 µL *P. entomophila*Δ*pipC*-pCEP*pipA* culture together with 50 µL *P. entomophila* Δ*pipE*-CEP*pipA* culture. As controls, each strain was also separately cultivated. All cultures were induced with 0.2 % L-arabinose (*w/v*). Cultivation was performed for 72 h at 28 °C (shaking at 200 rpm).

**Inserting point mutations**

Point mutation of the TE domain in *pipB* was introduced by mutating Ser into Ala in the catalytic center by converting the codon TCG>GCG on overlapping primers listed in Table S1.5. Point mutations in the amino acids of the catalytic triad of PipF and PipG were introduced via PCR with designed primers listed in Table 1.7 by exchanging the respective codon into a non-sense GCG codon for the incorporation of alanine. p*pipF* and p*pipFG* listed in Table 1.7 were used as template for PCR. Generated fragments were further ligated using KLD Enzyme Mix (NEB). The obtained plasmids were verified by sequencing and transformed via conjugation into Δ*pipF*-pCEP*pipA* (2362) and Δ*pipFG*-pCEP*pipA* (2307) respectively. Productions cultures of respective mutants were analyzed as described above.

**Supplementary Tables**

**Table S1.1.** Genes and putative functions in the *pip* gene cluster and other genes investigated in this study. Asterisk indicate adaptation suggested by antiSMASH.

| name | locus tag | lengh [bp] | no. of amino acids | protein id | function |
|---|---|---|---|---|---|
| *pipA* | PSEEN_RS12600 | 8973 | 2990 | WP_011533898.1 | NRPS |
| *pipB* | PSEEN_RS12605 | 4116 | 1371 | WP_011533899.1 | NRPS |
| *pipC* | PSEEN_RS12610 | 9036 | 3011 | WP_011533900.1 | NRPS-PKS |
| *pipD* | PSEEN_RS12615 | 675* | 224 | WP_158020252.1 | hypothetical protein |
| *pipE* | PSEEN_RS12620 | 870 | 289 | WP_011533902.1 | TauD/TfdA dioxygenase family protein |
| *pipF* | PSEEN_RS12625 | 723* | 240 | WP_158020253.1 | hypothetical protein |
| *pipG* | PSEEN_RS12630 | 1035 | 344 | WP_011533904.1 | GSCFA-domain containing protein |
| *pipH* | PSEEN_RS12635 | 519 | 172 | WP_011533905.1 | GNAT family N-acetyltransferase |
| *fabD* | PSEEN_RS07520 | 939 | 312 | WP_011532885.1 | [acyl-carrier-protein] S-malonyltransferase |
| *hfq* | PSEEN_RS22860 | 261 | 86 | WP_011535948.1 | RNA-binding protein Hfq |
| *pipA* | GHKPGMKM_08200 | 9066 | 3021 | - | NRPS |
| *pipA* | LMDDJHCF_09560 | 9711 | 3236 | - | NRPS |

**Table S1.2.** Strains used in this work.

| Name | Strain | Reference |
|---|---|---|
| HBLC 395/EB 7001 | *Pseudomonas entomophila* L48 DSM 28517 | [7] |
| PG 4253 | *Bacillus cereus* ATCC14579 | [5] |
| HBEC 089 | *E. coli* S17-1λpir (Tp, Smr, *recA*, *thi*, hsdRM+RP4::2-Tc::Mu::Km, Tn7, λpir phage lysogen) | Invitrogen |
| HBEC 118 | *E. coli* ST18 (*E. coli* S17-1λpir Δ*hemA*) | [3] |
| HBEC 004 | *E. coli* DH10B (F-mcrA, Δ(mrr-hsdRMSmcrBC) Φ80lacZΔM15, ΔlacX74, *recA1*, endA1, araD139, Δ(ara leu)7697 galU, galK, rpsL, nupG, λ-) | [8] |
| HBEC 029 | DH10B *entD*::*mtaA* | [9] |
| HBLC 386 | *Pseudomonas syringae* DSM 50274 | DSMZ |
| HBLC 388 | *Pseudomonas viridiflava* DSM 11124 | DSMZ |
| EB 849 | *P. entomophila*_Δ*pvsA*Δ*eltA*ΔRS12890Δ*psmDFG*ΔRS10025-10010 | This work |
| EB 862 | *P. entomophila*_Δ*pvsA*Δ*eltA*ΔRS12890Δ*psmDFG*ΔRS10025-10010_pCEP*pipA* | This work |
| 2506 | *P. viridiflava*-pCEP*pipA* (2369)+p*pipD-H* (pEB141) | This work |
| EB 7050 | *P. viridiflava*-pCEP*pipA*+pSEVA621 (empty vector) | This work |
| 2484 | *P. syringae*-pCEP*pipA*(2367)+p*pipD-H* (pEB141) | This work |
| 2486 | *P. syringae*-pCEP*pipA*+pSEVA621 (empty vector) | This work |
| EB 7018 | *P. entomophila* Δ*pipD*Δ*pipH*-pCEP*pipA* | This work |
| EB 7045 | *P. entomophila*_Δ*pipD*Δ*pipH*-pCEP*pipA* + pSEVA621 (empty vector) | This work |
| EB 7046 | *P. entomophila*_Δ*pipD*Δ*pipH*-pCEP*pipA* + p*pipH* (pEB132) | This work |
| EB 7048 | *P. entomophila*_Δ*pipD*Δ*pipH-pCEPpipA* + p*pipD* (pEB136) | This work |
| 2512 | *P. entomophila* Δ*pipDEFGH*_pCEP*pipA* + pSEVA621-GM (empty vector) | This work |
| 2514 | *P. entomophila* Δ*pipDEFGH*_pCEP*pipA* + p*pipH* (pEB132) | This work |
| 2516 | *P. entomophila* Δ*pipDEFGH*_pCEP*pipA* + p*pipFGH* (pEB133) | This work |
| 2520 | *P. entomophila* Δ*pipDEFGH*_pCEP*pipA* +p*pipD* (pEB136) | This work |
| 2522 | *P. entomophila* Δ*pipDEFGH*_pCEP*pipA* + p*pipG* (pEB137) | This work |
| 2524 | *P. entomophila* Δ*pipDEFGH*_pCEP*pipA* + p*pipFG* (pEB138) | This work |
| 2528 | *P. entomophila* Δ*pipDEFGH*_pCEP*pipA* + p*pipF* (pEB140) | This work |
| 2530 | *P. entomophila* Δ*pipDEFGH*_pCEP*pipA* + p*pipE* (pEB143) | This work |
| 2532 | *P. entomophila* Δ*pipDEFGH*_pCEP*pipA* + p*pipDE* (pEB144) | This work |

**Table S1.3.** Plasmids used in this work.

| Name | Description | Reference |
|---|---|---|
| pEB17-Km | pDS132 based, R6K *ori*, $Km^R$, *ori*, *sacB* | [1,10] |
| pCEP-Km | pDS132 based, R6K *ori*, $Km^R$, $P_{BAD}$, *oriT* | [1,2,10] |
| pCOLA_ara/tacI-Km | ColA *ori*, $Km^R$, *araC*-$P_{BAD}$, *tacI* | [11] |
| pACYC_ara/tacI-Km | p15A *ori*, $Cm^R$, *araC*-$P_{BAD}$ *tacI* | [12] |
| pCOLA_ara/tacI-Gm | ColA *ori*, $Gm^R$, *araC*-$P_{BAD}$ *tacI* | [11] |
| pSEVA621-Gm | oriV, $Gm^R$, $P_{BAD}$, mNeonGreen, *oriT,* RiboJ | [6] |

**Table S1.4.** General primers used in this work.

| Name | Sequence | Purpose |
|---|---|---|
| V_pCEP_fw | GCTATGCCATAGCATTTTTATCCATAAG | verification primer for pSEVA and pCEP constructs |
| VPEB-796-pSEVA-rv | CATCGTTGCTGCTGCGTAAC | verification primer for pSEVA constructs |
| V_pDS132-rv | ACATGTGGAATTGTGAGCGG | verification primer for pEB17 and pCEP constructs |
| V_pDS132_fw | GATCGATCCTCTAGAGTCGACCT | verification primer for pEB17 constructs |
| PEB_1028-fw | CAGCTTAATTAACCTAGGCTGCTG | backbone amplification pACYCara or pCOLA_ara_tac |
| PEB_1029-rv | GGAATTCCTCCTGTTAGCCCAA | |
| AR_900-fw | GTCGTGACTGGGAAAACCCT | backbone amplification pSEVA621-Km |
| AR_970-rv | CTAGTATTTCCCCTCTTTCTCTAGT | |

**Table S1.5**. Primer for the construction of respective *E. coli* strains, carrying either pCEP constructs for promoter exchange of selected genes and respective *P. entomophila*-pCEP strains. *E. coli* strains with pEB17 constructs for gene deletion and the resulting *P. entomophila* deletion mutants with promoter exchange.

| Name | Sequence | *E. coli* strain | Purpose | *P. entomophila* strain |
|---|---|---|---|---|
| PEB_518-fw | TTTGGGCTAACAGGAGGCTAGCAT_ATGCCCGACACTTCCTCCTTG | EB535 pCEP*pipA* (pEB98) | Promoter exchange to PSEEN_RS12600 | pCEP*pipA* (JB32) |
| PEB_519-rv | TCTGCAGAGCTCGAGCATGCACAT_CTGCAACGGGTACAGCCGTGC | | | |
| VPEB_520-rv | GCTGAACACATCGAACGAG | | | |
| PEB_580-fw | TTTGGGCTAACAGGAGGCTAGCAT_ATGAGCGTGTTCCAGGAAAC | EB592 pCEP*pipC* (pEB97) | Promoter exchange to PSEEN_RS12610 | pCEP*pipC* (EB593) |
| PEB_581-rv | TCTGCAGAGCTCGAGCATGCACAT_TCGACAGGCAGACAAATCGAT | | | |
| VPEB_582-rv | CACCGTGATGAACAAGGTATCGAC | | | |
| JUB_1_fw | CCTCTAGAGTCGACCTGCAG_CGTGACAGCACCGGTCGAAG | JB29 pEB17_Δ*pipD* (pJuB-01) | Deletion of PSEEN_RS12615 | Δ*pip*D-pCEP*pipA* (JB55) |
| JUB_2_rv | CTGATGGGTGTTGGCTGGCCACGTG_CTGTGCATATTCACCGGCCATG | | | |
| JUB_3_fw | GTTCATGGCCGGTGAATATGCACAG_CACGTGGCCAGCCAACACC | | | |
| JUB_4_rv | TCCCGGGAGAGCTCAGATCT_GTCGAGCTGGTCGATGTGCG | | | |
| V_JUB_5_fw | AGTCGGTCTGTCGACACAGGAAG | | | |
| V_JUB_6_rv | GCGGCGCAGACACATTTGATCG | | | |
| JUB_7-fw | CCTCTAGAGTCGACCTGCAG_AAACCACGGTCATCAACAGTATCTGG | JB45 pEB17_*pipB*ΔTE (pJuB-02) | Deletion of the TE Domain in PSEEN_RS12605 (*pipB*) | Δ*pipB*-TE-pCEP*pipA* (JB59) |
| JUB_8-rv | ATTTCCCCTCCTGGGCATCGAAACGTTGCTGGCCGTCCCCTGACCTGAA | | | |
| JUB_9-fw | ATCACCTTCAGGTCAGGGGACGGCCAGCAACGTTTCGATGCCCAGGAGG | | | |
| JUB_10-rv | TCCCGGGAGAGCTCAGATCT_AATAGGGTGGCACAGGGCGAAGG | | | |

| | | | | |
|---|---|---|---|---|
| V_JUB_11-fw | TACAGCCTCAAGGACGTATTCTGC | | | |
| V_JUB_12-rv | ATGCCGGTGTCCGGCAAGGGCTGG | | | |
| PEB_616-fw | CCTCTAGAGTCGACCTGCAG_GAGCTGACCCTCTACCAGATCTGG | JB20 pEB17_Δ*pipC* (pJuB-03) | Deletion of PSEEN_RS12610 | Δ*pipC*-pCEP*pipA* (JB31) |
| PEB_617-rv | CATATTCACCGGCCATGAACACCCTCCTCCACGGTTTCCTGGAACACGCTCAT | | | |
| PEB_618-fw | GGAAATGATGAGCGTGTTCCAGGAAACCGTGGAGGAGGGTGTTCATGGCCGG | | | |
| PEB_619-rv | TCCCGGGAGAGCTCAGATCT_GTCCCGAGAAACGCCTGCATC | | | |
| VPEB_620-fw | CTACCTGTGGCTCGACAGCCTG | | | |
| VPEB_621-rv | CCAGGGCATCTCGCCGTCAT | | | |
| VPEB_582-rv | GTCGATACCTTGTTCATCACGGTG | | | |
| PEB_622-fw | CCTCTAGAGTCGACCTGCAG_GTGAGTCGCCCAACAGCAGGAAG | JB24 pEB17_Δ*pipH* (pJuB-04) | Deletion PSEEN_RS12635 | Δ*pipH*-pCEP*pipA* (JB57) |
| PEB_623-rv | CCTTGCCCTCCACTCTCGCTAGGAGCACCCTCGACCTGCACGGATAG | | | |
| PEB_624-fw | GTTCATGGGCTATCCGTGCAGGTCGAGGGTGCTCCTAGCGAGAGTGGAG | | | |
| PEB_625-rv | TCCCGGGAGAGCTCAGATCT_CAATGCTATGCAGCAAGACCAG | | | |
| VPEB_626-fw | CACCGCTGAACGACAGCTGC | | | |
| VPEB_627-rv | GAGCAGACTCACCTAACAATCGG | | | |
| VPEB_626.1-fw | GTACAGCCGCTCGAGTTCG | | | |
| PEB_610-fw | CCTCTAGAGTCGACCTGCAG_GATCGCCGAGGCGATCAATAGC | JB22 pEB17_Δ*pipE* (pJuB-05) | Deletion of PSEEN_RS12620 | Δ*pipE*-pCEP*pipA* (JB56) |
| PEB_611-rv | CTGCATACCCTGAAACAAGCACCTGCGGCCTTCAGTGCCTGTTGATCCAT | | | |
| PEB_612-fw | GTATGGATCAACAGGCACTGAAGGCC_GCAGGTGCTTGTTTCAGGGT | | | |
| PEB_613-rv | TCCCGGGAGAGCTCAGATCT_GGAGGCGTGGCCTTCGTGATG | | | |
| VPEB_614-fw | CATGTGCCTGGCGTACAGGC | | | |

| | | | | |
|---|---|---|---|---|
| VPEB_615-rv | CTACTTCCCGTCCTACGAAATCATC | | | |
| JUB_13-fw | ATCGATCCTCTAGAGTCGACCTGCAG-TTTCATCGACAGCCCGTTCGTGC | JB65 pEB17_*pipB*-S(72)A (pJuB-06) | Point mutation S(72)A in the TE domain of PSEEN_RS12605 (*pipB*) | Δ*pipB*-TE*-pCEP*pipA* (JB67) |
| JUB_14-rv | GCCGAACGCCAGCCCGGTGAG | | | |
| JUB_15-fw | CTCACCGGGCTGGCGTTCGGC | | | |
| JUB_16-rv | TGGAATTCCCGGGAGAGCTCAGATCT-AAATCGATCGCCCTGCTGGATATCGAT | | | |
| JUB_14.1_rv | GTAGGCCACCAGGCCGCCGAACGCCAGCCCGGT GAGCACCAG | | | |
| JUB_15.1_fw | CGGCCACTGGTGCTCACCGGGCTGGCGTTCGGC GGCCTGGTGGC | | | |
| Seq_JUB_17 _fw | TCTGCTACCTGTCGCTG | | | |
| Seq_JUB_18 _rv | AGCTTGTCGCGGAACTC | | | |
| PEB-1002-fw | GATCCTCTAGAGTCGACCTGCAG_CGAGAGTCAG ATGCTGTTGAAC | 2318 pEB17_Δ*pip*G (pPH61) | Deletion of PSEEN_RS12630 (*pipG*) | Δ*pipG*-pCEP*pipA* (2334) |
| PEB-1003-rv | CATCTACCGCCTCGACCTGCACGGATAGCTTCCTG CTGTTGGGCGAC | | | |
| PEB-1004-fw | CTAGGTGTGAGTCGCCCAACAGCAGGAAGCTATC CGTGCAGGTCGAG | | | |
| PEB-1005-rv | GTGGAATTCCCGGGAGAGCTCAGATCT_CCAGTGT TCACATCGAGCTC | | | |
| VPEB-1006-fw | CACCTCGCATATCGTGCAG | | | |
| VPEB-1007-rv | GTTGTAGGCGTACGCCAAG | | | |
| PEB-1008-fw | GATCCTCTAGAGTCGACCTGCAG_GGATCAACAG GCACTGAAGG | 2319 pEB17_Δ*pipF* (pPH62) | Deletion of PSEEN_RS12625 (*pipF*) | Δ*pipF*-pCEP*pipA* (2362) |
| PEB-1009-rv | GGGATTTCGGGGTGGACTTCTTCACCCTGGACCG CATCGCGGATGAATC | | | |
| PEB-1010-fw | GTAGGAGCGGATTCATCCGCGATGCGGTCCAGGG TGAAGAAGTCCAC | | | |
| PEB-1011-rv | GGATGAACTGGCTCGATG | | | |

| | | | | |
|---|---|---|---|---|
| VPEB-1012-fw | GTTGCAACACGTGGCCAG | | | |
| VPEB-1013-rv | CAAGGACGCCATCGTCAC | | | |
| PEB_963-rv | CATCTACCGCCTCGACCTGCACGGATAGGACCGC ATCGCGGATGAATC | 2305 pEB17_Δ*pipFG* (pEB138) | Deletion of PSEEN_RS12625-12630 (*pipFG*) | Δ*pipFG*-pCEP*pipA* (2311) |
| PEB_964-fw | GTAGGAGCGGATTCATCCGCGATGCGGTCCTATC CGTGCAGGTCGAG | | | |
| PEB_965-rv | GTGGAATTCCCGGGAGAGCTCAGATCT_GTTGTGT TGTAGGCGTACGC | | | |
| VPEB-966-fw | GGCCATCGAACGTATTGCC | | | |
| VPEB-967-rv | CAGCTCACCCAATCCGATGG | | | |
| PEB-1008-fw | GATCCTCTAGAGTCGACCTGCAG_GGATCAACAG GCACTGAAGG | 2305 pEB17_Δ*pipFGH* (pEB142) | Deletion of PSEEN_RS12625-12635 (*pipFGH*) | Δ*pipFGH*-pCEP*pipA* (2471) |
| PEB-1087-rv | CTTGCCCTCCACTCTCGCTAGGAGCACCGGGACC GCATCGCGGATGAATC | | | |
| PEB-1088-fw | GTAGGAGCGGATTCATCCGCGATGCGGTCCCGGT GCTCCTAGCGAGAG | | | |
| PEB-625-rv | GTGGAATTCCCGGGAGAGCTCAGATCT_CAATGCT ATGCAGCAAGACCAG | | | |
| VPEB-1012-fw | GATCCTCTAGAGTCGACCTGCAG_CTCGATCATAA GACACTGCG | | | |
| VPEB-627-rv | GAGCAGACTCACCTAACAATCGG | | | |
| P-JuB-01-fw | CCTCTAGAGTCGACCTGCAG_CGTGACAGCACCG GTCGAAG | 2469 pEB17_Δ*pipDEFGH* (pEB134) | Deletion of PSEEN_RS12615-12635 (*pipDEFGH*) | Δpip*DEFGH*-pCEP*pipA* (2470) |
| PEB-1081-rv | GCACCGGATGCTGGAGCTGCACACCGTGCAACAG CCGCCAGTAC | | | |
| PEB-1082-fw | CCAGGCGTACTGGCGGCTGTTGCACGGTGTGCAG CTCCAGCAT | | | |
| PEB-1005-rv | GTGGAATTCCCGGGAGAGCTCAGATCT_CCAGTGT TCACATCGAGCTC | | | |
| V-JuB-5-fw | AGTCGGTCTGTCGACACAGGAAG | | | |
| VPEB-1007-rv | GTTGTAGGCGTACGCCAAG | | | |

**Table S1.6.** Plasmids generated for heterologous expression of *pipC* and *fabD* in *E. coli* DH10B::*mtaA*.

| Name | Sequenz | Plasmid | Strain |
|---|---|---|---|
| PEB_1026-fw | CGTTTTTTTGGGCTAACAGGAGGAATTCC_ATGTCTGCATCCCTCGCATTC | pEB114 | 2341 *E. coli* DH10B::*mtaA*-pACYC_ara_*fabD* |
| PEB_1027-rv | GTGGCAGCAGCCTAGGTTAATTAAGCTG_GCAAGCGTCTCCAGATTCAG | | |
| PEB_1030-fw | CGTTTTTTTGGGCTAACAGGAGGAATTCC_ATGAGCGTGTTCCAGGAAAC | pEB115 | 2342 *E. coli* DH10B::*mtaA* pCOLA_ara_*pipC* |
| PEB_1031-rv | GTGGCAGCAGCCTAGGTTAATTAAGCTG_TCACCGGCCATGAACACC | | |
| PEB_1030-fw | CGTTTTTTTGGGCTAACAGGAGGAATTCC_ATGAGCGTGTTCCAGGAAAC | pEB118 | 2356 *E. coli* DH10B::*mtaA* pCOLA_ara_*pipCDE* |
| PEB_1033-rv | GTGGCAGCAGCCTAGGTTAATTAAGCTG_ACCCTGAAACAAGCACCTGC | | |

**Table S1.7.** Primers, plasmids and strains generated to introduce point mutations in the catalytic centers of PipF and PipG using p*pipF* (pEB140) and p*pipFG* (pEB138) as template.

| Name | Sequence | Plasmid | *E. coli* strain | *P. entomophila* strain |
|---|---|---|---|---|
| P-Ser18-fw | GCGCACCTAGGCGTCGTCGG | pEB146 | pEB146-pSEVA621-GM-*pipF* Ser18*-Ala (2559) | Δ*pipF*-pCEP*pipA* + p*pipF* Ser18* (2565) |
| P-Ser18-rv | GTCGCCCAACAGCAGGAAG | | | |
| P-Asp208-fw | GCGCCGTTGCATGGCAACCTC | pEB147 | pEB147-pSEVA621-GM-*pipF* Asp208*-Ala (2560) | Δ*pipF*-pCEP*pipA* + p*pipF* Asp208* (2567) |
| P-Asp208-rv | GTCCACCTCGCAGAACTCG | | | |
| P-His211-fw | GCGGGCAACCTCGCGTTTG | pEB148 | pEB148-pSEVA621-GM-*pipF* His211*-Ala (2561) | Δ*pipF*-pCEP*pipA* + p*pipF* His211* (2569) |
| P-His211-rv | CAACGGGTCGTCCACCTCG | | | |
| P-Cys47-fw | GCGTTCGCCCAGCACATC | pEB149 | pEB149-pSEVA621-GM-*pipFG* Cys47*-Ala (2562) | Δ*pipFG*-pCEP*pipA* + p*pipFG* Cys47* (2571) |
| P-Cys47-rv | GGACCCGGCGGTGACGAT | | | |
| P-His51-fw | GCGATCGGCCGGGCCCTG | pEB150 | pEB150-pSEVA621-GM-*pipFG* His51*-Ala (2563) | Δ*pipFG*-pCEP*pipA* + p*pipFG* His51* (2573) |
| P-His51-rv | CTGGGCGAAACAGGACCC | | | |
| P-Thr238-fw | GCGGCCAGCGCCACCGG | pEB151 | pEB151-pSEVA621-GM-*pipFG* Thr238*-Ala (2564) | Δ*pipFG*-pCEP*pipA* + p*pipFG* Thr238* (2575) |
| P-Thr238-rv | GAGGGGAACCGGCGAGAC | | | |

**Table S1.8** Primer and plasmids for generated *E. coli* strains and respective complemented deletion strains.

| Name | Sequence | Plasmid | *E. coli* strain | Strain |
|---|---|---|---|---|
| PEB_1048-fw | GTTTAATACTAGAGAAAGAGGGGAAATACTAG_GTGCCCGCGTGGCTGGGC | pEB136 p*pipD* | pSEVA-ara_RS12615 (p*pipD*) (2444) | Δ*pipD*-pCEP*pipA* + p*pipD* (2520) |
| PEB_1049-rv | GTCGCCAGGGTTTTCCCAGTCACGAC_CGTTCGATGGCCTTCAGTGCC | | | |
| PEB_1051-fw | GTTTAATACTAGAGAAAGAGGGGAAATACTAG_ATGGATCAACAGGCACTGAAGG | pEB143 p*pipE* | pSEVA-ara_RS12620 (p*pipE*) (2504) | Δ*pipE*-pCEP*pipA* + p*pipE* (2530) |
| PEB_1050-rv | GTCGCCAGGGTTTTCCCAGTCACGAC_CTCGATCTCTAGAGCGCTGCATA | | | |
| PEB_1066-fw | GTTTAATACTAGAGAAAGAGGGGAAATACTAG_GTGAAGACGCCATTCTCGATTAC | pEB140 p*pipF* | pSEVA-ara_RS12625 (p*pipF*) (2448) | Δ*pipF*-pCEP*pipA* + p*pipF* (2528) |
| PEB_1067-rv | GTCGCCAGGGTTTTCCCAGTCACGAC_TCATGGGCTCCGCAACGC | | | |
| PEB_1068-fw | GTTTAATACTAGAGAAAGAGGGGAAATACTAG_ATGAACCCCTACCAATACCTGC | pEB137 p*pipG* | pSEVA-ara_RS12625 (p*pipG*) (2445 ) | Δ*pipG*-pCEP*pipA* + p*pipG* (2522) |
| PEB_1069-rv | GTCGCCAGGGTTTTCCCAGTCACGAC_CAGCAGGAAGCTAGGCGTAG | | | |
| PEB_1044-fw | GTTTAATACTAGAGAAAGAGGGGAAATACTAG_ATGCTGGAGCTGCACACC | pEB132 p*pipH* | pSEVA-ara_RS12635 (p*pipH*) (2424) | Δ*pipH*-pCEP*pipA* + p*pipH* (2514) |
| PEB_1047-rv | GTCGCCAGGGTTTTCCCAGTCACGAC_GTATTGGTAGGGGTTCATGGG | | | |
| PEB_1064-fw | GTTTAATACTAGAGAAAGAGGGGAAATACTAG_ATGGCCGGTGAATATGCACAG | pEB144 p*pipDE* | pSEVA-ara_RS12615-RS12620 (p*pipDE*) (2505) | Δ*pipDE*-pCEP*pipA* + p*pipDE* (2532) |
| PEB_1050-rv | GTCGCCAGGGTTTTCCCAGTCACGAC_CTCGATCTCTAGAGCGCTGCATA | | | |
| PEB_1068-fw | GTTTAATACTAGAGAAAGAGGGGAAATACTAG_ATGAACCCCTACCAATACCTGC | pEB138 p*pipFG* | pSEVA-ara_RS12625-RS12630 (p*pipFG*) (2446) | Δ*pipFG*-pCEP*pipA* + p*pipFG* (2524) |
| PEB_1067-rv | GTCGCCAGGGTTTTCCCAGTCACGAC_TCATGGGCTCCGCAACGC | | | |
| PEB_1044-fw | GTTTAATACTAGAGAAAGAGGGGAAATACTAG_ATGCTGGAGCTGCACACC | pEB133 p*pipFGH* | pSEVA-ara_RS12625-RS12635 (p*pipFGH*) (2425) | Δ*pipFGH*-pCEP*pipA* + p*pipFGH* (2516) |
| PEB_1046-rv | GTCGCCAGGGTTTTCCCAGTCACGAC_GATTCATCCGCGATGCGGTC | | | |

| | | | | |
|---|---|---|---|---|
| PEB_1083-fw | CCCGGCTTTGGCGTTGCGGAGCCC_ATGAGCGACCTGCTTGCGCGGGAG | pEB141 p*pipDFGH* | pSEVA-ara_RS12625-RS12635+RS12615 (p*pipDFGH*) (2481) | *P. viridiflava*-pCEP*pipA*+p*pipDFGH* (2488) *P. syringae*-pCEP*pipA*+p*pipDFGH* (2484) |
| PEB_1085-rv | CTAGTCGCCAGGGTTTTCCCAGTCACGAC_TCAGTGCCTGTTGATCCATACG | | | |
| AR_900-fw | GTCGTGACTGGGAAAACCCT | | | |
| PEB-1086-rv | GATTCATCCGCGATGCGGTC | | | |
| PEB-1083-fw | GCCCATGAGACCGCATCGCGGATGAATC_AGCGACCTGCTTGCGCGG | pEB145 p*pipDEFGH* | pSEVA-ara_RS12615-12635 (*pipFGH-DE*) (2552) | ΔpipD-H_pCEP*pipA* + p*pipDEFGH* #2 (PH2482xpEB145) (2553) |
| PEB-2031-rv | GTCGCCAGGGTTTTCCCAGTCACGAC_CTCTAGAGCGCTGCATACCCTG | | | |
| PEB-1065-fw | GTTTAATACTAGAGAAAGAGGGGAAATACTAG_GTGGACTTCTTCACCCTGGC | | | |
| PEB_796-rv | CATCGTTGCTGCTGCGTAAC | | | |

**Table S1.9.** Primers, plasmids and generated *E. coli* strains for promoter exchange strains in *P. syringae* and *P. viridiflava*

| Name | Sequence | *E. coli* strain | Purpose | Strain |
|---|---|---|---|---|
| PEB_1038-fw | TTTGGGCTAACAGGAGGCTAGCATATGCTCGACAGATCTTCCTC | 2365 pCEP_LMDDJHCF_09560 (pPH65) | Promoter exchange to LMDDJHCF_09560 | (2369) *P. viridiflava*-pCEP*pipA* |
| PEB_1039-rv | CCGTTTAAACATTTAAATCTGCAGCCTGGTGATGATCACTGAC | | | |
| VPEB_1040-rv | CTCGACAACCTGCTGCAATGTG | | | |
| PEB_1041-fw | TTTGGGCTAACAGGAGGCTAGCATATGGTCGACAAATCTTCCTC | 2366 pCEP_GHKPGMKM_08200 (pPH66) | Promoter exchange to GHKPGMKM_08200 | (2367) *P. syringae*-pCEP*pipA* |
| PEB_1042-rv | CCGTTTAAACATTTAAATCTGCAGCGAGGTGTAGATCACATACG | | | |
| VPEB_1043-rv | CTCGCTCAACAGGCGATACAG | | | |

**Table S1.8.** Sum formulas of all $[M+H]^+$ ions detected and calculated with Δppm.

| Compound | Sum formula | detected | calculated | Δppm |
|---|---|---|---|---|
| **1a** | $C_{13}H_{25}N_2O_4$ | 273.1805 | 273.1809 | 1.3 |
| **1b** | $C_{13}H_{25}N_2O_5$ | 289.1754 | 289.1758 | 1.4 |
| **1c** | $C_{15}H_{27}N_2O_6$ | 331.1863 | 331.1864 | 0.1 |
| **2a** | $C_{12}H_{23}N_2O_4$ | 259.1647 | 259.1652 | 2.1 |
| **3a** | $C_{19}H_{32}N_7O_4S_2$ | 486.1955 | 486.1951 | 0.7 |
| **3b** | $C_{21}H_{34}N_7O_5S_2$ | 528.2071 | 528.2075 | -2.7 |
| **3c** | $C_{21}H_{34}N_7O_6S_2$ | 544.2002 | 544.2007 | 0.8 |
| **4a** | $C_{32}H_{54}N_9O_7S_2$ | 740.3605 | 740.3582 | -3.1 |
| **4b** | $C_{32}H_{54}N_9O_8S_2$ | 756.3538 | 756.3531 | -0.9 |
| **4b'** | $C_{32}H_{54}N_9O_8S_2$ | 756.3547 | 756.3531 | -2.1 |
| **4c** | $C_{34}H_{56}N_9O_8S_2$ | 782.3697 | 782.3688 | -1.2 |
| **4d** | $C_{34}H_{56}N_9O_9S_2$ | 798.3646 | 798.3637 | -1.2 |
| **4d'** | $C_{34}H_{56}N_9O_9S_2$ | 798.3637 | 798.3637 | 0 |
| **4e** | $C_{36}H_{58}N_9O_{10}S_2$ | 840.376 | 840.3743 | -1.7 |
| **4f** | $C_{32}H_{54}N_9O_8S_2$ | 756.3536 | 756.3531 | -0.6 |
| **4g** | $C_{34}H_{56}N_9O_9S_2$ | 798.3656 | 798.3637 | -2.3 |
| **4h** | $C_{38}H_{60}N_9O_{11}S_2$ | 882.3848 | 882.3848 | 0 |
| **5a** | $C_{31}H_{52}N_9O_7S_2$ | 726.3445 | 726.3426 | -2.6 |
| **5b/5b'** | $C_{31}H_{52}N_9O_8S_2$ | 742.3394 | 742.3375 | -2.6 |
| **5c** | $C_{33}H_{54}N_9O_8S_2$ | 768.3548 | 768.3531 | -2.2 |
| **5d** | $C_{33}H_{54}N_9O_9S_2$ | 784.3473 | 784.3473 | 0.9 |
| **5d'** | $C_{33}H_{55}N_9O_9S_2$ | 392.6773 | 392.6777 | 1 |
| **5e** | $C_{35}H_{56}N_9O_{10}S_2$ | 826.358 | 826.3586 | -2.1 |
| **5g** | $C_{33}H_{55}N_9O_9S_2$ | 392.6775 | 392.6777 | 1 |
| **5h** | $C_{37}H_{58}N_9O_{11}S_2$ | 868.369 | 868.3692 | 0.2 |

### Supplementary Figures

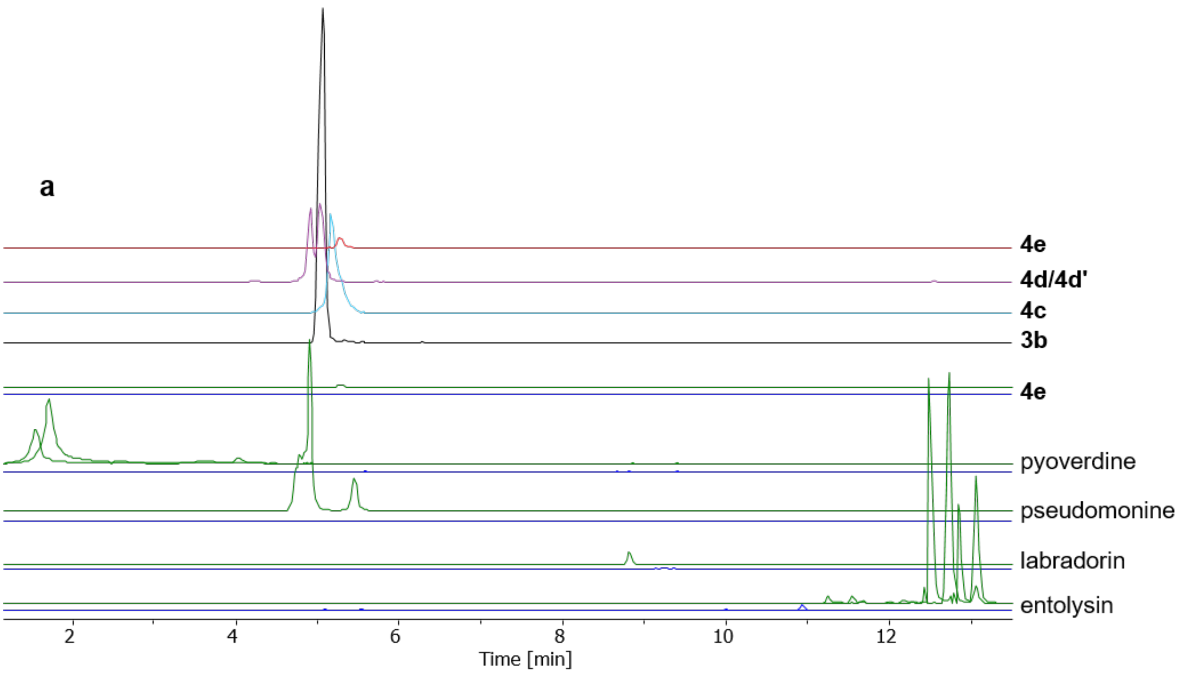


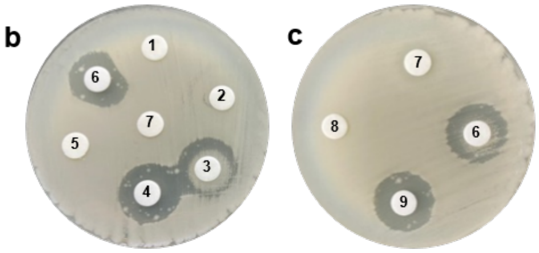


| disc | 862 | BCS | 7001 | 849 |
|---|---|---|---|---|
| 1 | 20 | 1 | | |
| 2 | 2 | 1 | | |
| 3 | 0.2 | 1 | | |
| 4 | 0.02 | 1 | | |
| 5 | 20 | | | |
| 6 | | 1 | | |
| 7 | MeOH | | | |
| 8 | | 1 | 20 | |
| 9 | | 1 | | 20 |

**Fig. S1.1.** Testing for anti-antibiotic activity. (**a**) EICs of crude extracts from ΔPELP4_pCEP*pipA* (top four lines), ΔPELP4 (blue) and WT (green). EICs for the WT are pyoverdines *m/z* 665.8 $[M+2H]^{2+}$ and 657.8 $[M+2H]^{2+}$, pseudomonine *m/z* 331.1 $[M+H]^{+}$, labradorin *m/z* 241.1 $[M+H]^{+}$ , and entolysines *m/z* 861.1 $[M+2H]^{2+}$, and 870.1 $[M+2H]^{2+}$. (**b** & **c**) Agar diffusion assay of different extract/blasicidin S combinations with *B. cereus* as test organism for anti-antibiotic activity testing. Concentrations of applied extracts in mg/mL are listed in the Table. 10 µL of extract and/or blasicidin S was applied on the discs and dried before placing it on the inoculated agar surface. Inhibition zones indicate the antibiotic activity of blasticidine S.

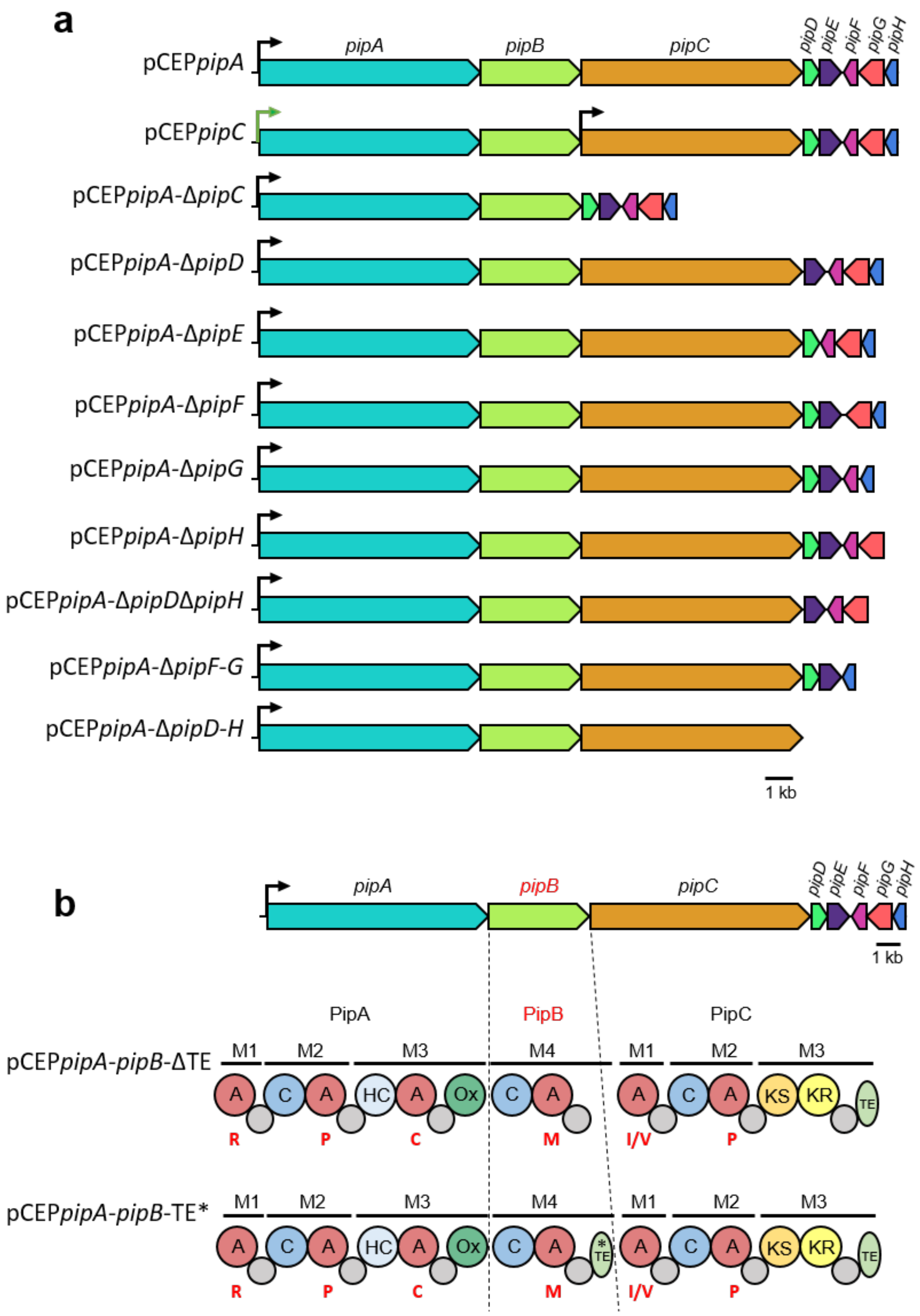


**Fig. S1.3.** (**a**) BGC of PipA-PipH and its genetic modifications via promoter exchanges and deletions of selected genes. Black arrow indicates the inserted promoter $P_{BAD}$, green arrow indicated the natural promoter. (**b**) BGC of pseudotetraivprolides focused on the modification of the TE domain in *pipB*. Top: Deletion of the TE domain, bottom: point mutation S(72)A within the TE domain indicated by an asterisk. A adenylation domain, C condensation domain, HC heterocyclisation domain, Ox oxidation domain, KS keto synthase domain, KR keto reductase domain, TE thioesterase domain, small grey cycle thiolation domain.

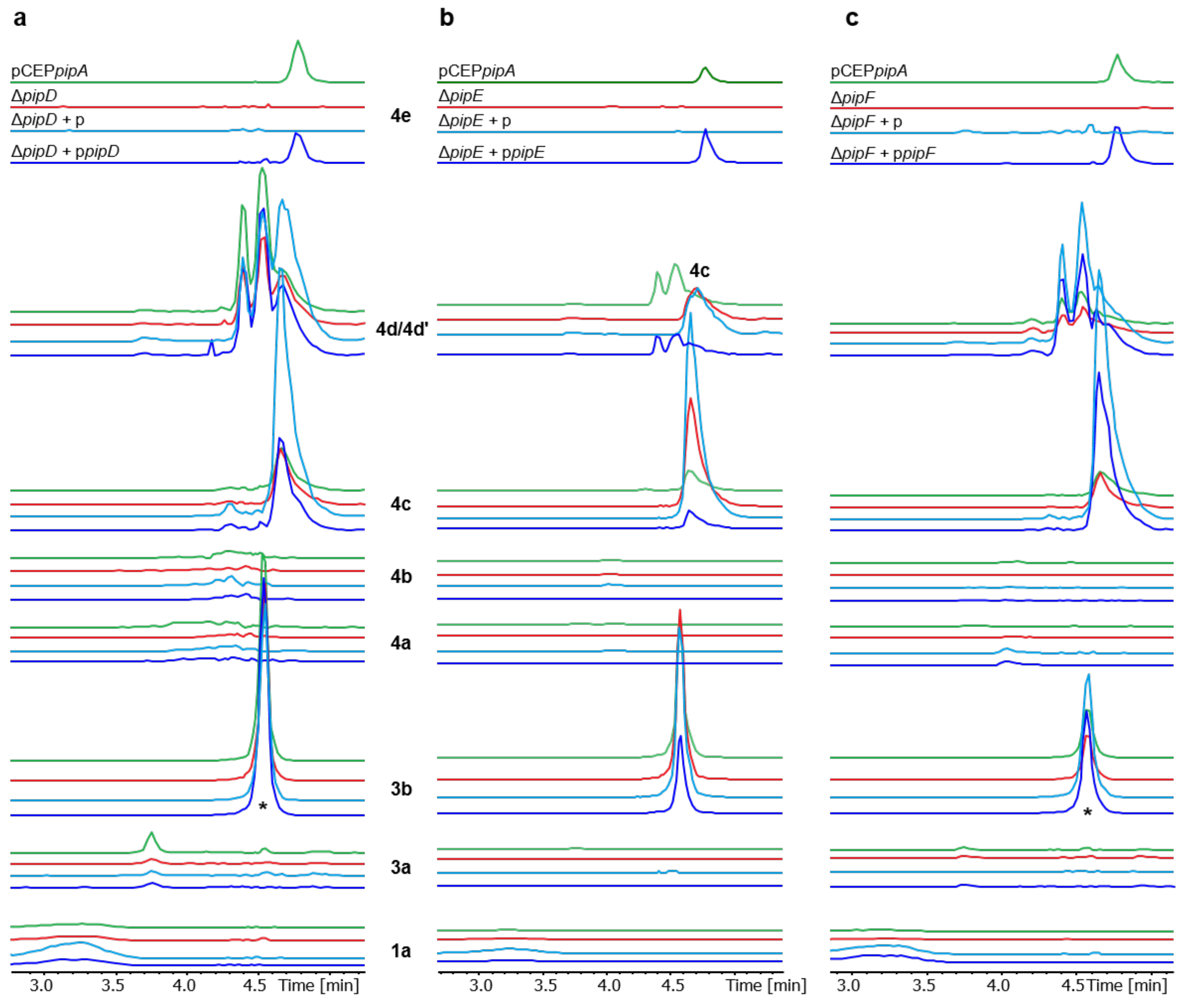


**Fig. S1.4.** Plasmid based complementation of deletion mutants. Depicted are the EICs of selected pip derivatives found in culture extracts of respective mutants. (**a**) Complementation of pCEP*pipA*-Δ*pipD* (red) with plasmid encoded p*pipD* (blue). (**b**) Complementation of of deletion mutant pCEP*pipA*-Δ*pipE* (red) with plasmid encoded p*pipE* (blue). (**c**) Complementation of of deletion mutant pCEP*pipA*-Δ*pipF* (red) with plasmid encoded p*pipF* Depicted are the EICs of selected pip derivatives found in culture extracts of respective mutants. Deletion mutants transformed with the empty vector pSEVA621-Gm as empty vector control (light blue) and compared with WT pCEP*pipA* (green). Asterisks below the peaks indicate a 10-fold increased signal.

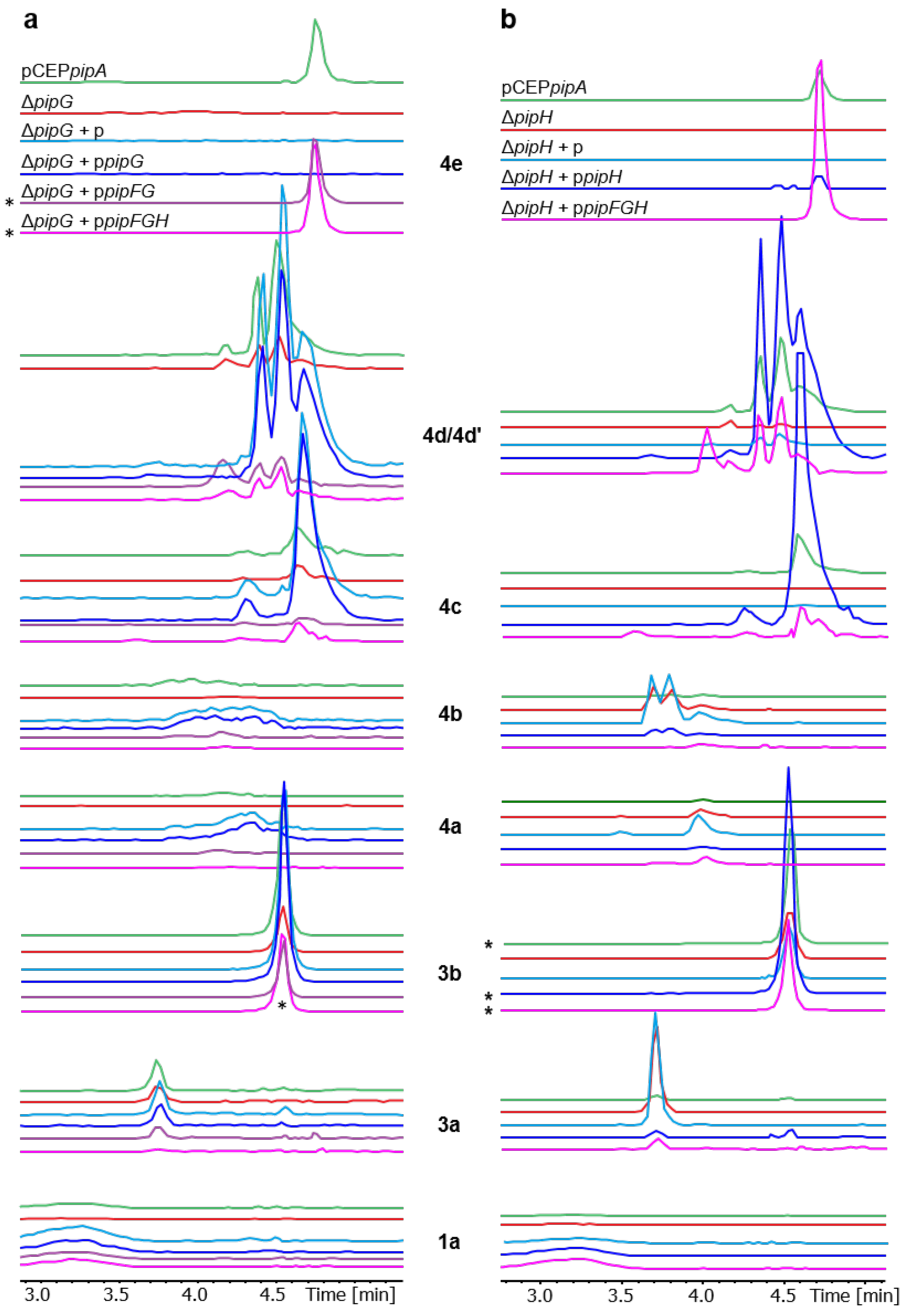


**Fig. S1.5.** (**a**) Plasmid based complementation of deletion mutant pCEP*pipA*-Δ*pipG* (red) with plasmid encoded p*pipG* (blue), p*pipFG* (violet) and p*pipFGH* (magenta), compared to pCEP*pipA* (green) and control strain Δ*pipG* + p (light blue). Depicted are the EICs of selected pip derivatives found in culture extracts of respective mutants (**a**). Plasmid based complementation of deletion mutant pCEP*pipA*-Δ*pipH* (red) with plasmid encoded p*pipH* (blue) and p*pipFGH* (magenta), compared to pCEP*pipA* (green) and control strain Δ*pipH* + p (light blue). Depicted are the EICs of selected pip derivatives found in culture extracts of respective mutants (**b**) Asterisk indicate a 10-fold decreased peak.

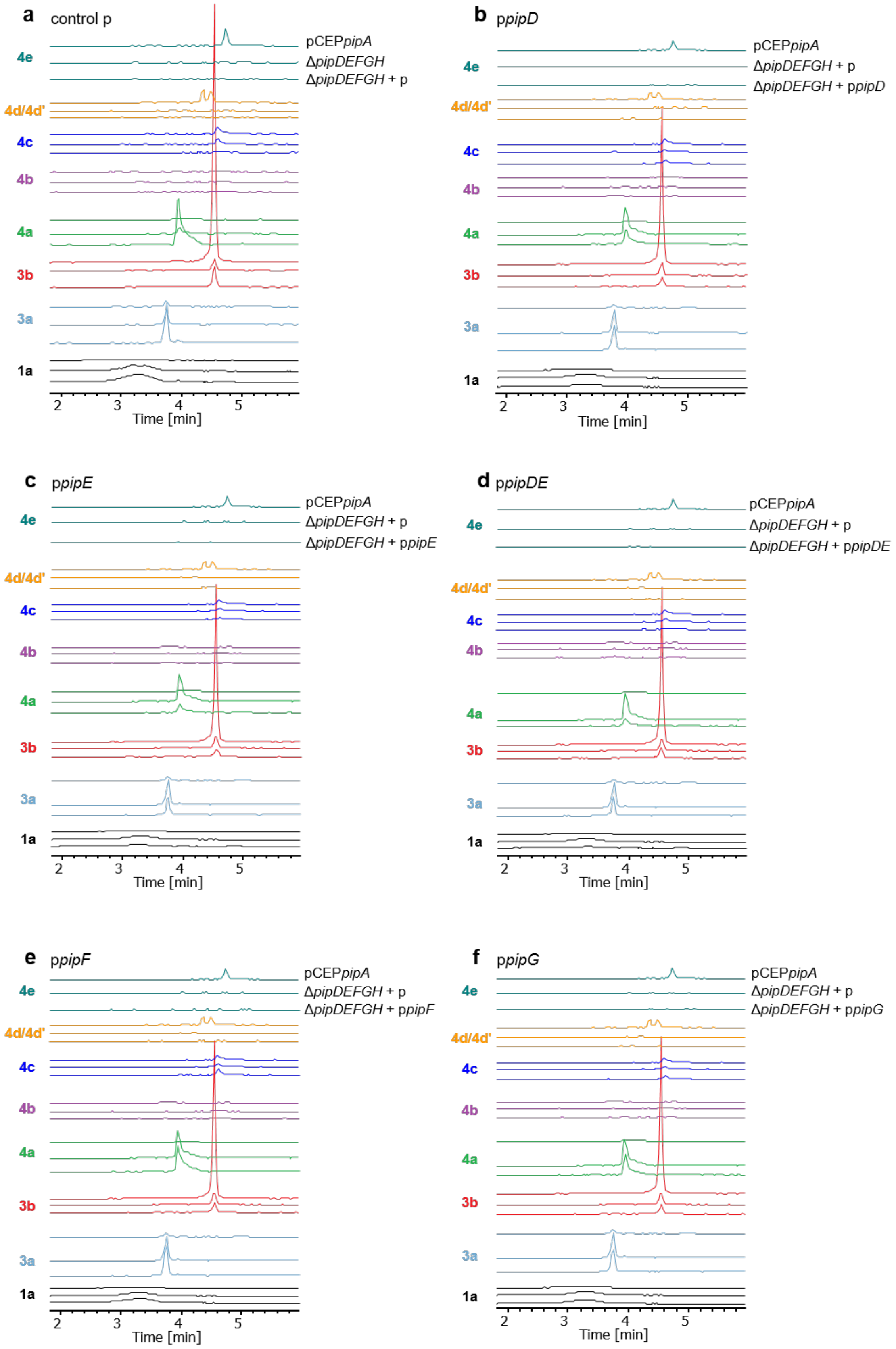
a
control p
pCEPpipA
ΔpipDEFGH
ΔpipDEFGH + p
b
ppipD
ΔpipDEFGH + p
ΔpipDEFGH + ppipD
c
ppipE
ΔpipDEFGH + ppipE
d
ppipDE
ΔpipDEFGH + ppipDE
e
ppipF
ΔpipDEFGH + ppipF
f
ppipG
ΔpipDEFGH + ppipG
4e
4d/4d'
4c
4b
4a
3b
3a
1a
2
3
4
5
Time [min]

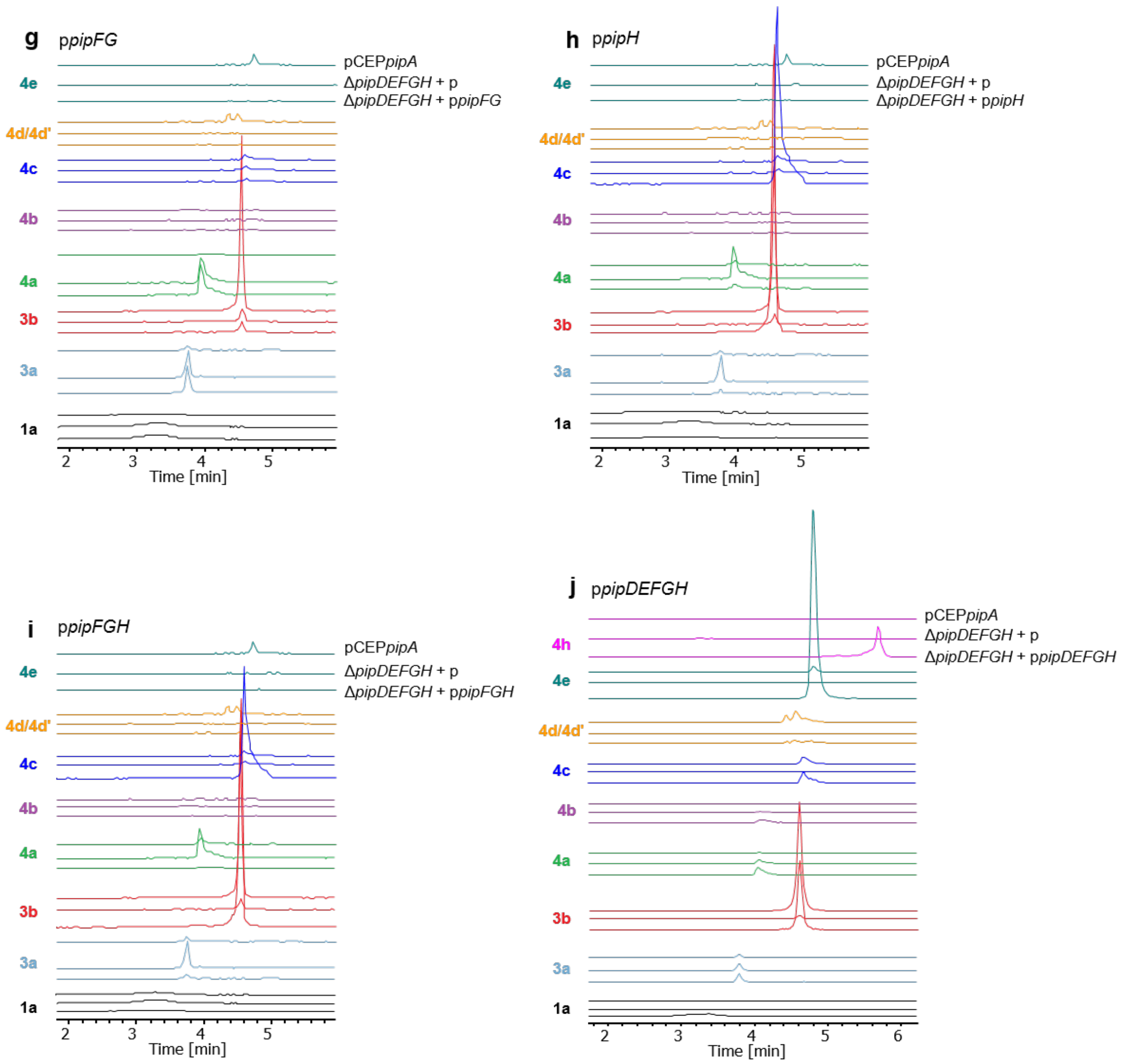


**Fig. S1.6.** Plasmid based complementation of deletion mutant Δ*pipDEFGH*-pCEP*pipA*. Depicted are the EICs of selected compounds from culture extracts of pCEP*pipA*Δ*pipDEFGH*-pCEP*pipA*+empty plasmid (p) after HPLC-MS analysis. (**a**) pCEP*pipA*, Δ*pipDEFGH*-pCEP*pipA* and empty vector control p (**b**) p*pipD*, (**c**) p*pipE*, (**d**) p*pipDE*, (**e**) p*pipF*, (**f**) p*pipG*, (**g**) p*pipFG*, (**h**) p*pipH*, (**i**) p*pipFGH*. (**j**) p*pipDEFGH*.

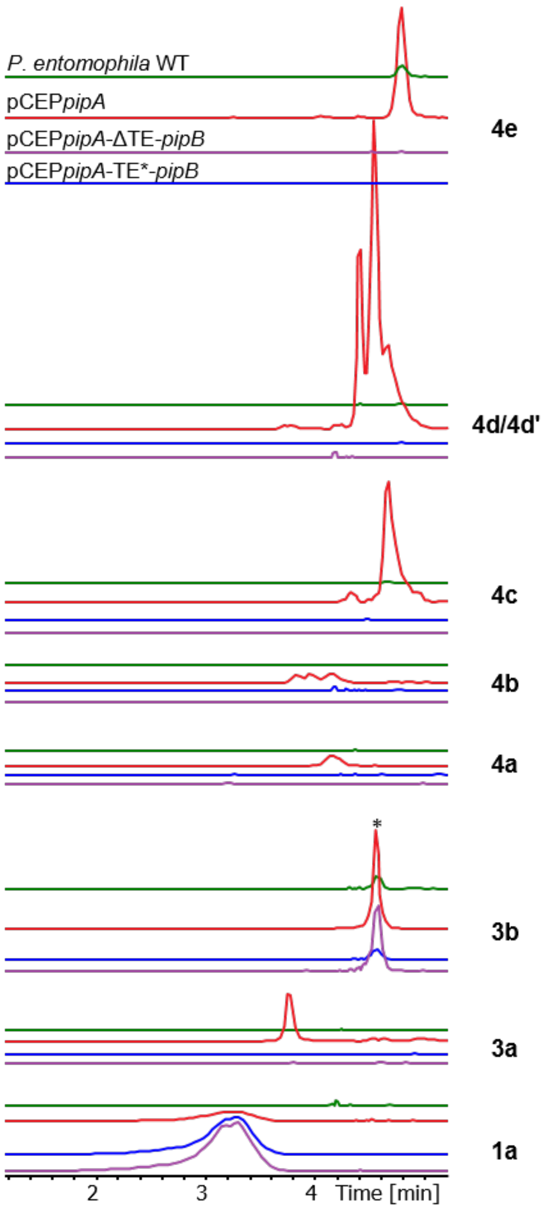


**Fig. S1.7.** Investigation of the TE domain of *pipB*. EICs of selected derivatives of pseudotetraivprolid detected in culture extracts of *P. entomophila* WT (green), pCEP*pipA* (red), pCEP*pipA*-ΔTE-*pipB* (blue) and pCEP*pipA*- TE*-*pipB* (violet). Asterisk indicate a 20-fold decreased signal intensity.

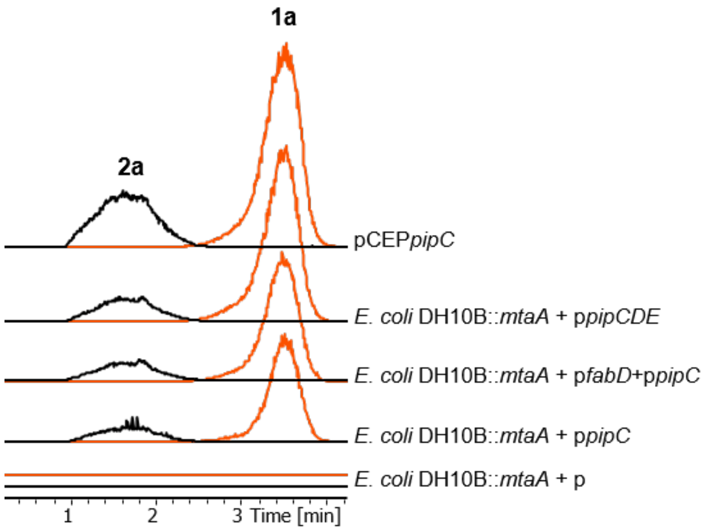


**Fig. S1.8.** Heterologous expression of p*pipCDE* and pPSEEN_RS07520 (p*fabD*) in *E. coli* DH10B::*mtaA*. EICs of **1a** and **2a** in HPLC-MS analysis of culture extracts from pCEP*pipC*, plasmid encoded p*pipC* and co-expression of plasmid encoded p*pipC* and p*fabD* in *E. coli* DH10B::*mtaA*. No hydroxylated or O-acetylated derivatives **1b** and **1c** have been detected.

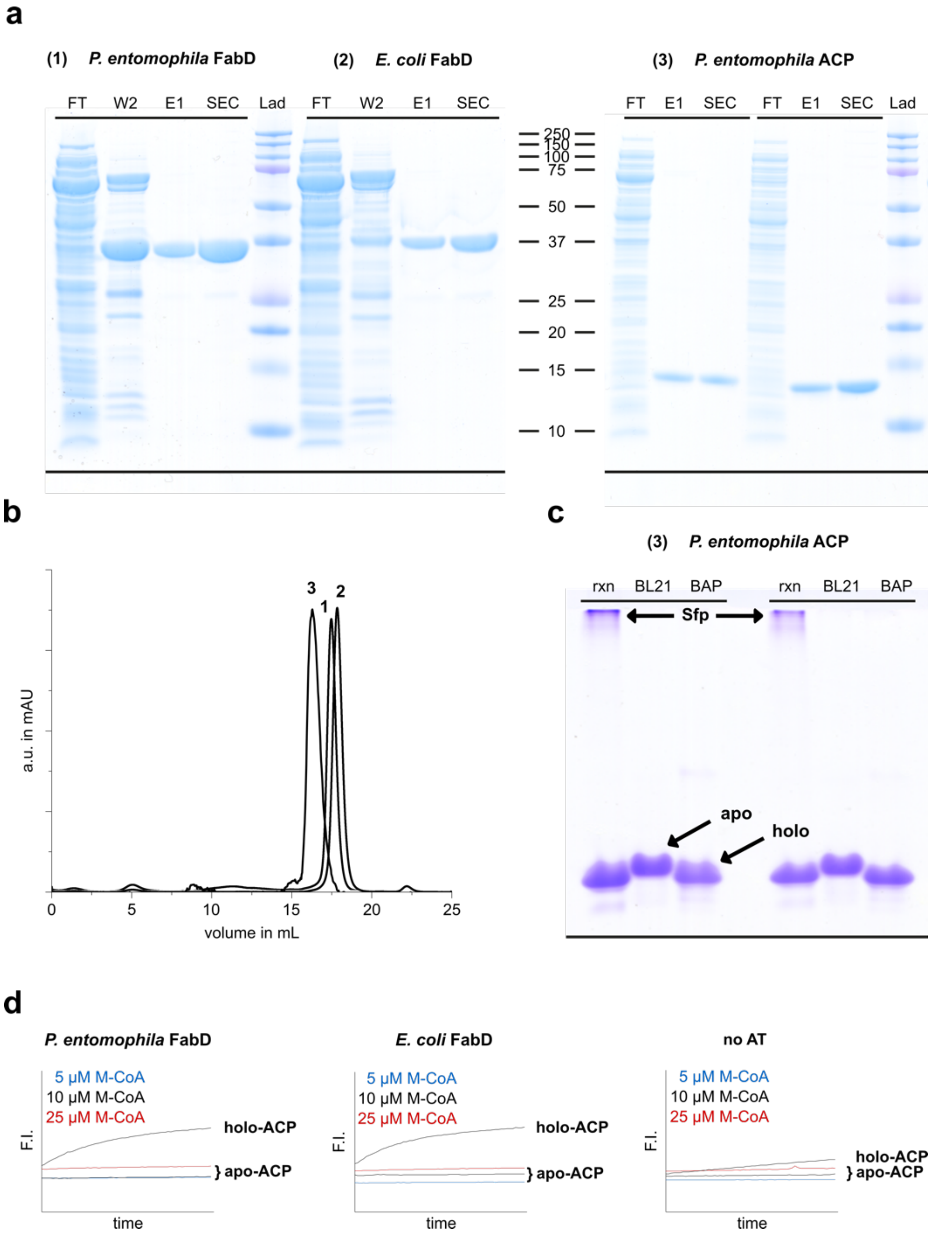


**Figure S1.9.** Protein Quality Control of *P. entomophila* FabD, *E. coli* FabD and PipC ACP. Highly pure protein was obtained for both AT domains as well as ACP domain (shown in biological duplicate) from heterologous expression in BAP1 cells. (**a**) SDS-PAGE to monitor purification protocol. Proteins were purified from 2-step protocol (Ni-chelating affinity chromatography followed by polishing via size exclusion chromatography). Protein sizes were observed as expected: *P. entom.* FabD: ≈ 36.9 kDa, *E. coli* FabD: ≈ 37.1 kDa, PipC ACP: ≈ 14.0 kDa (FT = flowthrough, E1 = Elution, SEC = Size Exclusion Chromatography, Lad = Precision Plus Protein Ladder). (**b**) Size exclusion chromatograms of purified proteins. *P. entom.* FabD (1; $V_E$ = 17.5 mL) and *E. coli* FabD (2; $V_E$ = 17.8 mL) were purified using a Superose$^{TM}$ 6 Increase 10/300 GL column, and PipC ACP (3; $V_E$ = 16.2 mL) was purified via a Superdex$^{TM}$ 200 Increase 10/300 GL column. (**c**) Conformation-sensitive urea PAGE for analysis of PipC ACP domain. PipC ACP was expressed in apo- (BL21 (DE3), middle) and holo-form (BAP1, genetically-encode 4'-phosphopantetheinyl transferase from *Bacillus subtilits* (Sfp), right). For comparison, phosphopantetheinylation was conducted *in vitro* using purified Sfp, CoA-SH and apo-ACP (expressed in BL21 (DE3) cells, left). Expression of PipC ACP in BAP1 cells solely provided holo-ACP. (**d**) Initial screening of enzyme activity at different substrate concentrations via in vitro assay. Holo-ACP was active with both FabD enzymes while apo-ACP was inactive (measurement performed in technical triplicates of biological duplicates).

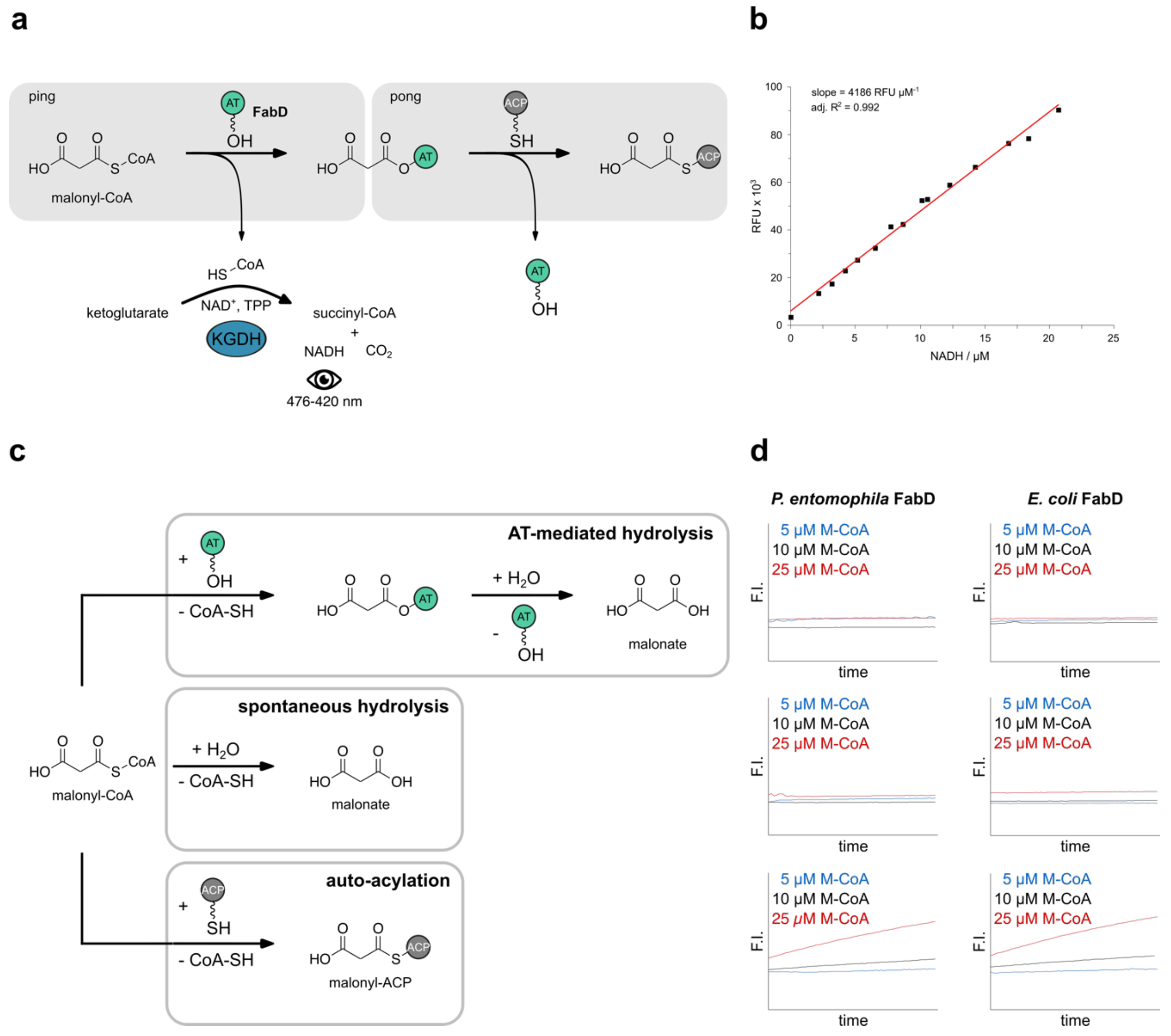


**Figure S1.10.** Overview of the alpha-KGDH assay. (**a**) AT-mediated transacylation reaction can be monitored via an enzyme-coupled assay. AT domains catalyze the transfer of an CoA-activated acyl-moiety (here: malonyl-CoA, Mal-CoA) to a holo-ACP domain via a double displacement reaction (ping-pong bi-bi mechanism). Upon release of CoA-SH, the acyl-moiety is loaded onto the AT domain (ping step). Next, the acyl-moiety is transferred onto the ACP-domain to give acyl-ACP (pong step). The release of CoA-SH can be couple to the KGDH-catalyzed formation of succinyl-CoA from ketoglutarate, CoA-SH and $NAD^+$. The reduction of $NAD^+$ can be monitored fluorometrically and enables quantification of the transacylation rate. (**b**) NADH calibration curve. To quantify AT activity, a calibration was conducted to correlate relative fluorescence units (RFU) to NADH concentration (µM). Measurement was performed in technical triplicates (**c**) Several side reactions can occur during AT-mediated transacylation that need to be taken into account when determining kinetic parameters. Eventually, formation of large quantities of CoA-SH can falsify kinetic constants. In the absence of an ACP domain, the acyl-X substrate is hydrolyzed (top: AT-mediated hydrolysis; middle: spontaneous hydrolysis). When ACP is present, the acyl-X substrate can be directly transferred onto the

ACP domain (bottom: auto-acylation). (**d**) Influence of CoA-SH forming side reactions was evaluated for *E. coli* FabD and *P. entomophila* FabD. For this, fluorescence intensity was measured in the absence of ACP (top), absence of AT and ACP (middle), and in the absence of AT (bottom). For both AT domains, hydrolysis (spontaneous or AT-mediated) does not influence the fluorescence intensity. In contrast, auto-acylation is a significant background reaction at higher substrate concentrations (≥25 µM). Consequently, F.I values of active measurements have been corrected by the background activity (Measurements were performed in technical triplicates. Blots are shown as average of biological duplicates).

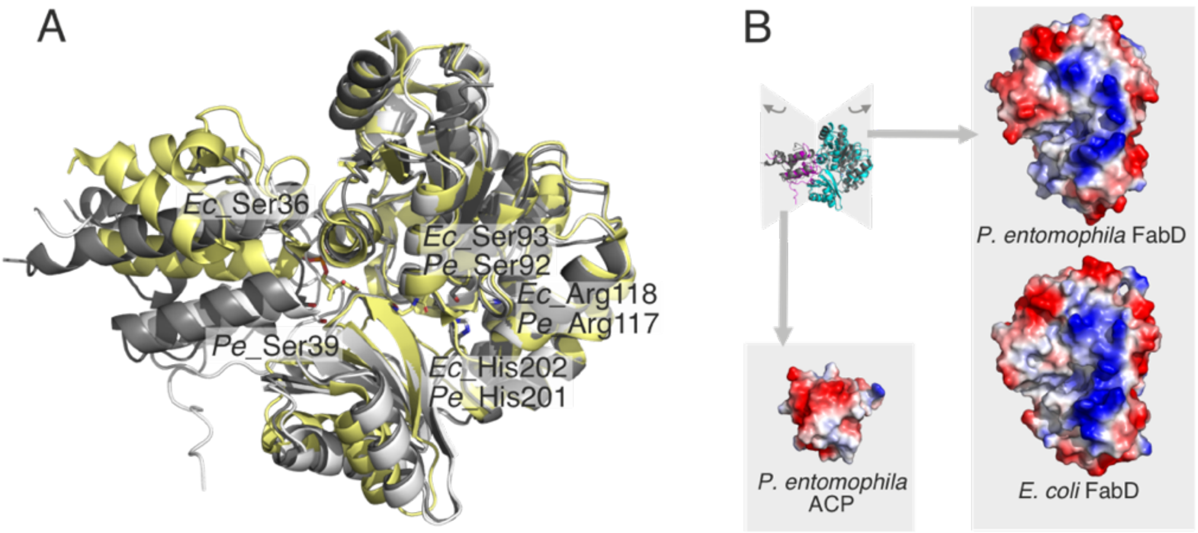


**Figure S1.11.** Structure of ACP-AT interaction. (A) ACP-AT complexes of P. entomophila ACP (PipC module 3) and P. entomophila AT (dark grey), P. entomophila ACP (PipC module 3) and E. coli FabD (white), both complexes modeled with AlphaFold3, and experimental structural data on the E. coli ACP-FabD complex (yellow) (PDB-ID 6u0j)[1]. Selected residues are highlighted in stick representation, as well as the covalent crosslinker in the FabD binding pocket. The predicted structures superimpose well in the ACP positioning, while the original data on the E. coli complex show a tilted position. The serine, carrying the phosphopantetheine after post-translational modification is plausible position, meaning that a phosphopantetheine could reach the AT active site. (B) Vacuum electrostatics of the ACP:AT interface. The binding interface is shown for both complexes (P. entomophila ACP-E. coli FabD complex colored superimposed with P. entomophila complex in grey). The ACP interface is negatively charged and the opposing AT interface positively charged, indicated molecular recognition that drives interaction. The figure is prepared with pymol. Structures have been superimposed with the pymol superposition tool and the charge distribution on the protein surfaces estimated with the pymol vacuum electrostatics tool (www.pymol.org).

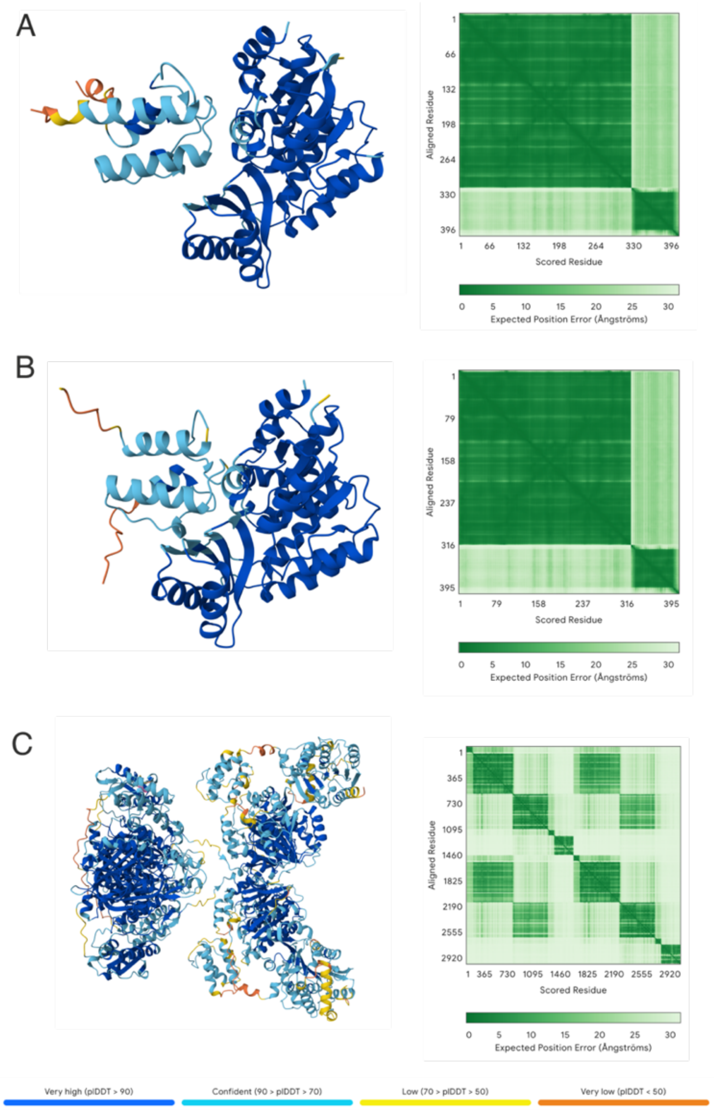


**Figure S1.12.** Assessment of model confidence and structural accuracy. Quality assessment for predicted structures of *P. entomophila* ACP (PipC module 3):*P. entomophila* AT (A), *P. entomophila* ACP:*E. coli* FabD (B), and PipC module 3 with N-terminal ACP domain (C). Orientation of structures as in Fig. 5. (Left panels) per-residue confidence scores (pLDDT) predicted by AlphaFold3, plotted across the sequence. Higher pLDDT values (>90) indicate regions of high structural reliability, while lower scores (<70) suggest flexible or uncertain regions. (Right panels) Predicted Alignment Error (PAE) plot showing the estimated positional error between residue pairs. Lower PAE values between residues from different domains indicate high confidence in their relative positioning, while higher values reflect greater uncertainty.

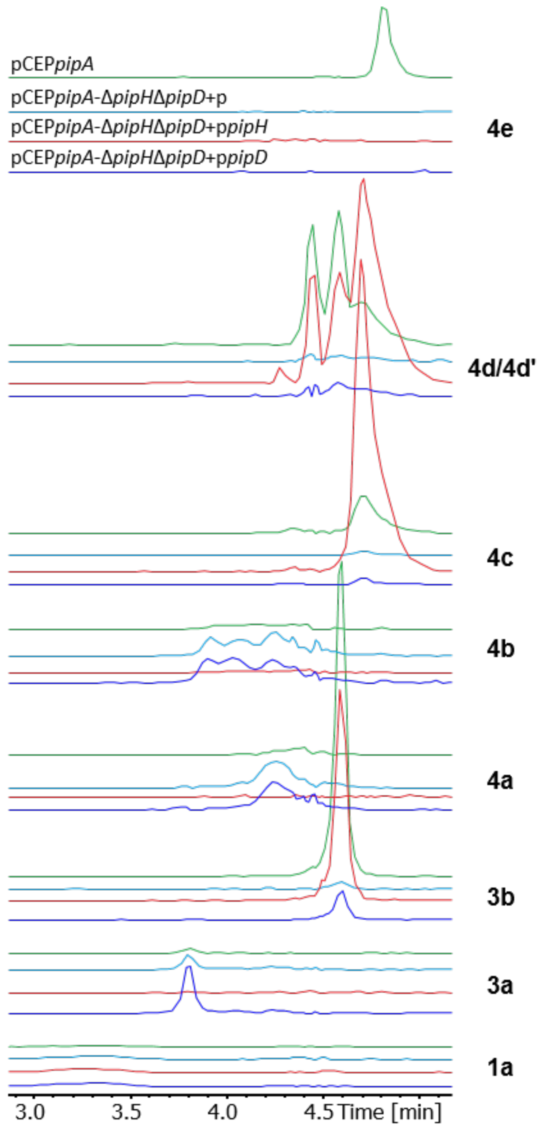


**Fig. S1.13.** Investigation on activities of PipH and PipD regarding N-terminal acetylation. EICs of selected derivatives of pip detected by HPLC-MS in culture extracts of pCEP*pipA* (green), pCEP*pipA*-Δ*pipD*Δ*pipH*+p (light blue) and complemented deletion mutants pCEP*pipA*-Δ*pipD*Δ*pipH*+p*pipH* (red) and pCEP*pipA*-Δ*pipD*Δ*pipH*+p*pipD* (blue).

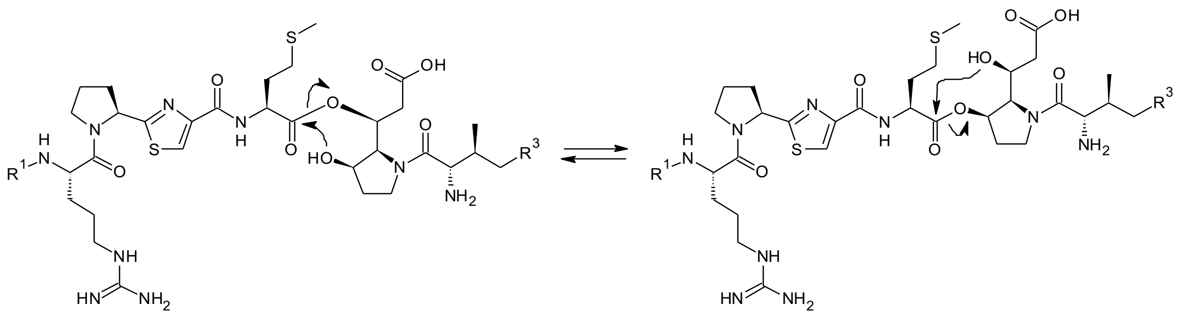

**Fig. S1.14.** Postulated transesterification of hydroxylated derivatives **4b** and **4d**, leading to two isobaric variants with an almost identical fragmentation pattern (see Fig. S2.7 & S2.9).

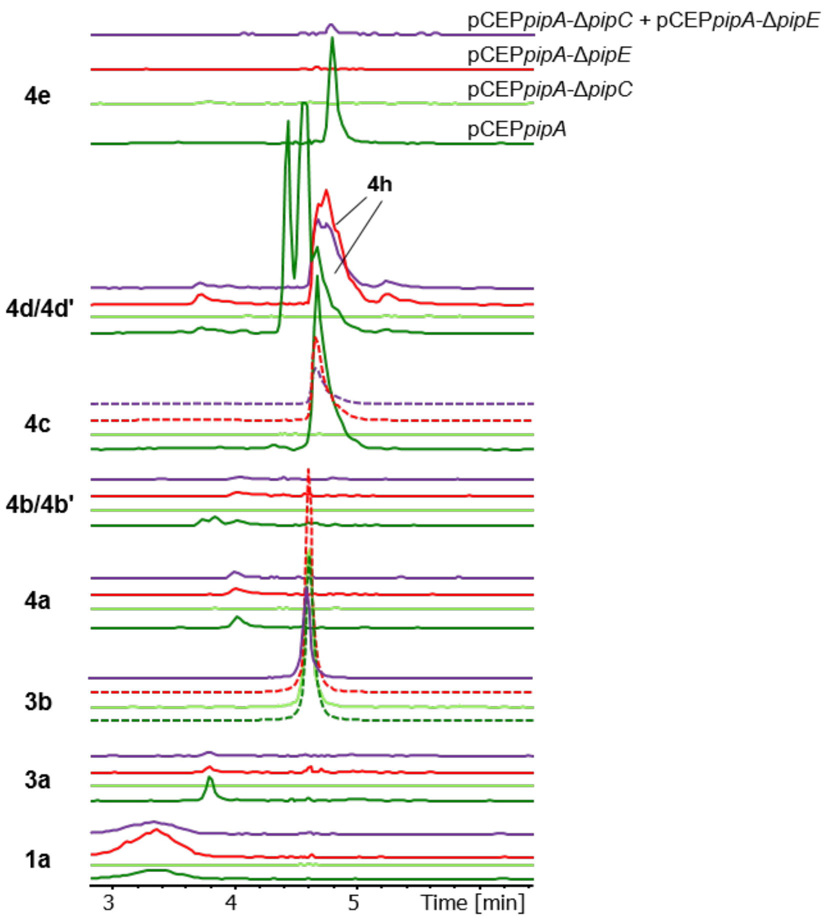


**Fig. S1.15.** Derivatives of pseudotetraivprolide detected in extracts of co-cultivated pCEP*pipA*-Δ*pipE* + pCEPpipA-Δ*pipC* (violet) in comparison to derivatives produced by single cultivated mutants pCEP*pipA* (green), pCEP*pipA*-Δ*pipE* (red), pCEP*pipA*-Δ*pipC* (neon green). dashed lines indicate a ten times reduced signal intensity.

**a**

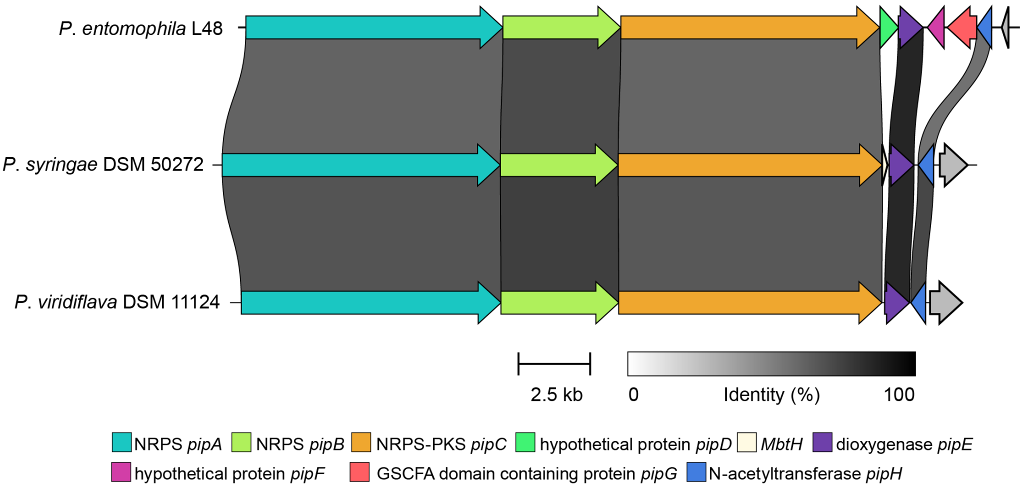


**Fig. S1.16.** Organization of pip BGC in *P. entomophila*, *P. syringae* and *P. viridiflava* analyzed via CAGECAT analysis.

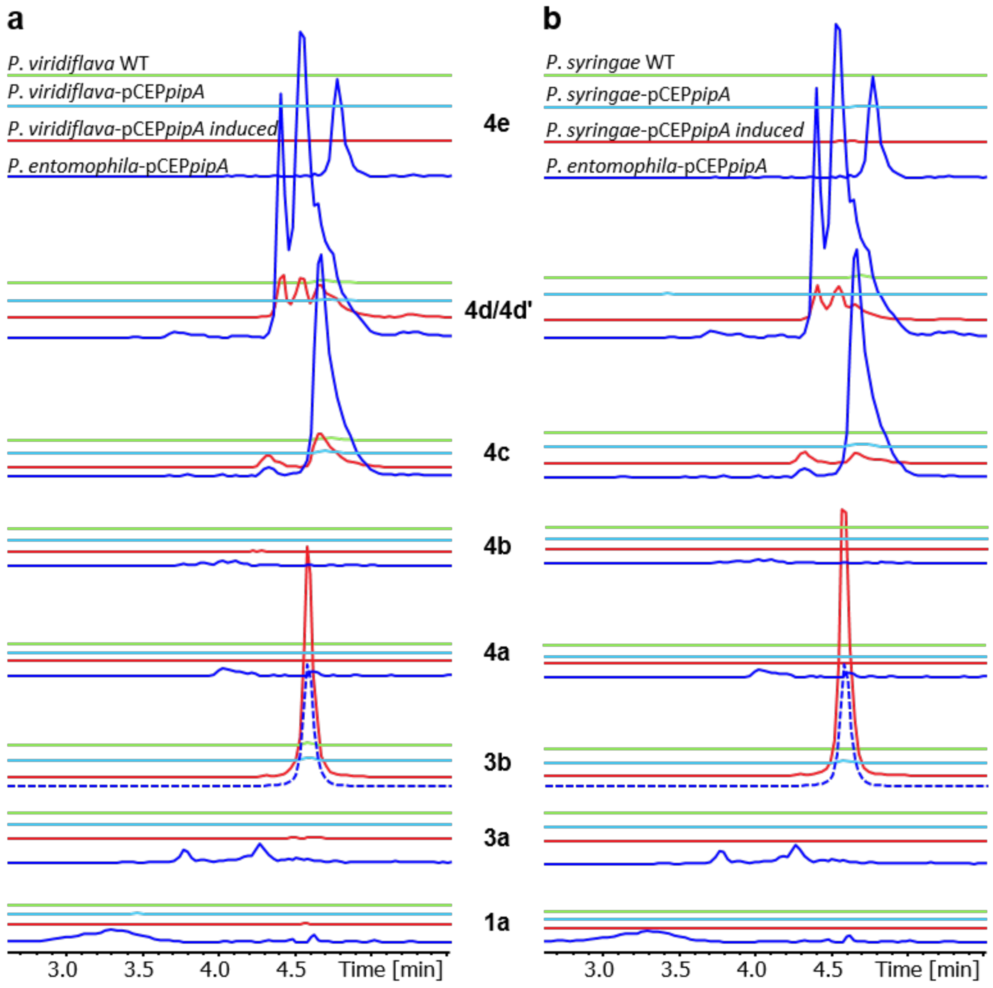


**Fig. S1.17.** (**a**) EICs of selected derivatives of pip detected via HPLC-MS analysis in culture extracts of *P. viridiflava* WT (green), *P. viridiflava*-pCEP*pipA* non induced (turquoise), induced (red) and *P. entomophila*-pCEP*pipA* induced (blue). (**b**) *P. syringae* WT (green), *P. syringae*-pCEP*pipA* non induced (turquoise) and *P. syringae*-pCEP*pipA* induced (red). dashed lines indicate a 10-fold decreased signal intensity.

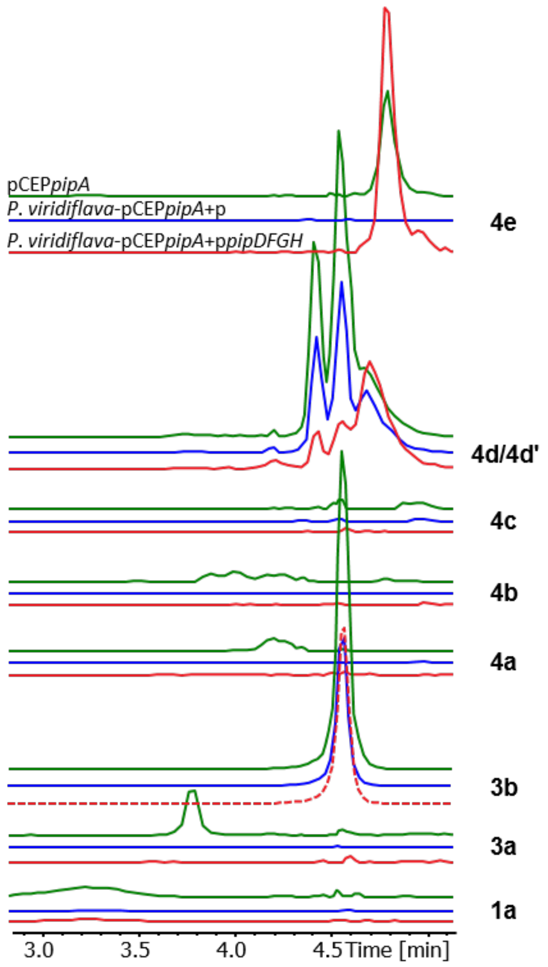


**Fig. S1.18.** Complementation of *P. viridiflava*-pCEPpipA with p*pipDFGH*. EICs of selected derivatives of pip detected via HPLC-MS analysis in culture extracts of pCEP*pipA* (green), *P. viridiflava*-pCEP*pipA*+p (blue) induced and *P. viridiflava*-pCEP*pipA*+p*pipDFGH* induced (red). Asterisk indicates a 10-fold increased signal intensity.

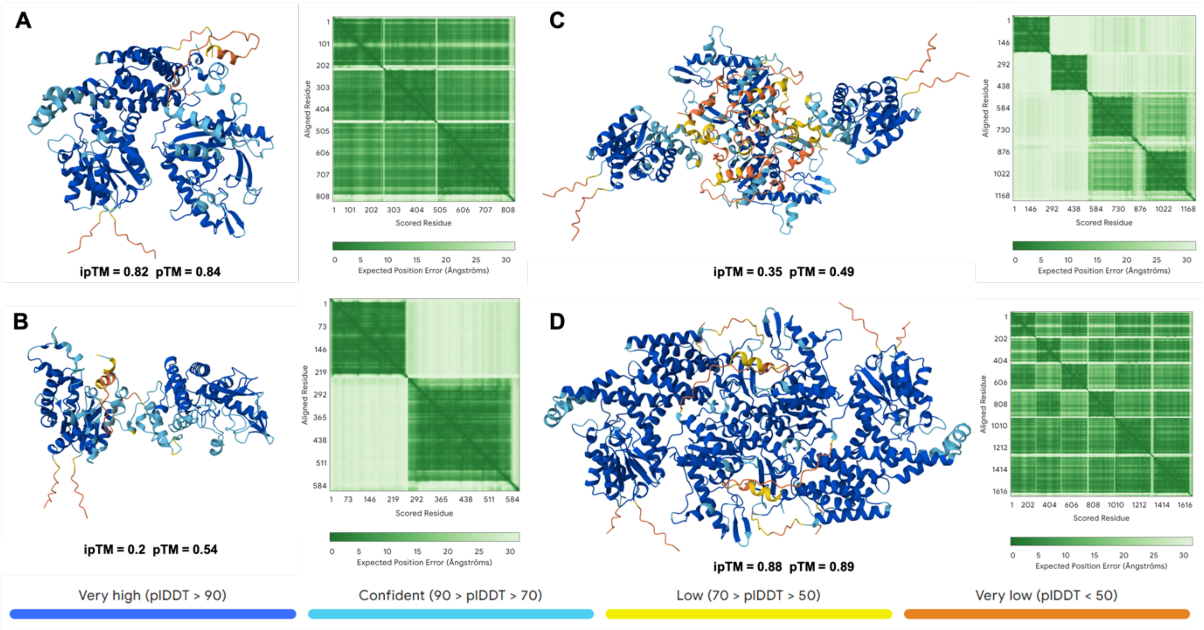


**Figure S1.19.** Dimeric, trimeric and higher order complexes structure predicted by AlphaFold3 and assessment of model confidence and structural accuracy. Quality assessment for predicted structures of PipDFG trimeric complexes (**A**), PipFG dimeric complexes (**B**), double copy dimeric complexes structure of PipFG (**C**) and double copy trimeric complexes structure of PipDFG (**D**). (Left panels for each sub-graph) per-residue confidence scores (pLDDT) predicted by AlphaFold3, plotted across the sequence. Higher pLDDT values (>90) indicate regions of high structural reliability, while lower scores (<70) suggest flexible or uncertain regions. (Bottom of right panels for each sub-graph) pTM and ipTM scores predicted by AlphaFold3, A pTM score above 0.5 means the overall predicted fold for the complex might be similar to the true structure. ipTM measures the accuracy of the predicted relative positions of the subunits within the complex. Values higher than 0.8 represent confident high-quality predictions, while values below 0.6 suggest likely a failed prediction. ipTM values between 0.6 and 0.8 are a gray zone where predictions could be correct or incorrect.
