## Supplementary Information SI-2 for "Identification of pseudotetraivprolide from *Pseudomonas entomophila* give novel insights into the biosynthesis of detoxin/rimosamide-like anti-antibiotics"

### Supporting information SI-2: Compound identification and structure elucidation

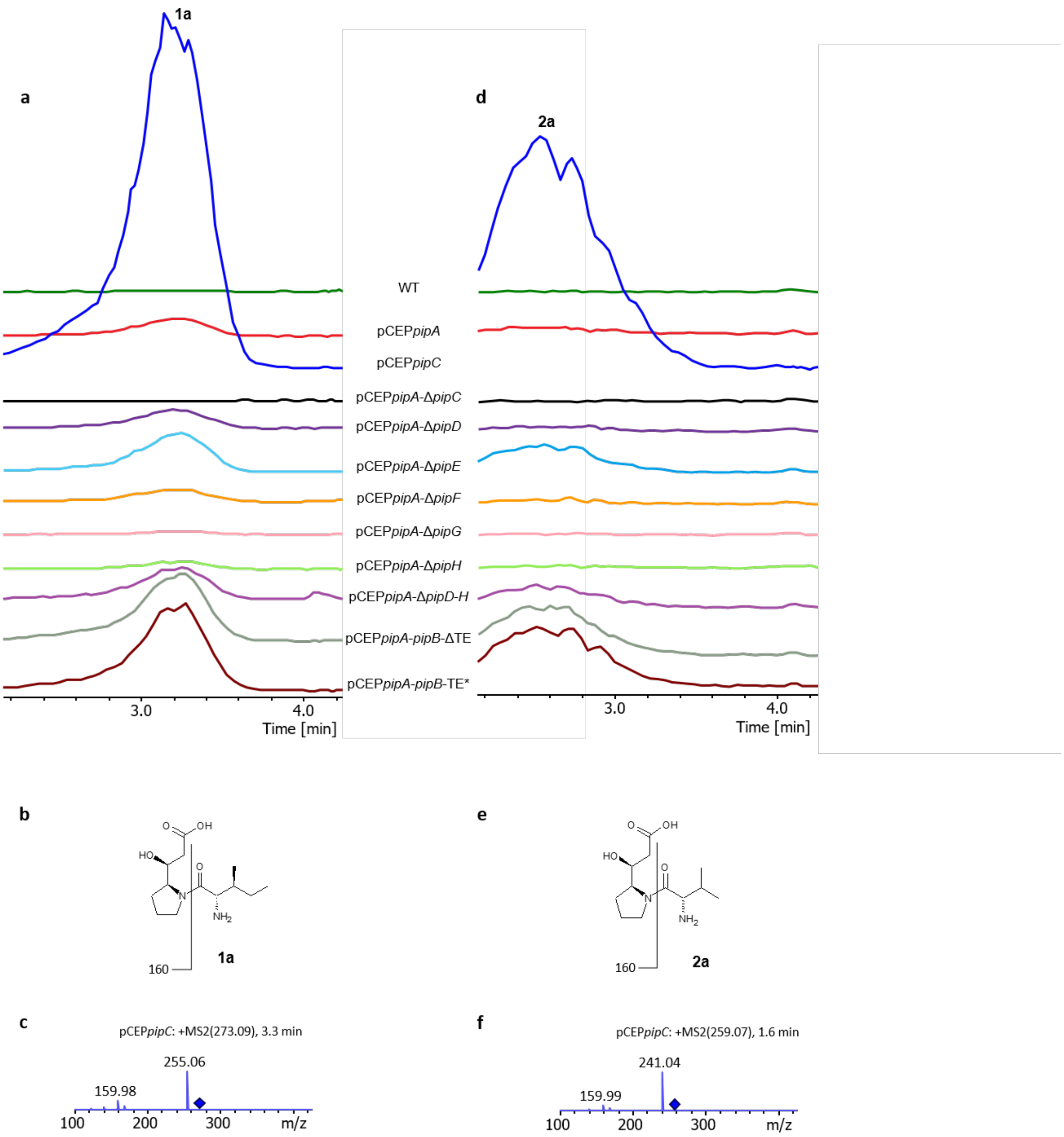


**Figure S2.1.** Extracted ion chromatograms (EICs) of compound **1a** after HPLC-MS analysis of selected culture extracts (**a**). Chemical structure of **1a** (**b**). $MS^2$ fragmentation pattern of **1a** (**c**). Extracted ion chromatograms (EICs) of compound **2a** after HPLC-MS analysis of culture extracts selected mutants (**d**). Chemical structure of **2a** (**e**). $MS^2$ fragmentation pattern of **2a** (**f**).

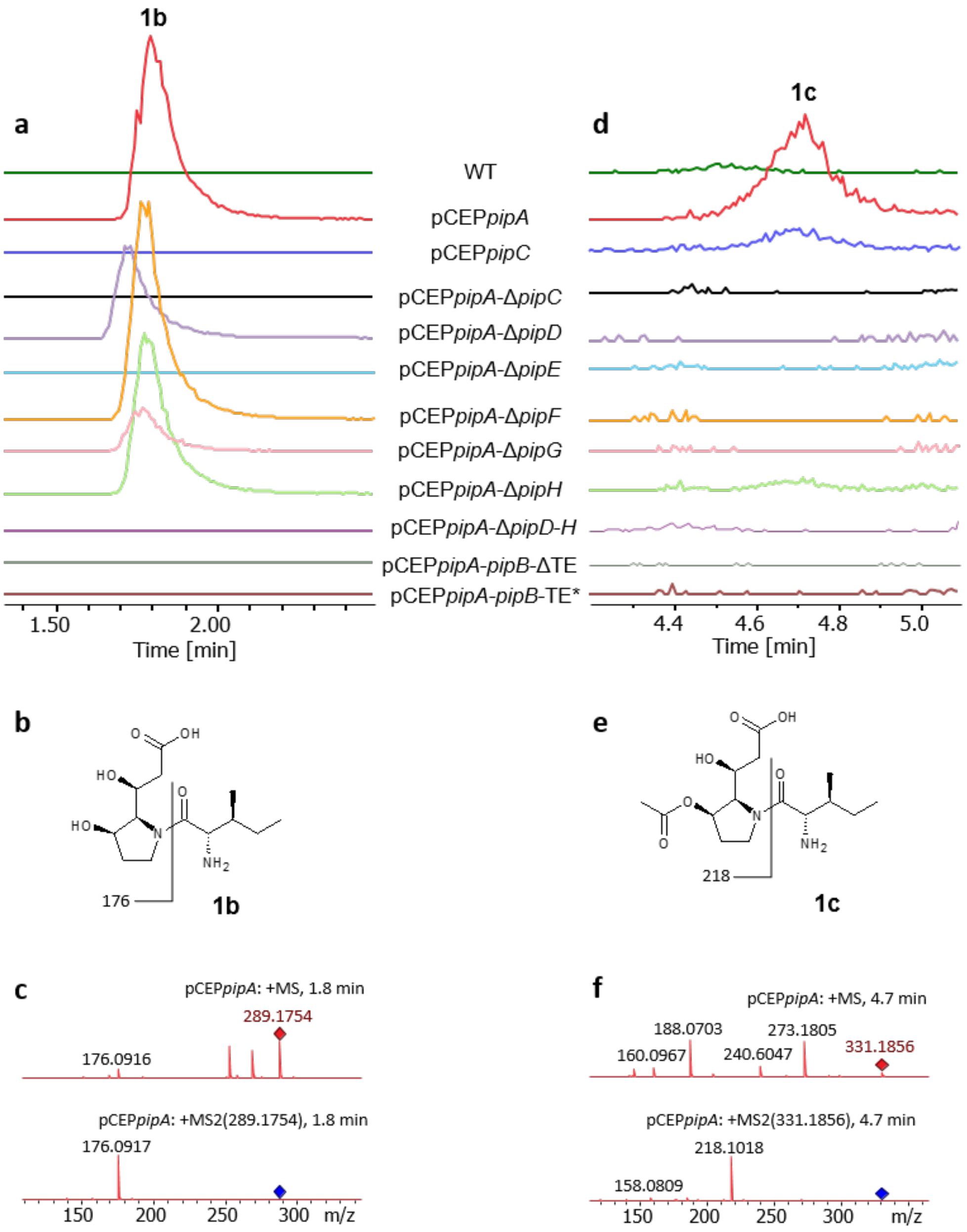


**Figure S2.2.** Extracted ion chromatograms (EICs) of compound **1b** after HPLC-MS analysis of selected culture extracts (**a**). Chemical structure of **1b** (**b**). MS$^2$ fragmentation pattern of **1b** (**c**). Extracted ion chromatograms (EICs) of compound **1c** after HPLC-MS analysis of culture extracts of selected mutants (**d**). Chemical structure of **1c** (**e**). MS$^2$ fragmentation pattern of **1c** (**f**).

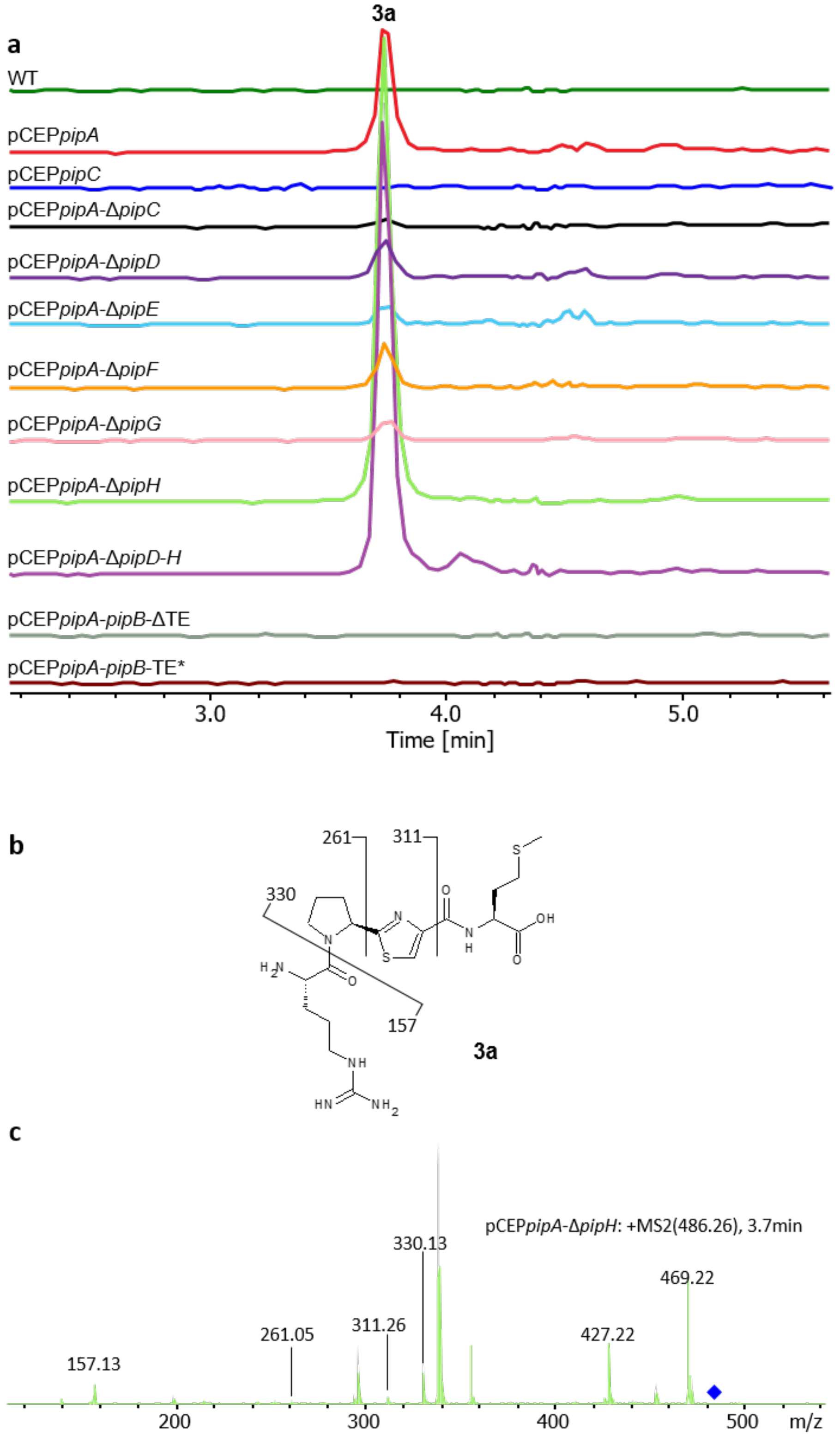


**Figure S2.3.** Extracted ion chromatograms (EICs) of compound **3a** after HPLC-MS analysis of selected culture extracts (**a**). Chemical structure of **3a** (**b**). $MS^2$ fragmentation pattern of **3a** (**c**).

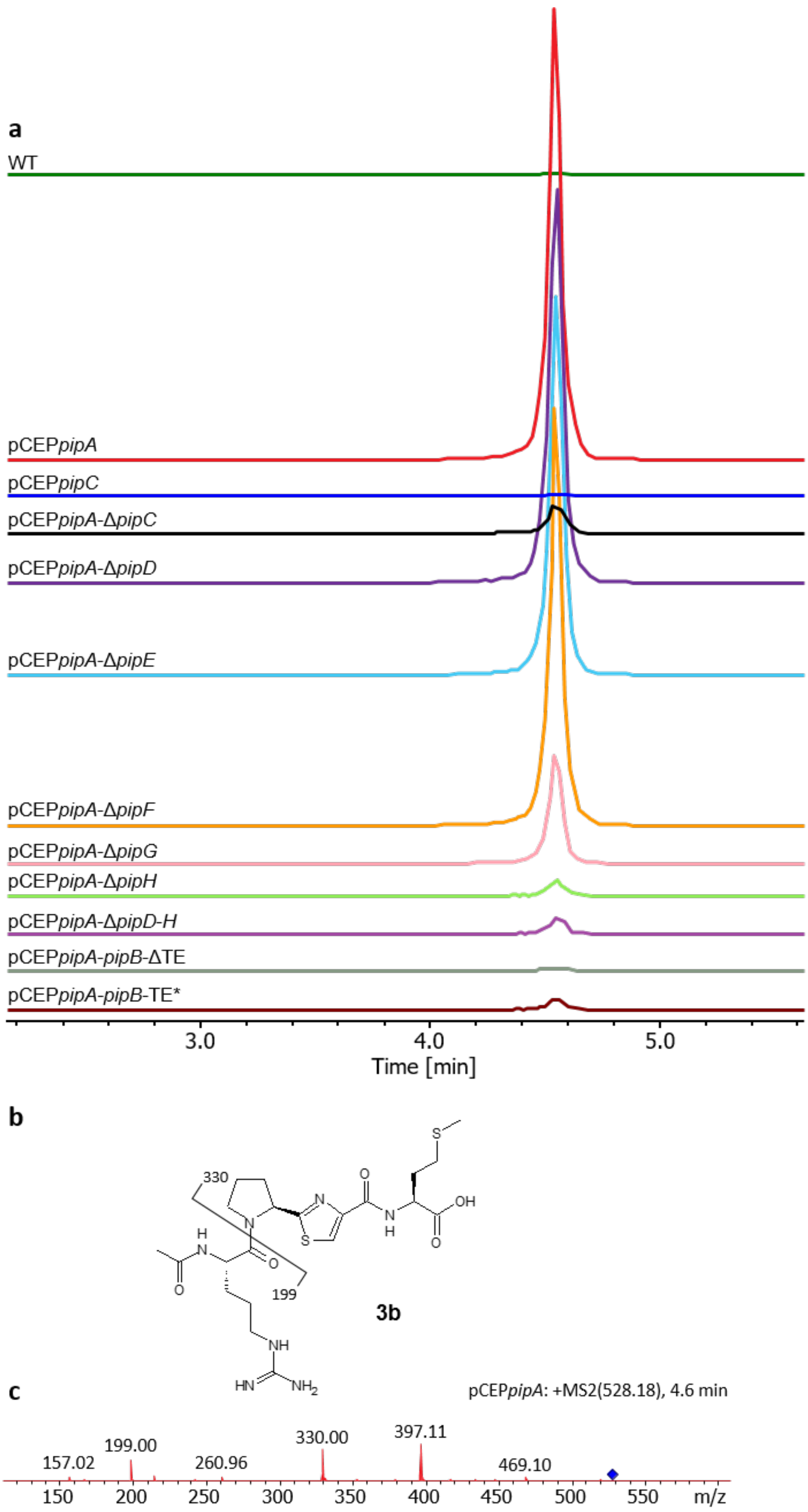


**Figure S2.4.** Extracted ion chromatograms (EICs) of compound **3b** after HPLC-MS analysis of selected culture extracts (**a**). Chemical structure of **3b** (**b**). $MS^2$ fragmentation pattern of **3b** (**c**). Dashed lines indicate a 10 times decreased signal.

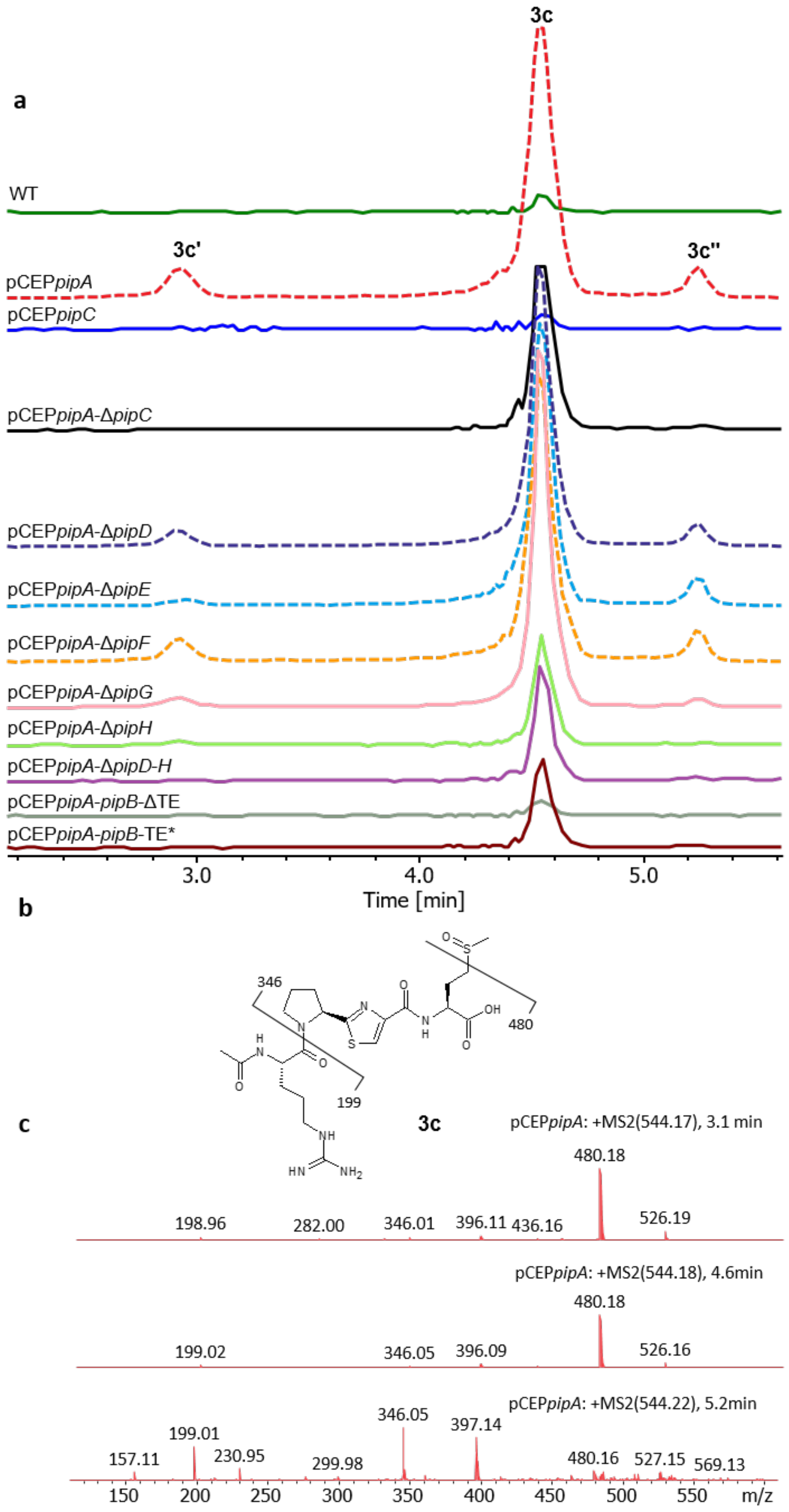


**Figure S2.5.** Extracted ion chromatograms (EICs) of compound **3c,** after HPLC-MS analysis of selected culture extracts (**a**). Chemical structure of **3c** (**b**). $MS^2$ fragmentation pattern of **3c** (**c**). Dashed lines indicate a 2x-decreased signal.

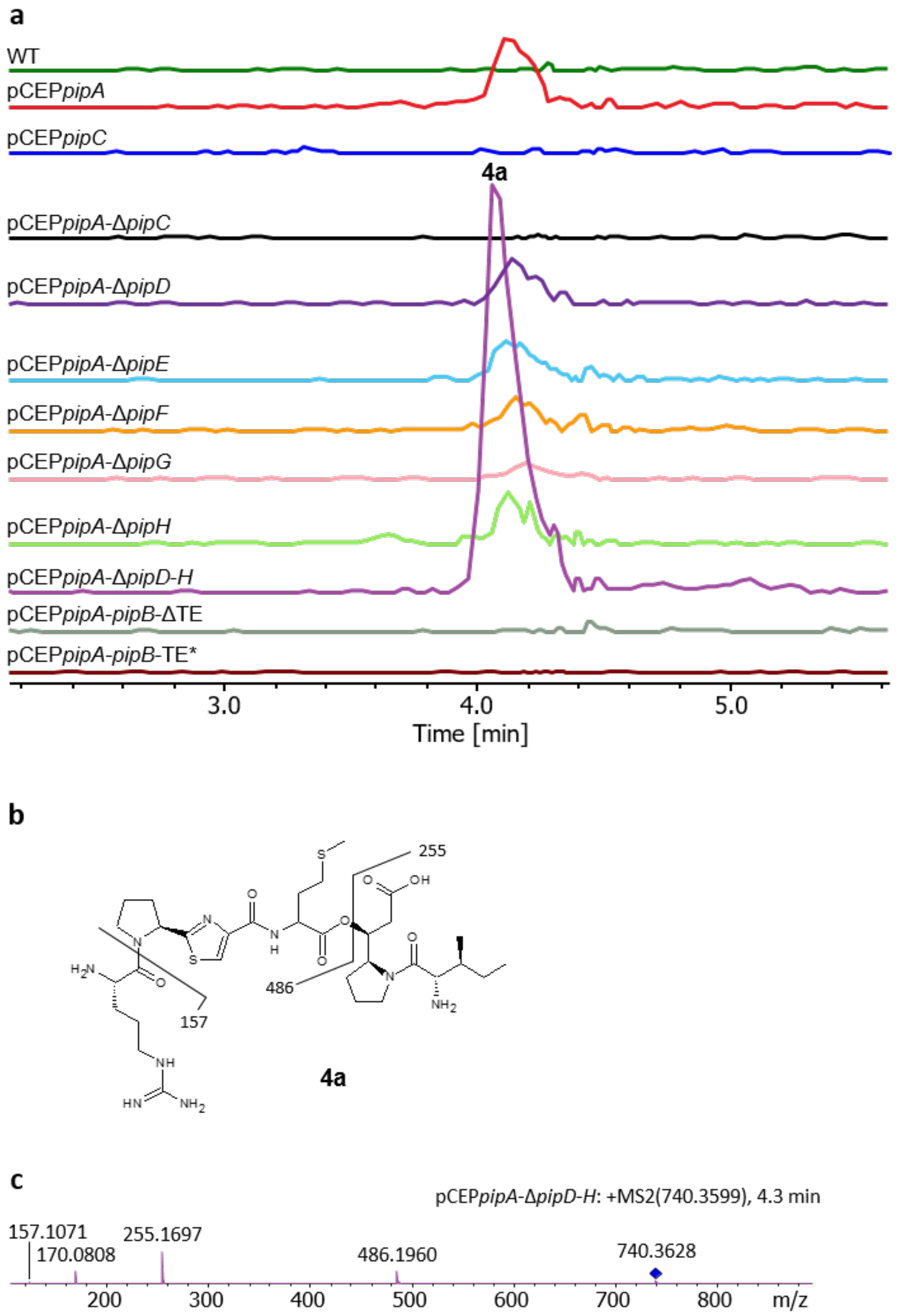


**Figure S2.6.** Extracted ion chromatograms (EICs) of compound **4a** after HPLC-MS analysis of selected culture extracts (**a**). Chemical structure of **4a** (**b**). $MS^2$ fragmentation pattern of **4a** (**c**).

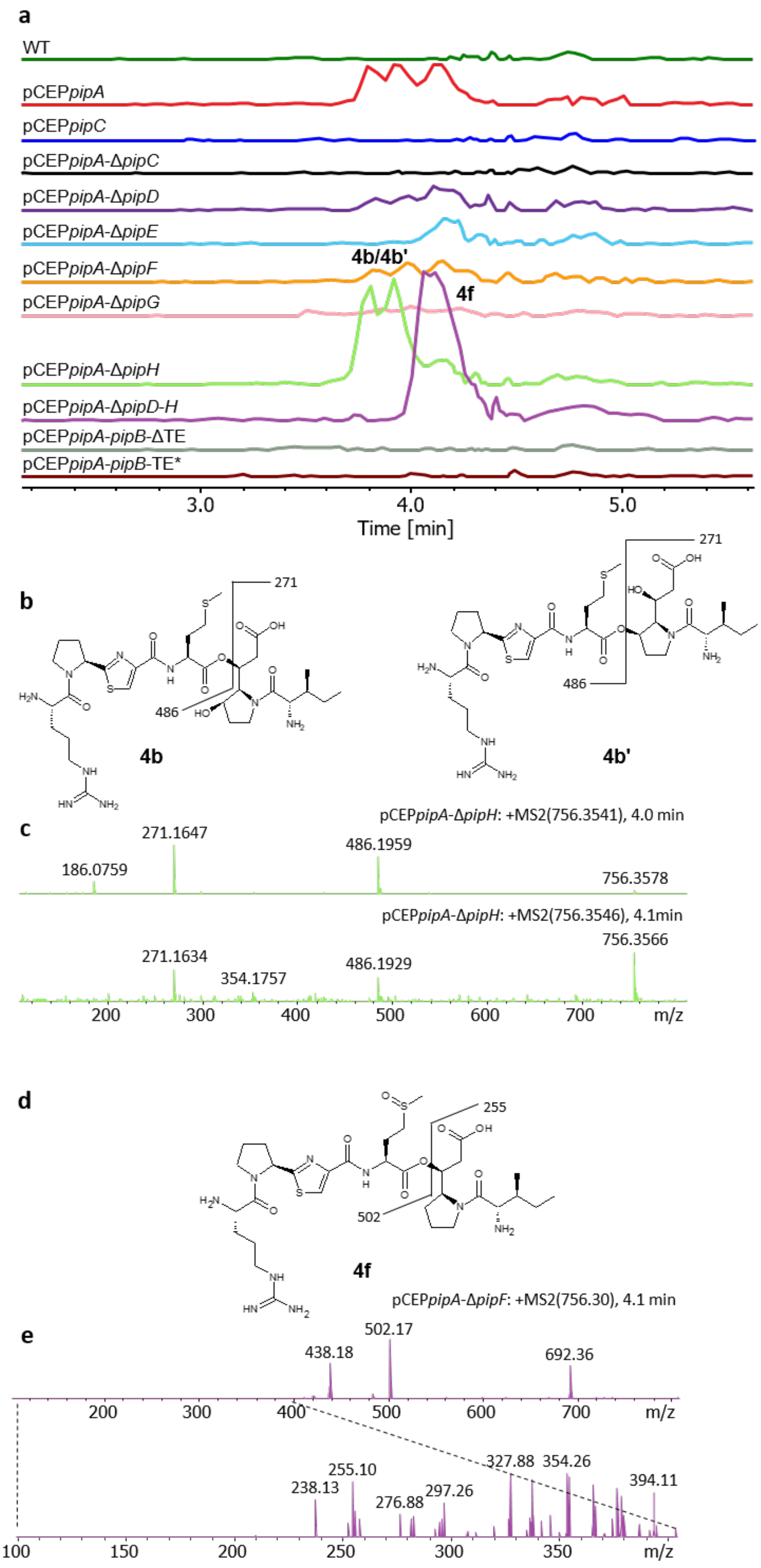


**Figure S2.7.** Extracted ion chromatograms (EICs) of compound **4b/4b'** and **4f** after HPLC-MS analysis of selected culture extracts (**a**). Chemical structure of **4b/4b'** (**b**). $MS^2$ fragmentation pattern of **4b/4b'** (**c**). Chemical structure of **4f** (**d**). $MS^2$ fragmentation pattern of **4f** (**e**).

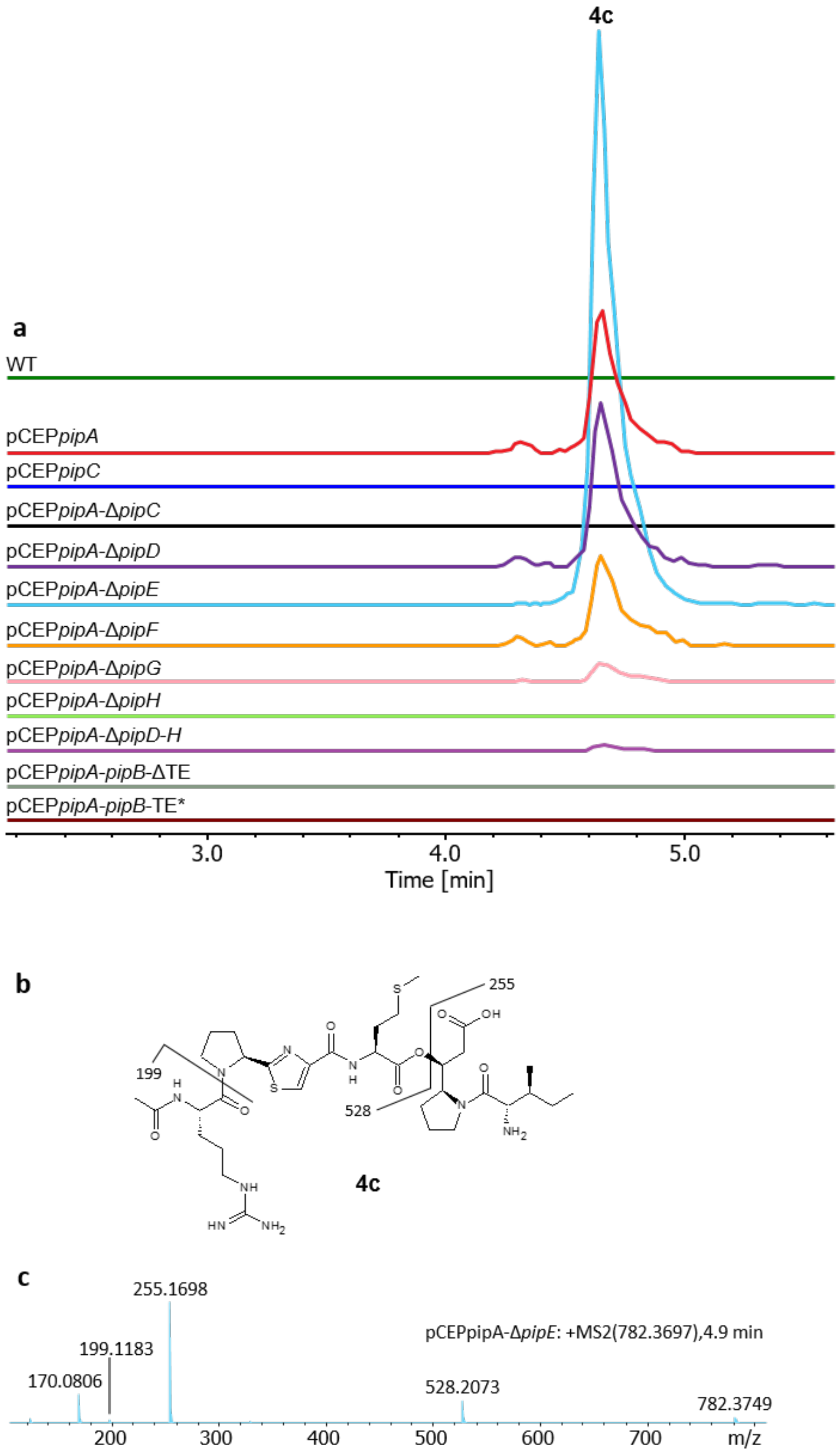


**Figure S2.8.** Extracted ion chromatograms (EICs) of compound **4c** after HPLC-MS analysis of selected culture extracts (**a**). Chemical structure of **4c** (**b**). $MS^2$ fragmentation pattern of **4c** (**c**).

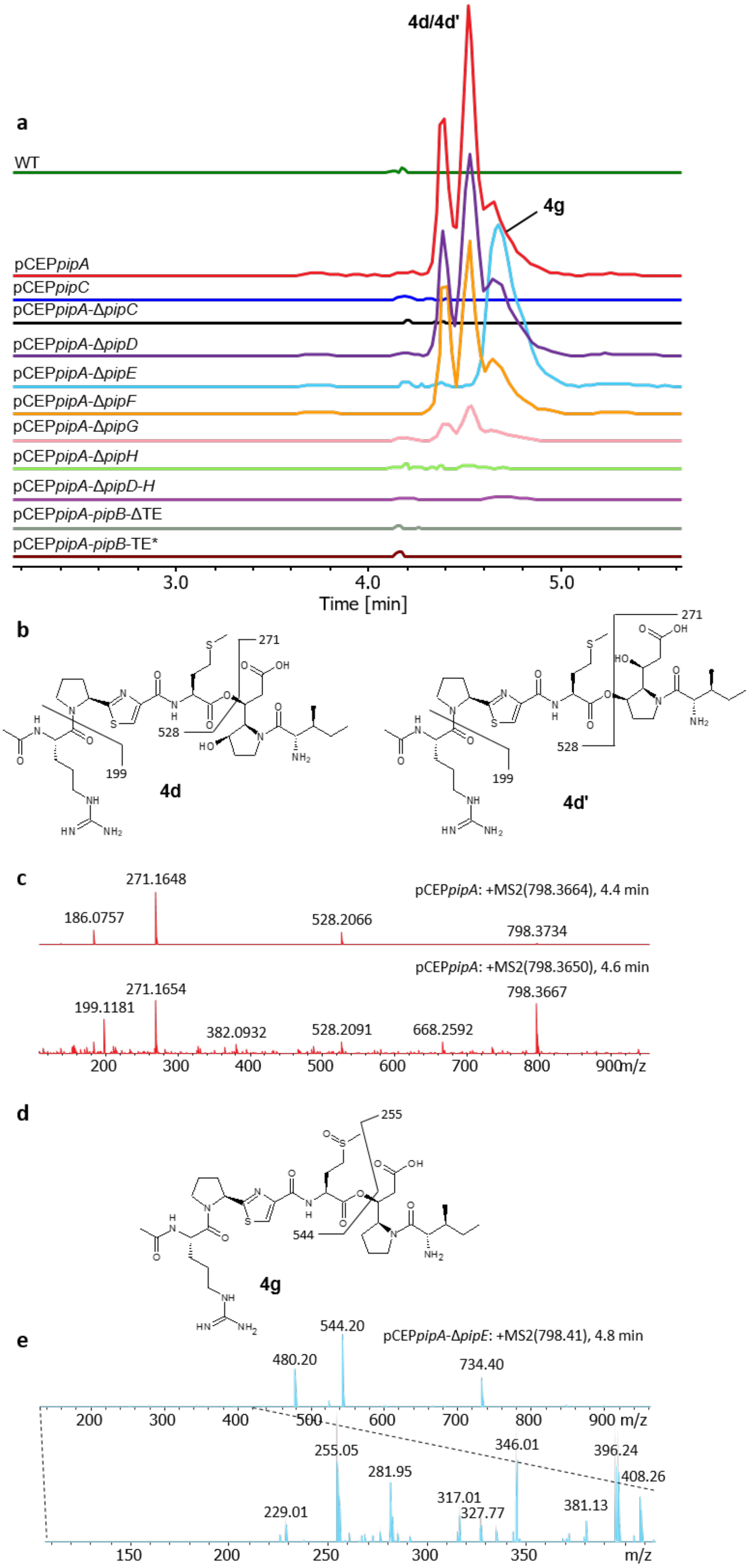


**Figure S2.9.** Extracted ion chromatograms (EICs) of compound **4d/4d'** and **4g** after HPLC-MS analysis of selected culture extracts (**a**). Chemical structure of **4d/4d'** (**b**). MS$^2$ fragmentation pattern of **4d/4d'** (**c**). Chemical structure of **4g** (**d**). MS$^2$ fragmentation pattern of **4g** (**e**).

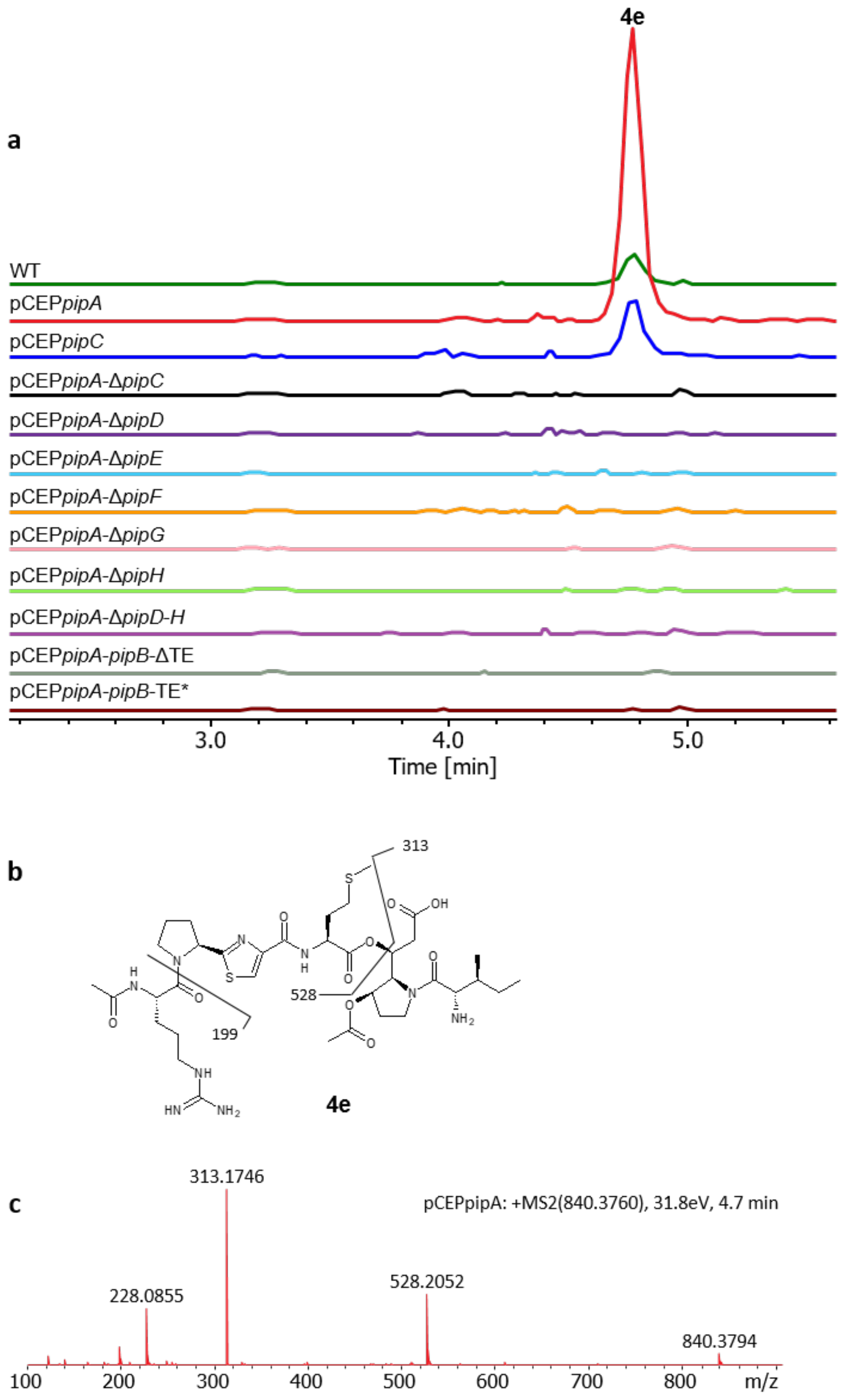


**Figure S2.10.** Extracted ion chromatograms (EICs) of compound **4e** after HPLC-MS analysis of selected culture extracts (**a**). Chemical structure of **4e** (**b**). $MS^2$ fragmentation pattern of **4e** (**c**).

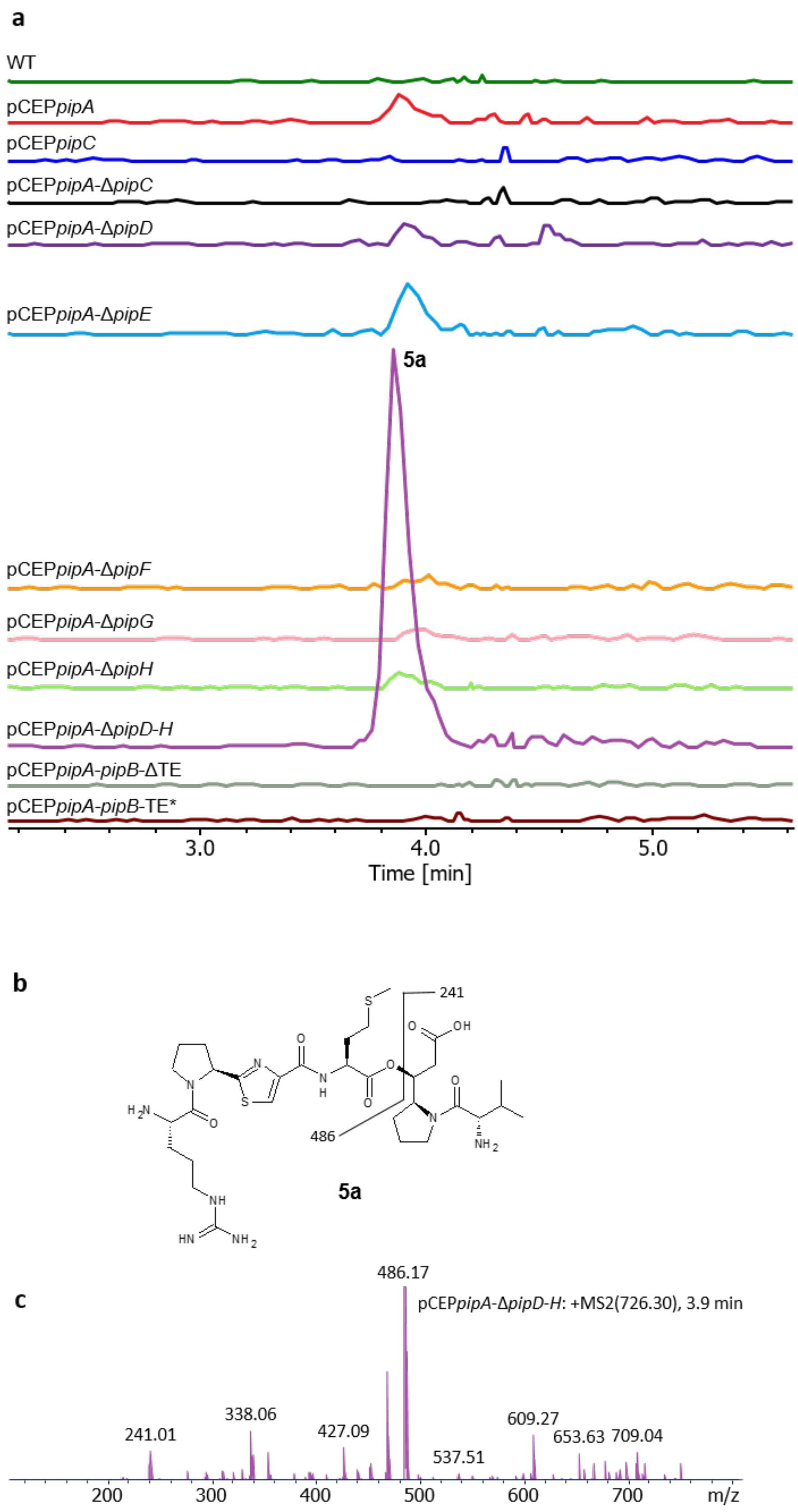


**Figure S2.11.** Extracted ion chromatograms (EICs) of compound **5a** after HPLC-MS analysis of selected culture extracts (**a**). Chemical structure of **5a** (**b**). $MS^2$ fragmentation pattern of **5a** (**c**).

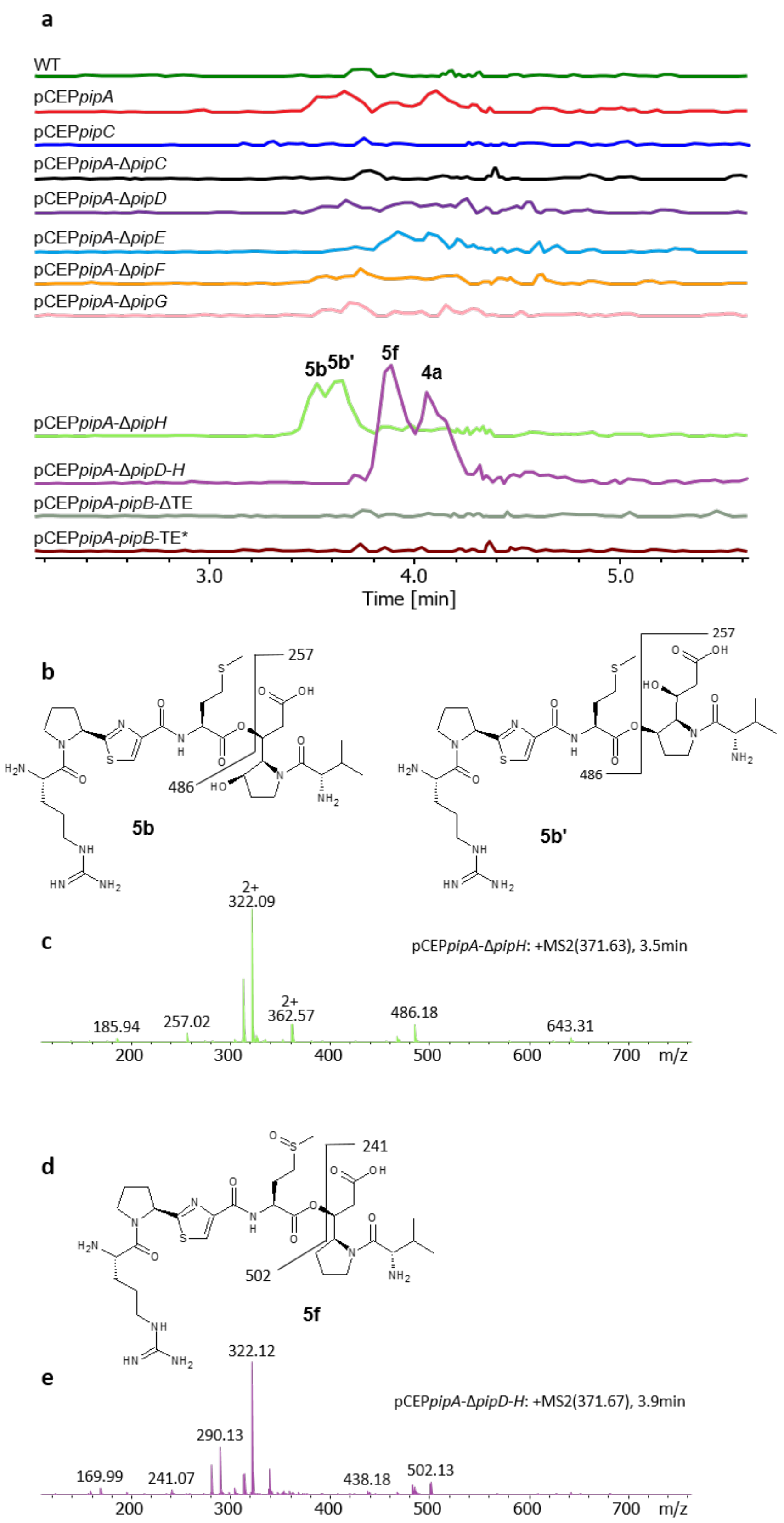


**Figure S2.12.** Extracted ion chromatograms (EICs) of compound **5b**/**5b'** and **5f** after HPLC-MS analysis of selected culture extracts (**a**). Chemical structure of **5b** and **5b'** (**b**). $MS^2$ fragmentation pattern of **5b/5b'** (**c**). Chemical structure of **5f** (**d**). $MS^2$ fragmentation pattern of **5f** (**e**).

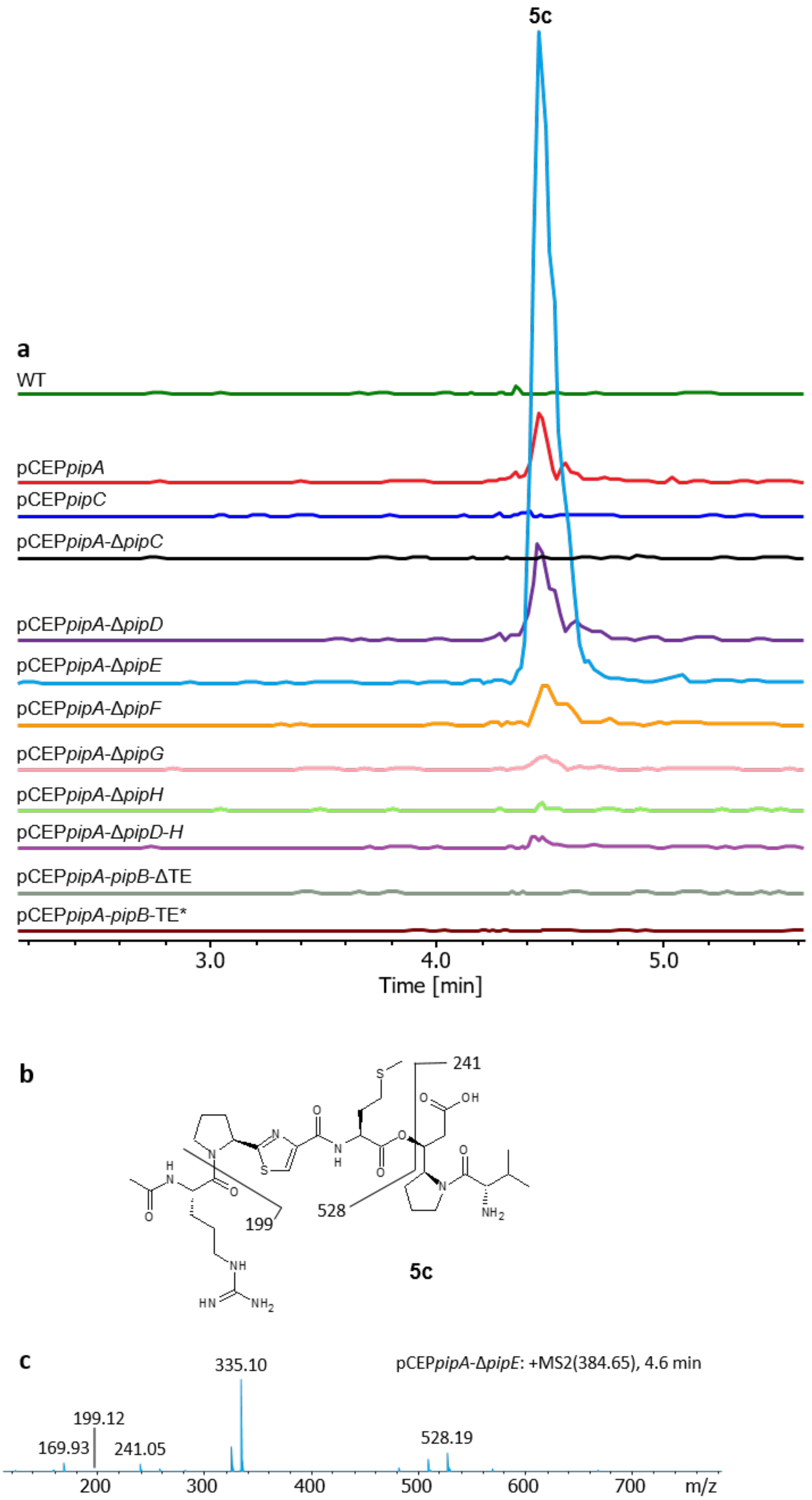


**Figure S2.13.** Extracted ion chromatograms (EICs) of compound **5c** after HPLC-MS analysis of selected culture extracts (**a**). Chemical structure of **5c** (**b**). $MS^2$ fragmentation pattern of **5c** (**c**).

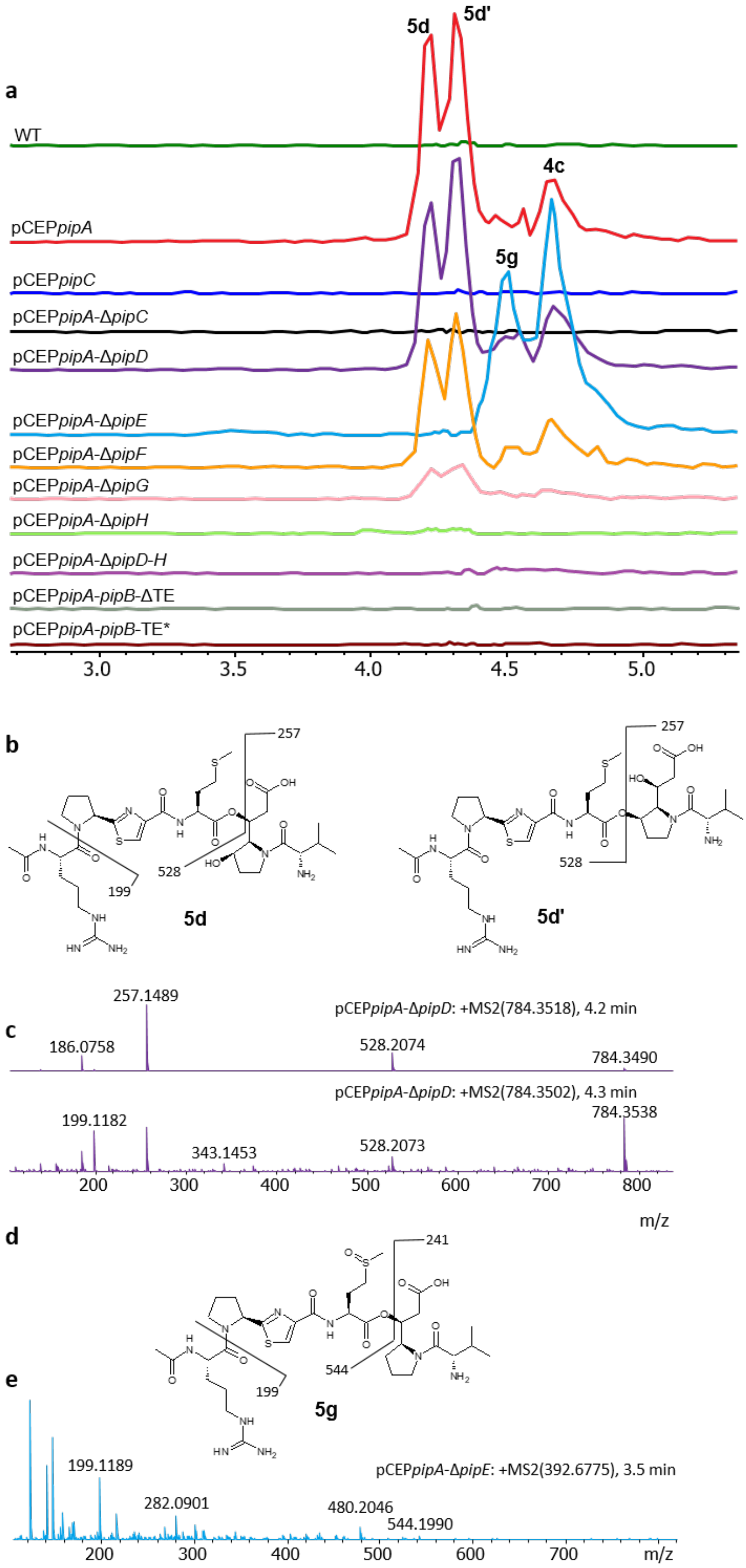


**Figure S2.14.** Extracted ion chromatograms (EICs) of compound **5d**/**5d'** and **5g** after HPLC-MS analysis of selected culture extracts (**a**). Chemical structure of **5d/5d'** (**b**). MS$^2$ fragmentation pattern of **5d/5d'** (**c**). Chemical structure of **5g** (**d**). MS$^2$ fragmentation pattern of **5g** (**e**).

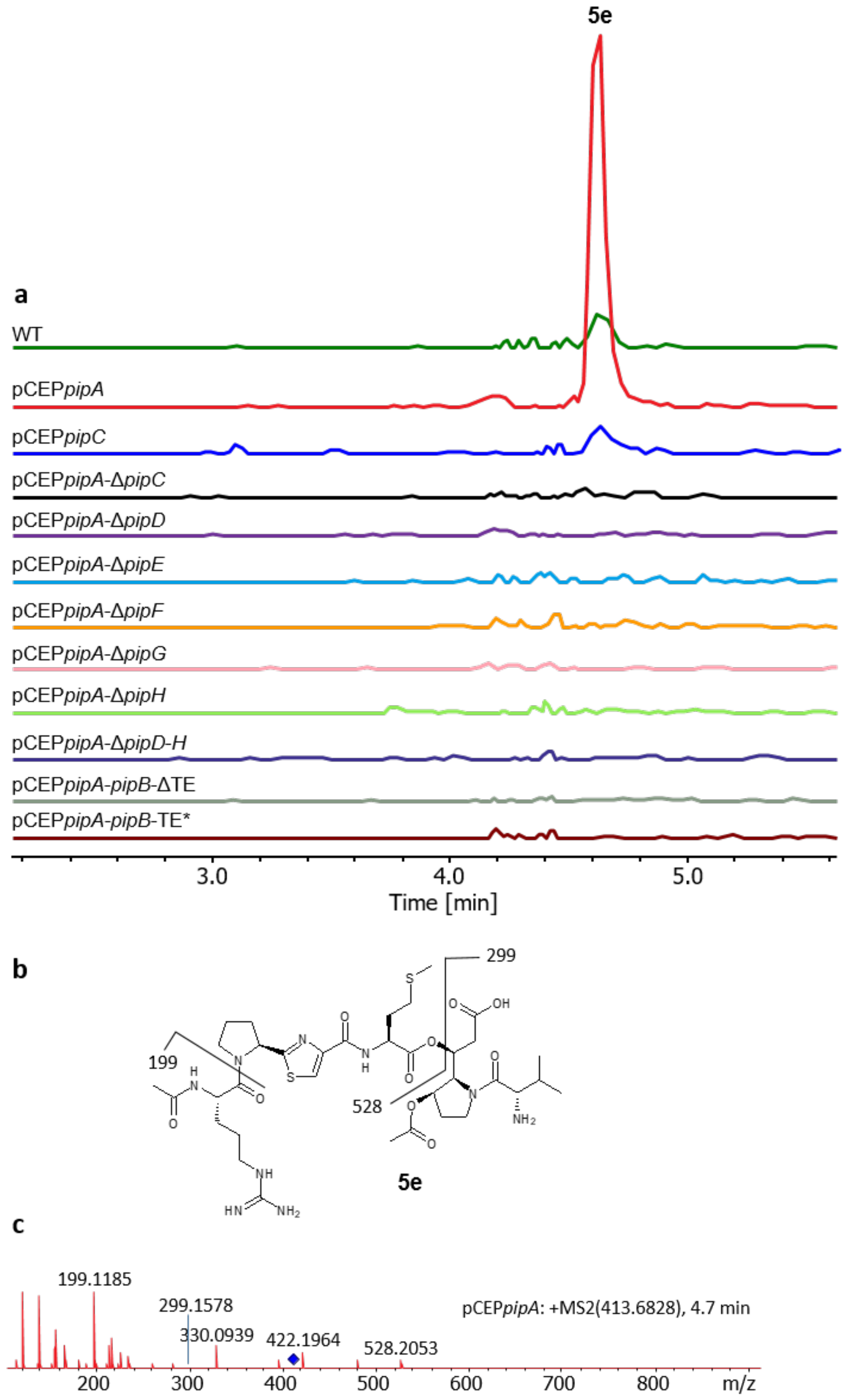


**Figure S2.15.** Extracted ion chromatograms (EICs) of compound **5e** after HPLC-MS analysis of selected culture extracts (**a**). Chemical structure of **5e** (**b**). MS$^2$ fragmentation pattern of **5e** (**c**).

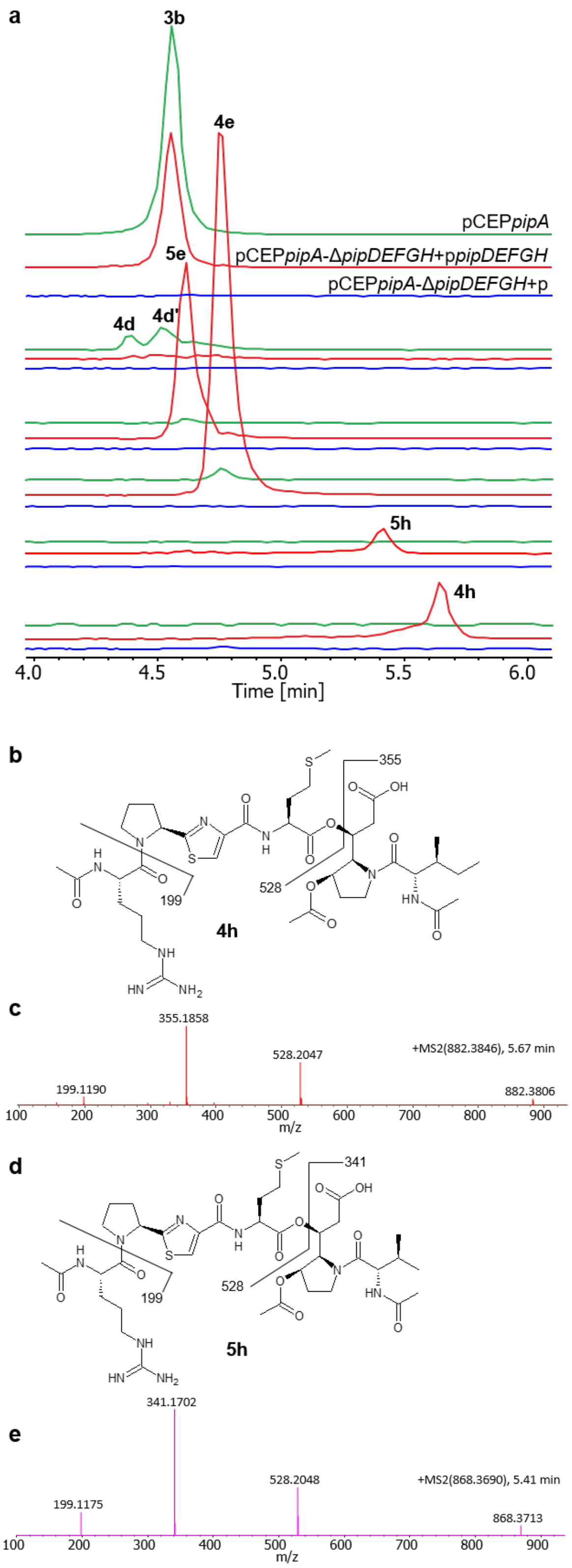


**Figure S2.16.** EICs of selected compounds produced in pCEP*pipA* (green), pCEP*pipA*-Δ*pipDEFGH*+p (blue) and pCEP*pipA*- Δ*pipDEFGH*+p*pipDEFGH* (red) (**a**). Chemical structure of **4h** (**b**) MS$^2$ fragmentation pattern of **4h** (**c**). Chemical structure of **5h** (**d**) MS$^2$ fragmentation pattern of **5h** (**e**).

### Isolation and structure elucidation

#### General experimental procedures

$^{1}H$, $^{13}C$, HSQC, HMBC, $^{1}H$-$^{1}H$ COSY, and ROESY spectra were measured on a Bruker AV600 spectrometer, using DMSO as solvent. Coupling constants are expressed in Hz and chemical shifts are given on a ppm scale. HRESIMS was performed on an UltiMate 3000 system (Thermo Fisher) coupled to an Impact II qTof mass spectrometer (Bruker Daltonik GmbH). Preparative HPLC was performed on an Agilent 1260 liquid chromatograph with a ZORBRAX StableBond 300 C18 (21.2 mm × 250 mm, 7.0μm, Agilent). Semi-preparative HPLC was performed on an Agilent 1260 liquid chromatograph with a ZORBRAX StableBond 300 C18 (9.4 mm × 250 mm, 5.0μm, Agilent).

#### Isolation and purification of 1a, 4c and 4e

For isolation of **1a** *P. entomophila*-pCEP*pipC* was cultivated in 6 x 1 L XPP medium containing 4% Amberlite® XAD-16 adsorber resin, induced with 0.2 % L-arabinose. The production cultures were incubated for 48 h at 30 °C shaking at 130 rpm. Subsequently the supernatant was separated from the resin by filtration. Extraction of the compound from the resin was performed by washing the resin with 6 x 500 ml methanol for 45 min. Via a sephadex column, the desired compound was separated from the crude extract and further purified via a semi preparative HPLC-system on a C18 column yielding 22 mg of **1a**.

For the isolation of **4c** the XAD-16 extract of *P. entomophila*Δ*hfq*-pCEP*pipA* (3.8 g) was directly subjected to preparative HPLC (ACN–$H_2O$, from 5% to 40% in 20 min, v/v) to obtain five fractions 1-5. Fraction 2 (163 mg) was purified by using semi-preparative HPLC (ACN–$H_2O$, 12:88, v/v) to yield compound **4c** (70.0 mg). Both of the ACN and $H_2O$ used in HPLC contained 0.1% formic acid. Similarily, **4e** was isolated from an XAD-16 extract of *P. entomophila* pCEP*pipA*-Δ*pipDEFGH*+p*pipDEFGH* (Fig. S2.16) yielding 12 mg of **4e**.

### Structure elucidation of 1a

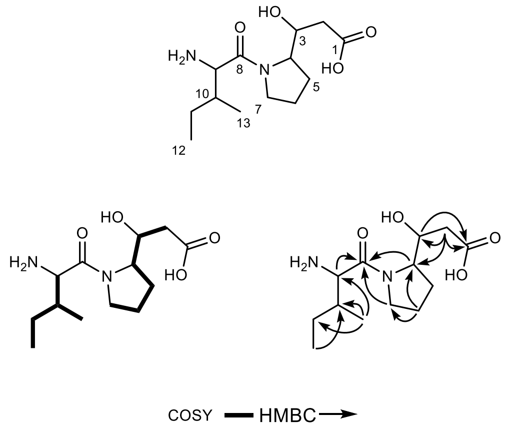


| no. | 1a | |
|---|---|---|
| | $\delta_H$ (mult., *J* in Hz) | $\delta_C$, type |
| 1 | | 174.2, C |
| 2 | 2.11 (brs) | 38.8, $CH_2$ |
| 3 | 4.19 (m) | 68.7, CH |
| 4 | 4.18 (m) | 61.1, CH |
| 5 | 1.89 (m) | 24.8, $CH_2$ |
| 6 | 1.81 (brs) | 25.6, $CH_2$ |
| 7 | 3.67 (dd, 14.0, 9.2)<br>3.38 (dd, 14.0, 6.4) | 47.9, $CH_2$ |
| 8 | | 171.6, C |
| 9 | 3.67 (overlap) | 56.5, CH |
| 10 | 1.64 (brs) | 37.5, CH |
| 11 | 1.46 (m)<br>1.10 (m) | 23.7, $CH_2$ |
| 12 | 0.83 (t, 7.3) | 11.8, $CH_3$ |
| 13 | 0.92 (d, 6.5) | 15.8, $CH_3$ |

**Figure S2.17.** NMR data for **1a**. $^1$H-$^1$H COSY- and HMBC-correlations (left). $^1$H (500 MHz) und $^{13}$C (125 MHz) NMR data of **1a** in DMSO-$d_6$ ($\delta$ in ppm und *J* in Hz) (right). Compound **1a** had a molecular formula of $C_{13}H_{24}N_2O_4$, as deduced from HR-ESI-MS at *m/z* 273.1805 $[M+H]^+$ (calcd for $C_{13}H_{25}N_2O_4$, 273.1809). indicating three degrees of unsaturation. The $^1$H and $^{13}$C NMR (Fig. S2.18 & S2.19) together with HSQC spectrum (Fig. S2.21) revealed the existence of two methyls ($\delta_C$ 11.8, 15.8), two methylenes ($\delta_C$ 23.7, 24.8, 25.6, 47.9 and 38.8), four sp$^3$ methines ($\delta_C$ 68.7, 61.1, 56.5 and 37.5), and two quaternary carbons ($\delta_C$ 174.2 and 171.6). The $^1$H-$^1$H COSY spectrum (Fig. S2.20) exhibits the correlations of H-2/H-3/H-4/H-5/H-6/H-7, of H-9/H-10/H-11/H-12 and of H-10/H-13, together with the observed HMBC correlations (Fig. S2.21) from H-2/H-3 to C-1, from H-4/H-7 to C-8, and from H-9 to C-8. It was noted that compound **1a** showed peak splitting from some protons and carbons in its NMR spectra. This phenomenon is caused by the C-N bond rotation in the solution which frequently occurs in compounds with an amide group. Thus, the plane structure of compound **1a** was established.

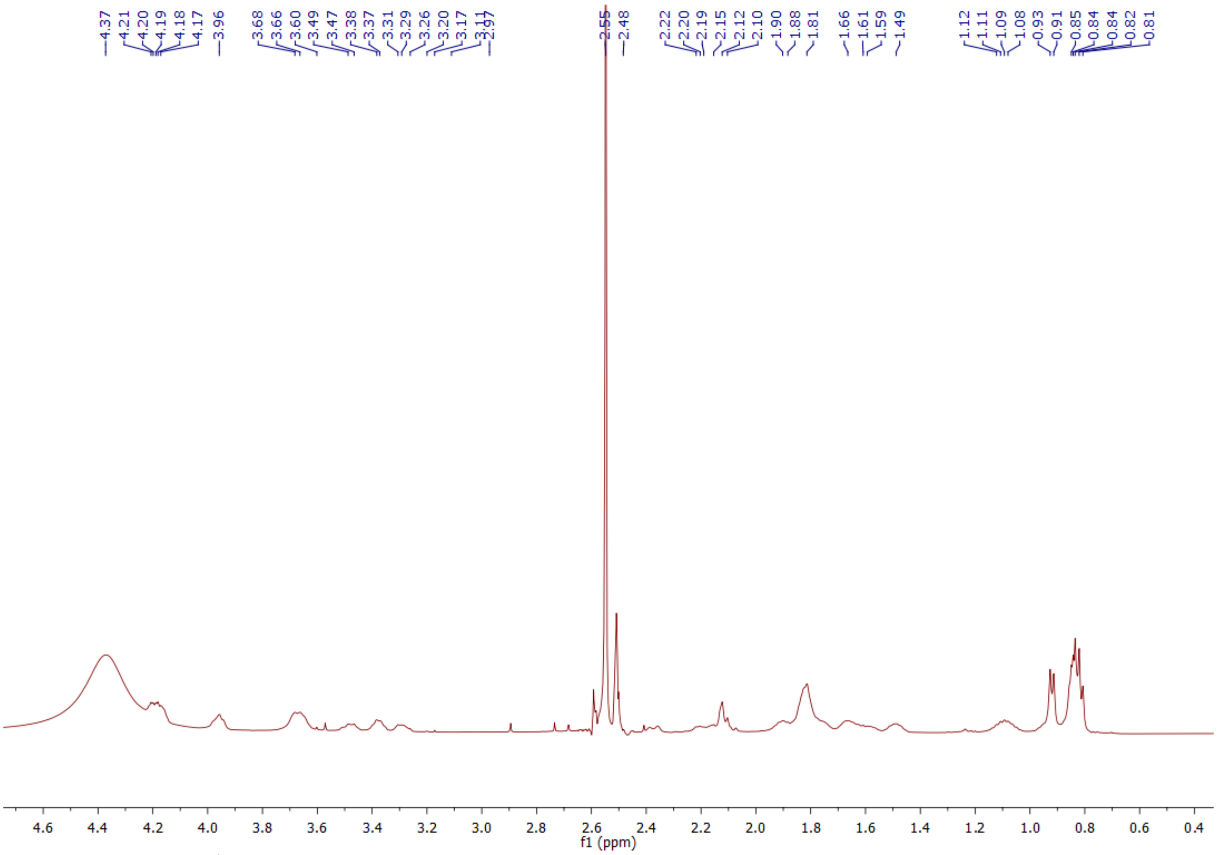


**Figure S2.18.** $^{1}$H NMR (600 MHz) spectrum of **1a**.

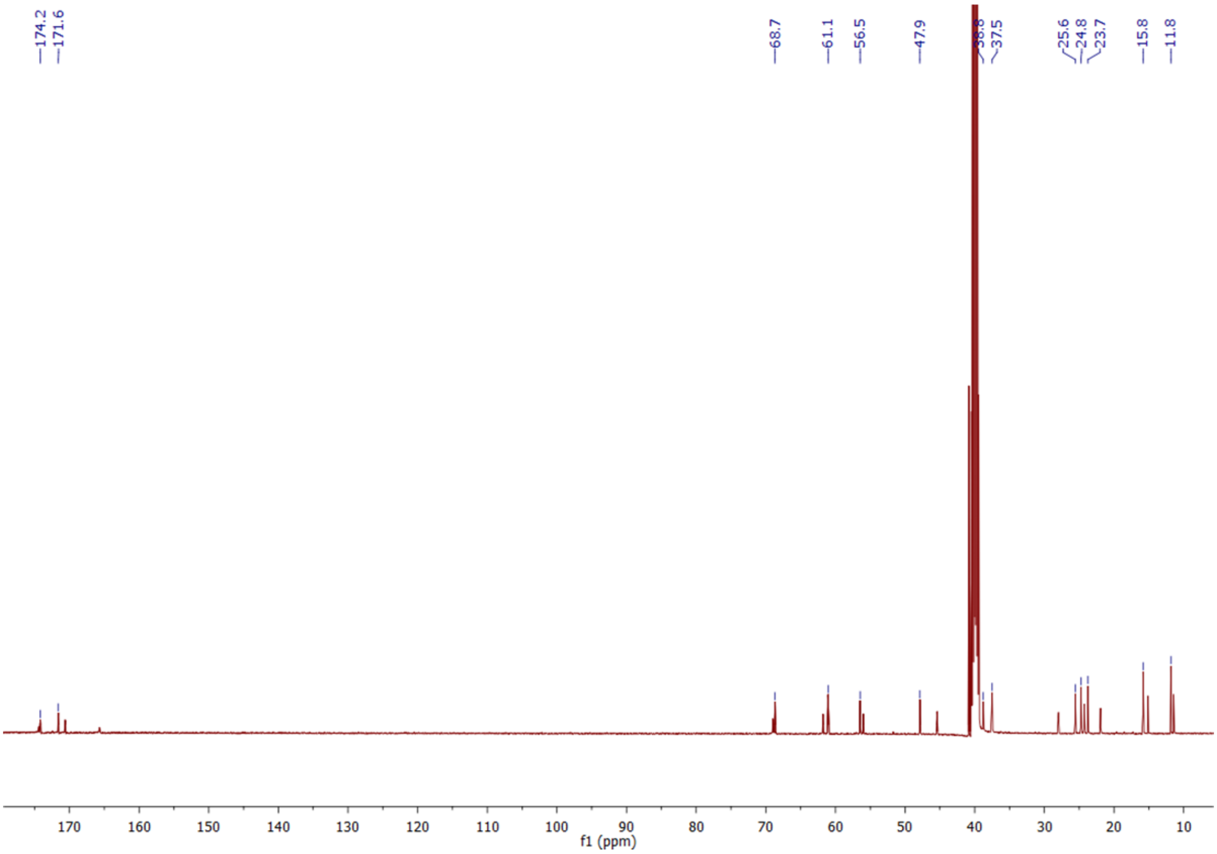


**Figure S2.19.** $^{13}$C NMR (150 MHz) spectrum of **1a**.

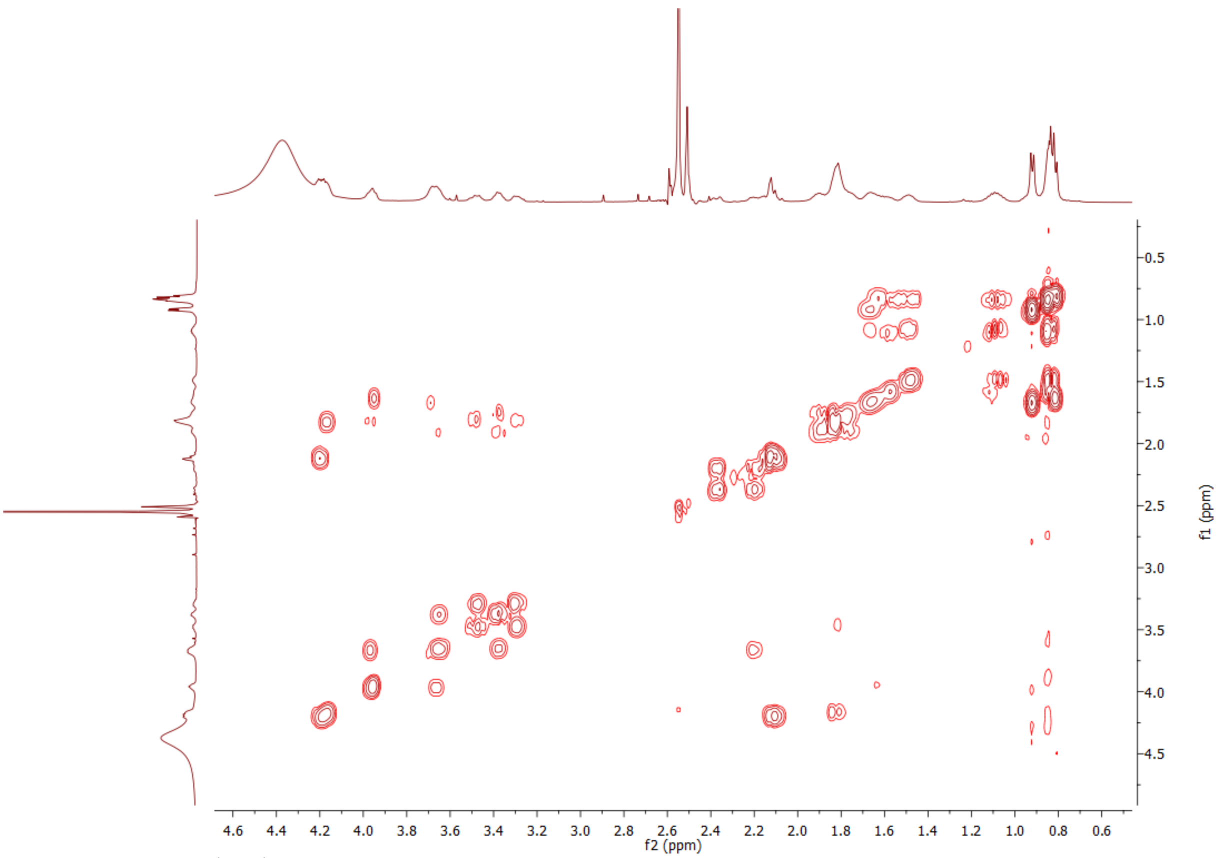


**Figure S2.20.** $^1H$-$^1H$ COSY NMR spectrum of **1a**.

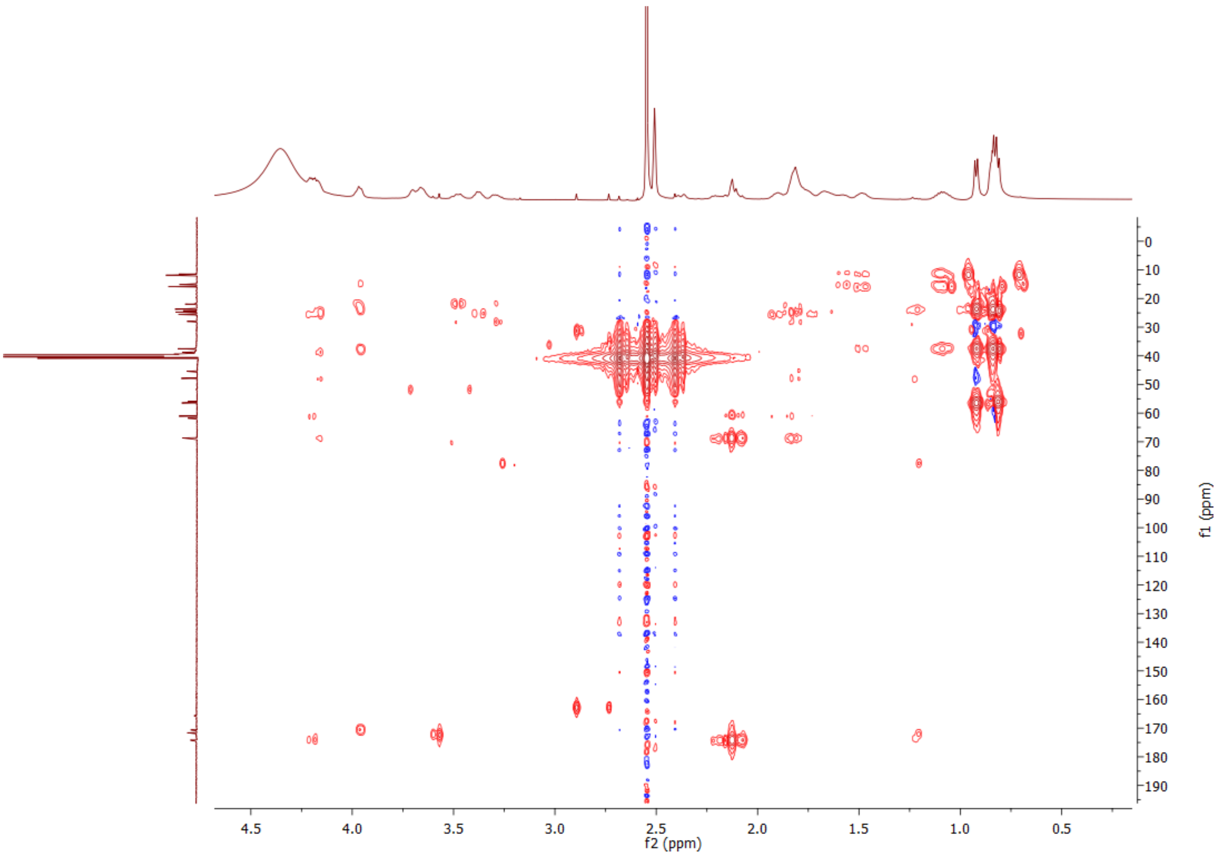


**Figure S2.21.** HMBC NMR spectrum of **1a**.

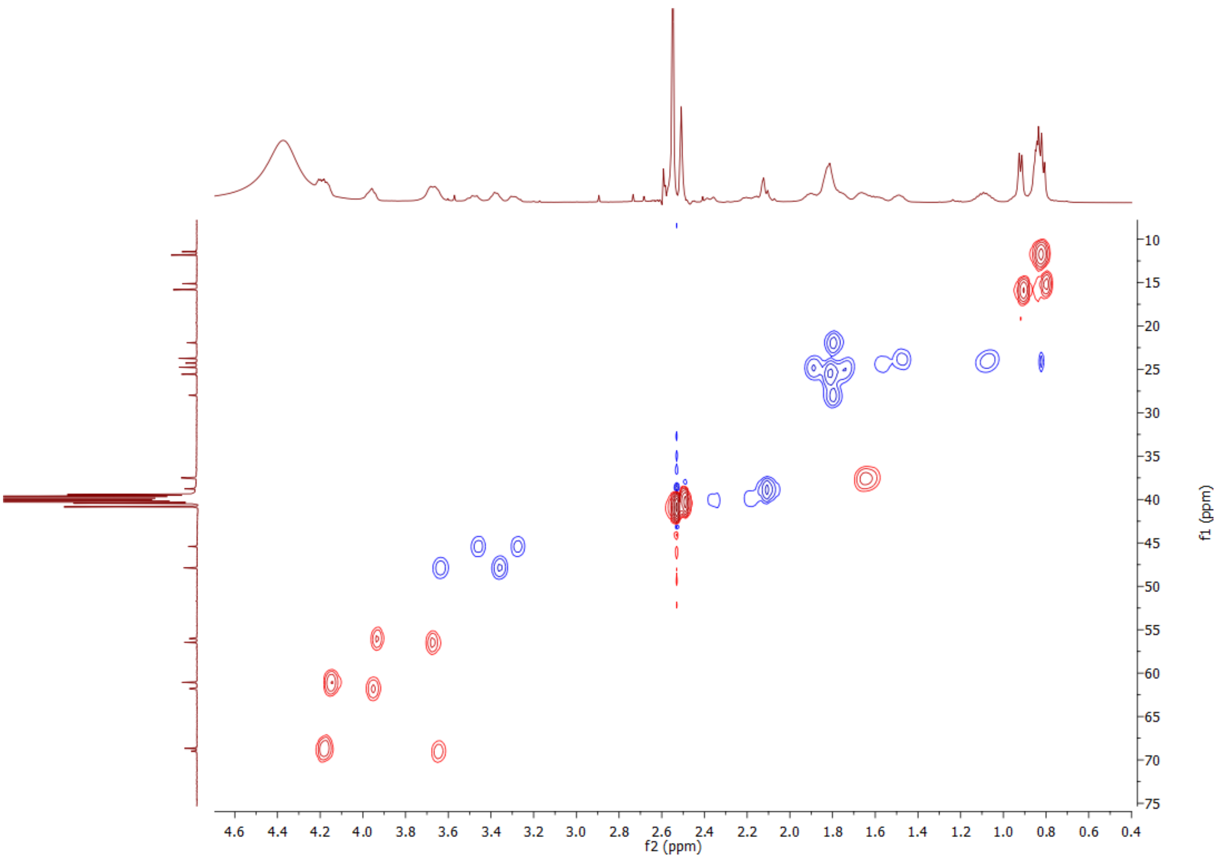


**Figure S2.21.** HSQC NMR spectrum of **1a**.

### Structure elucidation of 4c

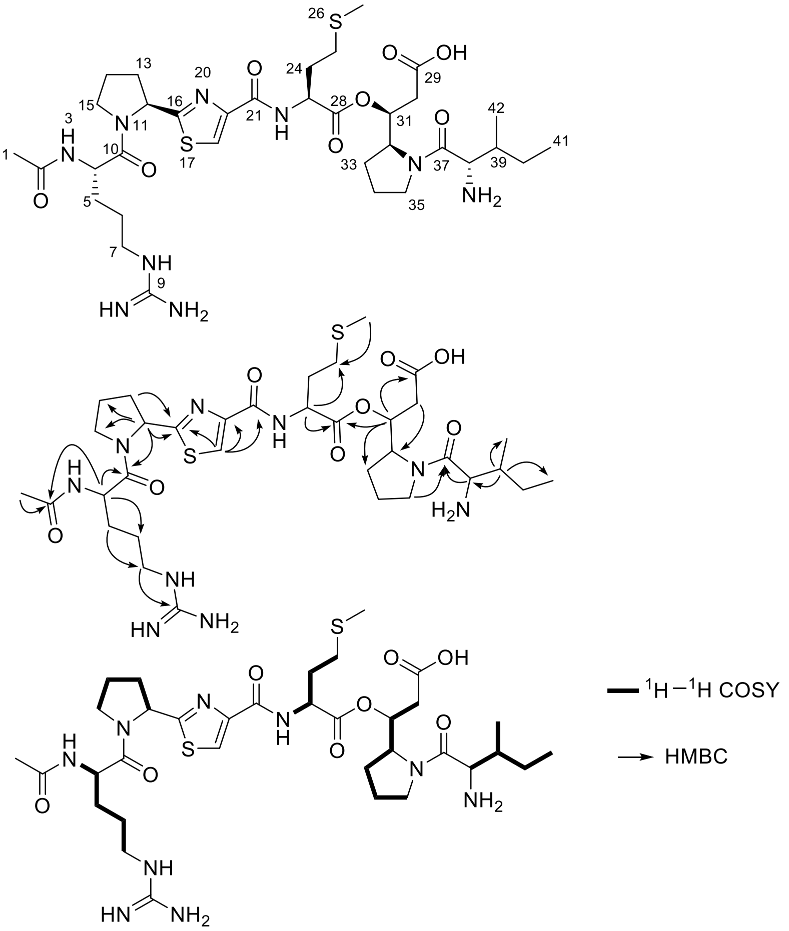


**Figure S2.22.** Key $^1H$-$^1H$ COSY and HMBC correlations of compound **4c**. Compound **4c** has a molecular formula of $C_{34}H_{55}N_9O_8S_2$, as determined by HR-ESI-MS (found, *m/z* 782.3675 $[M+H]^+$; calcd for $C_{34}H_{56}N_9O_8S_2$, *m/z* 782.3688 $[M+H]^+$; Δppm 1.3). The $^1H$ and $^{13}C$ NMR data (Table 4) together with HSQC data revealed the existence of four methyl groups, 13 methylenes, seven $sp^3$ methines, one olefinic methine, two olefinic quaternary carbon, six carbonyls and one of –NHCONH$_2$. The structure was established by $^1H$-$^1H$ COSY correlations of H-3/H-4/H-5/H-6/H-7, H-12/H-13/H-14/H-15, H-22/H-23/H-24/H-25, H-30/H-31/H-32/H-33/H-34/H-35, H-38/H-39/H-40/H-41 and H-39/H-42 in combination with HMBC correlations from H-1 to C-2, H-4 to C-2/C-6/C-10, H-7 to C-9, H-12 to C-10/C-14/C-15/C-16, H-13 to C-16, H-18 to C-16/C-19/C-21, H-23 to C-25/C-28, H-27 to C-25, H-31 to C-28/C-29/C-33, H-31 to C-29, H-35/H-38 to C-37. Thus, the structure of compound **4c** was determined to be that shown in Fig. 3.

**Table S2.1.** $^{1}H$ (600 MHz) and $^{13}C$ (150 MHz) NMR spectroscopic data in DMSO-$d_6$ ($\delta$ in ppm and $J$ in Hz).

| No | $\delta_H$ (mult., J) | $\delta_C$ |
|---|---|---|
| 1 | 1.85 (s) | 22.2 |
| 2 | | 169.3 |
| 3 | 8.37 (overlap) | |
| 4 | 4.51 (m) | 50.2 |
| 5 | 1.76 (m) | 28.1 |
| | 1.57 (m) | |
| 6 | 1.56 (m) | 24.9 |
| 7 | 3.07 (m) | 40.4 |
| 9 | | 157.4 |
| 10 | | 171.2 |
| 12 | 5.34 (dd, 8.0, 1.6) | 58.2 |
| 13 | 2.25 (m) | 31.5 |
| | 2.16 (m) | |
| 14 | 1.83 (overlap) | 24.0 |
| | 1.97 (overlap) | |
| 15 | 3.81 (m) | 46.7 |
| | 3.76 (m) | |
| 16 | | 173.6 |
| 18 | 8.19 (s) | 124.1 |
| 19 | | 148.8 |
| 21 | | 160.6 |
| 22 | 8.53 (d, 7.9) | |
| 23 | 4.60 (m) | 51.3 |
| 24 | 2.08 (m) | 30.1 |
| 25 | 2.47 (m) | 29.8 |
| | 2.51 (m) | |
| 27 | 2.04 (s) | 14.5 |
| 28 | | 170.3 |
| 29 | | 172.6 |
| 30 | 2.30 (m) | 36.1 |
| 31 | 5.49 (m) | 73.2 |
| 32 | 4.41 (m) | 57.2 |
| 33 | 1.73 (m) | 25.6 |
| | 1.82 (m) | |
| 34 | 1.71 (overlap) | 24.1 |
| | 2.04 (overlap) | |
| 35 | 3.29 (m) | 47.4 |
| | 3.67 (m) | |
| 37 | | 170.9 |
| 38 | 3.73 (m) | 55.9 |
| 39 | 1.64 (m) | 36.9 |
| 40 | 1.03 (m) | 23.3 |
| | 1.43 (m) | |
| 41 | 0.78 (t, 7.3) | 11.3 |
| 42 | 0.86 (d, 6.7) | 15.3 |

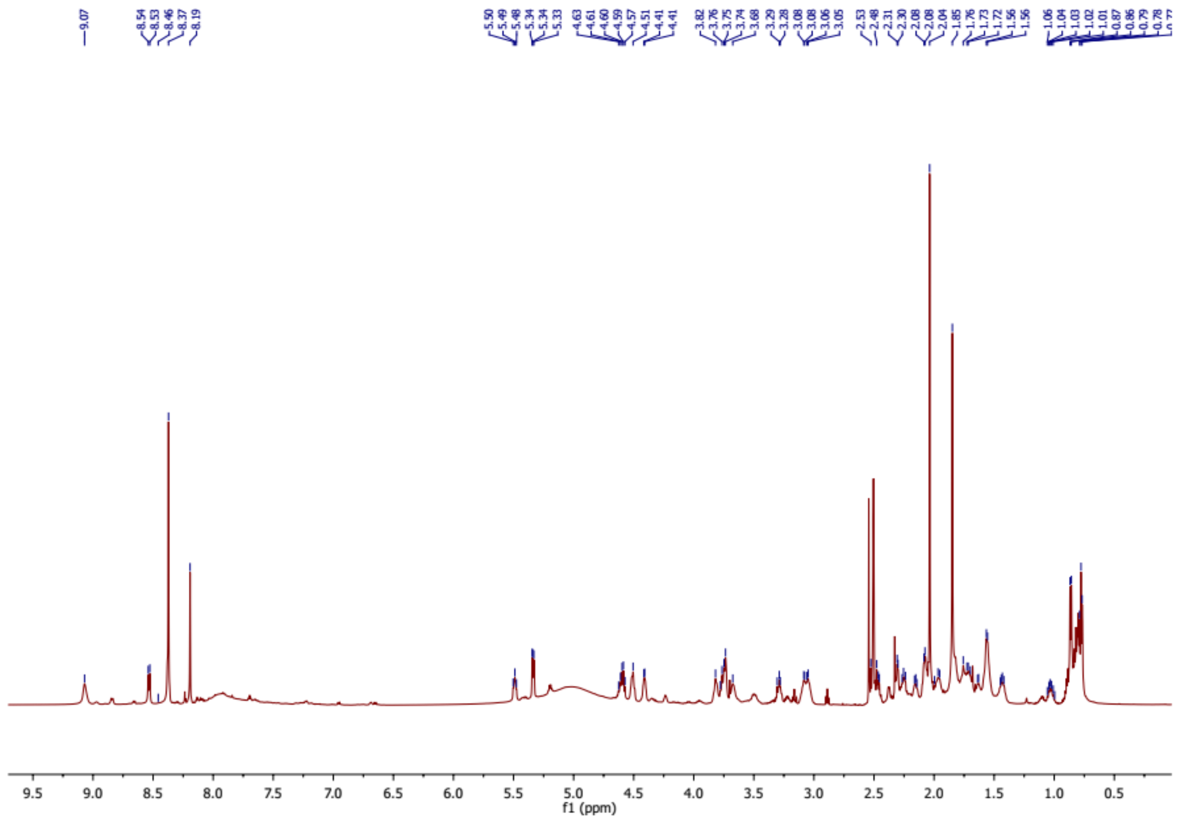


**Figure S2.23.** $^{1}$H NMR (600 MHz) spectrum of **4c**.

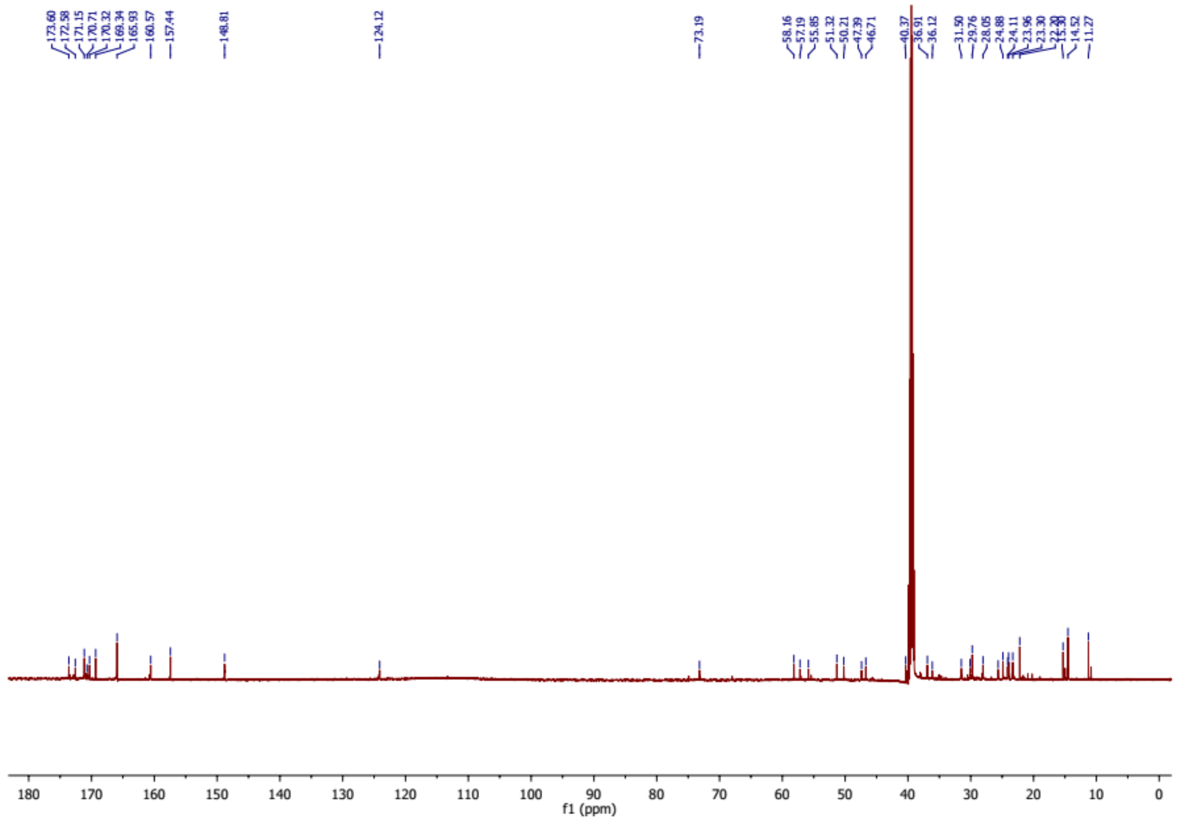


**Figure S2.24.**$^{13}$C NMR (150 MHz) spectrum of **4c**.

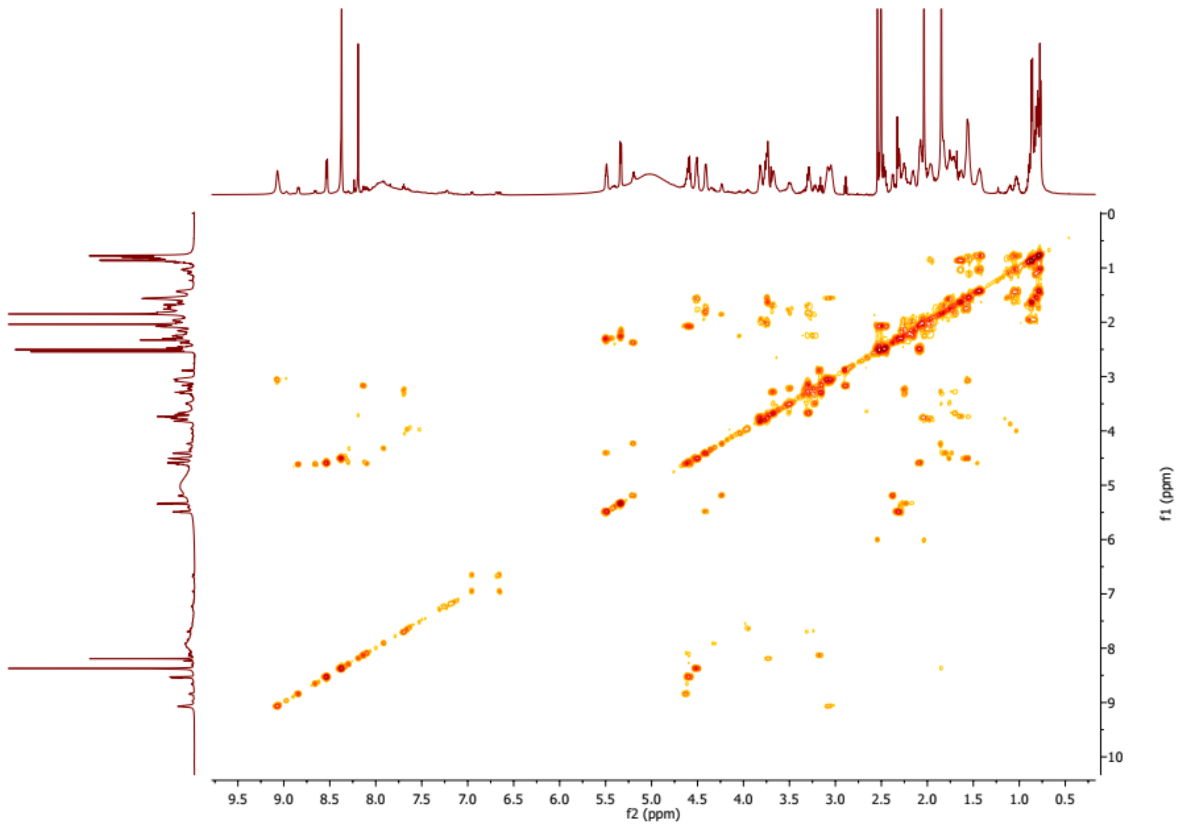


**Figure S2.25.** $^{1}H$-$^{1}H$ COSY spectrum of **4c**.

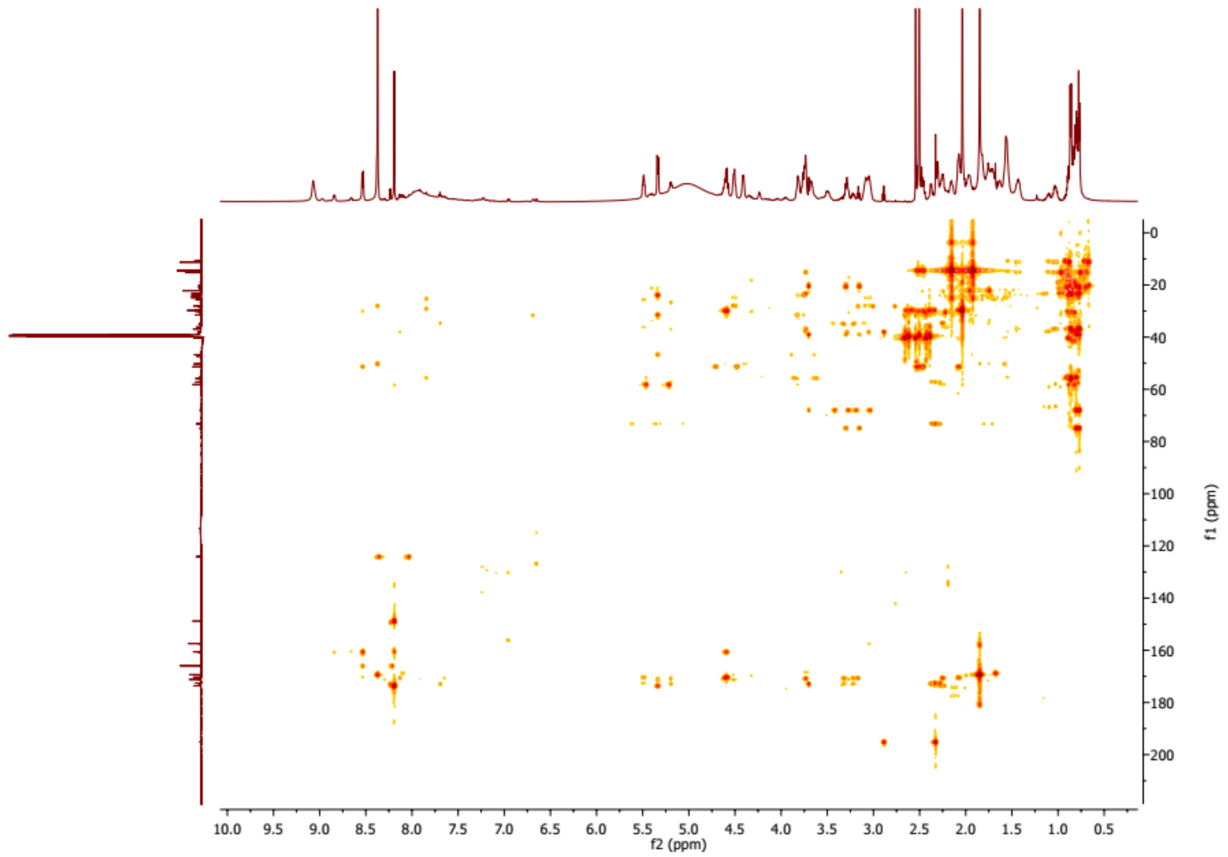


**Figure S2.26.** HMBC spectrum of **4c**.

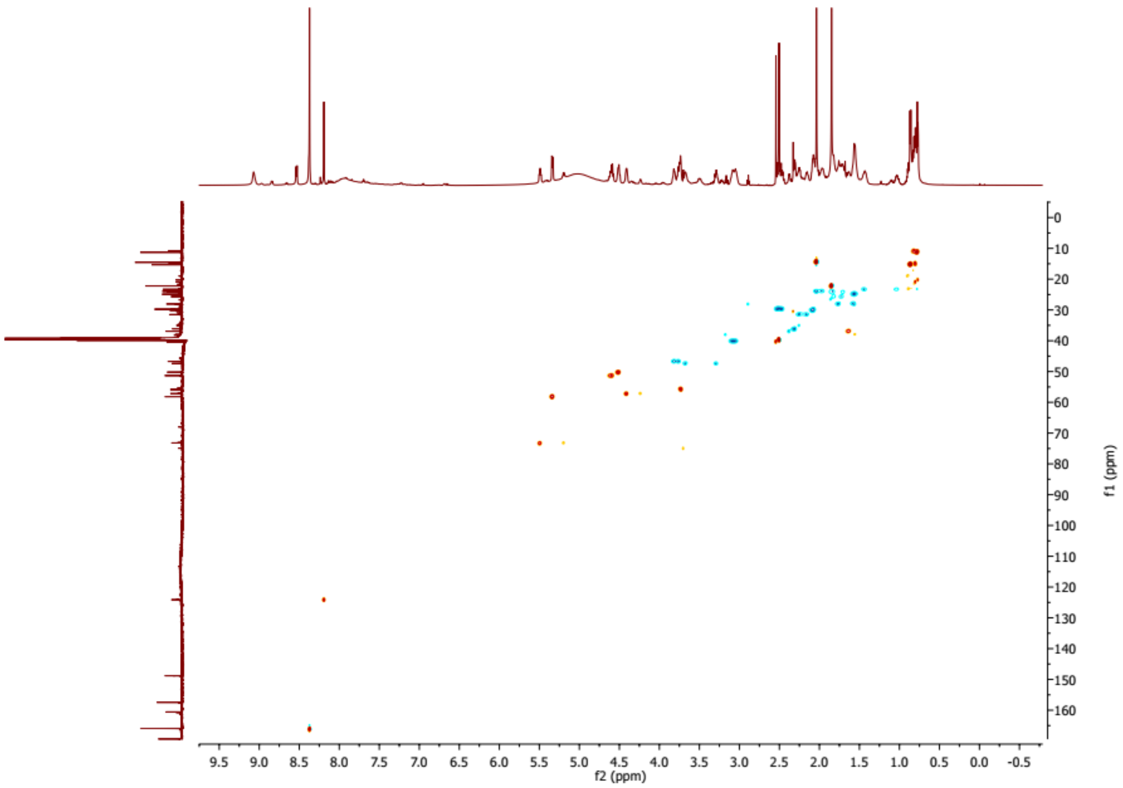


**Figure S2.27.** HSQC spectrum of **4c**.

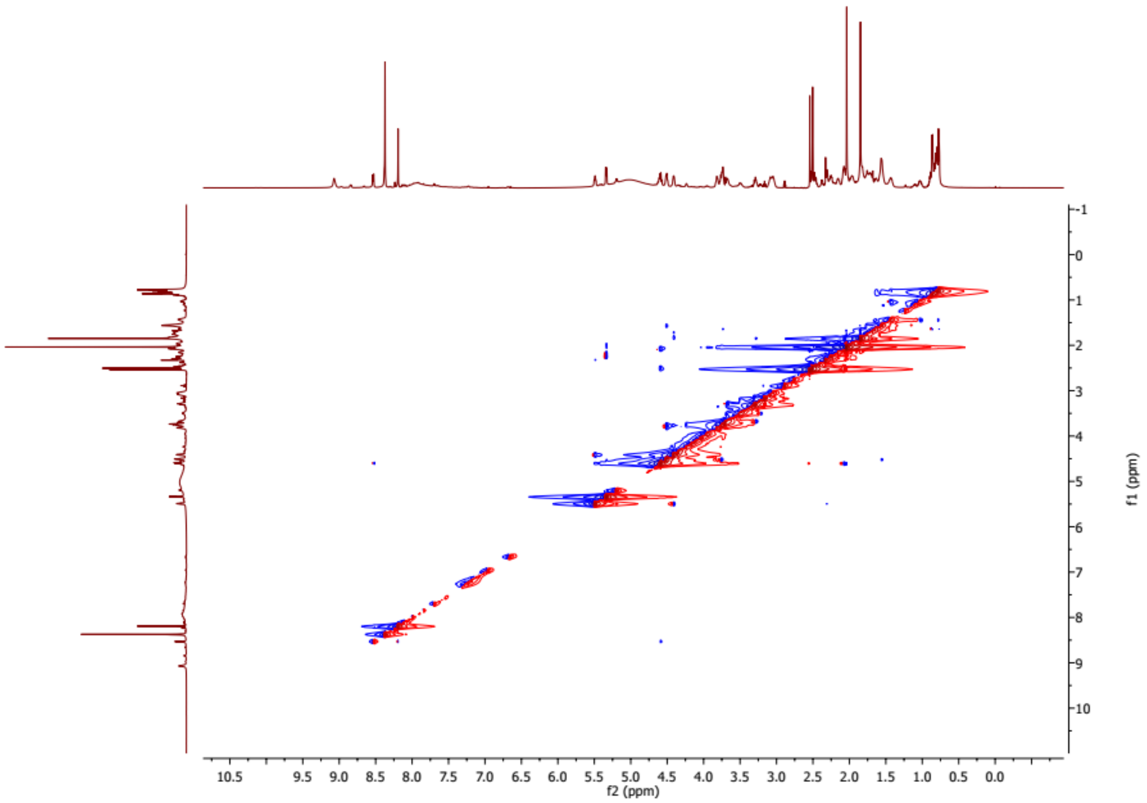


**Figure S2.28.** ROESY spectrum of **4c**.

**Structure elucidation of 4e**

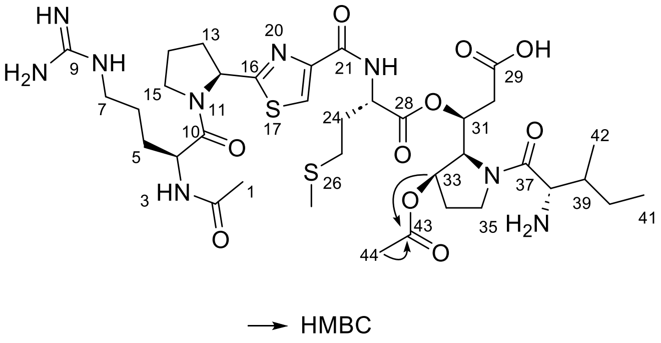


**Figure S2.29.** HMBC correlations of compound **4e**. Compound **4e** has a molecular formula of $C_{36}H_{57}N_9O_{10}S_2$, as determined by HR-ESI-MS (found, *m/z* 840.3760 $[M+H]^+$; calcd for $C_{34}H_{56}N_9O_8S_2$, *m/z* 840.3743 $[M+H]^+$; Δppm 2.1). By careful comparison of the HSQC and HMBC spectrum of **4c** and **4e**, an additional acetyl group which linked to C-33 in **4e** was established. This was deduced from the appearance of $\delta_{H\text{-}33}$ 5.09 with $\delta_{c\text{-}33}$ 71.2 in **4e** and disappearance of $\delta_{H\text{-}33}$ 1.73/1.82 and $\delta_{c\text{-}33}$ 25.6 in **4c**, additionally, $\delta_{H\text{-}44}$ 2.01 and $\delta_{c\text{-}44}$ 21.0 in **4e** in HSQC spectrum. In HMBC spectrum, correlations from H-33/H-44 to C-43 support the acetyl group attached to C-33. By comparing the chemical shifts and coupling constants of H-32 [*δ* 4.77 (d, 7.2)] and H-33 [*δ* 5.09 (dd, 15.1, 7.2)] in compound **4e** with those of chitinimide D[1] and rimosamide A[2], which share an identical hydroxyproline moiety, the relative configuration between H-32 and H-33 is assigned as *cis*. Given that the absolute configuration at C-32 is predicted to be *S* based on domain annotation, the configuration at C-33 is thereby determined to be *R*.

**Table S2.2.** $^{1}H$ (400 MHz) and $^{13}C$ (125 MHz) NMR spectroscopic data in DMSO-$d_6$ ($\delta$ in ppm and *J* in Hz).

| no. | $\delta_H$ (mult., J) | $\delta_C$ |
|---|---|---|
| 1 | 1.86 (s) | 22.8 |
| 2 | | 169.8 |
| 3 | 8.34 (overlap) | |
| 4 | 4.61 (m) | 51.6 |
| 5 | 1.76 (br s) | 28.6 |
| | 1.57 (br s) | |
| 6 | 1.57 (overlap) | 25.4 |
| 7 | 3.09 (br s) | 40.7 |
| 12 | 5.34 (m) | 58.7 |
| 13 | 2.28 (m) | 31.9 |
| | 2.16 (m) | |
| 14 | 2.06 (overlap) | 24.4 |
| | 1.96 (overlap) | |
| 15 | 3.80 (m) | 47.2 |
| 18 | 8.20 (br s) | 124.8 |
| 22 | 8.32 (d, 7.1) | |
| 23 | 4.62 (m) | 51.6 |
| 24 | 2.12 (m) | 30.1 |
| 25 | 2.52 (m) | 30.2 |
| 27 | 2.06 (s) | 15.1 |
| 30 | 2.42 (m) | 36.3 |
| 31 | 5.38 (m) | 70.2 |
| 32 | 4.77 (d, 7.2) | 57.9 |
| 33 | 5.09 (dd, 15.1, 7.2) | 71.2 |
| 34 | 2.00 (m) | 30.2 |
| | 2.22 (overlap) | |
| 35 | 3.30 (m) | 48.1 |
| | 3.72 (m) | |
| 38 | 3.88 (m) | 55.3 |
| 39 | 1.67 (m) | 37.1 |
| 40 | 1.05 (m) | 23.8 |
| | 1.45 (m) | |
| 41 | 0.77 (t, 7.0) | 11.6 |
| 42 | 0.86 (d, 7.1) | 15.5 |
| 43 | | 170.6 |
| 44 | 2.01 (s) | 21.0 |

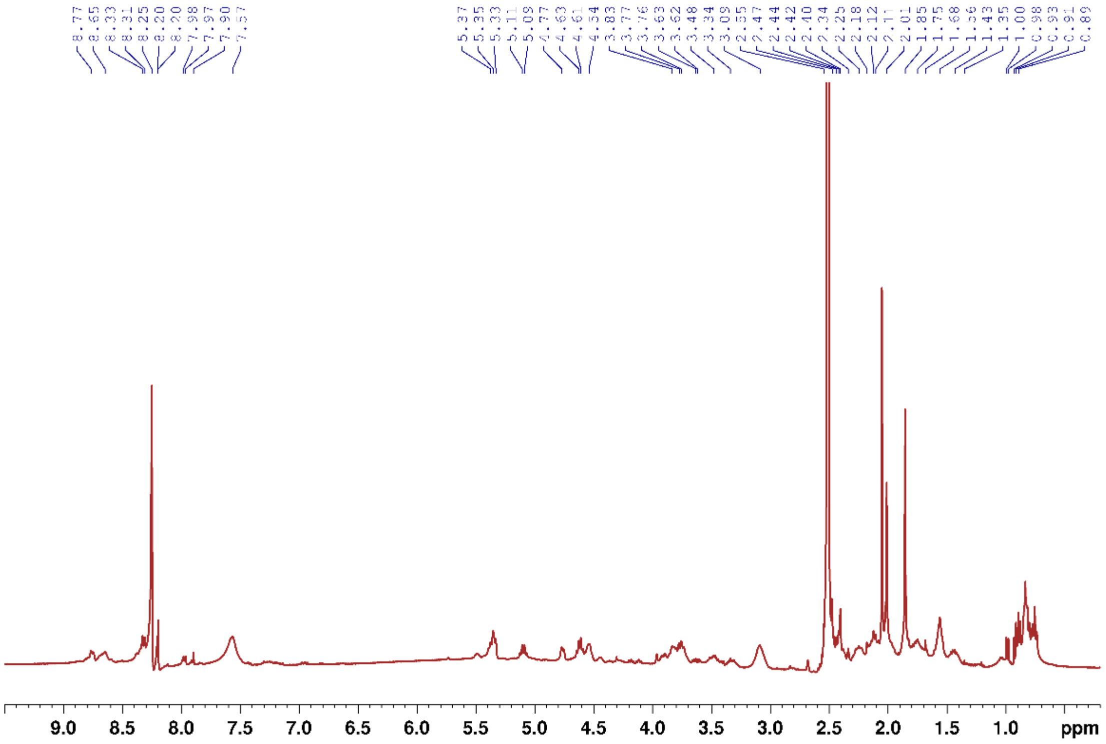

**Figure S2.30.** $^{1}$H NMR (400 MHz) spectrum of compound **4e**.

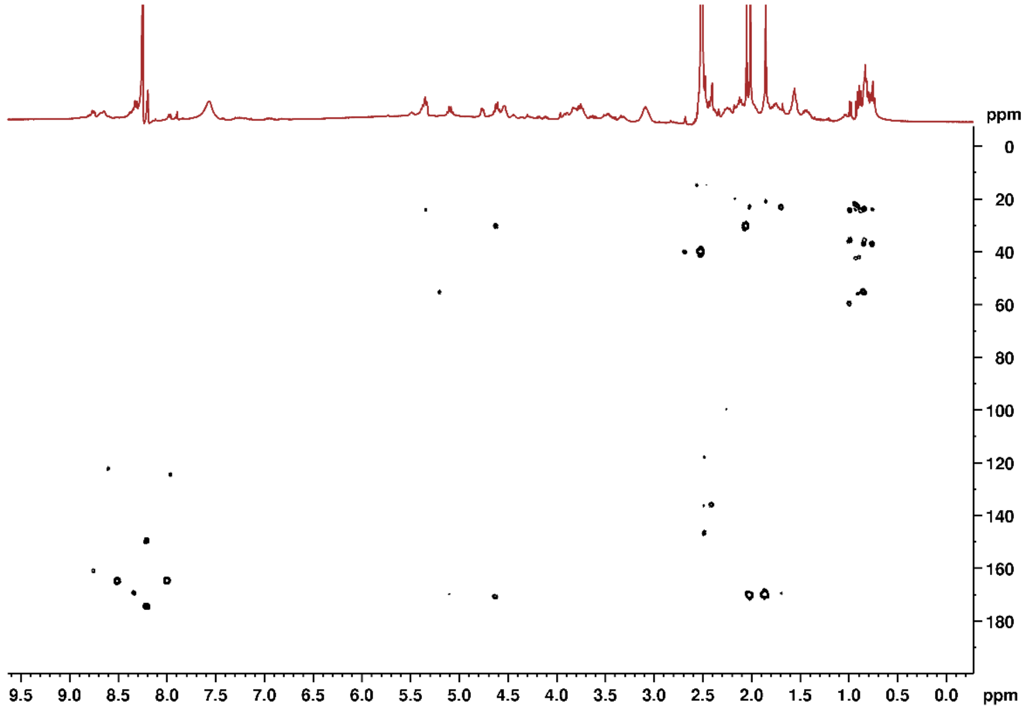

**Figure S2.31.** HMBC spectrum of compound **4e**.

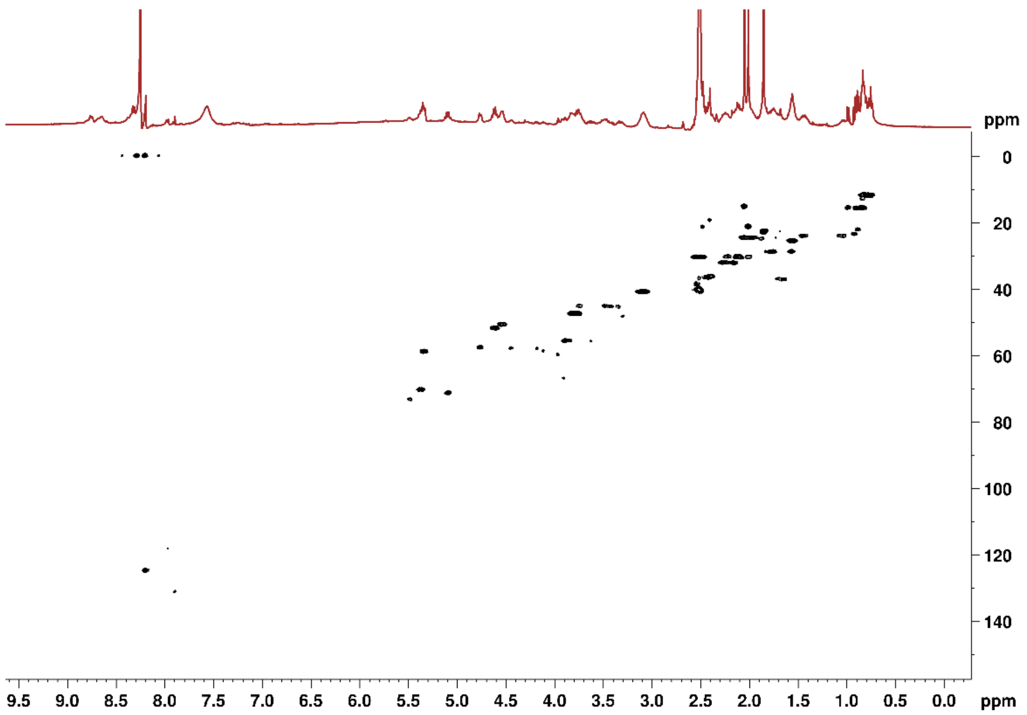


**Figure S2.32.** HSQC spectrum of compound **4e**.
