## Supplementary Information SI-3 for "Identification of pseudotetraivprolide from *Pseudomonas entomophila* give novel insights into the biosynthesis of detoxin/rimosamide-like anti-antibiotics"

### Supporting Information SI-3: Chemical synthesis

#### Abbreviation list

| | |
|---|---|
| Ac | acetyl |
| BINAP | 2,2'-bis(diphenylphosphino)-1,1'-binaphthyl |
| brsm | based on recovered starting material |
| Boc | *tert*-butyloxycarbonyl |
| Cbz | benzyloxycarbonyl |
| CI | chemical ionization |
| DCC | *N,N*'-dicyclohexylcarbodiimide |
| DCM | dichloromethane |
| DMAP | 4-dimethylaminopyridine |
| DME | 1,2-dimethoxyethane |
| DMF | *N,N*-dimethylformamide |
| DMSO | dimethyl sulfoxide |
| ECF | ethyl chloroformate |
| ESI | electrospray ionization |
| Et | ethyl |
| Fmoc | fluorenylmethyloxycarbonyl |
| HATU | 1-[Bis(dimethylamino)methylene]-1H-1,2,3-triazolo[4,5-b]pyridinium 3-oxide hexafluorophosphate |
| HBTU | 3-[Bis(dimethylamino)methyliumyl]-3H-benzotriazol-1-oxide hexafluorophosphate |
| HOBt | *N*-hydroxy benzotriazole |
| HPLC | high performance liquid chromatography |
| LC | liquid chromatography |
| Me | methyl |
| MS | mass spectrometry |
| NMM | *N*-methyl morpholine |
| NMR | nuclear magnetic resonance |
| PE | petroleum ether |
| Ph | phenyl |
| PyAOP | (7-Azabenzotriazol-1-yloxy)tripyrrolidinophosphonium hexafluorophosphate |
| $R_f$ | retention factor |
| sat. | saturated |
| TFA | trifluoroacetic acid |
| TFAA | trifluoroacetic anhydride |
| THF | tetrahydrofurane |
| $t_R$ | retention time |

#### General Information

All reactions were carried out in oven-dried glassware under an atmosphere of $N_2$, if the use of an anhydrous solvent is mentioned in the procedure. Anhydrous THF was prepared by distillation over sodium/benzophenone. Other anhydrous solvents were purchased from Acros Organics and Thermo Scientific.

**NMR spectra** were measured on a *Bruker* Avance II 400 (400 MHz, 5 mm BBO Probe, 298 K), a *Bruker* Avance I 500 (500 MHz, 5 mm TCI Probe, 298 K) or a *Bruker* Avance Neo 500 (500 MHz, 5 mm TCI Prodigy CryoProbe, 298 K). Spectra were calibrated on the solvent signals $CDCl_3$ ($^1H$ 7.27 ppm, $^{13}C$ 77.16 ppm) or DMSO-d6 ($^1H$ 2.50 ppm, $^{13}C$ 39.52 ppm). The spectral data was analyzed with MestReNova 14.2 from *MestreLabReasearch S.L.* Chemical shifts ($\delta$) are reported in ppm and coupling constants (J) in Hz. Multiplicities in $^1H$-NMR spectra are reported as singlet (s), doublet (d), triplet (t), quartet (q) and multiplet (m). $^{13}C$-NMR spectra were measured Broadband decoupled, theoretical multiplicity of the carbon is given as s (quaternary C-atom), d (tertiary C-Atom, t (secondary C-Atom) and q (primary C-Atom). Assignments were done by utilizing two-dimensional measurements like H,H-COSY, HSQCED, HMBC and TOCSY.

Reaction monitoring was done by thin-layer chromatography (**TLC**) on Polygram® SIL G/$UV_{254}$ plates by *Macherey-Nagel* ($KMnO_4$, cerium-molybdate or ninhydrin stains) and/or **LC-MS** analysis on a *Shimadzu* Prominence LC-2030 (*Phenomenex* Onyx® C18, 50 x 4.6 mm) coupled with *Shimadzu* LCMS-2020 (ESI ionization). All runs were performed at a flow rate of 4 mL/min with a column temperature of 40 °C and 0.1% $HCOOH_{aq}$/MeCN as mobile phase.

| **0.1% $HCOOH_{aq}$ : MeCN** | **short Method (time)** | **long Method (time)** |
|---|---|---|
| **90:10 to 1:99** | 1.5 min | 6.0 min |
| **1:99** | 1.0 min | 1.5 min |
| **90:10** | 0.7 min | 1.0 min |

For **column chromatography** silica gel 60M 40 – 63 µm by *Macherey Nagel* was used. Automated flash column chromatography was done on a *Büchi* Pure C815 Flash with *Teledyne* Isco RediSep $R_f$ cartridges. Automated reversed phase column chromatography was done on a *Büchi* Reveleris® Prep with *Büchi* FlashPure Select C18 (spherical) cartridges or *Kinesis* Telos C18 cartridges. Preparative HPLC was done with a *Büchi* Reveleris® Prep with a *Phenomenex* Luna® (C18, 5 µm, 21.2 x 250 mm).

Specific optical rotation ($[\boldsymbol{\alpha}]_D^{20}$) was measured on a P-8000-T polarimeter by *A. Krüss Optronic GmbH* with a PT80 thermostat by *A. Krüss Optronic GmbH* at 20 °C (λ = 589 nm). Concentrations are given in g/100 mL.

High-resolution mass spectra (**HRMS**) were recorded at Saarland University by Rudi Thomes on a *Finnigan* MAT 95 (CI, sector field) or at HIPS Saarland on a *Bruker* maXis 4G hr-ToF (ESI, ToF).

**Melting points** were measured in open glass capillaries on an M3000 from *Krüss*.

#### Overview of the Synthesis

The literature known thiazole fragment **9** (scheme 1) was synthesized analogous to the procedure from Deng and Taunton's synthesis of ceratospongamide.[1] Boc-Pro-$NH_2$ **SI-1** was prepared from Boc-Pro-OH by activation as mixed anhydride and quenching with aqueous ammonia. Thioamidation via Lawesson's reagent[2] followed by thiazole formation in a modified Hantzsch procedure by Meyers *et al.*[3] gave the thiazole fragment **9** in 73% yield over 3 steps. After saponification with lithium hydroxide, thiazole **SI-3** was coupled with L-methionine methyl ester in a mixed anhydride mediated peptide coupling with 81% yield.

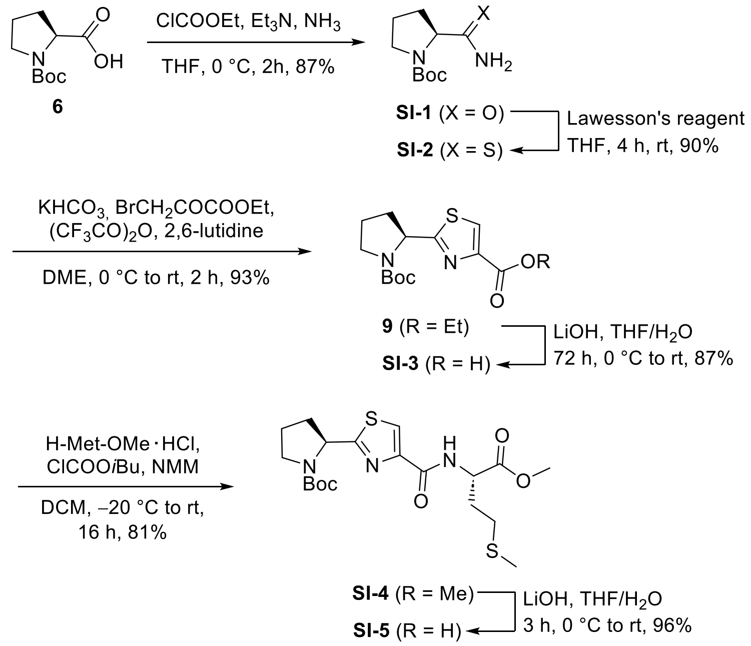


**Scheme 1.** Preparation of the thiazole building block.

Another saponification with lithium hydroxide gave the thiazole dipeptide **SI-5** in 96% yield. The β-hydroxyester **7** (scheme 2) was prepared by a procedure from Greck *et al.*[4] In the first step, the β-ketoester was formed via 1,1'-carbonyldiimidazole (CDI) activation and potassium methyl malonate (KMM) as nucleophile. The resulting β-ketoester was reduced in a Noyori asymmetric hydrogenation[5] with a in-situ formed ruthenium catalyst, first described by Genêt *et al.*[6] Boc-deprotection, followed by a HATU coupling with Cbz-Val-OH gave the dipeptide **SI-6** in 82% yield over 2 steps. In a next step, the esterification with the thiazole building block **SI-5** was investigated. All the tested conditions (Yamaguchi[7], Ghosez[8] and Steglich-type[9]) gave the depsipeptide **SI-7** in 12% to 42% yield with at least 30% α-epimerization at the methionine moiety. Additionally, several deprotection trials of the Cbz-carbamate, were unsuccesful due to thioether induced inhibition[10] during the catalytic hydrogenation. Saponification of the methyl ester was also troublesome due to the lability of the internal ester of depsipeptide **SI-7**.

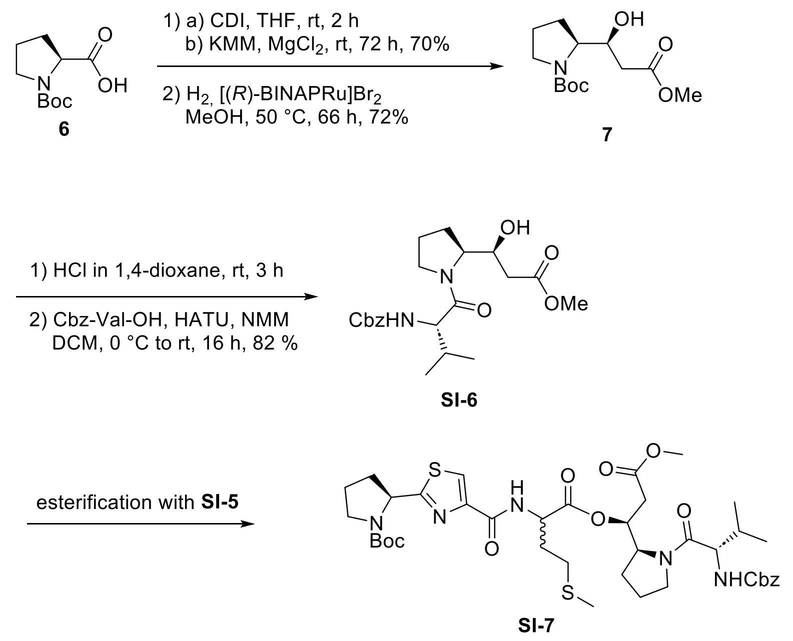


**Scheme 2.** Attempted esterification of the thiazole and ivprolid building blocks

These unsatisfying results led to a shift in the protecting group strategy to an acidic deprotection, and an alternative disconnection approach to avoid a late-stage esterification. The methyl ester **SI-6** was transformed into *tert*-butyl ester **SI-8** using *tert*-butyl 2,2,2-trichloroacetimidate (TBTA) in a protocol by Hutton *et al.*[11] (scheme 3). The Cbz-carbamate was replaced by a Boc-carbamate by catalytic hydrogenation in the presence of Boc-anhydride. The reprotected β-hydroxy ester **SI-9** was coupled with Fmoc-protected L-methionine in a standard Steglich esterification.

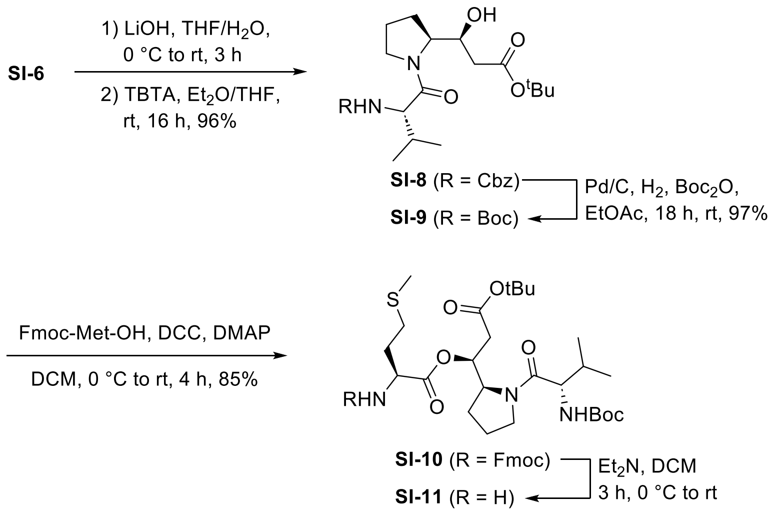


**Scheme 3.** Finalization of the valine depsipeptide building block **SI-11**.

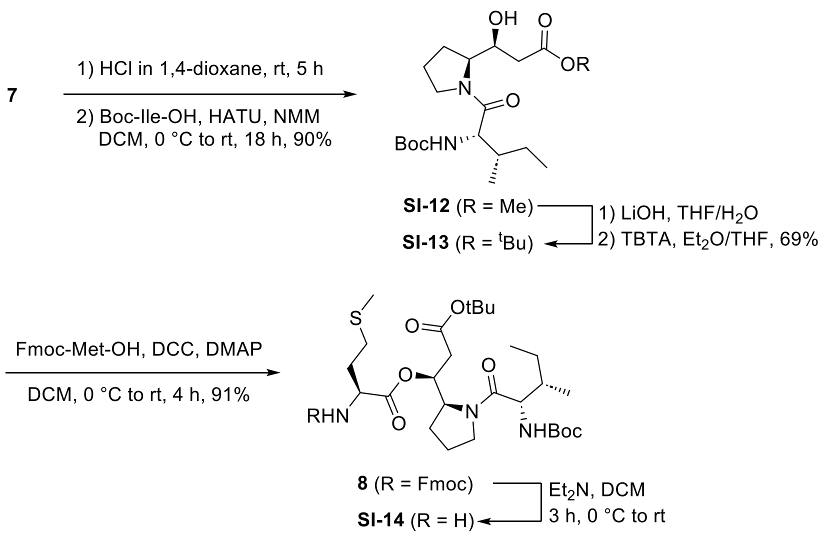


**Scheme 4.** Synthesis of the isoleucine depsipeptide building block **SI-14**

An excess of the protected L-methionine resulted in an improved yield and suppressed the epimerization at the α-position of the methionine. On the basis of the new protecting group strategy the isoleucine depispeptide building block was synthesized in a similar fashion (scheme 4). At first, the β-hydroxyester **7** was coupled with Boc-protected isoleucine in 90% yield. Transesterification to the *tert*-butyl ester and steglich esterification with Fmoc-Met-OH gave the depsipeptide **8** without observable epimerization.

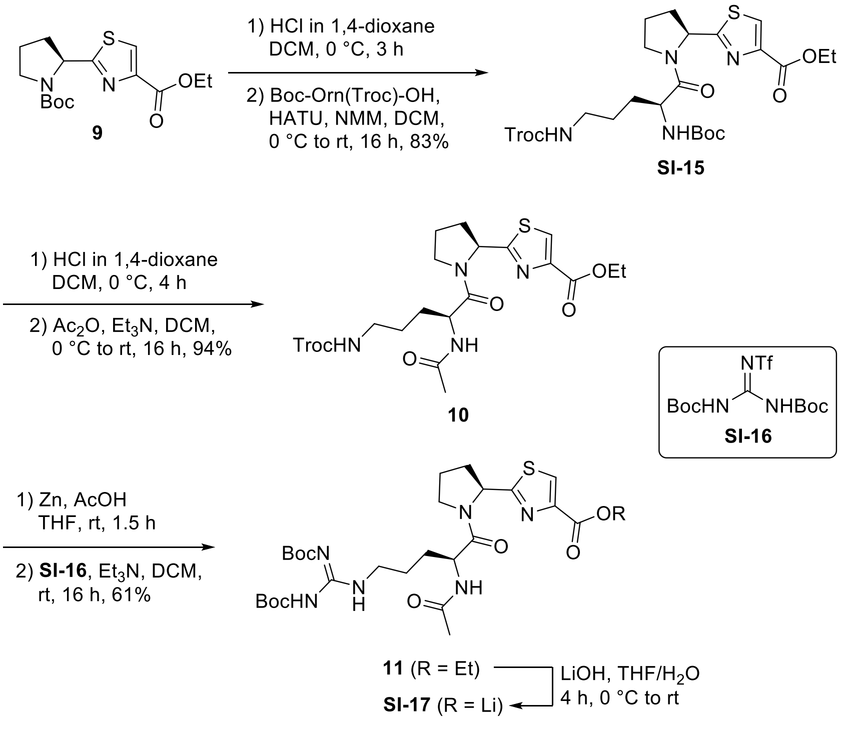


**Scheme 5.** Introduction of the arginine moiety.

With the two depsipeptide building blocks **SI-11** and **SI-14** in hand, we continued with the synthesis of the western fragment (scheme 5). After Boc-deprotection, the thiazole building block **9** was coupled with Boc-Orn(Troc)-OH in a HATU mediated peptide coupling with 83% yield.

Couplings with acetylated ornithine and arginine resulted in strong epimerization at the α-position, which is a known problem for acetylated amino acids.[12,13] After removal of the Boc-carbamate, acetylation with acetic anhydride gave the dipeptide **10** in 94% yield. In a last step, the Troc-carbamate was removed to introduce the Di-Boc-guanidine via its triflate **SI-16.**[14] Finally, The amino depsipeptides **SI-11** and **SI-14** were coupled with the lithium carboxylate of the western fragment **19** using HBTU as coupling agent. In both cases the yields were only moderate with 35% and 47%, respectively. This coupling wasn't optimized because of the limited amount of reactants. In the last step, a final deprotection of the protected pseudotetraivprolids **SI-18** and **12** was carried out with a cleavage cocktail to suppress side reactions induced by the *tert*-butyl cations.[15] Pseudotetraivprolid D **5a** was isolated in a excellent yield of 85% after preparative HPLC. In the case of pseudotetraivprolid B **4a** an incomplete cleavage resulted in only 37% yield with additional 32% of the corresponding *tert*-butyl ester.

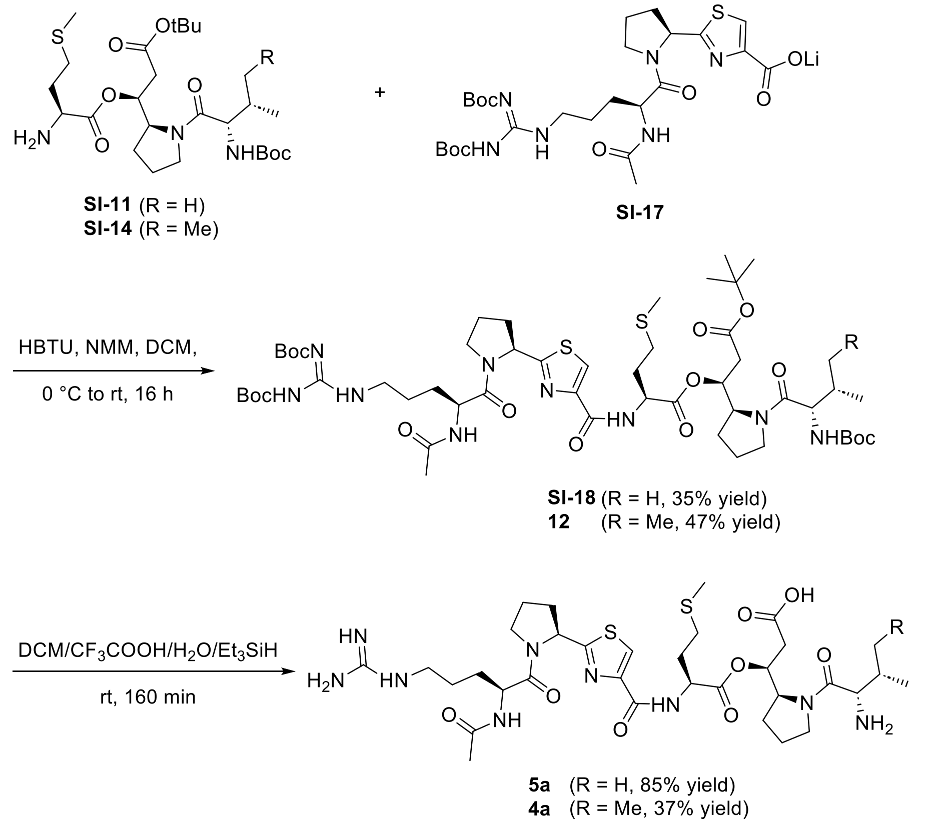


**Scheme 6.** Final coupling and deprotection to pseudotetraivprolid B and D (**4a** and **5a**).

#### Experimental Section

##### ***tert*-Butyl (*S*)-2-carbamoylpyrrolidine-1-carboxylate SI-1**

Ethyl chloroformate (578 µL, 6.02 mmol, 1.2 eq.) was dropwise added to a 0 °C cold solution of Boc-Pro-OH (1.08 g, 5.02 mmol) and triethylamine (839 µL, 6.02 mmol, 1.2 eq.) in anhydrous THF (20 mL). The resulting mixture was stirred for 1 h at 0 °C. Aqueous ammonia solution (3.33 mL, 60.2 mmol, 35 wt%, 12 eq.) was added to the reaction mixture, and stirring was continued for 1 h. The reaction mixture was diluted with EtOAc and water. The aqueous layer was extracted with EtOAc (3 x 100 mL). The combined organic extracts were dried over $MgSO_4$ and concentrated in vacuo to give Boc-Pro-$NH_2$ **SI-1** (939 mg, 4.38 mmol, 87%) as a white solid.

**TLC: $R_f$ (SI-1)** = 0.16 (silica, PE:EtOAc 1:1 + 1% AcOH)

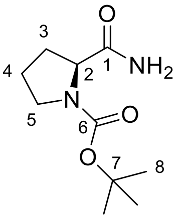


**SI-1**

**$^1$H-NMR** (500 MHz, DMSO-$d_6$): δ = 7.30 (s, 1 H, 1-$NH_2$), 6.91 (s, 1 H, 1-$NH_2$), 3.97 (dd, $^3J_{2,3}$ = 8.4 Hz, $^3J_{2,3'}$ = 3.6 Hz, 1 H, 2-H), 3.36 (m, 1 H, 5-H'), 3.25 (m, 1 H, 5-H), 2.09 (m, 1 H, 3-H'), 1.84 – 1.68 (m, 3 H, 4-H, 3-H), 1.34 (s, 9 H, 8-H).

**$^{13}$C-NMR** (125 MHz, DMSO-$d_6$): δ = 174.6 (s, C-1), 153.3 (s, C-6), 78.3 (s, C-7), 59.6 (d, C-2), 46.4 (t, C-5), 31.0 (t, C-3), 28.1 (q, C-8), 23.2 (t, C-4).

**Selected rotamer signals:**

**$^1$H-NMR** (500 MHz, DMSO-$d_6$): δ = 7.27 (s, 1 H, 1-$NH_2$), 6.86 (s, 1 H, 1-$NH_2$), 4.00 (dd, $^3J_{2,3}$ = 8.8 Hz, $^3J_{2,3'}$ = 2.9 Hz, 1 H, 2-H), 2.02 (m, 1 H, 3-H'), 1.39 (s, 9 H, 8-H).

**$^{13}$C-NMR** (125 MHz, DMSO-$d_6$): δ = 174.3 (s, C-1), 153.6 (s, C-6), 78.4 (s, C-7), 59.4 (d, C-2), 46.4 (t, C-5), 30.0 (t, C-3), 28.2 (q, C-8), 23.9 (t, C-4).

| | | |
|---|---|---|
| **Optical rotation:** | $[\alpha]_D^{20}$ = −99.4 (c = 0.5, $CHCl_3$) | |
| **HRMS (CI):** | calculated | found |
| $C_{10}H_{19}N_2O_3$ $[M+H]^+$: | 215.1390 | 215.1377 |
| **Melting point:** | 105 – 107 °C (lit: 103.6 – 107.7 °C)[16] | |

##### **Ethyl (*S*)-2-(1-(*tert*-butoxycarbonyl)pyrrolidin-2-yl)thiazole-4-carboxylate 9**[1]

According to Taunton *et al.*[1], Boc-Pro-$NH_2$ **SI-1** (1.81 g, 8.45 mmol) was dissolved in anhydrous THF (25 mL) under an atmosphere of $N_2$. Lawesson's reagent (1.71 g, 4.22 mmol, 0.5 eq.) was added, and the resulting solution was stirred for 5 h at room temperature. The reaction mixture was concentrated in vacuo and the crude product was purified by column chromatography (silica, 97:3 DCM:MeOH) to give thioamide **SI-2** (1.76 g, 7.64 mmol, 90%) as a white solid.

$KHCO_3$ (2.02 g, 10.1 mmol, 4.0 eq.) was added to a solution of above-prepared thioamide **SI-2** (580 mg, 2.52 mmol) in anhydrous DME (10 mL). The resulting suspension was stirred for 10 min at room temperature before ethyl bromopyruvate (1.05 mL, 7.55 mmol, 3.0 eq.) was added dropwise. The reaction mixture was stirred for 30 min at room temperature and then cooled to 0 °C. A preformed solution of TFAA (1.42 mL, 10.1 mmol, 4.0 eq.) and 2,6-lutidine (2.49 mL, 21.4 mmol, 8.5 eq.) in anhydrous DME (3.0 mL) was added dropwise to the reaction mixture over 10 min. The reaction mixture was stirred for 2 h while slowly reaching room temperature before being quenched by the addition of 1.0 M $HCl_{aq.}$ and EtOAc. After extraction of the aqueous layer with EtOAc, the combined organic extracts were dried over $MgSO_4$ and concentrated in vacuo. The crude product was purified twice by column chromatography (1. silica, PE:EtOAc 6:4; 2. silica, PE:EtOAc 75:25) to give thiazole **9** (763 mg, 2.34 mmol, 93%) as a white solid.

**TLC: $R_f$ (9)** = 0.39 (silica, PE:EtOAc 1:1)

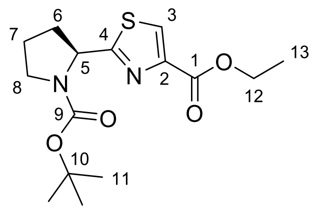


**9**

**$^1$H-NMR** (500 MHz, $CDCl_3$): δ = 8.07 (s, 1 H, 3-H), 5.20 (m, 1 H, 5-H), 4.42 (q, $^3J_{12,13}$ = 7.2 Hz, 2 H, 12-H), 3.62 (m, 1 H, 8-H‘), 3.52 (ddd, $^2J_{8,8'}$ = 9.1 Hz, $^3J_{8,7}$ = 8.9 Hz, $^3J_{8,7'}$ = 8.9 Hz, 1 H, 8-H), 2.35 (m, 1 H, 6-H‘), 2.23 (m, 1 H, 6-H), 1.92 (m, 2 H, 7-H), 1.40 (t, $^3J_{13,12}$ = 7.1 Hz, 3 H, 13-H), 1.32 (s, 9 H, 11-H).

**$^{13}$C-NMR** (125 MHz, DMSO-$d_6$, 373 K): δ = 174.7 (s, C-4), 160.2 (s, C-1), 153.1 (s, C-9), 145.6 (s, C-2), 127.4 (d, C-3), 78.8 (s, C-10), 60.0 (d, C-5), 58.3 (t, C-12), 46.1 (t, C-8), 32.4 (t, C-6), 27.5 (q, C-11), 22.6 (t, C-7), 13.6 (q, C-13).

**Selected rotamer signals:**

**$^1$H-NMR** (500 MHz, $CDCl_3$): δ = 5.27 (m, 1 H, 5-H), 3.44 (m, 1 H, 8-H), 1.48 (s, 9 H, 11-H).

| | | |
|---|---|---|
| **Optical rotation:** | $[\alpha]_D^{20}$= –75.6 (c = 1.0, $CHCl_3$) | |
| **HRMS (CI):** | calculated | found |
| $C_{15}H_{22}O_4N_2S$ $[M]^+$: | 326.1295 | 326.1299 |
| **Melting point:** | 98 – 101 °C | |

**tert-Butyl (S)-2-(4-(((S)-1-methoxy-4-(methylthio)-1-oxobutan-2-yl)carbamoyl)thiazol-2-yl)-pyrrolidine-1-carboxylate SI-4**

1.0 M $LiOH_{aq}$ (4.60 mL, 4.60 mmol, 1.2 eq.) was added dropwise to a 0 °C cold solution of thiazole **9** (1.25 g, 3.83 mmol) in THF (20 mL). The reaction mixture was stirred for 72 h before being diluted with EtOAc and acidified with 1.0 M $HCl_{aq}$. The aqueous layer was extracted with EtOAc, and the combined organic extracts were washed with brine. After drying over $MgSO_4$ and concentration in vacuo, the crude acid **SI-3** (1.10 g, 3.32 mmol, 87%, 90wt% purity) was isolated as a yellow solid.

The crude acid **SI-3** (1.08 g, 3.26 mmol, 90 wt%) and NMM (895 µL, 8.14 mmol, 2.5 eq.) were dissolved in anhydrous $CH_2Cl_2$ (25 mL) before being cooled to –20 °C. IBCF (513 µL, 3.91 mmol, 1.2 eq.) was added dropwise, and the resulting solution was stirred for 30 min at –20 °C. H-Met-OMe·HCl (781 mg, 3.91 mmol, 1.2 eq.) was added in one portion, and the reaction mixture was allowed to reach room temperature for 16 h. After dilution with EtOAc, the solution was washed with 1.0 M $HCl_{aq.}$, sat. $NaHCO_3$ solution and brine. Drying over $MgSO_4$ and concentration in vacuo led to the crude product, which was purified by column chromatography (silica, PE:EtOAc 65:35) to give dipeptide **SI-4** (1.17 g, 2.64 mmol, 81%) as a light-yellow resin.

**TLC: $R_f$ (SI-4)** = 0.13 (silica, PE:EtOAc 7:3)

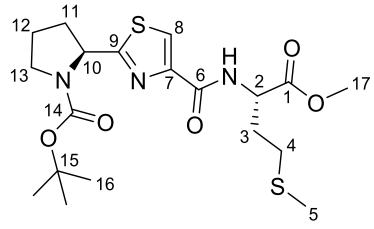


**SI-4**

**$^1$H-NMR** (500 MHz, DMSO-d$_6$, 373 K): δ = 8.19 (d, $^3J_{NH,2}$ = 8.0 Hz, 1 H, 6-NH), 8.15 (s, 1 H, 8-H), 5.12 (dd, $^3J_{10,11'}$ = 8.2 Hz, $^3J_{10,11}$ = 3.1 Hz, 1 H, 10-H), 4.66 (dt, $^3J_{2,NH}$ = 7.9 Hz, $^3J_{2,3}$ = 5.7 Hz, 1 H, 2-H), 3.69 (s, 3 H, 17-H), 3.52 – 3.42 (m, 2 H, 13-H, 13-H'), 2.55 (m, 2 H, 4-H), 2.37 (ddt, $^2J_{11',11}$ = 12.8 Hz, $^3J_{11',10}$ = 8.5 Hz, $^3J_{11',12}$ = 8.5 Hz, 1 H, 11'-H), 2.18 – 2.10 (m, 3 H, 11-H, 3-H), 2.07 (s, 3 H, 5-H), 1.94 (m, 2 H, 12-H), 1.32 (s, 9 H, 16-H).

**$^{13}$C-NMR** (125 MHz, DMSO-d$_6$, 373 K): δ = 174.7 (s, C-9), 171.2 (s, C-1), 160.0 (s, C-6), 153.1 (s, C-14), 148.5 (s, C-7), 123.2 (d, C-8), 78.8 (s, C-15), 58.3 (d , C-10), 51.4 (q, C-17), 50.9 (d, C-2), 46.1 (t, C-13), 32.5 (t, C-11), 30.4 (t, C-3), 29.5 (t, C-4), 27.5 (q, C-16), 22.7 (t, C-12), 14.2 (q, C-5).

| | | |
|---|---|---|
| **Optical rotation:** | $[\alpha]_D^{20}$ = –50.7 (c = 1.0, $CHCl_3$) | |
| **HRMS (CI):** | calculated | found |
| $C_{19}H_{30}N_3O_5S_2$ $[M+H]^+$: | 444.1621 | 444.1625 |

**(2-((S)-1-(tert-Butoxycarbonyl)pyrrolidin-2-yl)thiazole-4-carbonyl)-L-methionine SI-5**

Dipeptide **SI-4** (488 mg, 1.10 mmol) was dissolved in THF (4.0 mL). The solution was cooled to 0 °C, and 0.30 M $LiOH_{aq}$ (3.85 mL, 1.16 mmol, 1.05 eq.) was added. After stirring for 3 h, while slowly reaching room temperature, the reaction mixture was diluted with diethyl ether and acidified with 0.1 M $HCl_{aq}$. The aqueous layer was extracted with diethyl ether and the combined organic extracts where dried over $MgSO_4$. Concentration in vacuo led to carboxylic acid **SI-5** (452 mg, 1.05 mmol, 96%) as a colorless resin.

**TLC: $R_f$ (SI-5)** = 0.48 (silica, PE:EtOAc 4:6)

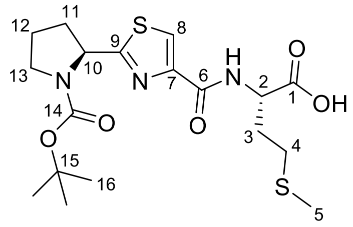


**SI-5**

**$^1$H-NMR** (500 MHz, DMSO-d$_6$, 373 K): δ = 12.47 (s, 1 H, 1-OH), 8.14 (s, 1 H, 8-H), 8.09 (d, $^3J_{NH,2}$ = 8.1 Hz, 1 H, 6-NH), 5.12 (dd, $^3J_{10,11'}$ = 8.2 Hz, $^3J_{10,11}$ = 3.2 Hz, 1 H, 10-H), 4.59 (dt, $^3J_{2,NH}$ = 8.0 Hz, $^3J_{2,3}$ = 5.2 Hz, 1 H, 2-H), 3.54 – 3.42 (m, 2 H, 13-H, 13-H‘), 2.55 (m, 2 H, 4-H), 2.36 (m, 1 H, 11-H‘), 2.18 – 2.08 (m, 3 H, 11-H, 3-H), 2.07 (s, 3 H, 5-H), 1.94 (dddd, $^3J_{12,11'}$ = 8.0 Hz, $^3J_{12,11}$ = 8.0 Hz, $^3J_{12,13}$ = 8.0 Hz, $^3J_{12,13'}$ = 6.3 Hz, 2 H, 12-H), 1.36 (s, 9 H, 16-H).

**$^{13}$C-NMR** (125 MHz, DMSO-d$_6$, 373 K): δ = 174.6 (s, C-9), 172.0 (s, C-1), 159.9 (s, C-6), 153.1 (s, C-14), 148.5 (s, C-7), 123.0 (d, C-8), 78.8 (s, C-15), 58.3 (d, C-10), 50.9 (d, C-2), 46.0 (t, C-13), 32.5 (t, C-11), 30.7 (t, C-3), 29.5 (t, C-4), 27.5 (q, C-16), 22.6 (t, C-12), 14.2 (q, C-5).

**Optical rotation:** $[\alpha]_D^{20}$ = –52.1 (c = 1.0, $CHCl_3$)

| **HRMS (CI):** | calculated | found |
|---|---|---|
| $C_{18}H_{28}N_3O_5S_2$ $[M+H]^+$: | 430.1465 | 430.1472 |

##### ***Tert*-butyl (*S*)-2-((*S*)-1-hydroxy-3-methoxy-3-oxopropyl)pyrrolidine-1-carboxylate 7[14]**

According to Greck *et al.*[14], *N,N*'-carbonyldiimidazole (2.58 g, 15.9 mmol, 1.2 eq.) was added to a solution of Boc-Pro-OH (2.85 g, 13.2 mmol) in anhydrous THF (100 mL). The reaction mixture was stirred for 2 h at room temperature before $MgCl_2$ (4.08 g, 19.9 mmol, 1.5 eq.) and potassium methyl malonate (3.13 g, 19.9 mmol, 1.5 eq.) were added. After stirring for another 72 h at room temperature, the mixture was concentrated in vacuo. The residue was partitioned between $Et_2O$ and 1.0 M $HCl_{aq,}$ and the aqueous layer was extracted with $Et_2O$. The combined organic layers were dried over $MgSO_4$ and concentrated in vacuo. The crude product was purified by column chromatography (silica, $CH_2Cl_2$:EtOAc 9:1) to give *tert*-butyl (*S*)-2-(3-methoxy-3-oxopropanoyl)-pyrrolidine-1-carboxylate (2.66 g, 9.80 mmol, 74%) as a white solid.

Before use, all the solvents in this reaction were degassed with argon. Bis-(2-methylallyl)-cycloocta-1,5-diene-ruthenium (II) complex (4.70 mg, 147 µmol, 2 mol%) and (*R*)-BINAP (9.20 mg,

147 µmol, 2 mol%) were dissolved in acetone (300 µL). The mixture was stirred for 1 h after adding 0.20 M HBr in MeOH (1.47 mL, 295 µmol, 4 mol%). The solvent was removed in vacuo, and a solution of the above-prepared β-ketoester (2.00 g, 7.37 mmol) in MeOH (12.5 mL) was added to the residue. After 5 min, the atmosphere was exchanged to $H_2$ (1 atm), and the solution was stirred for 66 h at 50 °C (sand bath). The solvent was removed in vacuo, and the crude was purified by column chromatography (silica, $CH_2Cl_2$:EtOAc 15% to 20% EtOAc) to give β-hydroxyester **7** (1.44 g, 5.27 mmol, 72%) as a colorless oil.

**TLC: $R_f$ (7)** = 0.28 (silica, DCM:EtOAc 85:15)

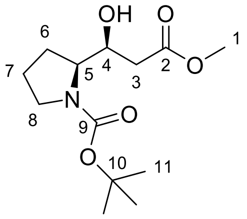


**7**

**$^1$H-NMR** (500 MHz, $CDCl_3$): δ = 5.06 (s, 1 H, 4-OH), 4.01 (m, 1 H, 4-H), 3.92 (dt, $^3J_{5,4}$ = 4.4 Hz, $^3J_{5,6}$ = 7.9 Hz, 1 H, 5-H), 3.71 (s, 3 H, 1-H), 3.49 (m, 1 H, 8-H‘), 3.30 (ddd, $^2J_{8,8'}$ = 10.9 Hz, $^3J_{8,7'}$ = 7.4 Hz, $^3J_{8,7}$ = 5.4 Hz, 1 H, 8-H), 2.52 (dd, $^2J_{3',3}$ = 15.3 Hz, $^3J_{3',4}$ = 3.3 Hz, 1 H, 3-H‘), 2.43 (dd, $^2J_{3,3'}$ = 15.2 Hz, $^3J_{3,4}$ = 8.7 Hz, 1 H, 3-H), 1.99 (ddt, $^2J_{6',6}$ = 12.5 Hz, $^3J_{6',5}$ = 7.8 Hz, $^3J_{6',7}$ = 7.8 Hz, 1 H, 6-H‘), 1.89 (dtt, $^2J_{7',7}$ = 12.7 Hz, $^3J_{7',6}$ = 7.5 Hz, $^3J_{7',8}$ = 7.2 Hz, 1 H, 7-H‘), 1.78 (m, 1 H, 7-H), 1.69 (m, 1 H, 6-H), 1.46 (s, 9 H, 11-H).

**$^{13}$C-NMR** (125 MHz, $CDCl_3$): δ = 172.2 (s, C-2), 157.3 (s, C-9), 80.4 (s, C-10), 72.6 (d, C-4), 61.8 (d, C-5), 51.7 (q, C-1), 47.3 (t, C-8), 40.0 (t, C-3), 28.3 (t, C-6, q, C-11), 24.1 (q, C-7).

**Optical rotation:** $[\alpha]_D^{20} = -83.4$ (c = 1.0, $CHCl_3$)

| **HRMS (CI):** | calculated | found |
|---|---|---|
| $C_{13}H_{24}O_5N$ $[M+H]^+$: | 274.1649 | 274.1646 |

**Methyl (*S*)-3-((*S*)-1-(((benzyloxy)carbonyl)-L-valyl)pyrrolidin-2-yl)-3-hydroxypropanoate SI-6**

β-Hydroxyester **7** (494 mg, 1.81 mmol) was dissolved in anhydrous $CH_2Cl_2$ (3.0 mL) and cooled to 0 °C. 4.0 M HCl in 1,4-dioxane (4.52 mL, 18.1 mmol, 10 eq.) was added, and the resulting solution was stirred for 3 h while slowly reaching room temperature. The reaction mixture was concentrated in vacuo to give the crude amine as hydrochloride.

A solution of the above-prepared hydrochloride salt and Cbz-Val-OH (477 mg, 1.90 mmol, 1.05 eq.) in anhydrous $CH_2Cl_2$ (15 mL) was cooled to 0 °C. NMM (596 µL, 5.42 mmol, 3.0 eq.) and HATU (720 mg, 1.90 mmol, 1.05 eq.) were added, and the resulting yellow solution was stirred for 16 h while slowly reaching room temperature. The reaction mixture was diluted with EtOAc and washed with 1.0 M $HCl_{aq}$, sat. $NaHCO_3$ solution and brine. The organic layer was dried over $MgSO_4$

and concentrated in vacuo. The crude product was purified by column chromatography (silica, PE:EtOAc 1:1) to give dipeptide **SI-6** (677 mg, 1.57 mmol, 87%) as a colorless resin.

**TLC: $R_f$ (SI-6)** = 0.51 (silica, PE:EtOAc 1:1)

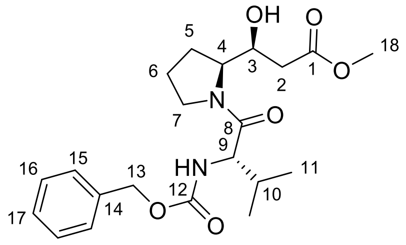


**SI-6**

**$^1$H-NMR** (500 MHz, $CDCl_3$): δ = 7.38 – 7.29 (m, 5 H, 15-H, 16-H, 17-H), 5.50 (d, $^3J_{NH,9}$ = 9.4 Hz, 1 H, 12-NH), 5.08 (m, 2 H, 13-H), 4.63 (d, $^3J_{OH,3}$ = 4.4 Hz, 1 H, 3-OH), 4.39 (dd, $^3J_{9,NH}$ = 9.2 Hz, $^3J_{9,10}$ = 6.1 Hz, 1 H, 9-H), 4.31 (ddd, $^3J_{4,3}$ = 7.7 Hz, $^3J_{4,5}$ = 7.5 Hz, $^3J_{4,5'}$ = 4.5 Hz, 1 H, 4-H), 4.01 (m, 1 H, 3-H), 3.85 (ddd, $^2J_{7',7}$ = 10.3 Hz, $^3J_{7',6}$ = 6.6 Hz, $^3J_{7',6'}$ = 6.6 Hz, 1 H, 7-H'), 3.72 (s, 3 H, 18-H), 3.53 (ddd, $^2J_{7,7'}$ = 10.3 Hz, $^3J_{7,6}$ = 6.5 Hz, $^3J_{7,6'}$ = 6.5 Hz, 1 H, 7-H), 2.52 (dd, $^2J_{2',2}$ = 15.3 Hz, $^3J_{2',3}$ = 3.4 Hz, 1 H, 2-H'), 2.46 (dd, $^2J_{2,2'}$ = 15.3 Hz, $^3J_{2,3}$ = 8.8 Hz, 1 H, 2-H), 2.07 – 1.95 (m, 3 H, 5-H', 6-H', 10-H), 1.92 (m, 1 H, 6-H), 1.70 (m, 1 H, 5-H), 1.03 (d, $^3J_{11',10}$ = 6.8 Hz, 3 H, 11-H'), 0.94 (d, $^3J_{11,10}$ = 6.8 Hz, 3 H, 11-H).

**$^{13}$C-NMR** (125 MHz, $CDCl_3$): δ = 174.1 (s, C-8), 172.3 (s, C-1), 156.6 (s, C-12), 136.4 (s, C-14), 128.7 (d, C-16), 128.3 (d, C-17), 128.2 (d, C-15), 72.8 (d, C-3), 67.1 (t, C-13), 62.3 (d, C-4), 57.8 (d, C-9), 52.0 (q, C-18), 48.2 (t, C-7), 40.3 (t, C-2), 31.7 (d, C-10), 27.8 (t, C-5), 24.7 (t, C-6), 19.6 (q, C-11'), 17.5 (q, C-11).

| | | |
|---|---|---|
| **Optical rotation:** | $[\alpha]_D^{20}$ = –58.1 (c = 1.0, $CHCl_3$) | |
| **HRMS (ESI):** | calculated | found |
| $C_{21}H_{31}O_6N_2$ $[M+H]^+$: | 407.2182 | 407.2191 |

##### *tert*-Butyl (*S*)-3-((*S*)-1-(((benzyloxy)carbonyl)-L-valyl)pyrrolidin-2-yl)-3-hydroxypropanoate SI-8

0.20 M $LiOH_{aq.}$ (1.27 mL, 254 µmol, 1.1 eq.) was added to a 0 °C cold solution of dipeptide **SI-6** (100 mg, 231 µmol) in THF (1.2 mL). The resulting solution was stirred for 3 h while slowly reaching room temperature. The reaction mixture was acidified with 1.0 M $HCl_{aq.}$ and extracted with diethyl ether. The combined organic layers were dried over $MgSO_4$ and concentrated in vacuo to give the crude carboxylic acid as a white foam.

A solution of *tert*-butyl 2,2,2-trichloroacetimidate (76.0 mg, 348 µmol, 1.5 eq.) in anhydrous $Et_2O$ (500 µL) was added to a solution of the above-prepared carboxylic acid in anhydrous THF (1.0 mL). The reaction mixture was stirred for 16 h before the mixture was diluted with EtOAc and washed with water, sat. $NaHCO_3$ solution and brine. The organic layer was dried over $MgSO_4$ and concentrated in vacuo. The crude product was purified by automated reversed phase column chromatography (C18 spherical, $H_2O$:MeCN 10% to 90% MeCN) to give *tert*-butyl ester **SI-8** (100 mg, 223 µmol, 96%) as a colorless oil.

**TLC: $R_f$ (SI-8)** = 0.45 (silica, PE:EtOAc 6:4)

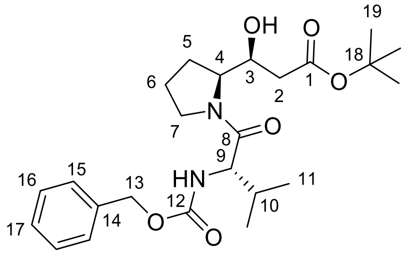

**SI-8**

**$^1$H-NMR** (400 MHz, $CDCl_3$): δ = 7.39 – 7.29 (m, 5 H, 15-H, 16-H, 17-H), 5.51 (d, $^3J_{NH,9}$ = 9.2 Hz, 1 H, 12-NH), 5.08 (m, 2 H, 13-H), 4.42 – 4.27 (m, 3 H, 4-H, 9-H, 3-OH), 3.99 (m, 1 H, 3-H), 3.83 (ddd, $^2J_{7',7}$ = 10.1 Hz, $^3J_{7',6}$ = 6.6 Hz, $^3J_{7',6'}$ = 6.6 Hz, 1 H, 7-H'), 3.52 (ddd, $^2J_{7,7'}$ = 10.3 Hz, $^3J_{7,6}$ = 6.6 Hz, $^3J_{7,6'}$ = 6.6 Hz, 1 H, 7-H), 2.45 (dd, $^2J_{2',2}$ = 15.5 Hz, $^3J_{2',3}$ = 3.5 Hz, 1 H, 2-H'), 2.35 (dd, $^2J_{2,2'}$ = 15.5 Hz, $^3J_{2,3}$ = 8.4 Hz, 1 H, 2-H), 2.09 – 1.95 (m, 3 H, 5-H', 6-H', 10-H), 1.90 (m, 1 H, 6-H), 1.73 (m, 1 H, 5-H), 1.46 (s, 9 H, 19-H), 1.03 (d, $^3J_{11',10}$ = 6.7 Hz, 3 H, 11'-H), 0.94 (d, $^3J_{11,10}$ = 6.7 Hz, 3 H, 11-H).

**$^{13}$C-NMR** (100 MHz, $CDCl_3$): δ = 173.6 (s, C-8), 171.2 (s, C-1), 156.4 (s, C-12), 136.3 (s, C-14), 128.5 (d, C-16), 128.1 (d, C-17), 128.0 (d, C-15), 81.0 (s, C-18), 72.3 (d, C-3), 66.9 (t, C-13), 61.9 (d, C-4), 57.6 (d, C-9), 48.0 (t, C-7), 41.2 (t, C-2), 31.5 (d, C-10), 28.1 (q, C-19), 27.4 (t, C-5), 24.5 (t, C-6), 19.5 (q, C-11'), 17.4 (q, C-11).

| | | |
|---|---|---|
| **Optical rotation:** | $[\alpha]_D^{20}$ = –50.4 (c = 1.0, $CHCl_3$) | |
| **HRMS (CI):** | calculated | found |
| $C_{24}H_{37}N_2O_6$ $[M+H]^+$: | 449.2646 | 449.2652 |

**_tert_-Butyl (_S_)-3-((_S_)-1-((_tert_-butoxycarbonyl)-L-valyl)pyrrolidin-2-yl)-3-hydroxypropanoate SI-9**

Benzyl carbamate **SI-8** (178 mg, 397 µmol), $Boc_2O$ (97 µL, 417 µmol, 1.05 eq.), and Pd/C (20.0 mg, 19.1 µmol, 10 wt% Pd, 5 mol%) were suspended in EtOAc (4.0 mL). The resulting suspension was stirred for 18 h under an atmosphere of $H_2$ (1 atm). The reaction mixture was filtered through a plug of celite® and concentrated in vacuo. The crude was purified by column chromatography (silica, PE:EtOAc 1:1) to give dipeptide **SI-9** (160 mg, 386 µmol, 97%) as a colorless oil.

**TLC: $R_f$ (SI-9)** = 0.22 (silica, PE:EtOAc 6:4)

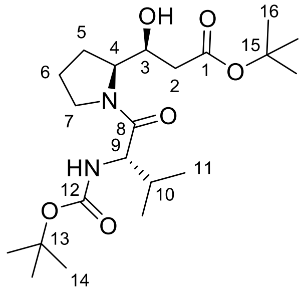


**SI-9**

**$^{1}$H-NMR** (500 MHz, $CDCl_3$): δ = 5.25 (d, $^{3}J_{NH,9}$ = 9.4 Hz, 1 H, 12-NH), 4.42 – 4.27 (m, 2 H, 4-H, 9-H), 3.99 (td, $^{3}J_{3,4}$ = 8.2 Hz, $^{3}J_{3,2}$ = 8.2 Hz, $^{3}J_{3,2'}$ = 3.5 Hz, 1 H, 3-H), 3.82 (ddd, $^{2}J_{7',7}$ = 10.1 Hz, $^{3}J_{7',6}$ = 6.7 Hz, $^{3}J_{7',6'}$ = 6.7 Hz, 1 H, 7-H'), 3.50 (ddd, $^{2}J_{7,7'}$ = 10.2 Hz, $^{3}J_{7,6}$ = 6.4 Hz, $^{3}J_{7,6'}$ = 6.4 Hz, 1 H, 7-H), 2.44 (dd, $^{2}J_{2',2}$ = 15.5 Hz, $^{3}J_{2',3}$ = 3.6 Hz, 1 H, 2-H'), 2.35 (dd, $^{2}J_{2,2'}$ = 15.5 Hz, $^{3}J_{2,3}$ = 8.4 Hz, 1 H, 2-H), 2.09 – 1.95 (m, 3 H, 5-H', 6-H', 10-H), 1.90 (m, 1 H, 6-H), 1.71 (m, 1 H, 5-H), 1.46 (s, 9 H, 16-H), 1.43 (s, 9 H, 14-H), 1.01 (d, $^{3}J_{11',10}$ = 6.7 Hz, 3 H, 11'-H), 0.93 (d, $^{3}J_{11,10}$ = 6.7 Hz, 3 H, 11-H).

**$^{13}$C-NMR** (100 MHz, $CDCl_3$): δ = 174.2 (s, C-8), 171.4 (s, C-1), 156.0 (s, C-12), 81.1 (s, C-15), 79.7 (s, C-13), 72.5 (d, C-3), 62.1 (d, C-4), 57.2 (d, C-9), 48.1 (t, C-7), 41.3 (t, C-2), 31.7 (d, C-10), 28.5 (q, C-14), 28.2 (q, C-16), 27.6 (t, C-5), 24.7 (t, C-6), 19.6 (q, C-11'), 17.5 (q, C-11).

| | | |
|---|---|---|
| **Optical rotation:** | $[\alpha]_D^{20} = -48.8$ (c = 1.0, $CHCl_3$) | |
| **HRMS (CI):** | calculated | found |
| $C_{21}H_{39}O_6N_2$ $[M+H]^+$: | 415.2803 | 415.2799 |

**(*S*)-3-(*tert*-Butoxy)-1-((*S*)-1-((*tert*-butoxycarbonyl)-L-valyl)pyrrolidin-2-yl)-3-oxopropyl(((9*H*-fluoren-9-yl)methoxy)carbonyl)-L-methioninate SI-10**

Dipeptide **SI-9** (139 mg, 335 µmol), Fmoc-Met-OH (374 mg, 1.01 mmol, 3.0 eq.), and DMAP (123 mg, 1.01 mmol, 3.0 eq.) were dissolved in anhydrous $CH_2Cl_2$ (3.0 mL). The resulting solution was cooled to 0 °C before DCC (208 mg, 1.01 mmol, 3.0 eq.) was added in one portion. The reaction mixture was stirred for 4 h while slowly reaching room temperature. After the addition of MeCN (4.0 mL), most of the urea was removed by filtration. The filtrate was diluted with EtOAc (50 mL) and washed with sat. $NH_4Cl$ solution, sat. $NaHCO_3$ solution and brine. Drying over $MgSO_4$ and concentration in vacuo followed by column chromatography (silica, PE:EtOAc 75:25) led to depsipeptide **SI-10** (219 mg, 285 µmol, 85%) as a white solid.

**TLC: $R_f$ (SI-10)** = 0.20 (silica, PE:EtOAc 7:3)

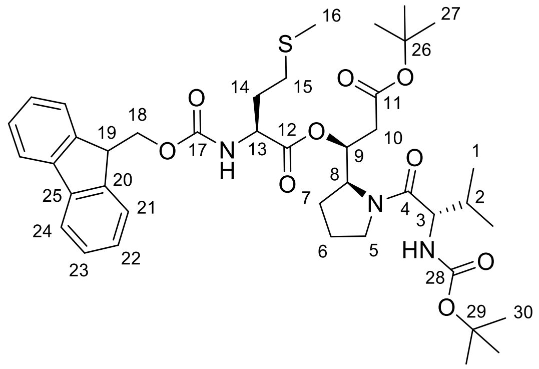


**SI-10**

**$^1$H-NMR** (500 MHz, $CDCl_3$): δ = 7.77 (d, $^3J_{24,23}$ = 7.5 Hz, 2 H, 24-H), 7.62 (m, 2 H, 21-H), 7.41 (dd, $^3J_{23,24}$ = 7.5 Hz, $^3J_{23,22}$ = 7.5 Hz, 2 H, 23-H), 7.32 (dd, $^3J_{22,21}$ = 7.4 Hz, $^3J_{22,23}$ = 7.4 Hz, 2 H, 22-H), 5.61 (ddd, $^3J_{9,8}$ = 8.9 Hz, $^3J_{9,10}$ = 4.2 Hz, $^3J_{9,10'}$ = 4.2 Hz, 1 H, 9-H), 5.53 (d, $^3J_{NH,13}$ = 8.2 Hz, 1 H, 17-NH), 5.28 (d, $^3J_{NH,3}$ = 9.2 Hz, 1 H, 28-NH), 4.53 (m, 1 H, 8-H), 4.48 (m, 1 H, 13-H), 4.44 (dd, $^2J_{18',18}$ = 10.5 Hz, $^3J_{18',19}$ = 7.3 Hz, 1 H, 18-H'), 4.37 (dd, $^2J_{18,18'}$ = 10.9 Hz, $^3J_{18,19}$ = 7.0 Hz, 1 H, 18-H), 4.31 (dd, $^3J_{3,NH}$ = 9.2 Hz, $^3J_{3,2}$ = 6.0 Hz, 1 H, 3-H), 4.22 (dd, $^3J_{19,18}$ = 7.1 Hz, $^3J_{19,18'}$ = 7.1 Hz, 1 H, 19-H), 3.77 (dt, $^2J_{5',5}$ = 9.9 Hz, $^3J_{5',6}$ = 6.8 Hz, 1 H, 5-H'), 3.41 (dt, $^2J_{5,5'}$ = 10.1 Hz, $^3J_{5,6}$ = 6.7 Hz, 1 H, 5-H), 2.58 – 2.48 (m, 3 H, 10-H', 15-H), 2.43 (dd, $^2J_{10,10'}$ = 15.9 Hz, $^3J_{10,9}$ = 9.0 Hz, 1 H, 10-H), 2.17 (m, 1 H, 14-H'), 2.10 (s, 3 H, 16-H), 2.00 – 1.84 (m, 5 H, 2-H, 6-H, 6-H', 7-H', 14-H), 1.79 (m, 1 H, 7-H), 1.42 (s, 9 H, 30-H), 1.41 (s, 9 H, 27-H), 0.97 (d, $^3J_{1',2}$ = 6.7 Hz, 3 H, 1'-H), 0.88 (d, $^3J_{1,2}$ = 6.8 Hz, 3 H, 1-H).

**$^{13}$C-NMR** (125 MHz, $CDCl_3$): δ = 172.5 (s, C-4), 170.6 (s, C-12), 169.4 (s, C-11), 156.0 (s, C-28), 155.9 (s, C-17), 143.9 (s, C-20'), 143.8 (s, C-20), 141.43 (s, C-25), 141.40 (s, C-25'), 127.9 (d, C-23), 127.21 (d, C-22), 127.18 (d, C-22'), 125.2 (d, C-21), 125.1 (d, C-21'), 120.12 (d, C-24), 120.09 (d, C-24'), 81.4 (s, C-26), 79.6 (s, C-29), 73.3 (d, C-9), 67.1 (t, C-18), 58.1 (d, C-8), 57.1 (d, C-3), 53.5 (d, C-13), 47.9 (t, C-5), 47.3 (d, C-19), 37.5 (t, C-10), 32.3 (t, C-14), 31.6 (d, C-2), 30.0 (t, C-15), 28.5 (q, C-27), 28.1 (q, C-30), 26.4 (t, C-7), 24.7 (t, C-6), 19.8 (q, C-1'), 17.2 (q, C-1), 15.5 (q, C-16).

| | | |
|---|---|---|
| **Optical rotation:** | $[\alpha]_D^{20} = -20.9$ (c = 1.0, $CHCl_3$) | |
| **HRMS (CI):** | calculated | found |
| $C_{41}H_{58}O_9N_3S$ $[M+H]^+$: | 768.3888 | 768.3894 |

**Methyl (*S*)-3-((*S*)-1-((*tert*-butoxycarbonyl)-L-isoleucyl)pyrrolidin-2-yl)-3-hydroxypropanoate SI-12**

β-Hydroxyester **7** (479 mg, 1.75 mmol) was dissolved in anhydrous $CH_2Cl_2$ (4.0 mL) and cooled to 0 °C. 4.0 M HCl in 1,4-dioxane (4.38 mL, 17.5 mmol, 10 eq.) was added, and the resulting solution was stirred for 5 h while slowly reaching room temperature. The reaction mixture was concentrated in vacuo to give the crude amine as hydrochloride.

A solution of the above-prepared hydrochloride salt and Boc-Ile-OH (445 mg, 1.93 mmol, 1.1 eq.) in anhydrous $CH_2Cl_2$ (18 mL) was cooled to 0 °C. NMM (616 µL, 5.60 mmol, 3.2 eq.) and HATU (732 mg, 1.93 mmol, 1.1 eq.) were added, and the resulting yellow solution was stirred for 18 h while slowly reaching room temperature. The reaction mixture was diluted with EtOAc and washed with 1.0 M $HCl_{aq}$, sat. $NaHCO_3$ solution and brine. The organic layer was dried over $MgSO_4$ and concentrated in vacuo. The crude was purified by column chromatography (silica, PE:EtOAc 55:45) to give dipeptide **SI-12** (642 mg, 1.58 mmol, 90%) as a colorless resin.

**TLC: $R_f$ (SI-12)** = 0.20 (silica, PE:EtOAc 6:4)

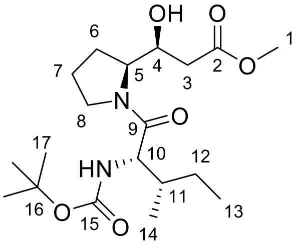


**SI-12**

**$^1$H-NMR** (500 MHz, $CDCl_3$): δ = 5.17 (d, $^3J_{NH,10}$ = 9.4 Hz, 1 H, 15-NH), 4.75 (d, $^3J_{OH,4}$ = 4.4 Hz, 1 H, 4-OH), 4.36 – 4.27 (m, 2 H, 5-H, 10-H), 4.02 (m, 1 H, 4-H), 3.89 (m, 1 H, 8-H‘), 3.72 (s, 3 H, 1-H), 3.51 (dt, $^2J_{8,8'}$ = 10.2 Hz, $^3J_{8,7}$ = 6.7 Hz, 1 H, 8-H), 2.54 (dd, $^2J_{3',3}$ = 15.2 Hz, $^3J_{3',4}$ = 3.6 Hz, 1 H, 3-H‘), 2.46 (dd, $^2J_{3,3'}$ = 15.2 Hz, $^3J_{3,4}$ = 8.5 Hz, 1 H, 3-H), 2.05 – 1.97 (m, 2 H, 6-H‘, 7-H‘), 1.90 (m, 1 H, 7-H), 1.77 – 1.66 (m, 2 H, 6-H, 11-H), 1.56 (m, 1 H, 12‘-H), 1.43 (s, 9 H, 17-H), 1.14 (m, 1 H, 12-H), 0.98 (d, $^3J_{14,11}$ = 6.7 Hz, 3 H, 14-H), 0.90 (t, $^3J_{13,12}$ = 7.4 Hz, 3 H, 13-H).

**$^{13}$C-NMR** (100 MHz, $CDCl_3$): δ = 174.7 (s, C-9), 172.3 (s, C-2), 155.9 (s, C-15), 79.8 (s, C-16), 72.8 (d, C-4), 62.4 (d, C-5), 56.7 (d, C-10), 52.0 (q, C-1), 48.3 (t, C-8), 40.3 (t, C-3), 38.2 (d, C-11), 28.5 (q, C-17), 27.9 (t, C-6), 24.7 (t, C-7), 24.3 (t, C-12), 15.6 (q, C-14), 11.4 (q, C-13).

| | | |
|---|---|---|
| **Optical rotation:** | $[\alpha]_D^{20}$= –63.6 (c = 1.0, $CHCl_3$) | |
| **HRMS (CI):** | calculated | found |
| $C_{19}H_{35}O_6N_2$ $[M+H]^+$: | 387.2490 | 387.2496 |

##### ***tert*-Butyl (*S*)-3-((*S*)-1-((*tert*-butoxycarbonyl)-L-isoleucyl)pyrrolidin-2-yl)-3-hydroxypropanoate SI-13**

0.20 M $LiOH_{aq.}$ (7.97 mL, 1.59 mmol, 1.1 eq.) was added to a 0 °C cold solution of dipeptide **SI-12** (560 mg, 1.45 mmol) in THF (7.0 mL). The resulting solution was stirred for 2 h while slowly reaching room temperature. Another portion of 0.20 M $LiOH_{aq.}$ (2.90 mL, 0.58 mmol, 0,4 eq.) was added, and the reaction mixture was stirred for two more hours. The reaction mixture was acidified with 1.0 M $HCl_{aq.}$ and extracted with EtOAc. The combined organic layers were dried over $MgSO_4$ and concentrated in vacuo to give the crude carboxylic acid.

A solution of *tert*-butyl 2,2,2-trichloroacetimidate (380 mg, 1.74 mmol, 1.2 eq.) in anhydrous $Et_2O$ (3.5 mL) was added to a solution of the above-prepared carboxylic acid (540 mg, 1.45 mmol) in anhydrous THF (7.0 mL). The reaction mixture was stirred for 3 h before another portion of *tert*-butyl 2,2,2-trichloroacetimidate (95.0 mg, 0.435 mmol, 0.3 eq.) was added. After one more hour, the mixture was diluted with EtOAc, washed with water, sat. $NaHCO_3$ and brine. The organic layer was dried over $MgSO_4$ and concentrated in vacuo. The crude product was purified by automated reversed phase column chromatography (C18, $H_2O$:MeCN 10% to 90% MeCN) to give *tert*-butyl ester **SI-13** (430 mg, 1.01 mmol, 70%, 80% brsm) as a colorless oil.

**TLC: $R_f$ (SI-13)** = 0.31 (silica, PE:EtOAc 6:4)

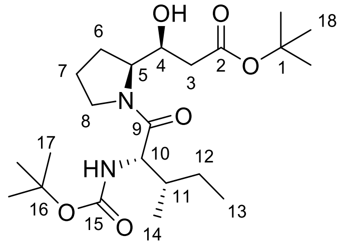


**SI-13**

**$^1$H-NMR** (400 MHz, $CDCl_3$): δ = 5.18 (d, $^3J_{NH,10}$ = 9.4 Hz, 1 H, 15-NH), 4.47 (s, 1 H, 4-OH), 4.36 – 4.28 (m, 2 H, 5-H, 10-H), 4.02 (ddd, $^3J_{4,3}$ = 8.0 Hz, $^3J_{4,5}$ = 8.0 Hz, $^3J_{4,3'}$ = 3.5 Hz, 1 H, 4-H), 3.86 (dt, $^2J_{8',8}$ = 10.0 Hz, $^3J_{8',7}$ = 6.7 Hz, 1 H, 8-H‘), 3.50 (dt, $^2J_{8,8'}$ = 10.3 Hz, $^3J_{8,7}$ = 6.4 Hz, 1 H, 8-H), 2.44 (dd, $^2J_{3',3}$ = 15.5 Hz, $^3J_{3',4}$ = 3.6 Hz, 1 H, 3-H‘), 2.35 (dd, $^2J_{3,3'}$ = 15.5 Hz, $^3J_{3,4}$ = 8.2 Hz, 1 H, 3-H), 2.07 – 1.93 (m, 2 H, 6-H‘, 7-H‘), 1.90 (m, 1 H, 7-H), 1.77 – 1.67 (m, 2 H, 6-H, 11-H), 1.57 (m, 1 H, 12-H‘), 1.43 (s, 9 H, 18-H), 1.43 (s, 9 H, 17-H), 1.14 (m, 1 H, 12-H), 0.98 (d, $^3J_{14,11}$ = 6.7 Hz, 3 H, 14-H), 0.90 (t, $^3J_{13,12}$ = 7.4 Hz, 3 H, 13-H).

**$^{13}$C-NMR** (100 MHz, $CDCl_3$): δ = 174.4 (s, C-9), 171.4 (s, C-2), 155.9 (s, C-15), 81.1 (s, C-1), 79.7 (s, C-16), 72.3 (d, C-4), 62.1 (d, C-5), 56.7 (d, C-10), 48.2 (t, C-8), 41.2 (t, C-3), 38.3 (d, C-11), 28.5 (q, C-17), 28.2 (q, C-18), 27.6 (t, C-6), 24.7 (t, C-7), 24.3 (t, C-12), 15.7 (q, C-14), 11.5 (q, C-13).

| | | |
|---|---|---|
| **Optical rotation:** | $[\alpha]_D^{20}$ = –54.9 (c = 1.0, $CHCl_3$) | |
| **HRMS (CI):** | calculated | found |
| $C_{22}H_{41}N_2O_6$ $[M+H]^+$: | 429.2959 | 429.2957 |

##### (*S*)-3-(*tert*-Butoxy)-1-((*S*)-1-((*tert*-butoxycarbonyl)-L-isoleucyl)pyrrolidin-2-yl)-3-oxopropyl (((9*H*-fluoren-9-yl)methoxy)carbonyl)-L-methioninate 8

A solution of Fmoc-Met-OH (208 mg, 560 µmol, 3.0 eq.), *tert*-butyl ester **SI-13** (80.1 mg, 187 µmol), and DMAP (68.4 mg, 560 µmol, 3.0 eq.) in anhydrous $CH_2Cl_2$ (3.0 mL) was cooled to 0 °C. DCC (116 mg, 560 µmol, 3.0 eq.) was added, and the resulting solution was stirred for 4 h while slowly reaching room temperature. The reaction mixture was diluted with MeCN (2.0 mL) and filtrated. The residue was washed with MeCN (3 x 2.0 mL), and the combined organic layers were diluted with EtOAc before being washed with sat. $NH_4Cl$ solution, sat. $NaHCO_3$ solution and brine. The organic layer was dried over $MgSO_4$ and concentrated in vacuo. The crude was purified by column chromatography (silica, PE:EtOAc 7:3) to give compound **8** (133 mg, 170 µmol, 91%) as a white foam.

**TLC: R$_f$ (8)** = 0.51 (silica, PE:EtOAc 1:1)

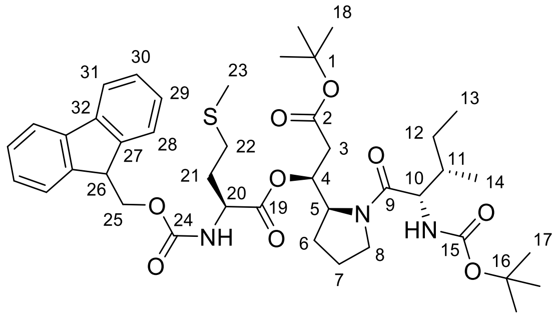


**8**

**$^1$H-NMR** (500 MHz, $CDCl_3$): δ = 7.78 (m, 2 H, 28-H), 7.62 (m, 2 H, 30-H), 7.33 (m, 2 H, 29-H), 7.33 (m, 2 H, 31-H), 5.64 (ddd, $^3J_{4,5}$ = 8.8 Hz, $^3J_{4,3}$ = 4.3 Hz, $^3J_{4,3'}$ = 4.3 Hz, 1 H, 4-H), 5.52 (d, $^3J_{NH,20}$ = 8.1 Hz, 1 H, 24-NH), 5.23 (d, $^3J_{NH,10}$ = 9.4 Hz, 1 H, 15-NH), 4.53 (m, 1 H, 5-H), 4.48 (m, 1 H, 20-H), 4.45 (dd, $^2J_{25',25}$ = 10.6 Hz, $^3J_{25',26}$ = 7.3 Hz, 1 H, 25-H), 4.37 (dd, $^2J_{25,25'}$ = 10.7 Hz, $^3J_{25,26}$ = 6.7 Hz, 1 H, 25-H), 4.32 (dd, $^3J_{10,NH}$ = 9.3 Hz, $^3J_{10,11}$ = 6.0 Hz, 1 H, 10-H), 4.22 (dd, $^3J_{26,25}$ = 7.2 Hz, $^3J_{26,25'}$ = 7.2 Hz, 1 H, 26-H), 3.81 (dt, $^2J_{8',8}$ = 9.8 Hz, $^3J_{8',7}$ = 7.2 Hz, 1 H, 8-H'), 3.42 (dt, $^2J_{8,8'}$ = 10.1 Hz, $^3J_{8,7}$ = 6.8 Hz, 1 H, 8-H), 2.54 (m, 2 H, 22-H), 2.45 (m, 2 H, 3-H), 2.17 (m, 1 H, 21-H'), 2.10 (s, 3 H, 23-H), 2.02 – 1.87 (m, 4 H, 6-H', 7-H, 21-H), 1.82 (m, 1 H, 6-H), 1.69 (m, 1 H, 11-H), 1.51 (m, 1 H, 12-H'), 1.44 – 1.39 (m, 18 H, 17-H, 18-H), 1.10 (m, 1 H, 12-H), 0.96 (d, $^3J_{14,11}$ = 6.6 Hz, 3 H, 14-H), 0.87 (t, $^3J_{13,12}$ = 7.4 Hz, 3 H, 13-H).

**$^{13}$C-NMR** (125 MHz, $CDCl_3$): δ = 172.5 (s, C-19), 170.4 (s, C-29), 169.2 (s, C-2), 155.8 (s, C-24), 155.7 (s, C-15), 143.8 (s, C-27'), 143.6 (s, C-27), 141.3 (s, C-32), 141.3 (s, C-32'), 127.7 (d, C-30), 127.1 (d, C-29'), 127.0 (d, C-29), 125.1 (d, C-28'), 125.0 (d, C-28), 120.0 (d, C-31), 120.0 (d, C-31'), 81.3 (s, C-16), 79.5 (s, C-1), 72.9 (d, C-4), 66.9 (t, C-25), 57.9 (d, C-5), 56.6 (d, C-10), 53.3 (d, C-20), 47.9 (t, C-8), 47.1 (d, C-26), 38.1 (d, C-11), 37.2 (t, C-3), 32.1 (t, C-21), 29.8 (t, C-22), 28.3 (q, C-17), 27.9 (q, C-18), 26.2 (t, C-6), 24.6 (t, C-7), 23.9 (t, C-12), 15.8 (q, C-14), 15.4 (q, C-23), 11.4 (q, C-13).

| | | |
|---|---|---|
| **Optical rotation:** | $[\alpha]_D^{20}$= −21.2 (c = 1.0, $CHCl_3$) | |
| **HRMS (CI):** | calculated | found |
| $C_{42}H_{60}N_3O_9S$ $[M+H]^+$: | 782.4045 | 782.4065 |

**Ethyl 2-((*S*)-1-((*S*)-2-((*tert*-butoxycarbonyl)amino)-5-(((2,2,2-trichloroethoxy)carbonyl)amino) pentanoyl)pyrrolidin-2-yl)thiazole-4-carboxylate SI-15**

Thiazole **9** (200 mg, 613 µmol) was dissolved in anhydrous $CH_2Cl_2$ (1.5 mL) and cooled to 0 °C. 4.0 M HCl in 1,4-dioxane (1.53 mL, 6.13 mmol, 10 eq.) was added, and the resulting solution was stirred for 2 h while slowly reaching room temperature. Another portion of 4.0 M HCl in 1,4-dioxane (0.31 mL, 1.23 mmol, 2.0 eq.) was added, and the stirring continued for 1 h. The reaction mixture was concentrated in vacuo to give the crude amine as hydrochloride.

A solution of the above-prepared hydrochloride salt and Boc-Orn(Troc)-OH **SI-19** (275 mg, 674 µmol, 1.1 eq.) in anhydrous $CH_2Cl_2$ (6.0 mL) was cooled to 0 °C. NMM (216 µL, 1.96 mmol, 3.2 eq.) and HATU (256 mg, 674 µmol, 1.1 eq.) were added, and the resulting yellow solution was stirred for 18 h while slowly reaching room temperature. The reaction mixture was diluted with EtOAc and washed with 1.0 M $HCl_{aq}$, sat. $NaHCO_3$ solution and brine. The organic layer was dried over $MgSO_4$ and concentrated in vacuo. The crude product was purified by automated reversed phase column chromatography (C18 spherical, $H_2O$:MeCN 10% to 90% MeCN) to give dipeptide **SI-15** (315 mg, 511 µmol, 83%) as a white foam.

**TLC: $R_f$ (SI-15)** = 0.35 (silica, DCM:MeOH 96:4)

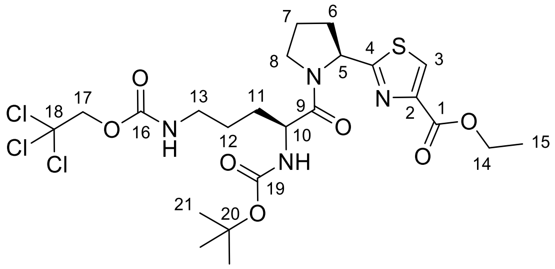


**SI-15**

**$^1$H-NMR** (500 MHz, $CDCl_3$): δ = 8.05 (s, 1 H, 3-H), 5.64 (t, $^3J_{NH,13}$ = 5.7 Hz, 1 H, 16-NH), 5.49 (t, $^3J_{5,6}$ = 5.7 Hz, 1 H, 5-H), 5.35 (d, $^3J_{NH,10}$ = 8.6 Hz, 1 H, 19-NH), 4.74 (d, $^2J_{17',17}$ = 12.0 Hz, 1 H, 17'-H), 4.69 (d, $^2J_{17,17'}$ = 12.0 Hz, 1 H, 17-H), 4.55 (dt, $^3J_{10,NH}$ = 8.4 Hz, $^3J_{10,11}$ = 4.1 Hz, 1 H, 10-H), 4.40 (q, $^3J_{14,15}$ = 7.1 Hz, 2 H, 14-H), 3.79 (m, 2 H, 8-H), 3.28 (m, 2 H, 13-H), 2.32 (m, 2 H, 6-H), 2.08 (m, 2 H, 7-H), 1.84 (m, 1 H, 11-H'), 1.73 – 1.60 (m, 3 H, 11-H, 12-H), 1.43 (s, 9 H, 21-H), 1.38 (t, $^3J_{15,14}$ = 7.1 Hz, 3 H, 15-H).

**$^{13}$C-NMR** (125 MHz, $CDCl_3$): δ = 172.9 (s, C-4), 171.5 (s, C-9), 161.4 (s, C-1), 155.6 (s, C-19), 154.9 (s, C-16), 147.0 (s, C-2), 127.4 (d, C-3), 95.9 (s, C-18), 80.0 (s, C-20), 74.5 (t, C-17), 61.6 (t, C-14), 58.9 (d, C-5), 51.6 (d, C-10), 47.4 (t, C-8), 41.1 (t, C-13), 32.3 (t, C-6), 30.3 (t, C-11), 28.5 (q, C-21), 25.2 (t, C-12), 24.7 (t, C-7), 14.5 (q, C-15).

| | | |
|---|---|---|
| **Optical rotation:** | $[\alpha]_D^{20}$ = –45.8 (c = 1.0, $CHCl_3$) | |
| **HRMS (CI):** | calculated | found |
| $C_{23}H_{34}O_7N_4Cl_3S$ $[M+H]^+$: | 615.1208 | 615.1210 |

**Ethyl 2-((*S*)-1-((*S*)-2-acetamido-5-(((2,2,2-trichloroethoxy)carbonyl)amino)pentanoyl) pyrrolidin-2-yl)thiazole-4-carboxylate 10**

Dipeptide **10** (496 mg, 805 µmol) was dissolved in anhydrous $CH_2Cl_2$ (3.0 mL) and cooled to 0 °C. 4.0 M HCl in 1,4-dioxane (2.01 mL, 8.05 mmol, 10 eq.) was added, and the resulting solution was stirred for 2 h while slowly reaching room temperature. The reaction mixture was concentrated in vacuo to give the crude amine as hydrochloride.

A solution of the above-prepared hydrochloride salt and triethylamine (236 µL, 1.69 mmol, 2.1 eq.) in anhydrous $CH_2Cl_2$ (7.0 mL) was cooled to 0 °C. Acetic anhydride (152 µL, 1.61 mmol, 2.0 eq.) was added, and the reaction mixture was stirred for 2 h while slowly reaching room temperature. After dilution with EtOAc, the mixture was washed with sat. $NH_4Cl$ solution, sat. $NaHCO_3$ solution and brine before being dried over $MgSO_4$ and concentrated in vacuo. The crude product was purified by automated reversed phase column chromatography (C18 spherical, $H_2O$:MeCN 10% to 90% MeCN) to give the acetylated dipeptide **10** (421 mg, 754 µmol, 94%) as a white foam.

**TLC: $R_f$ (10)** = 0.17 (silica, DCM:MeOH 95:5)

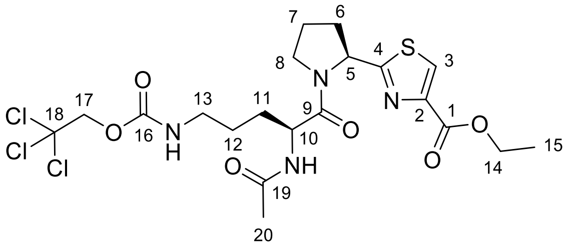

**10**

**$^1$H-NMR** (500 MHz, $CDCl_3$): δ = 8.06 (s, 1 H, 3-H), 6.62 (d, $^3J_{NH,10}$ = 8.0 Hz, 1 H, 19-NH), 5.81 (t, $^3J_{NH,13}$ = 6.1 Hz, 1 H, 16-NH), 5.46 (dd, $^3J_{5,6}$ = 7.3 Hz, $^3J_{5,6'}$ = 4.3 Hz, 1 H, 5-H), 4.86 (m, 1 H, 10-H), 4.73 (d, $^2J_{17',17}$ = 12.2 Hz, 1 H, 17-H'), 4.69 (d, $^2J_{17,17'}$ = 12.1 Hz, 1 H, 17-H), 4.39 (q, $^3J_{14,15}$ = 7.1 Hz, 2 H, 14-H), 3.81 (m, 2 H, 8-H), 3.26 (m, 2 H, 13-H), 2.31 (m, 2 H, 6-H), 2.09 (m, 2 H, 7-H), 1.96 (s, 3 H, 20-H), 1.86 (m, 1 H, 11-H'), 1.75 – 1.55 (m, 3 H, 11-H, 12-H), 1.37 (t, $^3J_{15,14}$ = 7.1 Hz, 3 H, 15-H).

**$^{13}$C-NMR** (125 MHz, $CDCl_3$): δ = 172.5 (s, C-4), 171.1 (s, C-9), 169.9 (s, C-19), 161.2 (s, C-1), 154.8 (s, C-16), 146.8 (s, C-2), 127.2 (d, C-3), 95.7 (s, C-18), 74.3 (t, C-17), 61.5 (t, C-14), 58.8 (d, C-5), 50.3 (d, C-10), 47.4 (t, C-8), 40.9 (t, C-13), 32.3 (t, C-6), 29.7 (t, C-11), 25.0 (t, C-12), 24.5 (t, C-7), 23.1 (q, C-20), 14.3 (q, C-15).

| | | |
|---|---|---|
| **Optical rotation:** | $[\alpha]_D^{20}$ = −63.6 (c = 0.5, $CHCl_3$) | |
| **HRMS (CI):** | calculated | found |
| $C_{20}H_{28}O_6N_4Cl_3S$ [M+H]$^+$: | 557.0795 | 557.0802 |

***N*-,*N*'-Bis(*tert*-Butoxycarbonyl)-*N*''-trifluoromethanesulfonyl-guanidine[14] SI-16**

Preparation according to Goodman *et al.*[14]

**TLC: $R_f$ (SI-16)** = 0.48 (silica, DCM:MeOH 98:2)

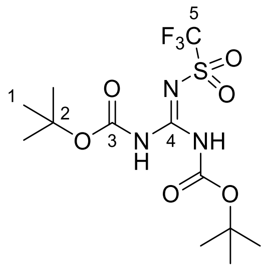

**SI-16**

**$^{1}$H-NMR** (400 MHz, DMSO-$d_6$): δ = 10.10 (s, 2 H, 3-NH), 1.54 (s, 18 H, 1-H).

**$^{13}$C-NMR** (100 MHz, DMSO-$d_6$): δ = 151.4 (s, C-3), 119.2 (q, $^{1}J_{C5,F}$ = 320.2 Hz, C-5), 86.0 (s, C-2), 27.8 (q, C-1), C-4 was not observed.

| **HRMS (CI):** | calculated | found |
|---|---|---|
| $C_{12}H_{21}F_3N_3O_6S$ [M+H]$^+$: | 392.1098 | 392.1090 |

**Melting point:** 110 – 113 °C (lit: 115 °C)[14]

**Ethyl 2-((*S*)-1-(*N*$^2$-acetyl-*N*$^ω$,*N*$^{ω'}$-bis(*tert*-butoxycarbonyl)-L-arginyl)pyrrolidin-2-yl)thiazole-4-carboxylate 11**

Acetylated dipeptide **10** (51 mg, 91.0 µmol) was dissolved in anhydrous THF (1.0 mL). After the addition of AcOH (10.5 µL, 183 µmol, 2.0 eq.) and zinc dust (120 mg, 1.83 mmol, 20 eq.), the reaction was initiated by the addition of 1,2-dibromoethane and TMS-Cl (2 µL each) and stirred at room temperature for 90 min. Sat. $NaHCO_3$ solution was added, and the mixture was extracted with $CHCl_3$/$^{i}$PrOH (3:1). After concentration in vacuo, the residue was dissolved in $CHCl_3$ and dried over $MgSO_4$. Concentration in vacuo led to the crude amine.

The above-prepared crude amine and triethylamine (14.0 µL, 101 µmol, 1.1 eq.) were dissolved in anhydrous $CH_2Cl_2$ (1.0 mL). Di-Boc-guanidyl triflate **SI-16** (39.4 mg, 101 µmol, 1.1 eq.) was added, and the reaction mixture was stirred for 16 h at room temperature. After dilution with EtOAc, the mixture was washed with sat. $NH_4Cl$ solution, sat. $NaHCO_3$ solution and brine. The organic layer was dried over $MgSO_4$ and concentrated in vacuo. The crude product was purified by automated reversed phase column chromatography (C18 spherical, $H_2O$:MeCN 10% to 90% MeCN) to give dipeptide **11** (35.0 mg, 56.0 µmol, 61%) as a white foam.

**TLC: $R_f$ (11)** = 0.44 (silica, DCM:MeOH 95:5)

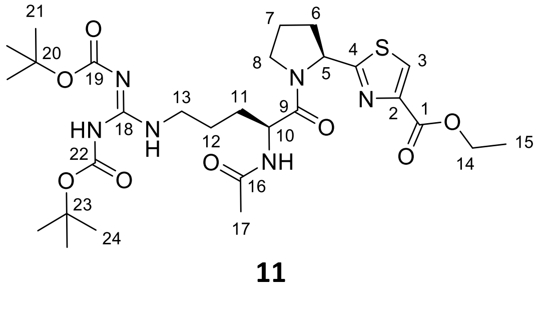

**11**

**$^1$H-NMR** (500 MHz, $CDCl_3$): δ = 11.49 (s, 1 H, 22-NH), 8.34 (t, $^3J_{NH,13}$ = 5.5 Hz, 1 H, 13-NH), 8.06 (s, 1 H, 3-H), 6.60 (d, $^3J_{NH,10}$ = 8.0 Hz, 1 H, 19-NH), 5.52 (dd, $^3J_{5,6}$ = 8.1 Hz, $^3J_{5,6'}$ = 2.8 Hz, 1 H, 5-H), 4.85 (ddd, $^3J_{10,NH}$ = 7.6 Hz, $^3J_{10,11}$ = 7.4 Hz, $^3J_{10,11'}$ = 4.6 Hz, 1 H, 10-H), 4.40 (q, $^3J_{14,15}$ = 7.1 Hz, 2 H, 14-H), 3.79 (m, 2 H, 8-H), 3.49 (m, 1 H, 13-H‘), 3.41 (m, 1 H, 13-H), 2.40 (m, 1 H, 6-H‘), 2.29 (dddd, $^2J_{6,6'}$ = 12.9 Hz, $^3J_{6,7}$ = 12.9 Hz, $^3J_{6,7'}$ = 8.1 Hz, $^3J_{6,5}$ = 8.1 Hz, 1 H, 6-H), 2.08 (m, 2 H, 7-H), 2.00 (s, 3 H, 17-H), 1.88 (m, 1 H, 11-H‘), 1.76 – 1.56 (m, 3 H, 11-H, 12-H), 1.49 (s, 9 H, 21-H), 1.47 (s, 9 H, 24-H), 1.38 (t, $^3J_{15,14}$ = 7.2 Hz, 3 H, 15-H).

**$^{13}$C-NMR** (125 MHz, $CDCl_3$): δ = 173.2 (s, C-4), 171.4 (s, C-9), 169.9 (s, C-16), 163.5 (s, C-19), 161.2 (s, C-1), 156.2 (s, C-18), 153.2 (s, C-22), 147.1 (s, C-2), 127.1 (d, C-3), 83.1 (s, C-20), 79.3 (s, C-23), 61.4 (t, C-14), 59.0 (d, C-5), 50.2 (d, C-10), 47.3 (t, C-8), 40.1 (t, C-13), 31.9 (t, C-6), 29.4 (t, C-11), 28.2 (q, C-21), 28.0 (q, C-24), 25.1 (t, C-12), 24.4 (t, C-7), 23.1 (q, C-17), 14.3 (q, C-15).

| | | |
|---|---|---|
| **Optical rotation:** | $[\alpha]_D^{20} = -43.8$ (c = 0.5, $CHCl_3$) | |
| **HRMS (CI):** | calculated | found |
| $C_{28}H_{45}N_6O_8S$ $[M+H]^+$: | 625.3014 | 625.3022 |

**(*S*)-3-(*tert*-Butoxy)-1-((*S*)-1-((*tert*-butoxycarbonyl)-L-valyl)pyrrolidin-2-yl)-3-oxopropyl (2-((*S*)-1-(N$^2$-acetyl-N$^{\omega}$,N$^{\omega'}$-bis(*tert*-butoxycarbonyl)-L-arginyl)pyrrolidin-2-yl)thiazole-4-carbonyl)-L-methioninate SI-18**

Dipeptide **11** (50.0 mg, 80.1 µmol) was dissolved in 420 µL THF and cooled to 0 °C. After the dropwise addition of 0.20 M $LiOH_{aq.}$ (420 µL, 84.2 µmol, 1.05 eq.) the resulting solution was stirred for 4 h while slowly reaching room temperature. The solution was neutralized by the addition of 1.0 M $HCl_{aq}$ (84.2 µL, 84.2 µmol, 1.05 eq.) and concentrated in vacuo to give the crude carboxylic acid.

Depsipeptide **SI-10** (67.0 mg, 87.3 µmol, 1.09 eq.) was dissolved in $CH_2Cl_2$ (0.8 mL) and cooled to 0 °C. Diethyl amine (182 µL, 1.75 mmol, 20 eq.) was added, and the mixture was stirred for 3 h while slowly reaching room temperature. The solution was concentrated in vacuo to give the crude amine.

The crude amine and the crude carboxylic acid were dissolved in anhydrous DMF (0.8 mL) before being cooled to 0 °C. NMM (9.69 µL, 88.0 µmol, 1.1 eq.) and HATU (30.4 mg, 80.0 µmol, 1.0 eq.) were added subsequently. The resulting solution was stirred for 16 h while slowly reaching room temperature. After dilution with EtOAc, the mixture was washed with 5 wt% $LiCl_{aq}$, sat. $NH_4Cl$ solution, sat. $NaHCO_3$ solution and brine. The organic layer was dried over $MgSO_4$ and concentrated in vacuo. The crude product was purified by automated reversed phase column chromatography (C18 spherical, $H_2O$:MeCN 10% to 90% MeCN) to give protected pseudotetraivprolid D **SI-18** (31.4 mg, 28.0 µmol, 35%) as a white foam.

**LC-MS: $t_R$ (SI-18)** = 1.59 min (short method)

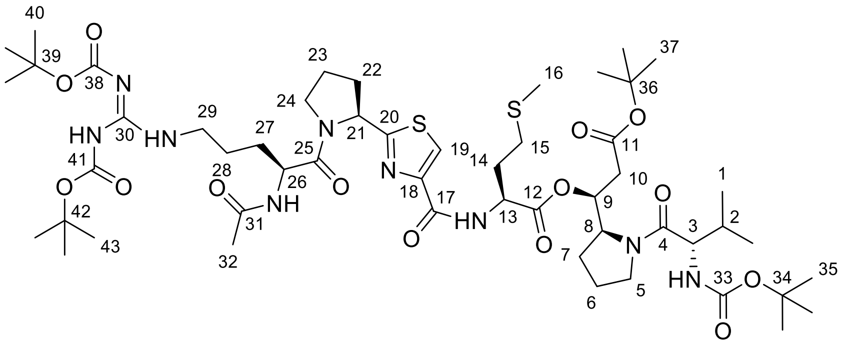

**SI-18**

**$^1$H-NMR** (500 MHz, $CDCl_3$): δ = 11.50 (s, 1 H, 41-NH), 8.35 (t, $^3J_{NH,29}$ = 5.5 Hz, 1 H, 29-NH), 7.98 (s, 1 H, 19-H), 7.80 (d, $^3J_{NH,13}$ = 8.4 Hz, 1 H, 17-NH), 6.54 (d, $^3J_{NH,26}$ = 8.3 Hz, 1 H, 31-NH), 5.65 (ddd, $^3J_{9,10}$ = 8.5 Hz, $^3J_{9,10'}$ = 4.1 Hz, $^3J_{9,8}$ = 4.1 Hz, 1 H, 9-H), 5.46 (dd, $^3J_{21,22}$ = 7.9 Hz, $^3J_{21,22'}$ = 2.8 Hz, 1 H, 21-H), 5.29 (d, $^3J_{NH,3}$ = 9.2 Hz, 1 H, 33-NH), 4.90 – 4.81 (m, 2 H, 13-H, 26-H), 4.56 (ddd, $^3J_{8,7'}$ = 8.4 Hz, $^3J_{8,7}$ = 3.9 Hz, $^3J_{8,9}$ = 3.9 Hz, 1 H, 8-H), 4.31 (dd, $^3J_{3,NH}$ = 9.2 Hz, $^3J_{3,2}$ = 5.3 Hz, 1 H, 3-H), 3.83 (m, 1 H, 24-H'), 3.79 – 3.72 (m, 2 H, 5-H', 24-H), 3.50 (m, 1 H, 29-H), 3.47 – 3.40 (m, 2 H, 5-H, 29-H), 2.58 (m, 2 H, 15-H), 2.50 (dd, $^3J_{10',10}$ = 16.1 Hz, $^3J_{10',9}$ = 4.7 Hz, 1 H, 10-H'), 2.44 (dd, $^2J_{10,10'}$ = 16.1 Hz, $^3J_{10,9}$ = 8.7 Hz, 1 H, 10-H), 2.35 (m, 1 H, 22-H'), 2.31 – 2.24 (m, 2 H, 14-H, 22-H), 2.13 – 1.79 (m, 9 H, 2-H, 6-H, 7-H, 14-H, 23-H, 27-H'), 2.11 (s, 3 H, 16-H), 2.03 (s, 3 H, 32-H), 1.72 – 1.65 (m, 3 H, 27-H, 28-H), 1.49 (s, 9 H, 40-H), 1.48 (s, 9 H, 43-H), 1.41 (s, 9 H, 37-H), 1.40 (s, 9 H, 35-H), 0.97 (d, $^3J_{1',2}$ = 6.8 Hz, 3 H, 1'-H), 0.86 (d, $^3J_{1,2}$ = 6.8 Hz, 3 H, 1-H).

**$^{13}$C-NMR** (125 MHz, $CDCl_3$): δ = 172.9 (s, C-20), 172.4 (s, C-4), 171.7 (s, C-25), 170.7 (s, C-12), 170.1 (s, C-31), 169.3 (s, C-11), 163.6 (s, C-38), 160.7 (s, C-17), 156.4 (s, C-30), 156.0 (s, C-33), 153.4 (s, C-41), 149.3 (s, C-18), 123.6 (d, C-19), 83.4 (s, C-39), 81.3 (s, C-42), 79.6 (s, C-36), 79.4 (s, C-34), 73.5 (d, C-9), 59.0 (d, C-21), 57.9 (d, C-8), 57.0 (d, C-3), 51.6 (d, C-13), 50.5 (d, C-26), 48.0 (t, C-5), 47.5 (t, C-24), 40.3 (t, C-29), 37.3 (t, C-10), 32.2 (t, C-14), 31.8 (t, C-22), 31.6 (d, C-2), 30.1 (t, C-15), 29.5 (t, C-27), 28.4 (q, C-43), 28.4 (q, C-40), 28.2 (q, C-37), 28.1 (q, C-35), 26.4 (t, C-7), 25.5 (t, C-28), 24.8 (t, C-6), 24.6 (t, C-23), 23.4 (q, C-32), 19.8 (q, C-1), 17.1 (q, C-1'), 15.6 (q, C-16).

**Optical rotation:** $[\alpha]_D^{20}$ = –80.9 (c = 1.0, MeOH)

| | | |
|---|---|---|
| **HRMS (ESI):** | calculated | found |
| $C_{52}H_{86}N_9O_{14}S_2$ $[M+H]^+$: | 1124.5730 | 1124.5742 |

**(*S*)-3-(*tert*-Butoxy)-1-((*S*)-1-((*tert*-butoxycarbonyl)-L-isoleucyl)pyrrolidin-2-yl)-3-oxopropyl (2-((*S*)-1-($N^2$-acetyl-$N^{\omega}$,$N^{\omega'}$-bis(*tert*-butoxycarbonyl)-L-arginyl)pyrrolidin-2-yl)thiazole-4-carbonyl)-L-methioninate 12**

Dipeptide **11** (24.0 mg, 38.4 µmol) was dissolved in THF (250 µL) and cooled to 0 °C. After the dropwise addition of 0.20 M $LiOH_{aq}$ (202 µL, 40.4 µmol, 1.05 eq.), the resulting solution was stirred for 4 h while slowly reaching room temperature. The solution was neutralized by the addition of

1.0 M $HCl_{aq}$ (40.4 µL, 40.4 µmol, 1.05 eq.) and concentrated in vacuo to give the crude carboxylic acid.

Fmoc amine **8** (30.0 mg, 38.4 µmol, 1.0 eq.) was dissolved in $CH_2Cl_2$ (500 µL) and cooled to 0 °C. Diethyl amine (80.0 µL, 767 µmol, 20 eq.) was added, and the mixture was stirred for 4 h while slowly reaching room temperature. Another portion of diethyl amine (80.0 µL, 767µmol, 20 eq.) was added, and stirring continued for another hour. The reaction mixture was concentrated in vacuo to give the crude amine.

The above-prepared crude amine and carboxylic acid were dissolved in anhydrous $CH_2Cl_2$ (400 µL) before being cooled to 0 °C. NMM (4.54 µL, 41.0 µmol, 1.05 eq.) and HBTU (14.5 mg, 38.4 µmol) were added subsequently. The resulting solution was stirred for 16 h while slowly reaching room temperature. After dilution with EtOAc, the mixture was washed with sat. $NH_4Cl$ solution, sat. $NaHCO_3$ solution and brine. The organic layer was dried over $MgSO_4$ and concentrated in vacuo. The crude product was purified by column chromatography (silica, $CH_2Cl_2$:MeOH 97:3) followed by preparative HPLC ($H_2O$:MeCN 10 % to 100% MeCN) to give protected pseudotetraivprolid B **12** (20.4 mg, 17.9 µmol, 47%) as a white foam.

**TLC: $R_f$ (12)** = 0.78 (silica, DCM:MeOH 95:5)

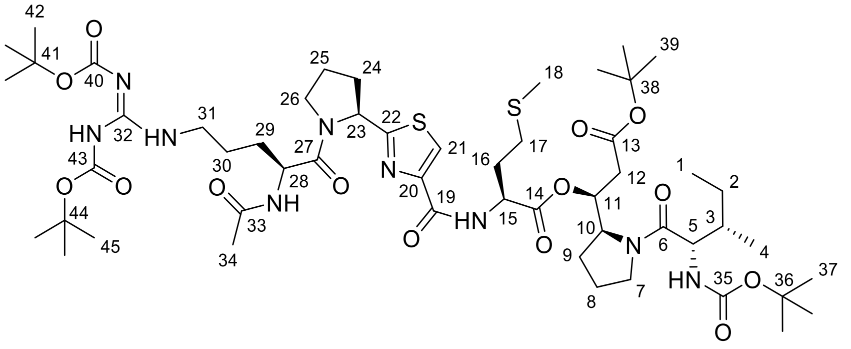

**12**

**$^1$H-NMR** (500 MHz, $CDCl_3$): δ = 11.50 (s, 1 H, 43-NH), 8.36 (t, $^3J_{NH,31}$ = 5.5 Hz, 1 H, 31-NH), 7.99 (s, 1 H, 21-H), 7.81 (d, $^3J_{NH,15}$ = 8.3 Hz, 1 H, 19-NH), 6.53 (d, $^3J_{NH,28}$ = 8.2 Hz, 1 H, 33-NH), 5.69 (ddd, $^3J_{11,12}$ = 8.7 Hz, $^3J_{11,12'}$ = 4.3 Hz, $^3J_{11,10}$ = 4.3 Hz, 1 H, 11-H), 5.47 (dd, $^3J_{23,24}$ = 7.9 Hz, $^3J_{23,24'}$ = 2.8 Hz, 1 H, 23-H), 5.24 (d, $^3J_{NH,5}$ = 9.3 Hz, 1 H, 35-NH), 4.90 – 4.82 (m, 2 H, 15-H, 28-H), 4.56 (ddd, $^3J_{10,9'}$ = 8.4 Hz, $^3J_{10,11}$ = 4.0 Hz, $^3J_{10,9}$ = 4.0 Hz, 1 H, 10-H), 4.33 (dd, $^3J_{5,NH}$ = 9.3 Hz, $^3J_{5,3}$ = 6.0 Hz, 1 H, 5-H), 3.88 – 3.72 (m, 3 H, 7-H‘, 26-H), 3.55 – 3.40 (m, 3 H, 7-H, 31-H), 2.58 (m, 2 H, 17-H), 2.47 (m, 2 H, 12-H), 2.31 (m, 2 H, 24-H), 2.15 – 1.82 (m, 9 H, 8-H, 9-H, 16-H, 25-H, 29-H'), 2.11 (s, 3 H, 18-H), 2.03 (s, 3 H, 34-H), 1.74 – 1.63 (m, 4 H, 3-H, 29-H, 30-H), 1.52 (m, 1 H, 2-H‘), 1.49 (s, 9 H, 42-H), 1.47 (s, 9 H, 45-H), 1.41 (s, 9 H, 39-H), 1.40 (s, 9 H, 37-H), 1.09 (m, 1 H, 2-H), 0.95 (d, $^3J_{4,3}$ = 6.7 Hz, 3 H, 4-H), 0.87 (t, $^3J_{1,2}$ = 7.3 Hz, 3 H, 1-H).

**$^{13}$C-NMR** (125 MHz, $CDCl_3$): δ = 172.7 (s, C-22), 172.4 (s, C-6), 171.5 (s, C-27), 170.5 (s, C-14), 170.0 (s, C-33), 169.1 (s, C-13), 163.5 (s, C-40), 160.5 (s, C-19), 156.3 (s, C-32), 155.8 (s, C-35), 153.3 (s, C-43), 149.1 (s, C-20), 123.5 (d, C-21), 83.2 (s, C-41), 81.1 (s, C-44), 79.5 (s, C-38), 79.4 (s, C-36),

73.1 (d, C-11), 58.8 (d, C-23), 57.7 (d, C-10), 56.5 (d, C-5), 51.4 (d, C-15), 50.3 (d, C-28), 47.9 (t, C-7), 47.3 (t, C-26), 40.1 (t, C-31), 38.2 (d, C-3), 36.9 (t, C-12), 32.1 (t, C-24), 31.6 (t, C-16), 30.0 (t, C-17), 29.4 (t, C-29), 28.30 (q, C-45), 28.26 (q, C-42), 28.0 (q, C-39), 27.9 (q, C-37), 26.2 (t, C-9), 25.3 (t, C-30), 24.7 (t, C-8), 24.4 (t, C-25), 23.9 (t, C-2), 23.2 (q, C-34), 15.9 (q, C-4), 15.5 (q, C-18), 11.5 (q, C-1).

**Optical rotation:** $[\alpha]_D^{20}$ = –62.9 (c = 1.0, MeOH)

| **HRMS (ESI):** | calculated | found |
|---|---|---|
| $C_{53}H_{88}N_9O_{14}S_2$ $[M+H]^+$: | 1138.5887 | 1138.5901 |

**Pseudotetraivprolid D 5a**

Protected pseudotetraivprolid D **SI-18** (31.4 mg, 28.0 µmol) was dissolved in anhydrous $CH_2Cl_2$ (150 µL). The deprotection mixture TFA:$H_2O$:TIPS (150 µL) in a ratio of 185:5:10 was added, and the solution was stirred for 160 min. The reaction was dried in high vacuum, and the residue was purified by automated reversed phase column chromatography (C18, 0.1% $HCOOH_{aq}$:MeCN 10% to 90% MeCN) followed by preparative HPLC (0.1% $HCOOH_{aq}$:MeCN 0% to 50% MeCN) to give pseudeotetraivprolid D **5a** (19.0 mg, 23.4 µmol, 85%) as an amorphous solid.

**LC-MS: $t_R$ (5a)** = 0.58 min (short method)

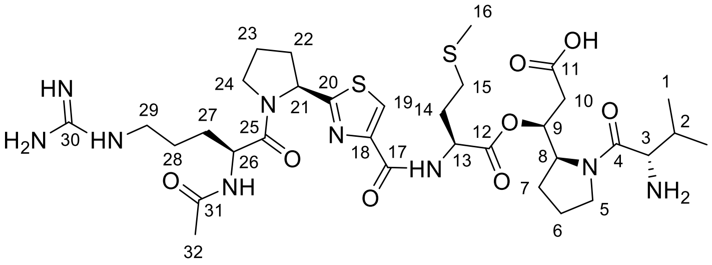

**5a**

**$^1$H-NMR** (500 MHz, $D_2O$): δ = 8.16 (s, 1 H, 19-H), 5.58 (ddd, $^3J_{9,10}$ = 7.0 Hz, $^3J_{9,10'}$ = 6.7 Hz, $^3J_{9,8}$ = 3.7 Hz, 1 H, 9-H), 5.41 (dd, $^3J_{21,22}$ = 8.2 Hz, $^3J_{21,22'}$ = 3.0 Hz, 1 H, 21-H), 4.80 (m, 1 H, 13-H), 4.64 (dd, $^3J_{26,27}$ = 8.3 Hz, $^3J_{26,27'}$ = 5.1 Hz, 1 H, 26-H), 4.51 (ddd, $^3J_{8,7}$ = 8.2 Hz, $^3J_{8,9}$ = 3.8 Hz, $^3J_{8,7'}$ = 3.8 Hz, 1 H, 8-H), 4.16 (d, $^3J_{3,2}$ = 5.1 Hz, 1 H, 3-H), 3.88 (m, 2 H, 24-H), 3.68 (dt, $^2J_{5,5'}$ = 10.3 Hz, $^3J_{5,6}$ = 7.2 Hz, 1 H, 5-H), 3.43 (dt, $^2J_{5',5}$ = 10.2 Hz, $^3J_{5',6}$ = 7.0 Hz, 1 H, 5-H'), 3.21 (m, 2 H, 29-H), 2.73 – 2.55 (m, 4 H, 10-H, 15-H), 2.41 (m, 1 H, 22-H), 2.26 (m, 1 H, 14-H), 2.21 – 2.11 (m, 5 H, 2-H, 14-H', 22-H', 23-H), 2.09 (s, 3 H, 16-H), 2.01 (s, 3 H, 32-H), 2.01 – 1.80 (m, 5 H, 6-H, 7-H, 27-H), 1.76 – 1.60 (m, 3 H, 27-H', 28-H), 0.99 (d, $^3J_{1',2}$ = 7.0 Hz, 3 H, 1'-H), 0.90 (d, $^3J_{1,2}$ = 6.9 Hz, 3 H, 1-H).

**$^{13}$C-NMR** (125 MHz, $D_2O$): δ = 175.0 (s, C-11), 174.71 (s, C-20), 174.68 (s, C-31), 173.3 (s, C-25), 172.6 (s, C-12), 169.8 (s, C-4), 163.5 (s, C-17), 157.3 (s, C-30), 148.1 (s, C-18), 125.7 (d, C-19), 74.0 (d, C-9), 59.9 (d, C-21), 59.1 (d, C-8), 57.6 (d, C-3), 52.5 (d, C-13), 51.9 (d, C-26), 49.0 (t, C-5), 48.3 (t, C-24), 41.0 (t, C-29), 36.8 (t, C-10), 32.5 (t, C-22), 30.0 (t, C-15), 29.81 (t, C-14), 29.78 (d, C-2),

28.0 (t, C-27), 26.5 (t, C-7), 24.9 (t, C-28), 24.53 (t, C-23), 24.50 (t, C-6), 22.0 (q, C-32), 18.7 (q, C-1), 16.6 (q, C-1‘), 14.6 (q, C-16).

**Selected rotamer signals:**

**$^1$H-NMR** (500 MHz, $D_2O$): δ = 8.23 (s, 1 H, 19-H), 5.50 (m, 1 H, 21-H), 5.28 (m, 1 H, 9-H), 4.57 (dd, $^3J_{26,27}$ = 9.0 Hz, $^3J_{26,27'}$ = 4.7 Hz, 1 H, 26-H), 4.28 (d, $^3J_{3,2}$ = 5.1 Hz, 1 H, 3-H), 3.78 (m, 2 H, 24-H), 3.58 (m, 2 H, 5-H), 3.17 (m, 2 H, 29-H), 2.50 (m, 1 H, 22-H), 1.75 (s, 1 H, 32-H).

| | | |
|---|---|---|
| **Optical rotation:** | $[\alpha]_D^{20}$= –63.4 (c = 0.5, DMSO) | |
| **HRMS (ESI):** | calculated | found |
| $C_{33}H_{54}N_9O_8S_2$ $[M+H]^+$: | 768.3531 | 768.3533 |

**Pseudotetraivprolid B 4a**

Protected pseudotetraivprolid B **12** (17.1 mg, 15.0 µmol) was dissolved in anhydrous $CH_2Cl_2$ (80 µL). The deprotection mixture TFA:$H_2O$:TIPS (80 µL) in a ratio of 185:5:10 was added, and the solution was stirred for 160 min. The reaction was dried in high vacuum, and the residue was purified by automated reversed phase column chromatography (C18, 0.1% $HCOOH_{aq}$: MeCN 10% to 90% MeCN) followed by preparative HPLC (0.1% $HCOOH_{aq}$:MeCN 0% to 50% MeCN) to give pseudeotetraivprolid B **4a** (4.3 mg, 5.5 µmol, 37%) as an amorphous solid well as the corresponding *tert*-butyl ester (4.0 mg, 4.8 µmol, 32%) as an amorphous solid, due to incomplete cleavage.

LC-MS: $t_R$ (**4a**) = 0.58 min (short method)

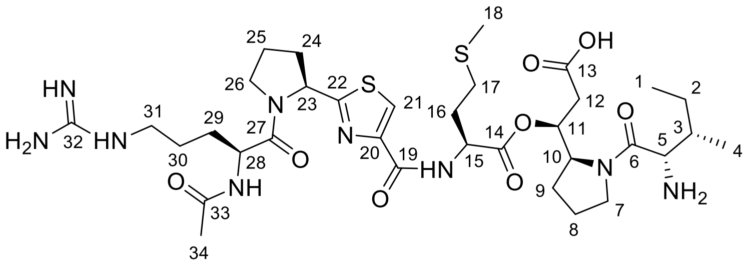


**4a**

**$^1$H-NMR** (500 MHz, $D_2O$): δ = 8.17 (s, 1 H, 21-H), 5.60 (m, 1 H, 11-H), 5.42 (dd, $^3J_{23,24}$ = 8.3 Hz, $^3J_{23,24'}$ = 3.0 Hz, 1 H, 23-H), 4.81 (m, 1 H, 15-H), 4.65 (dd, $^3J_{28,29}$ = 8.3 Hz, $^3J_{28,29'}$ = 5.1 Hz, 1 H, 28-H), 4.52 (ddd, $^3J_{10,9}$ = 8.3 Hz, $^3J_{10,11}$ = 4.0 Hz, $^3J_{10,9'}$ = 4.0 Hz, 1 H, 10-H), 4.20 (d, $^3J_{5,3}$ = 4.7 Hz, 1 H, 5-H), 3.89 (m, 2 H, 26-H), 3.69 (dt, $^2J_{7,7'}$ = 10.7 Hz, $^3J_{7,8}$ = 6.8 Hz, 1 H, 7-H), 3.46 (dt, $^2J_{7',7}$ = 10.6 Hz, $^3J_{7',8}$ = 7.3 Hz, 1 H, 7-H‘), 3.22 (m, 2 H, 31-H), 2.70 – 2.56 (m, 4 H, 12-H, 17-H), 2.42 (m, 1 H, 24-H), 2.27 (m, 1 H, 16-H), 2.21 – 2.11 (m, 4 H, 16-H‘, 24-H‘, 25-H), 2.10 (s, 3 H, 18-H), 2.02 (s, 3 H, 34-H), 2.02 – 1.81 (m, 6 H, 3-H, 8-H, 9-H, 29-H), 1.76 – 1.60 (m, 3 H, 29-H‘, 30-H), 1.43 (m, 1 H, 2-H), 1.14 (m, 1 H, 2-H‘), 1.00 (d, $^3J_{4,3}$ = 7.0 Hz, 3 H, 4-H), 0.85 (t, $^3J_{1,2}$ = 7.3 Hz, 3 H, 1-H).

**$^{13}$C-NMR** (125 MHz, $D_2O$): δ = 175.0 (s, C-13), 174.84 (s, C-22), 174.81 (s, C-33), 173.4 (s, C-27), 172.7 (s, C-14), 170.0 (s, C-6), 163.6 (s, C-19), 157.4 (s, C-32), 148.2 (s, C-20), 125.8 (d, C-21), 74.0

(d, C-11), 60.1 (d, C-23), 59.2 (d, C-10), 57.4 (d, C-5), 52.6 (d, C-15), 52.0 (d, C-28), 49.1 (t, C-7), 48.4 (t, C-26), 41.2 (t, C-31), 36.7 (t, C-12), 36.5 (d, C-3), 32.6 (t, C-24), 30.1 (t, C-17), 30.0 (t, C-16), 28.2 (t, C-29), 26.5 (t, C-9), 25.0 (t, C-30), 24.7 (t, C-25), 24.6 (t, C-8), 23.9 (t, C-2), 22.1 (q, C-34), 15.4 (q, C-4), 14.8 (q, C-18), 11.4 (q, C-1).

**Selected rotamer signals:**

**$^1$H-NMR** (500 MHz, $D_2O$): δ = 8.24 (s, 1 H, 21-H), 5.50 (m, 1 H, 23-H), 4.57 (dd, $^3J_{28,29}$ = 8.9 Hz, $^3J_{28,29'}$ = 4.7 Hz, 1 H, 28-H), 4.31 (d, $^3J_{5,4}$ = 6.1 Hz, 1 H, 5-H), 3.79 (m, 1 H, 26-H), 3.59 (m, 2 H, 7-H), 3.18 (m, 2 H, 31-H), 1.76 (s, 3 H, 34-H), 0.91 (m, 3 H, 1-H).

**Optical rotation:** $[\alpha]_D^{20}$= –68.5 (c = 0.2, DMSO)

| | | |
|---|---|---|
| **HRMS (ESI):** | calculated | found |
| $C_{34}H_{56}N_9O_8S_2$ $[M+H]^+$: | 782.3688 | 782.3678 |

**(*S*)-2-((*tert*-Butoxycarbonyl)amino)-5-(((2,2,2-trichloroethoxy)carbonyl)amino)pentanoic acid SI-19**

A solution of Troc-Cl (711 µL, 5.17 mmol, 1.2 eq.) in $Et_2O$ (5.0 mL) was added to a solution of $K_2CO_3$ (1.49 g, 10.8 mmol, 2.5 eq.) and Boc-Orn-OH (1.00 g, 4.31 mmol) in $H_2O$ (20 mL). The reaction mixture was stirred for 16 h at room temperature. The aqueous layer was washed twice with $Et_2O$ (10 mL), and the organic layer was discarded. The aqueous layer was acidified with 1.0 M $HCl_{aq}$ (pH 2) before being extracted thrice with $Et_2O$ (25 mL). The combined organic layers were dried with $MgSO_4$ and concentrated in vacuo. The crude was purified by column chromatography (silica, $CH_2Cl_2$:MeOH 95:5) to give Boc-Orn(Troc)-OH **SI-19** (1.60 g, 3.93 mmol, 91%) as a white foam.

**TLC: $R_f$ (SI-19)** = 0.31 (silica, $CH_2Cl_2$:MeOH 95:5)

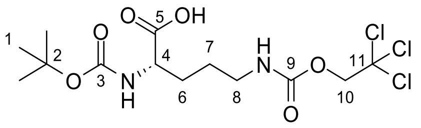


**SI-19**

**$^1$H-NMR** (500 MHz, DMSO-$d_6$): δ = 7.67 (t, $^3J_{NH,8}$ = 5.6 Hz, 1 H, 9-NH), 6.65 (m, 1 H, 3-NH), 4.77 (m, 2 H, 10-H), 3.78 (m, 1 H, 4-H), 3.00 (m, 2 H, 8-H), 1.66 (m, 1 H, 6-H), 1.55 – 1.40 (m, 3 H, 6-H‘, 7-H), 1.37 (s, 9 H, 1-H).

**$^{13}$C-NMR** (125 MHz, DMSO-$d_6$): δ = 155.3 (s, C-3), 154.4 (s, C-9), 96.4 (s, C-11), 77.8 (s, C-2), 73.3 (t, C-10), 53.9 (d, C-4), 40.4 (t, C-8), 29.0 (t, C-6), 28.3 (q, C-1), 25.9 (t, C-7), C-5 was not observed.

**Optical rotation:** $[\alpha]_D^{20}$= +5.5 (c = 1.0, $CHCl_3$)

| | | |
|---|---|---|
| **HRMS (ESI):** | calculated | found |
| $C_{13}H_{22}O_6N_2Cl_3$ $[M+H]^+$: | 407.0583 | 407.0543 |

***tert*-Butyl (*S*)-2-carbamoylpyrrolidine-1-carboxylate SI-1**

**$^{1}$H-NMR** (500 MHz, DMSO-$d_6$):

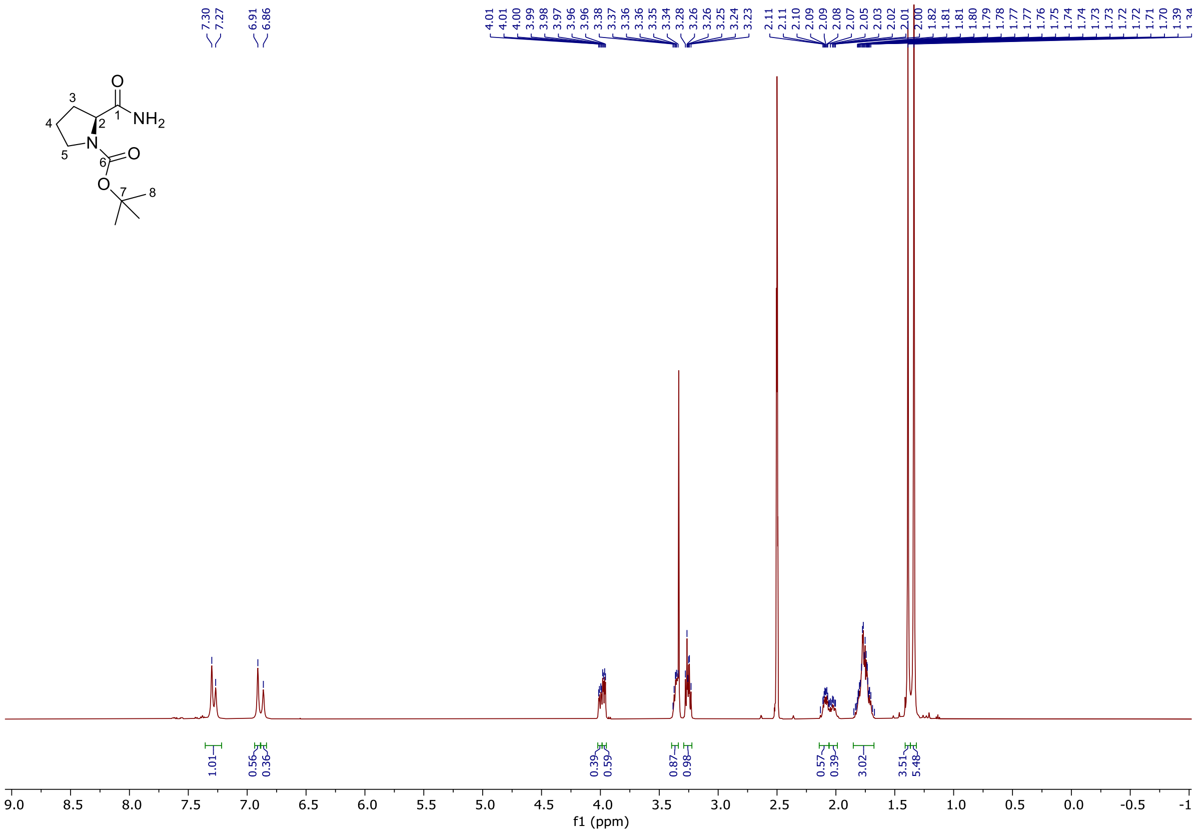


**$^{13}$C-NMR** (125 MHz, DMSO-$d_6$):

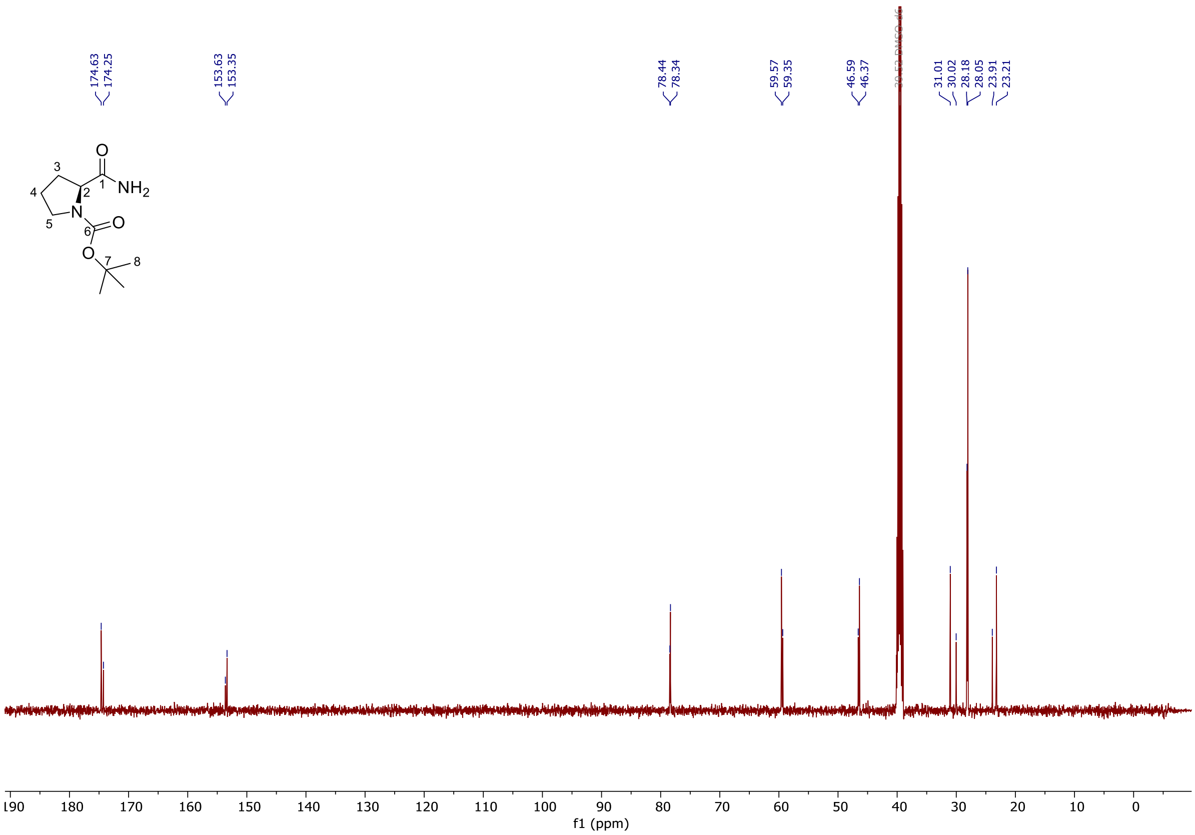

**Ethyl (*S*)-2-(1-(*tert*-butoxycarbonyl)pyrrolidin-2-yl)thiazole-4-carboxylate 9**

**[1]H-NMR** (500 MHz, $CDCl_3$):

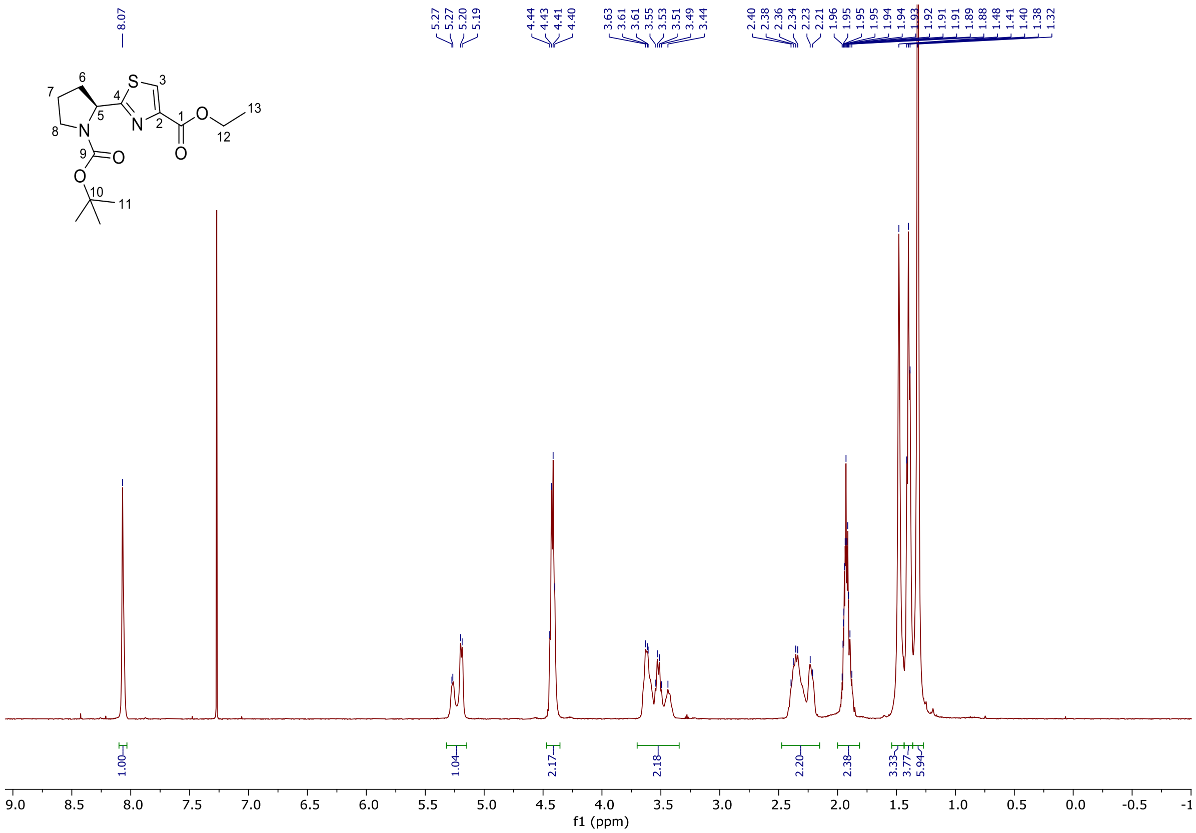


**[13]C-NMR** (125 MHz, DMSO-d$_6$, 373 K):

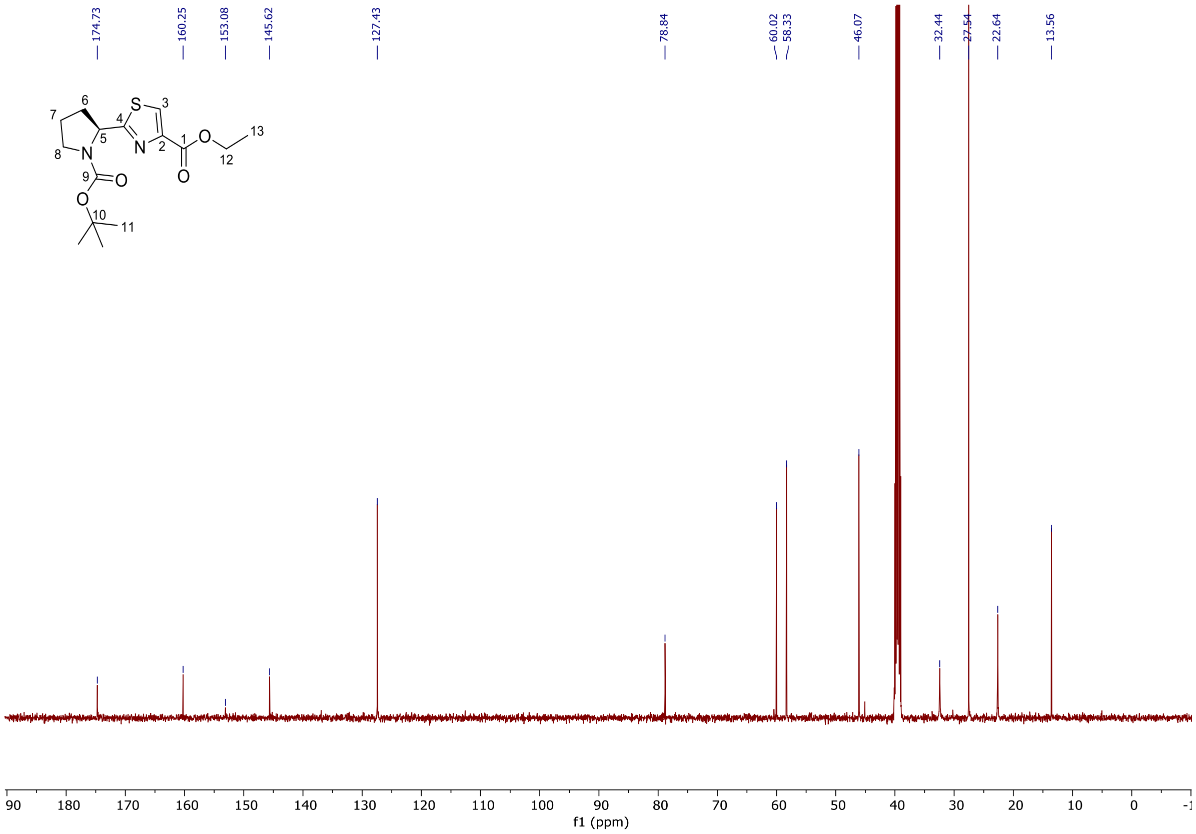

***tert*-Butyl (*S*)-2-(4-(((*S*)-1-methoxy-4-(methylthio)-1-oxobutan-2-yl)carbamoyl)thiazol-2-yl)-pyrrolidine-1-carboxylate SI-4**

**$^{1}$H-NMR** (500 MHz, DMSO-d$_6$, 373 K): (contains < 1% DCM and EtOAc as impurity)

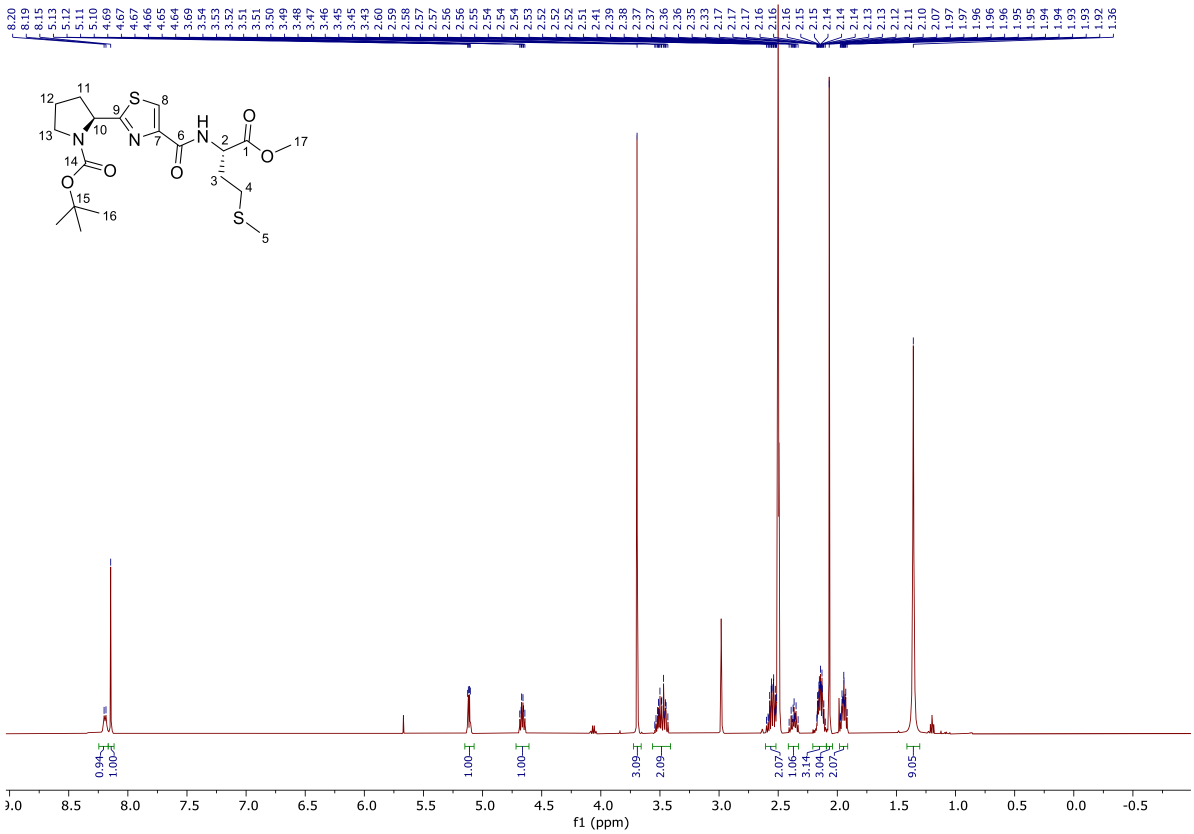


**$^{13}$C-NMR** (125 MHz, DMSO-d$_6$, 373 K):

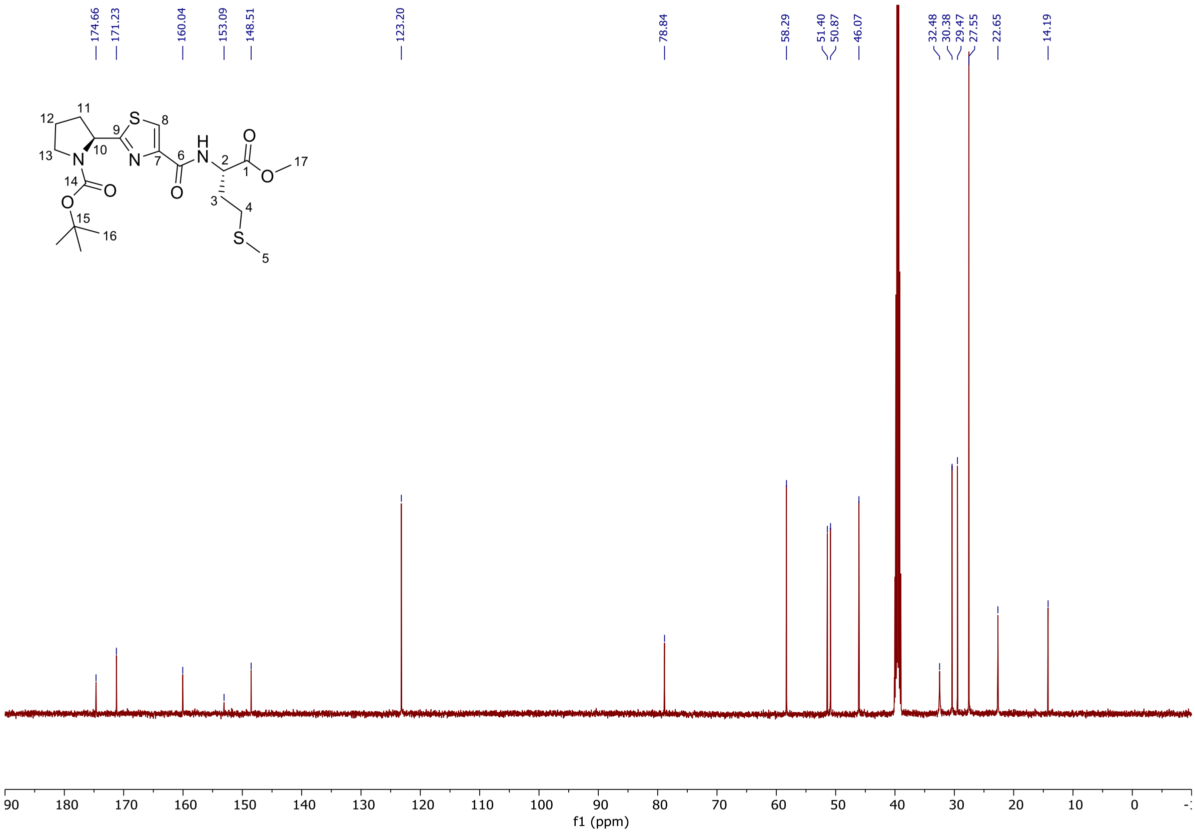

#### (2-((*S*)-1-(*tert*-Butoxycarbonyl)pyrrolidin-2-yl)thiazole-4-carbonyl)-L-methionine SI-5

**$^{1}$H-NMR** (500 MHz, DMSO-d$_6$, 373 K):

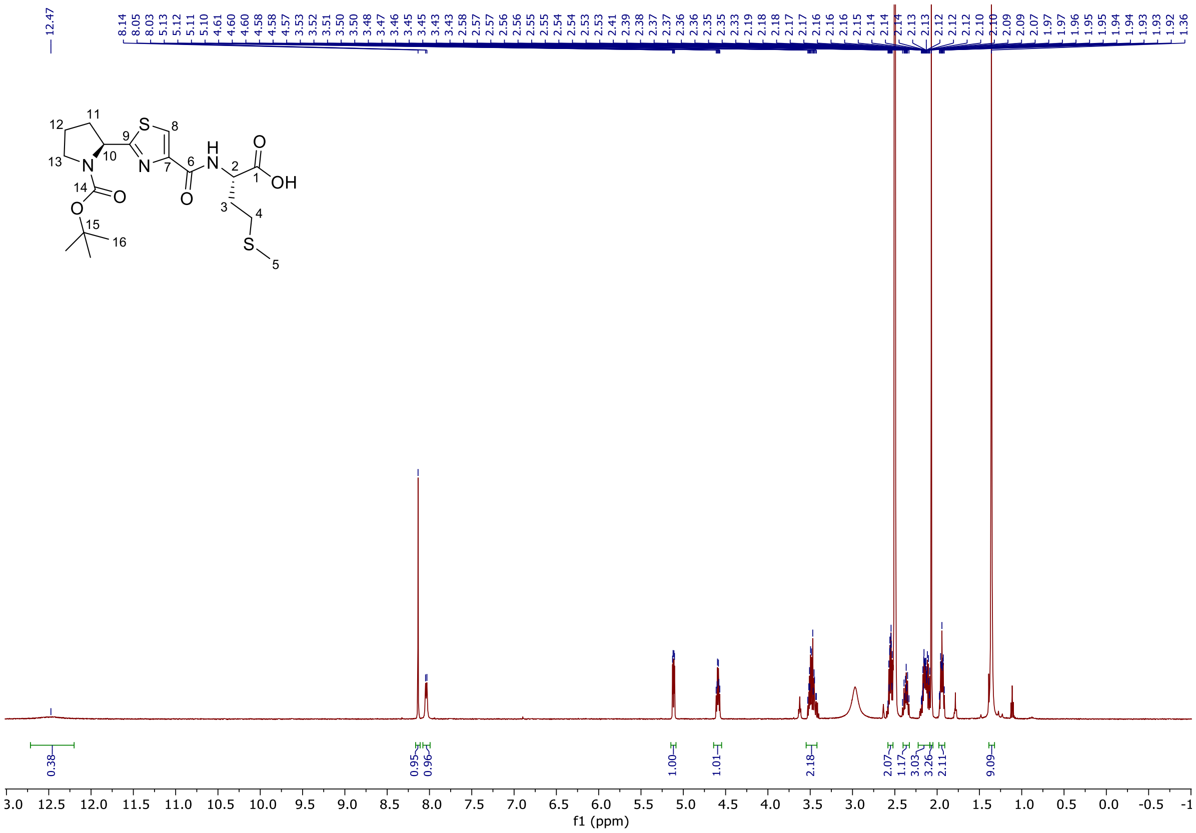

**$^{13}$C-NMR** (125 MHz, DMSO-d$_6$, 373 K):

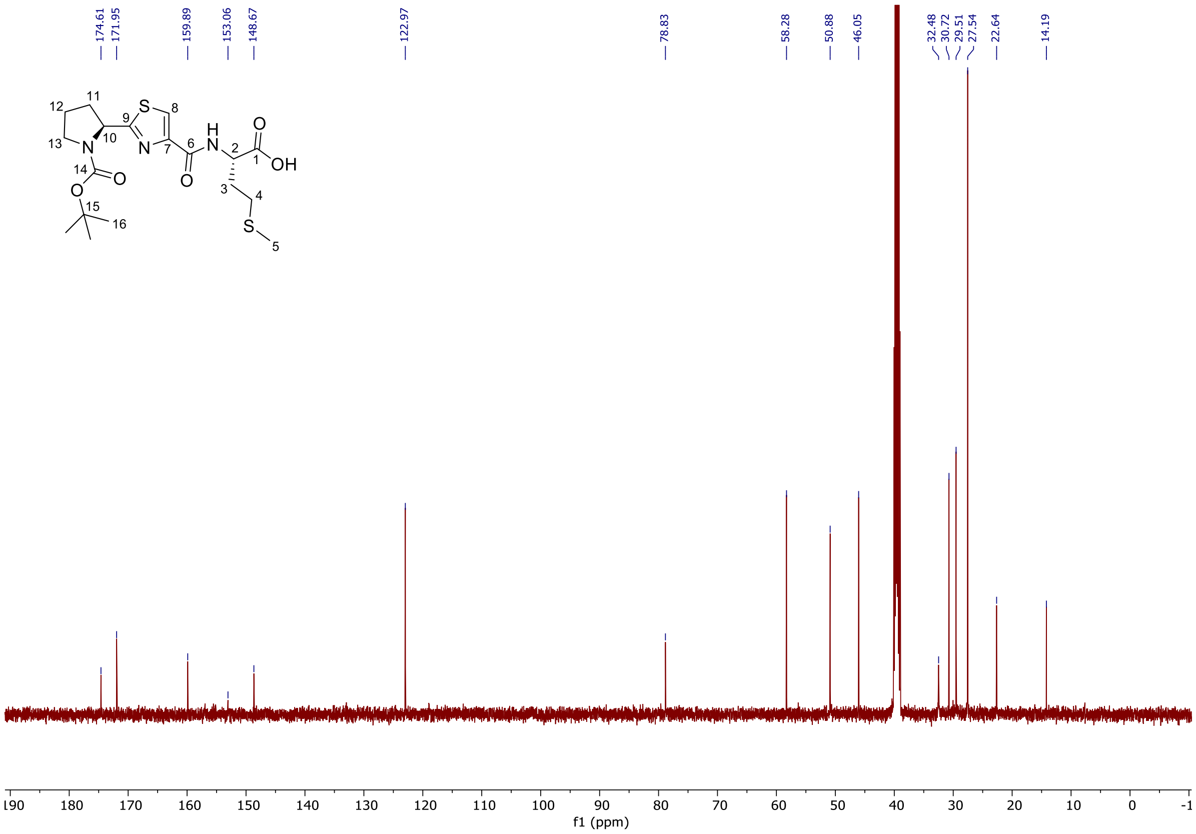

**tert-Butyl (S)-2-((S)-1-hydroxy-3-methoxy-3-oxopropyl)pyrrolidine-1-carboxylate 7**

**$^{1}$H-NMR** (500 MHz, $CDCl_3$):

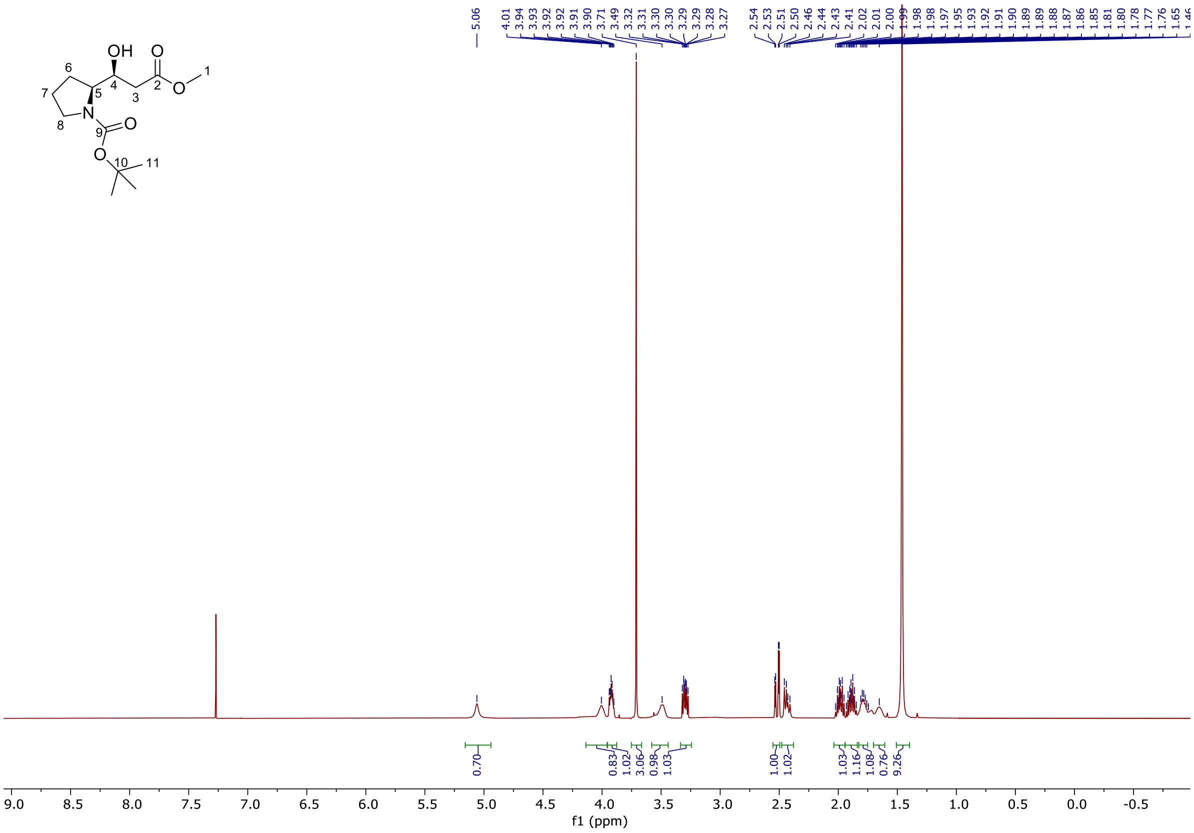


**$^{13}$C-NMR** (125 MHz, $CDCl_3$):

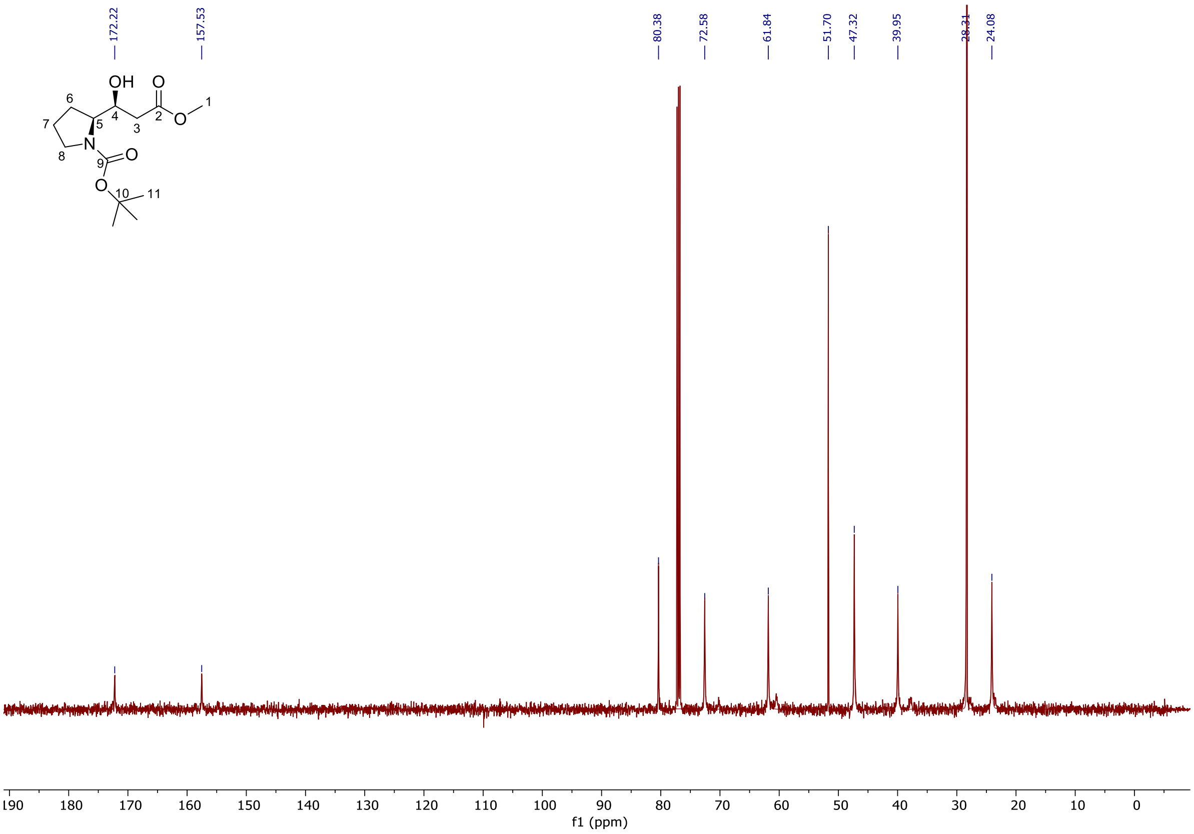

**Methyl (*S*)-3-((*S*)-1-(((benzyloxy)carbonyl)-L-valyl)pyrrolidin-2-yl)-3-hydroxypropanoate SI-6**

**$^1$H-NMR** (500 MHz, CDCl$_3$):

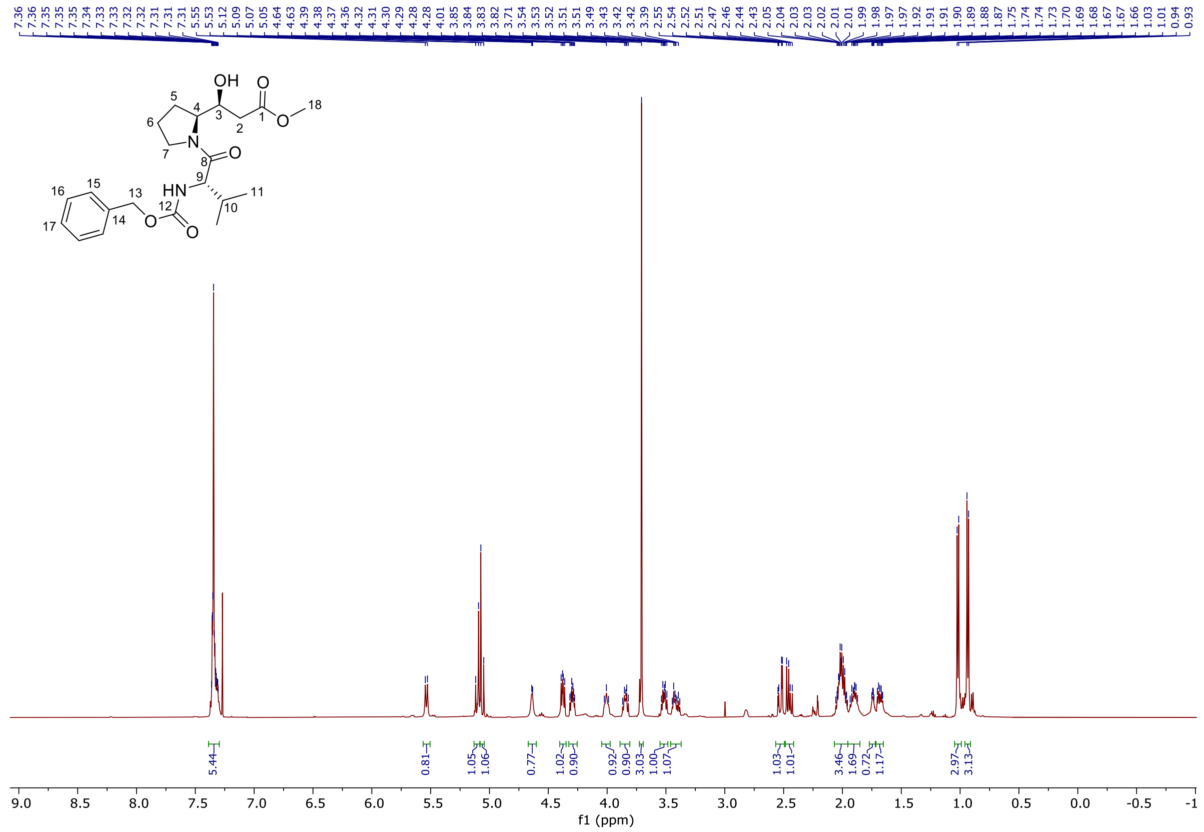


**$^{13}$C-NMR** (125 MHz, CDCl$_3$):

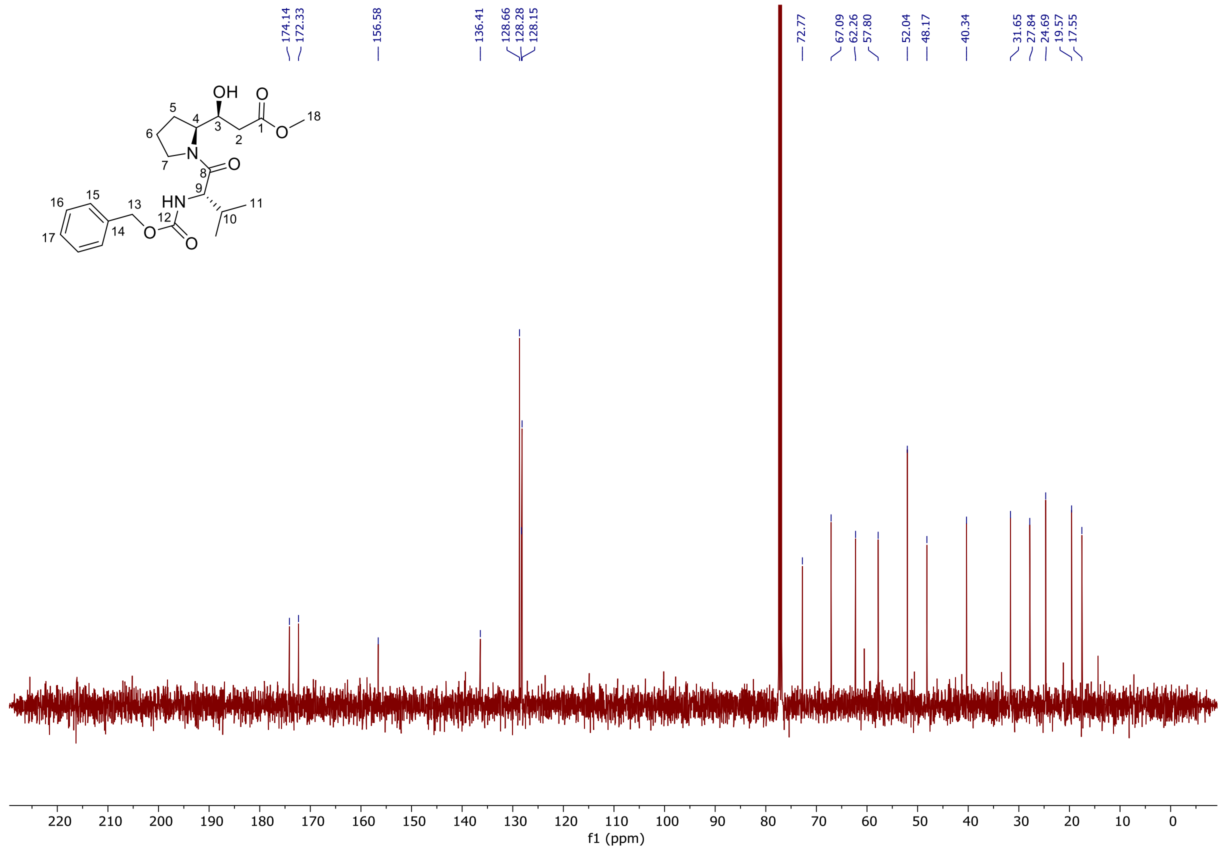

**_tert_-Butyl (_S_)-3-((_S_)-1-(((benzyloxy)carbonyl)-L-valyl)pyrrolidin-2-yl)-3-hydroxypropanoate SI-8**

**[1]H-NMR** (400 MHz, $CDCl_3$): (contains 2% DCM as impurity)

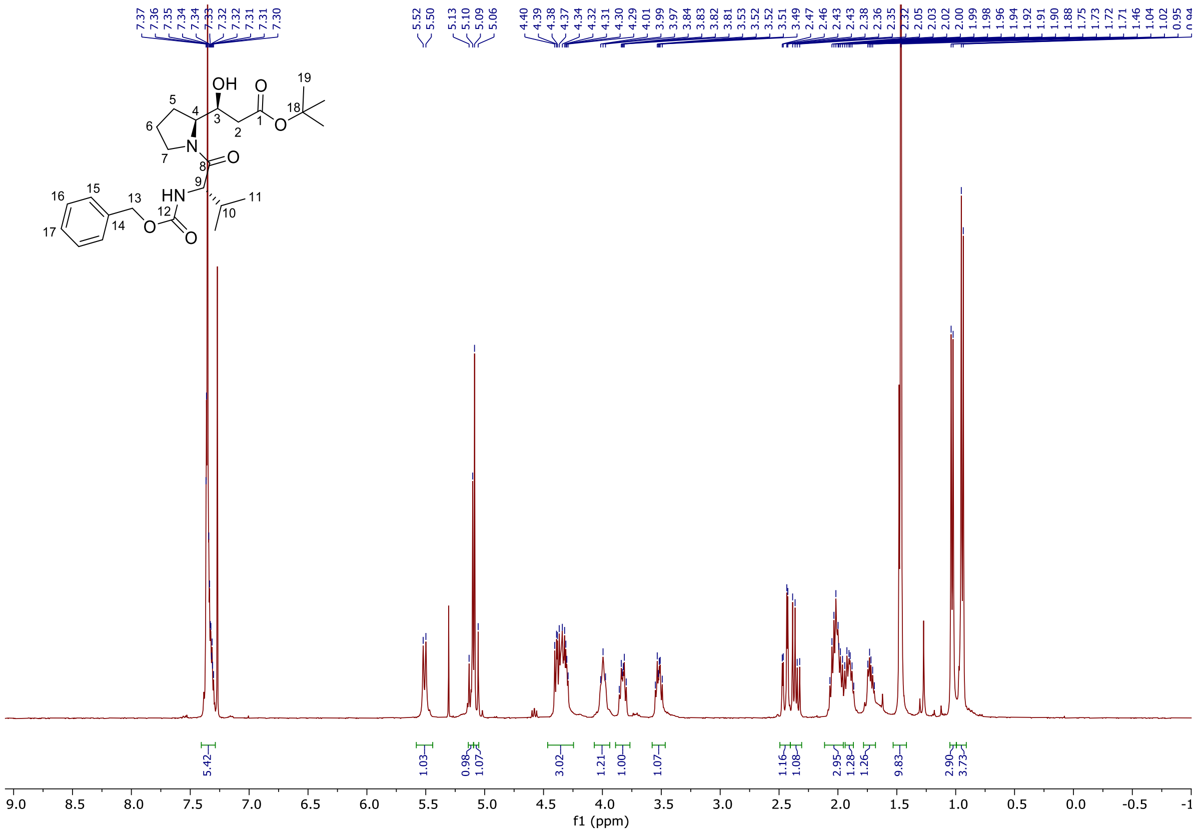

**[13]C-NMR** (100 MHz, $CDCl_3$):

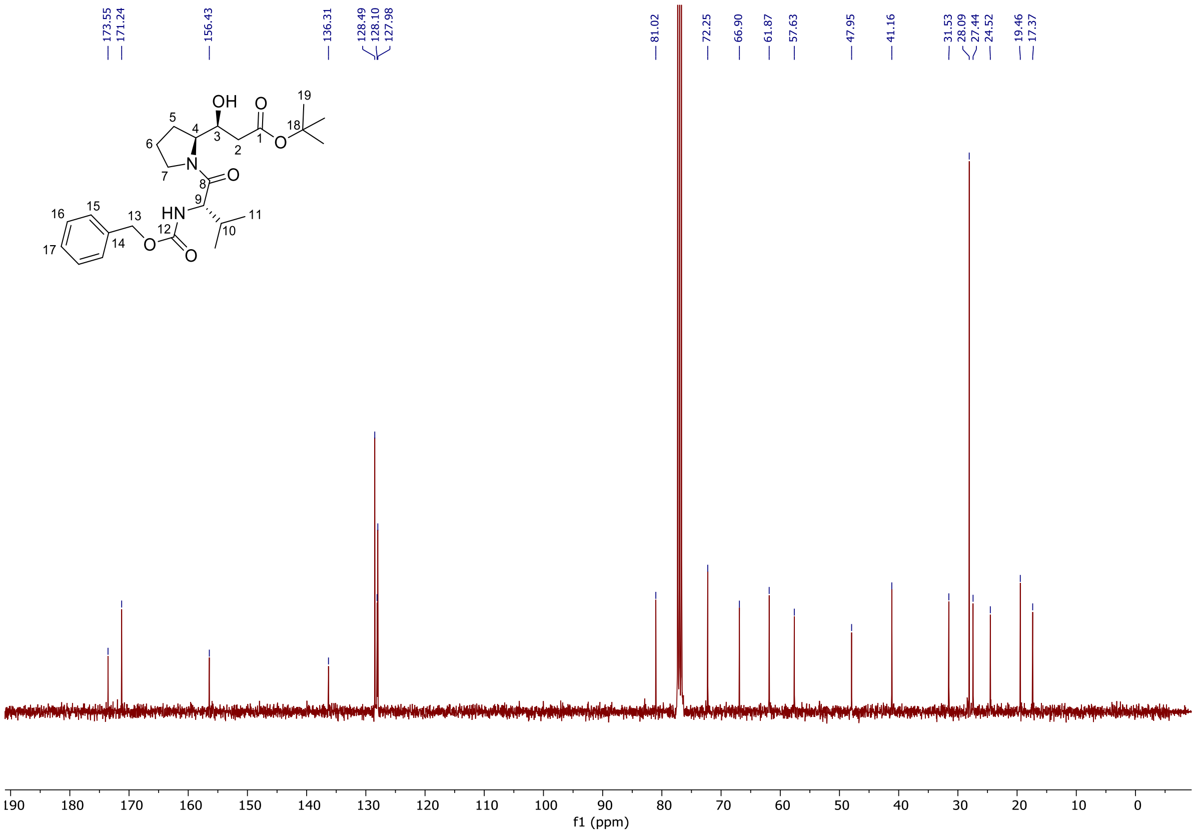

**_tert_-Butyl (_S_)-3-((_S_)-1-((_tert_-butoxycarbonyl)-L-valyl)pyrrolidin-2-yl)-3-hydroxypropanoate SI-9**

**$^{1}$H-NMR** (500 MHz, $CDCl_3$):

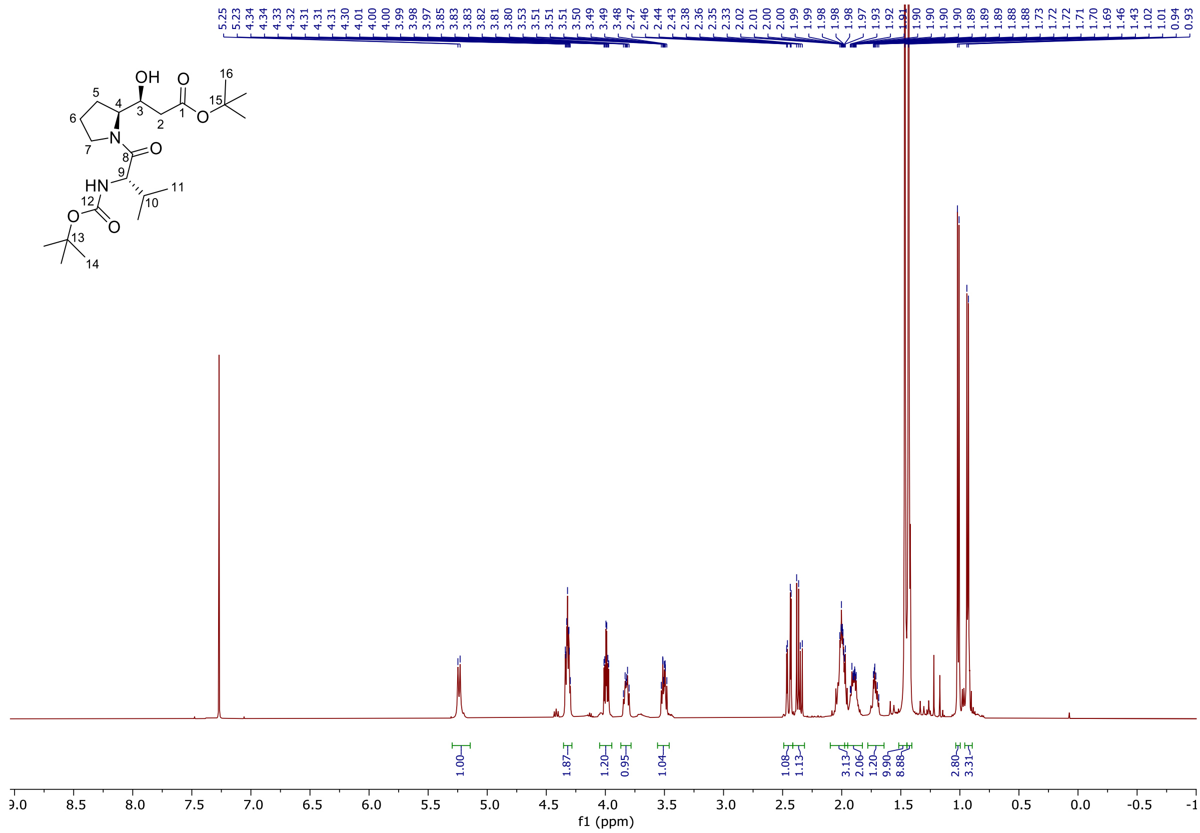


**$^{13}$C-NMR** (100 MHz, $CDCl_3$):

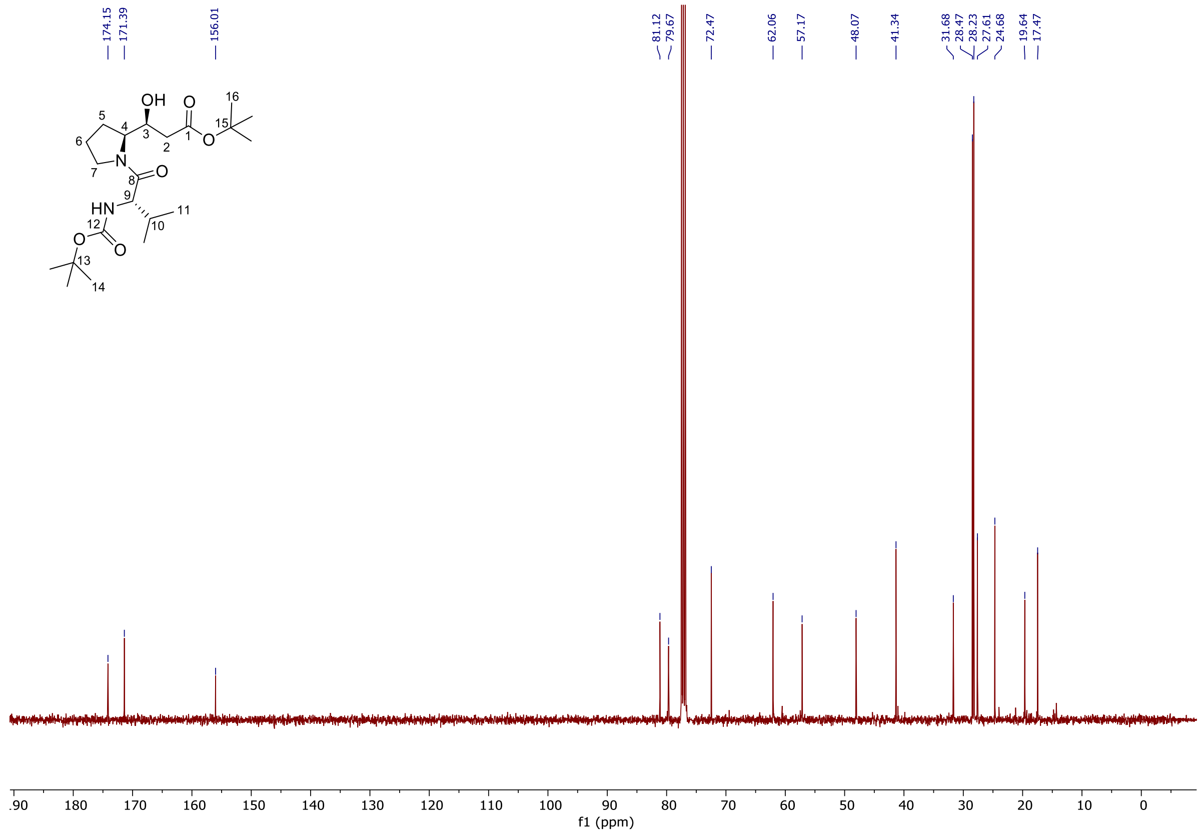

**(*S*)-3-(*tert*-Butoxy)-1-((*S*)-1-((*tert*-butoxycarbonyl)-L-valyl)pyrrolidin-2-yl)-3-oxopropyl(((9*H*-fluoren-9-yl)methoxy)carbonyl)-L-methioninate SI-10**

**$^1$H-NMR** (500 MHz, $CDCl_3$):

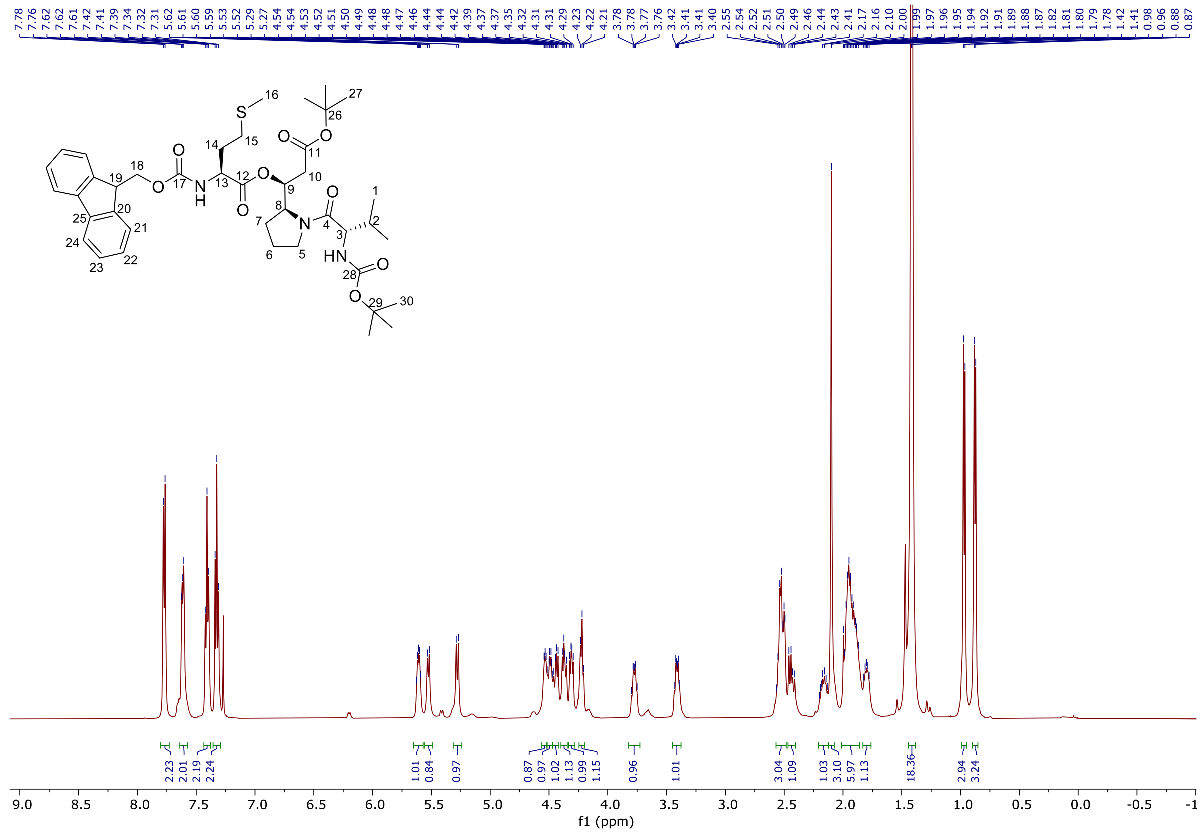


**$^{13}$C-NMR** (125 MHz, $CDCl_3$):

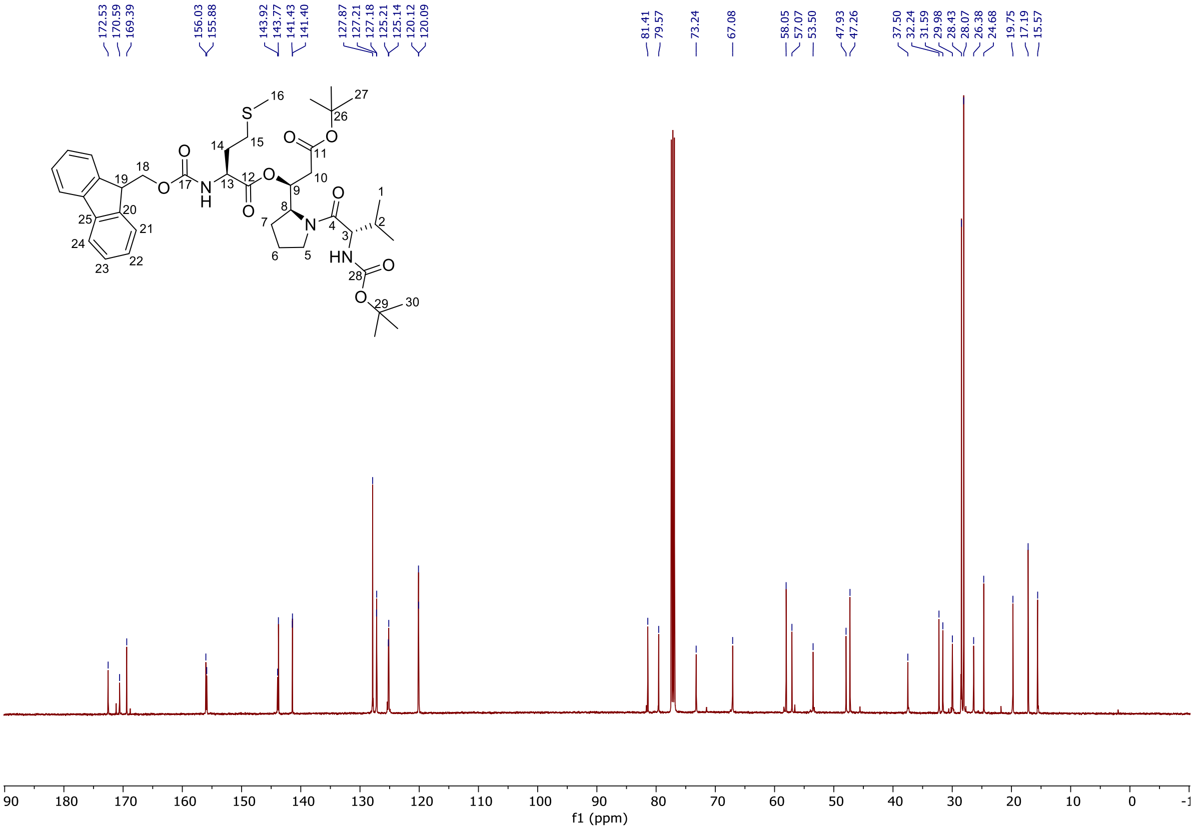

**Methyl (*S*)-3-((*S*)-1-((*tert*-butoxycarbonyl)-L-isoleucyl)pyrrolidin-2-yl)-3-hydroxypropanoate SI-12**

**$^{1}$H-NMR** (500 MHz, $CDCl_3$): (contains 4% EtOAc as impurity)

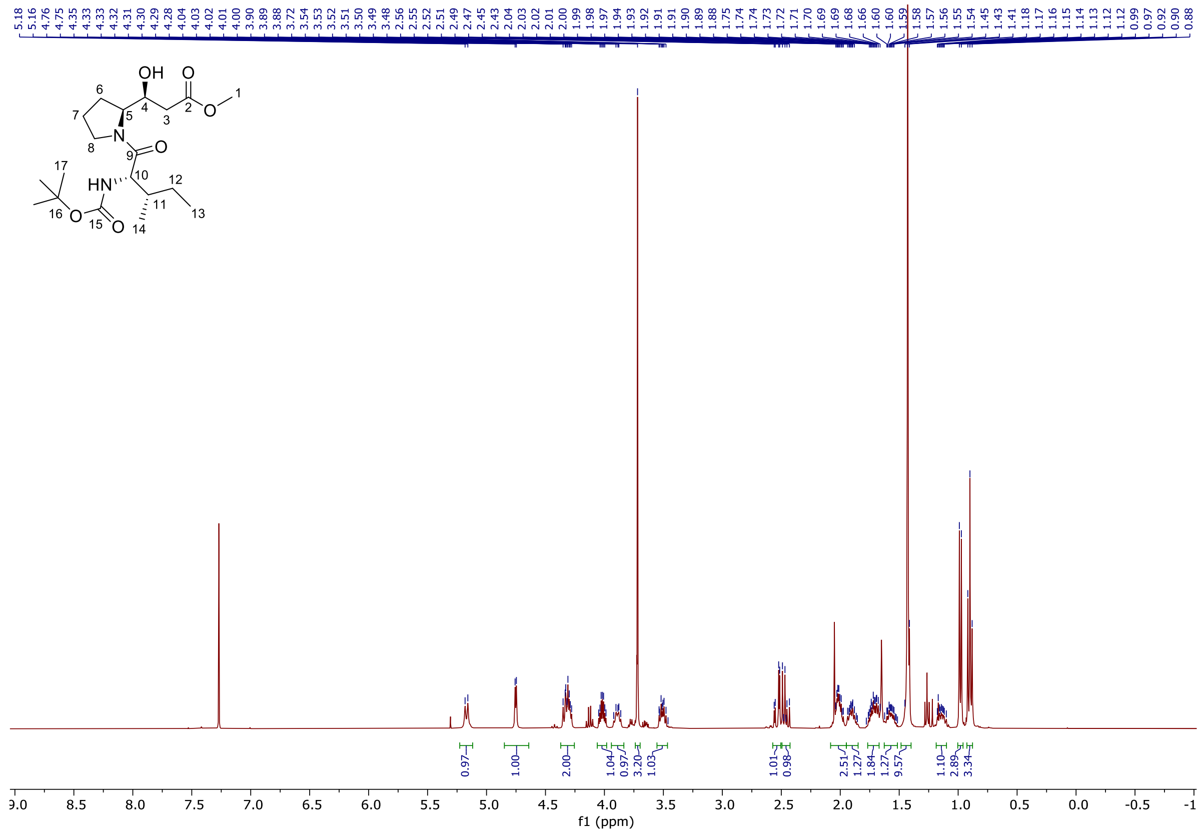


**$^{13}$C-NMR** (100 MHz, $CDCl_3$):

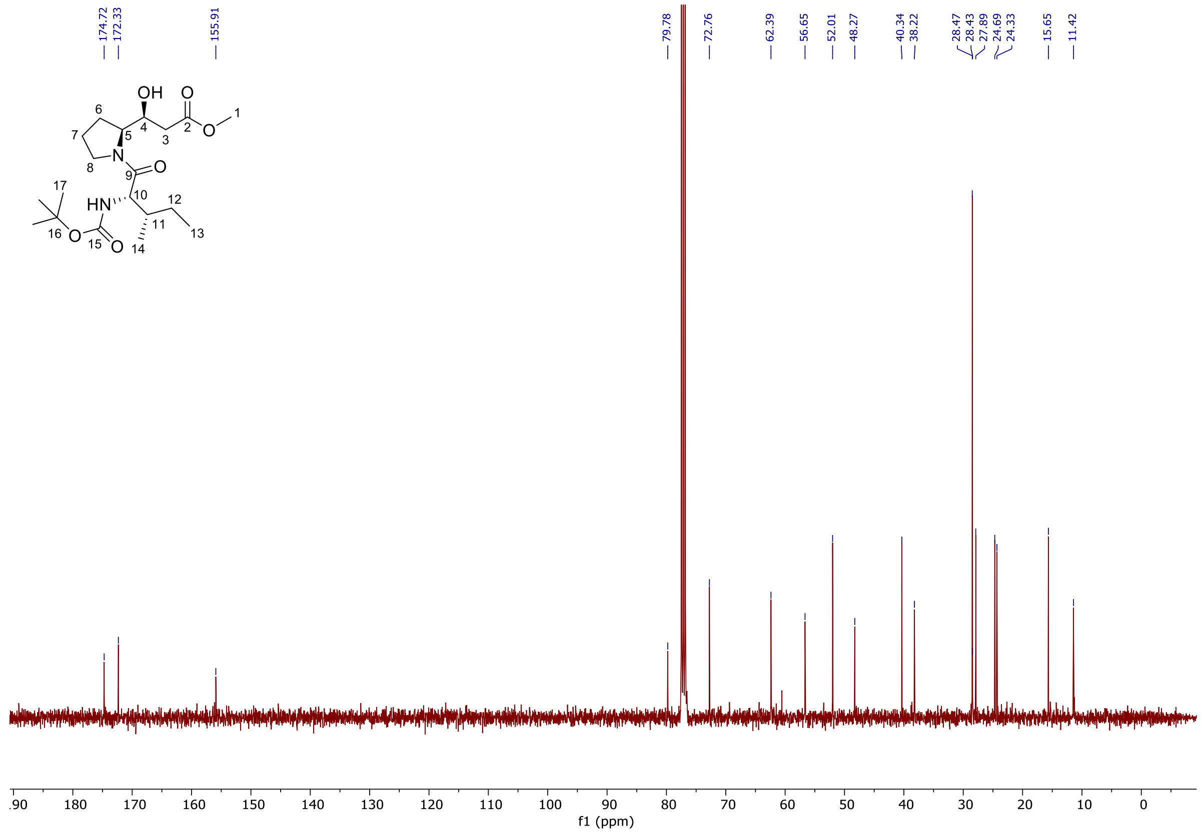

**tert-Butyl (S)-3-((S)-1-((tert-butoxycarbonyl)-L-isoleucyl)pyrrolidin-2-yl)-3-hydroxypropanoate SI-13**

**[1]H-NMR** (400 MHz, $CDCl_3$): (contains 6% DCM as impurity)

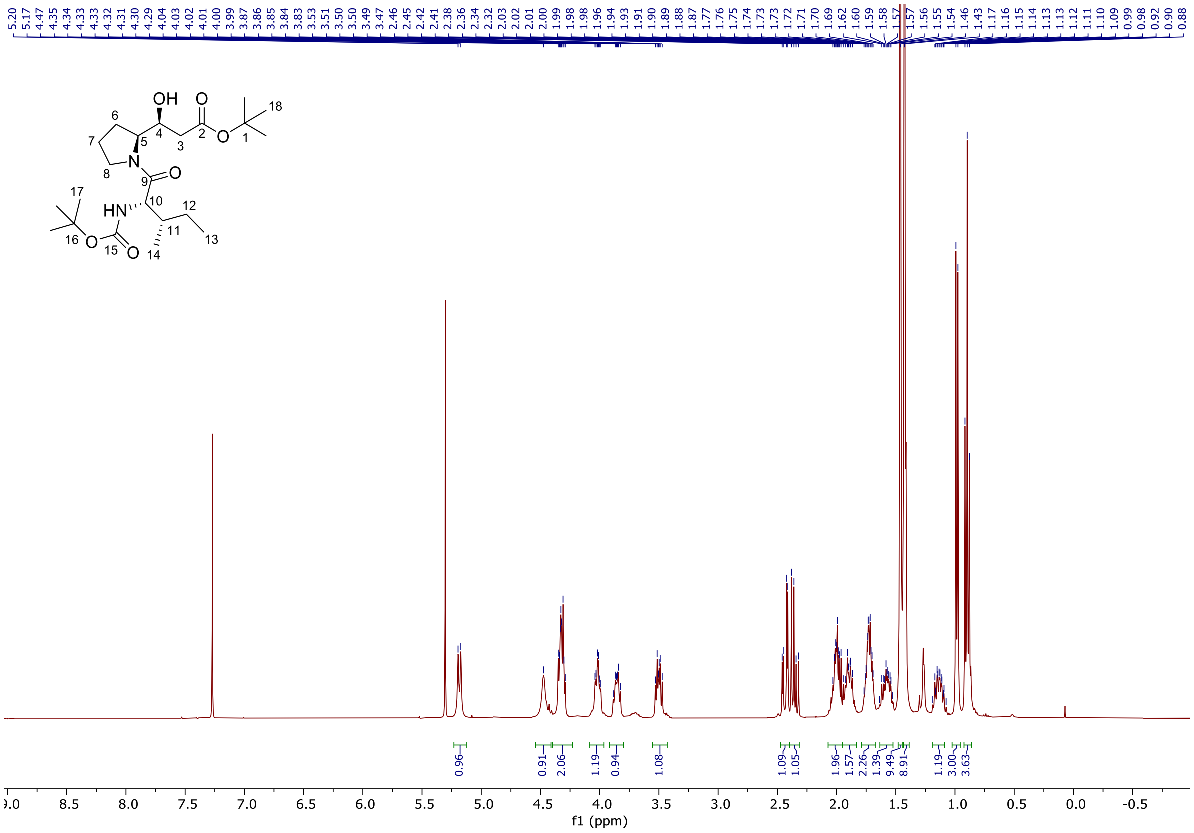


**[13]C-NMR** (100 MHz, $CDCl_3$):

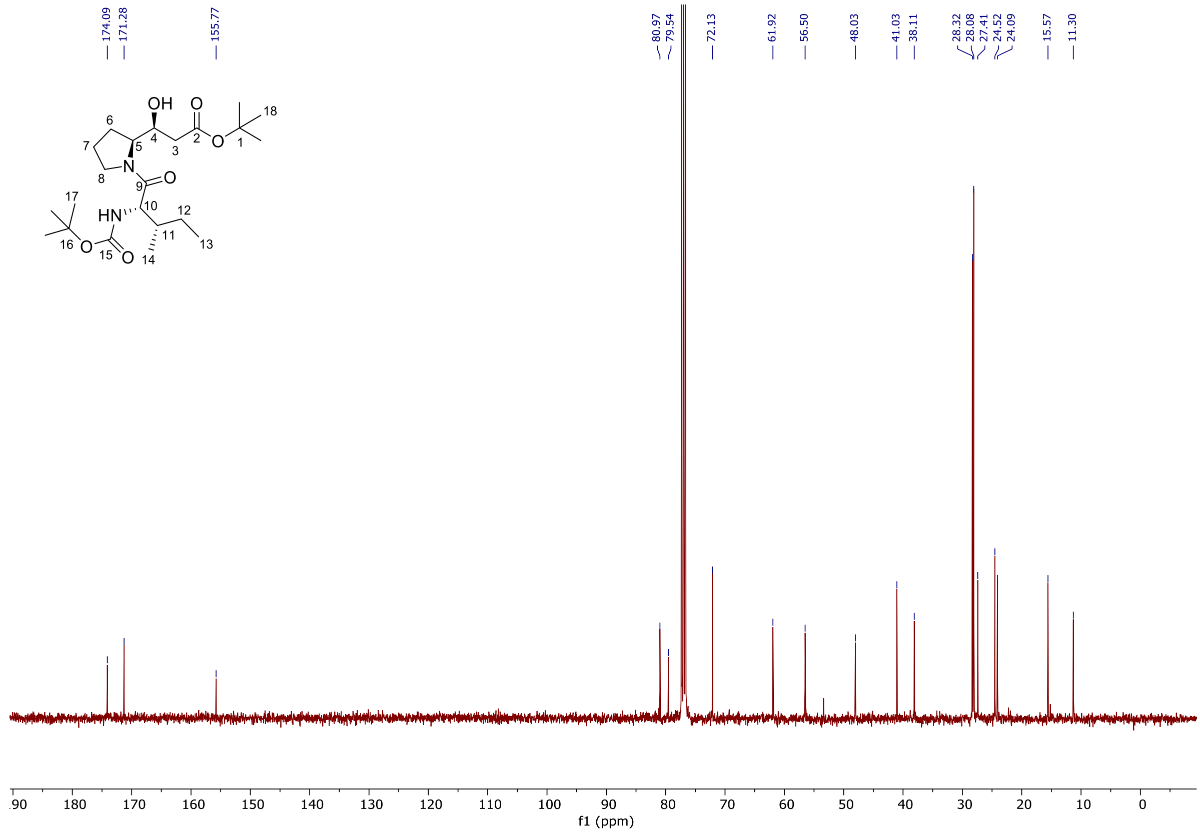

**(*S*)-3-(*tert*-Butoxy)-1-((*S*)-1-((*tert*-butoxycarbonyl)-L-isoleucyl)pyrrolidin-2-yl)-3-oxopropyl-(((9H-fluoren-9-yl)methoxy)carbonyl)-L-methioninate 8**

**$^{1}$H-NMR** (500 MHz, $CDCl_3$):

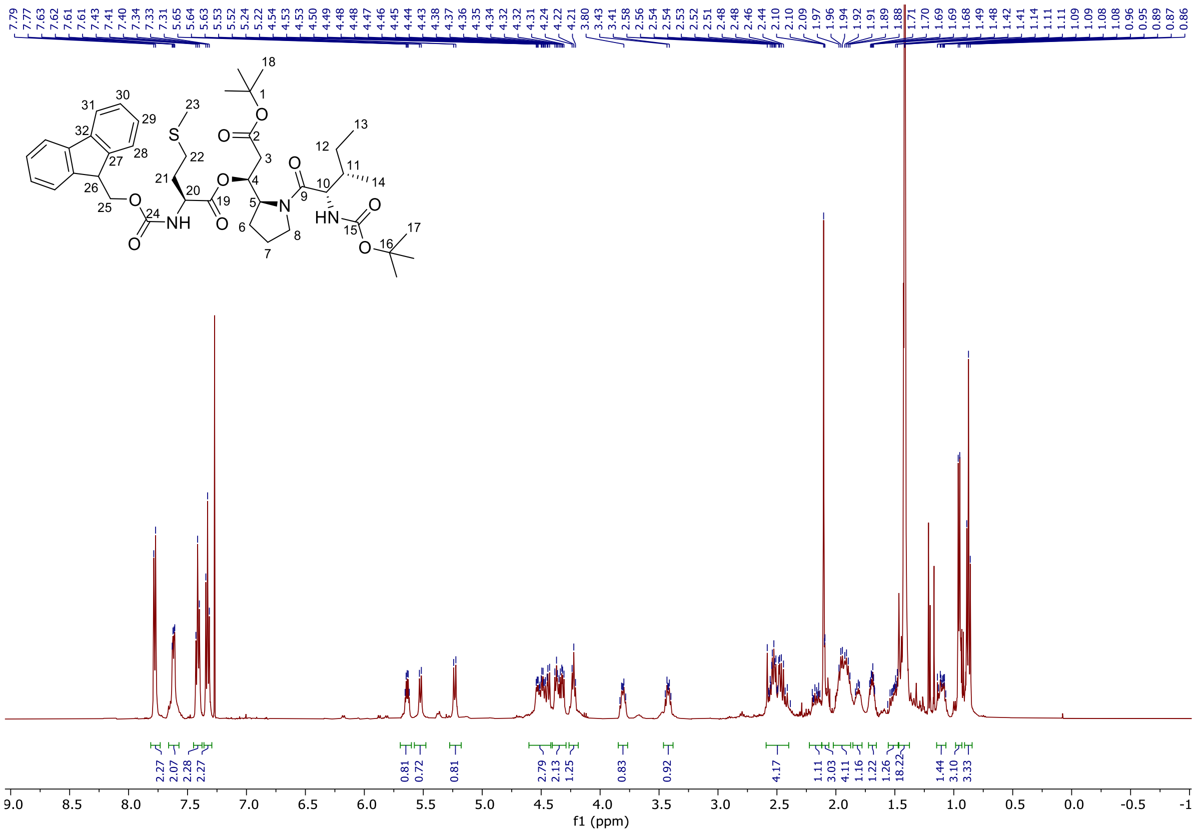


**$^{13}$C-NMR** (125 MHz, $CDCl_3$):

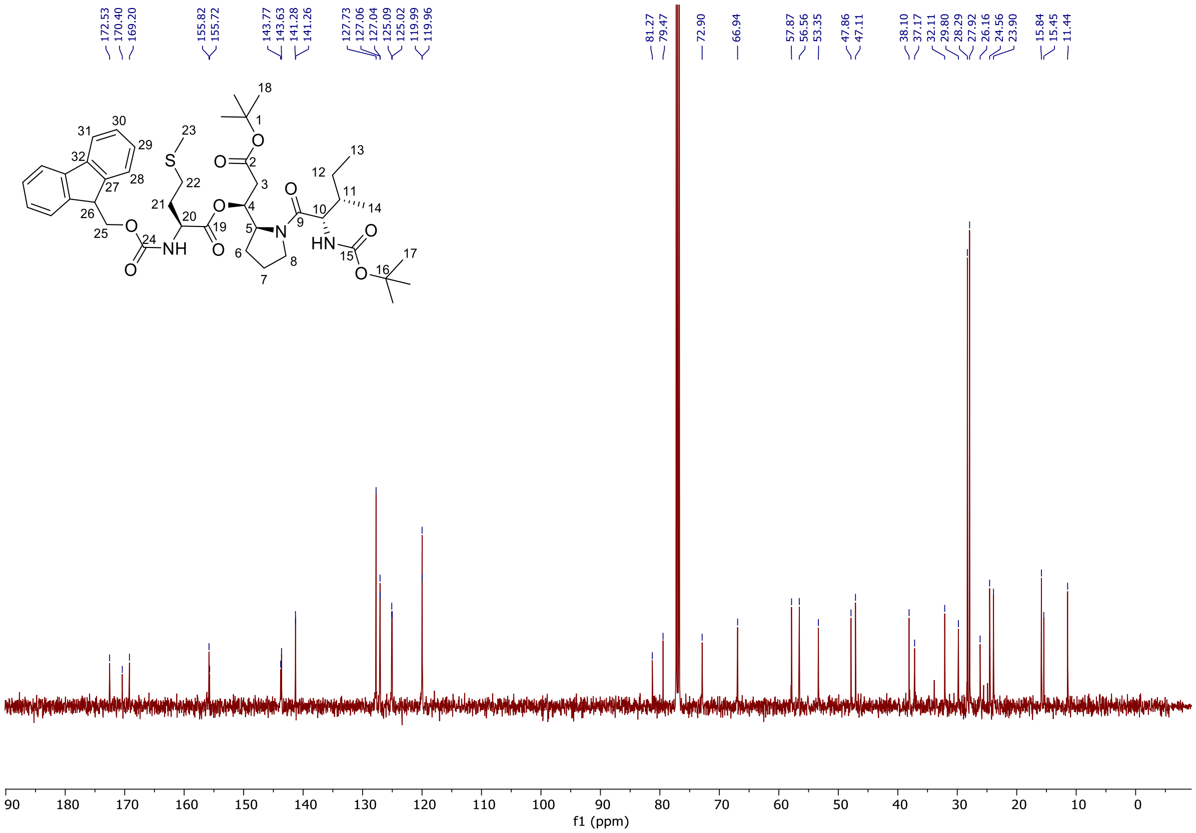

**Ethyl 2-((*S*)-1-((*S*)-2-((*tert*-butoxycarbonyl)amino)-5-(((2,2,2-trichloroethoxy)carbonyl)amino) pentanoyl)pyrrolidin-2-yl)thiazole-4-carboxylate SI-15**

**$^{1}$H-NMR** (500 MHz, $CDCl_3$): (contains 1% DCM as impurity)

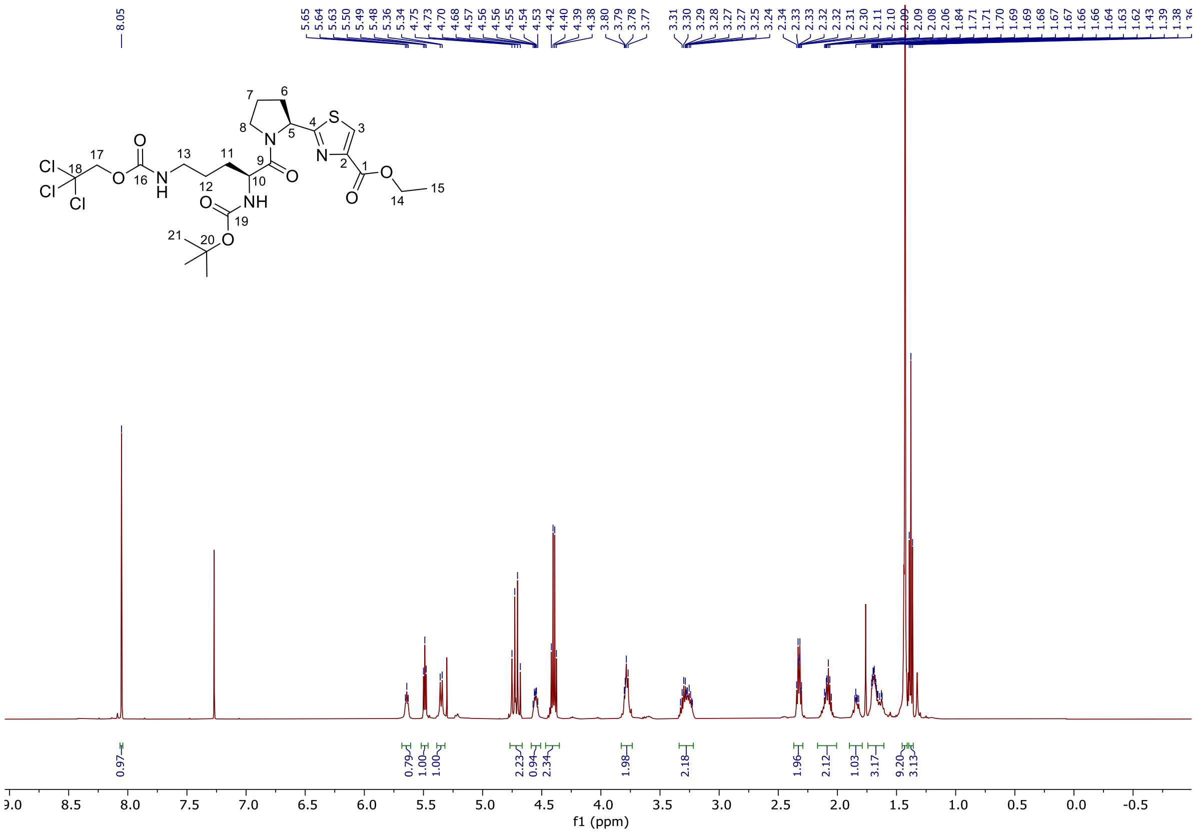


**$^{13}$C-NMR** (125 MHz, $CDCl_3$):

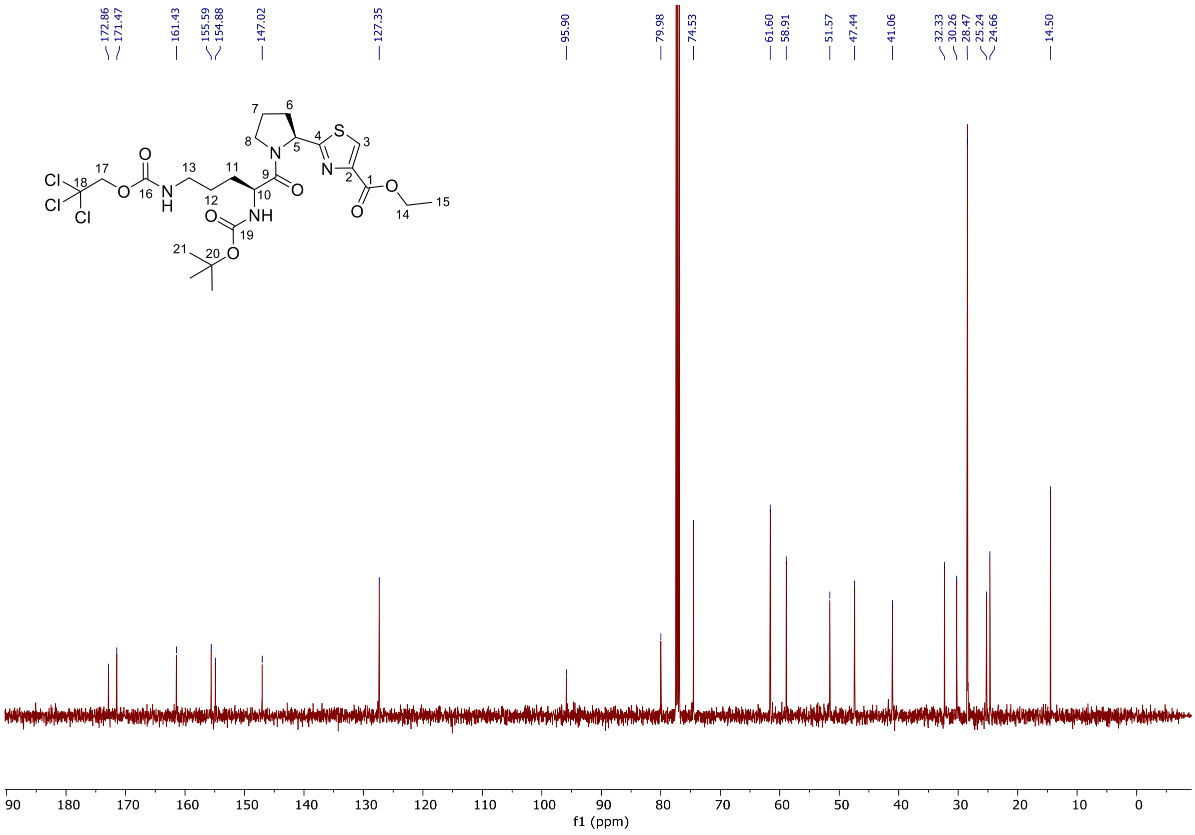

**Ethyl 2-((*S*)-1-((*S*)-2-acetamido-5-(((2,2,2-trichloroethoxy)carbonyl)amino)pentanoyl) pyrrolidin-2-yl)thiazole-4-carboxylate 10**

**[1]H-NMR** (500 MHz, $CDCl_3$):

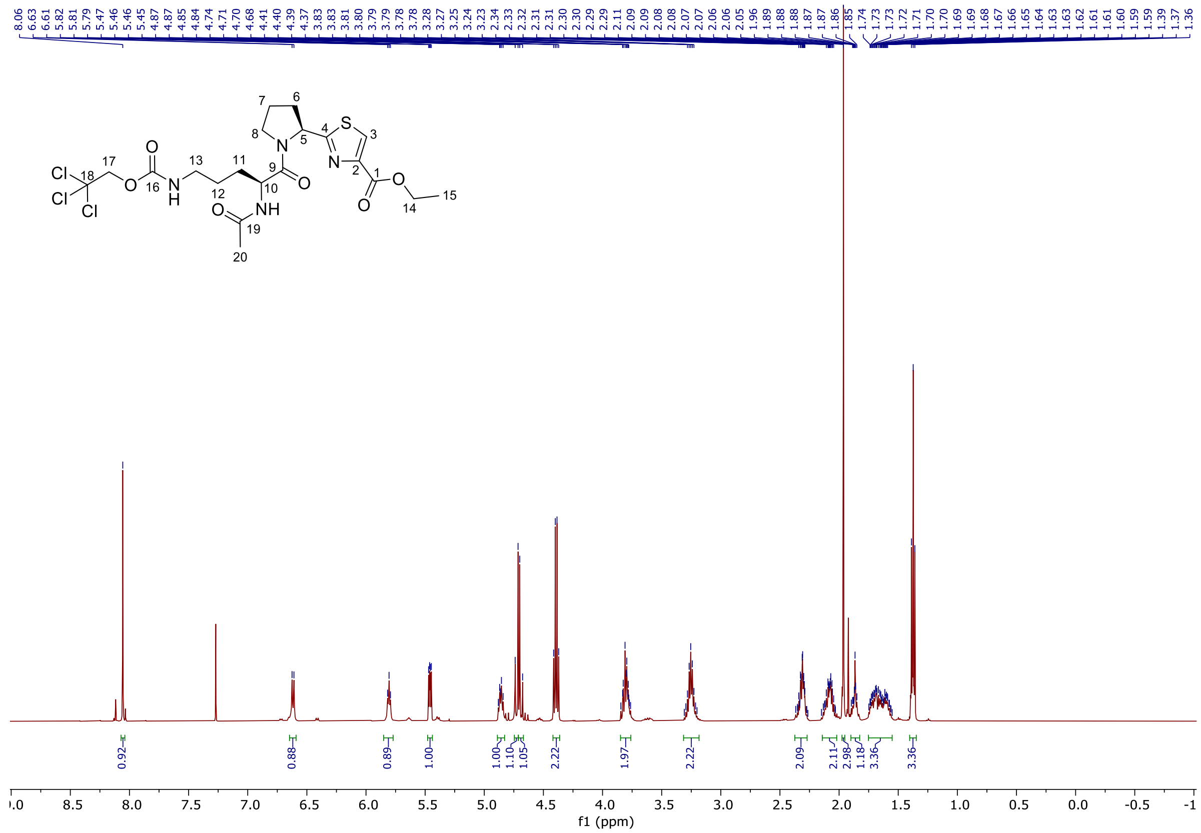


**[13]C-NMR** (125 MHz, $CDCl_3$):

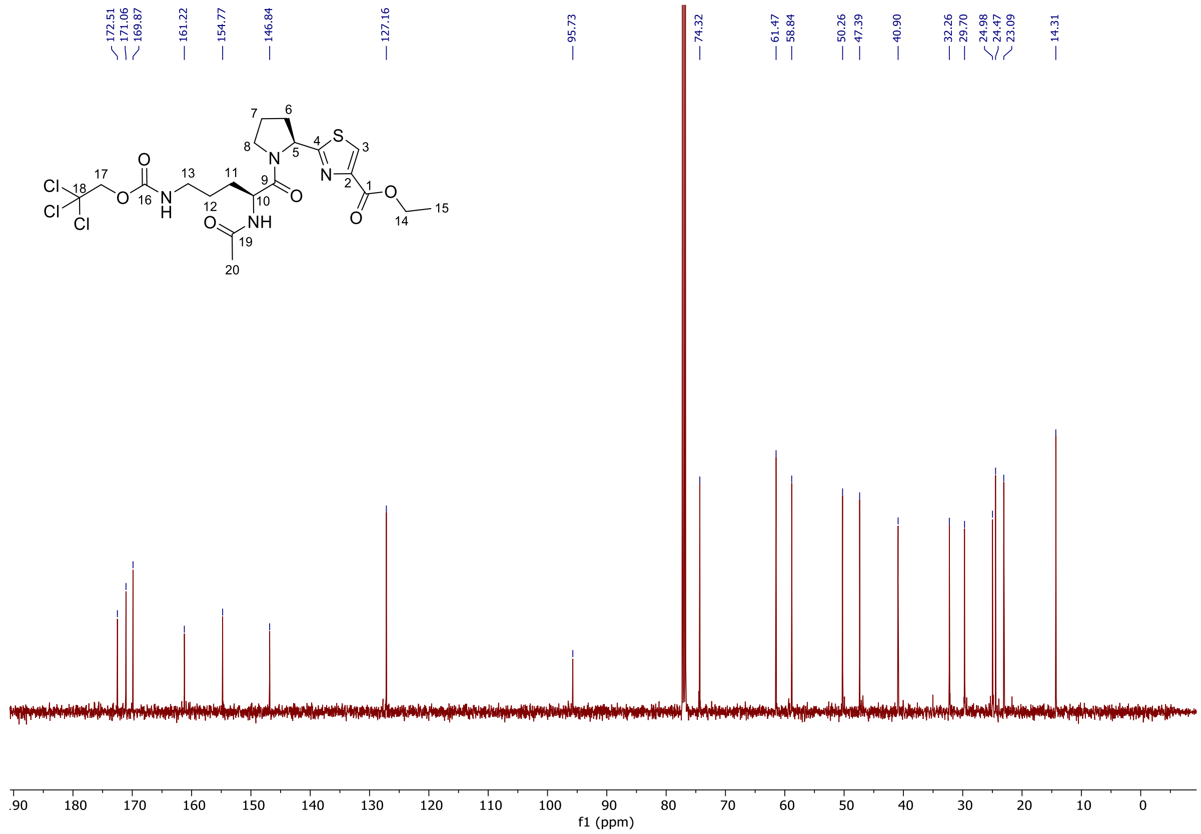

**Ethyl 2-((*S*)-1-($N^2$-acetyl-$N^{\omega}$,$N^{\omega'}$-bis(*tert*-butoxycarbonyl)-L-arginyl)pyrrolidin-2-yl)thiazole-4-carboxylate 11**

**$^1$H-NMR** (500 MHz, $CDCl_3$):

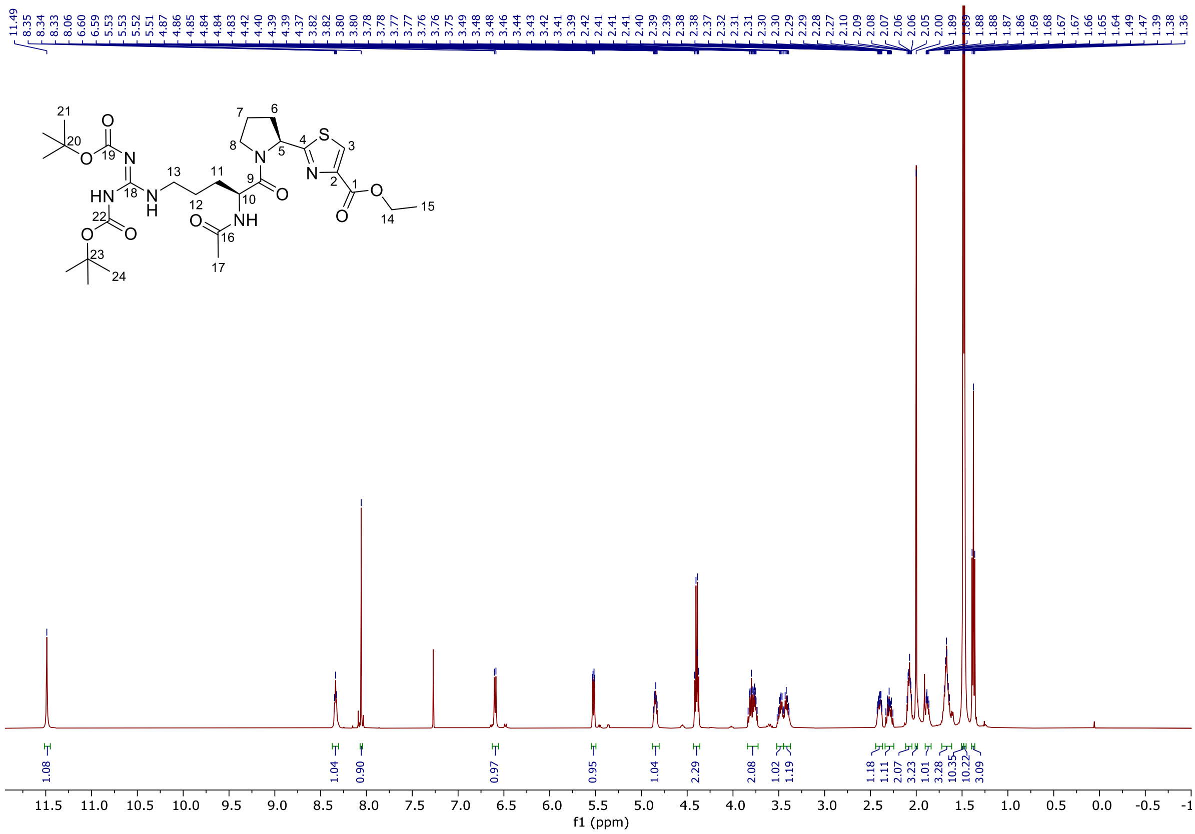


**$^{13}$C-NMR** (125 MHz, $CDCl_3$):

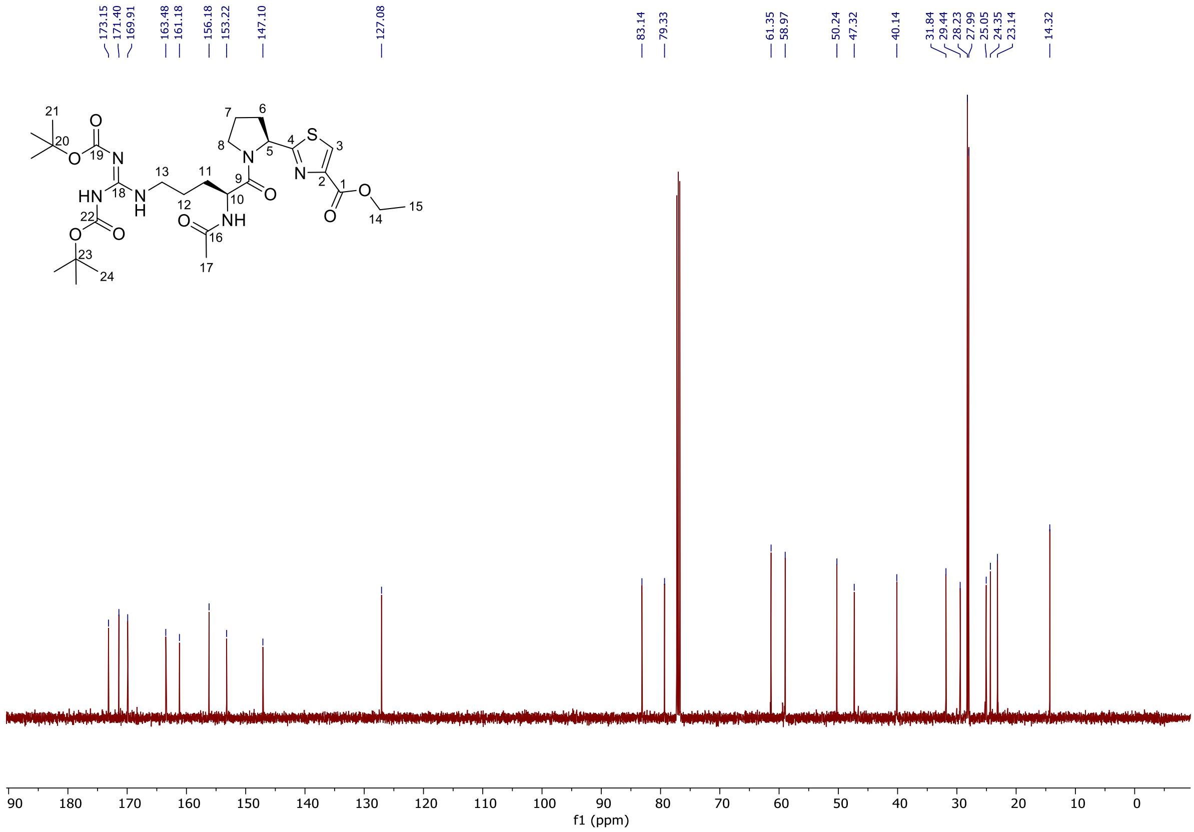

**(*S*)-3-(*tert*-Butoxy)-1-((*S*)-1-((*tert*-butoxycarbonyl)-L-valyl)pyrrolidin-2-yl)-3-oxopropyl (2-((*S*)-1-($N^2$-acetyl-$N^{\omega}$,$N^{\omega'}$-bis(*tert*-butoxycarbonyl)-L-arginyl)pyrrolidin-2-yl)thiazole-4-carbonyl)-L-methioninate SI-18**

**$^1$H-NMR** (500 MHz, $CDCl_3$):

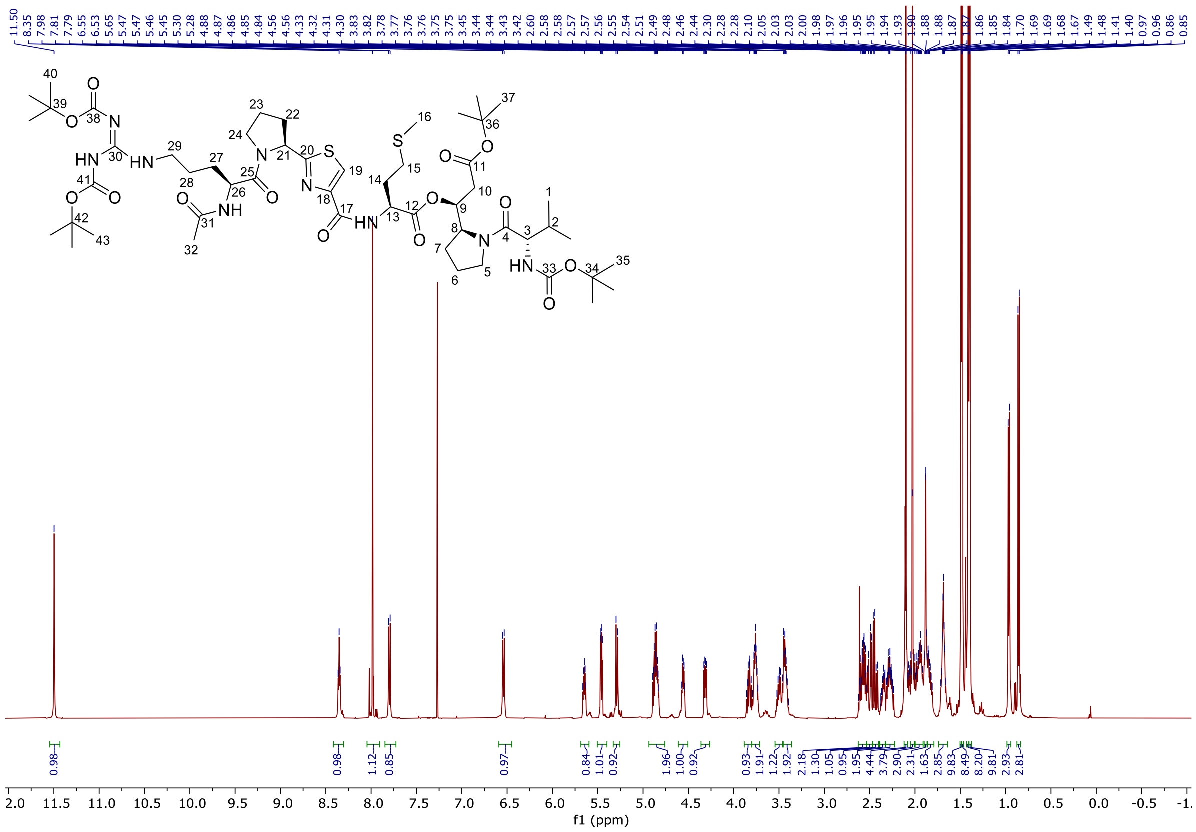


**$^{13}$C-NMR** (125 MHz, $CDCl_3$):

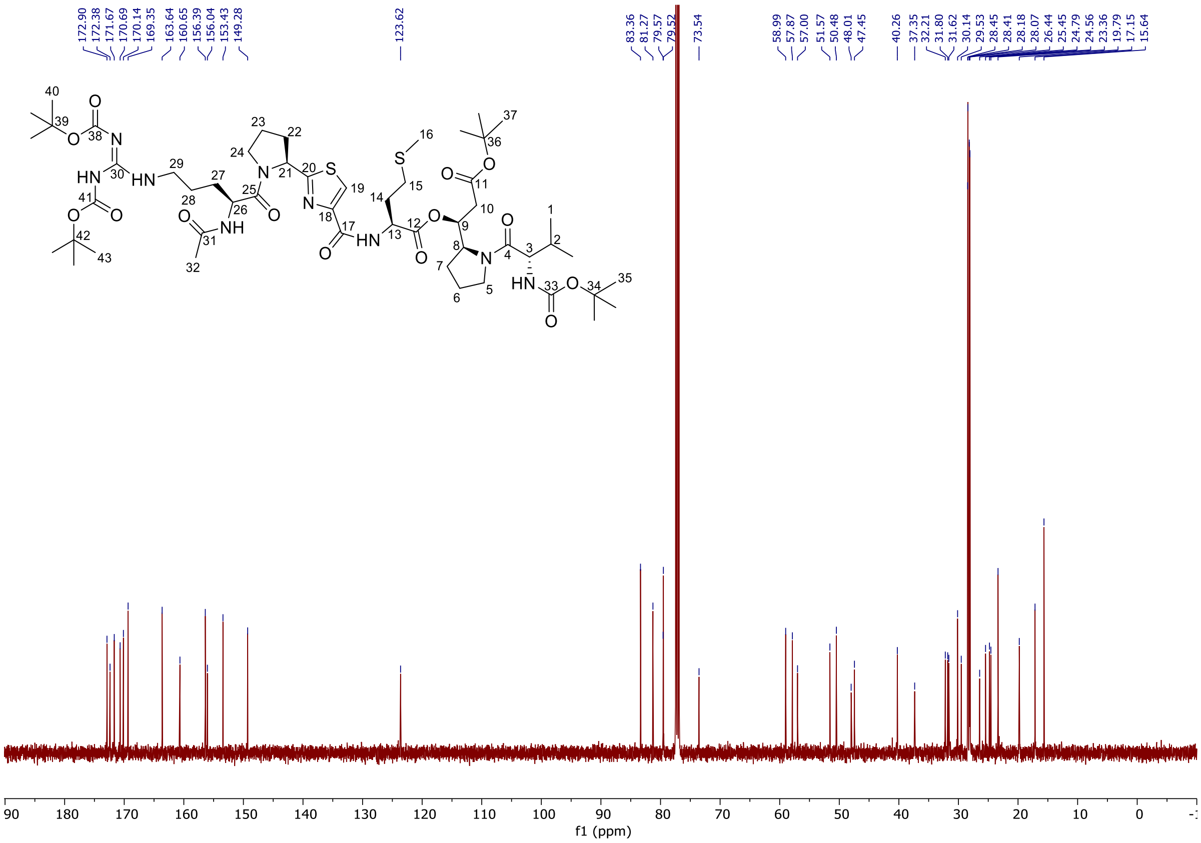

**(*S*)-3-(*tert*-Butoxy)-1-((*S*)-1-((*tert*-butoxycarbonyl)-L-isoleucyl)pyrrolidin-2-yl)-3-oxopropyl (2-((*S*)-1-($N^2$-acetyl-$N^{\omega}$,$N^{\omega'}$-bis(*tert*-butoxycarbonyl)-L-arginyl)pyrrolidin-2-yl)thiazole-4-carbonyl)-L-methioninate 12**

**$^1$H-NMR** (500 MHz, $CDCl_3$):

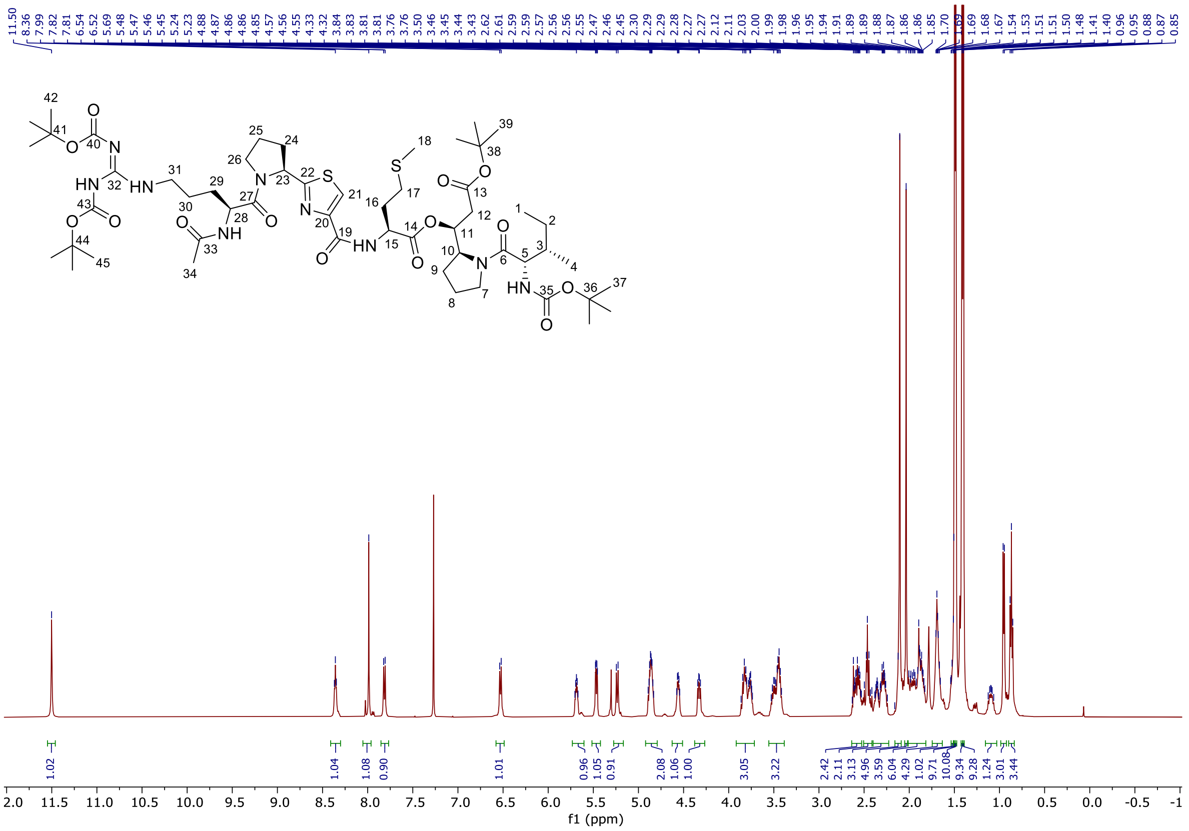


**$^{13}$C-NMR** (125 MHz, $CDCl_3$):

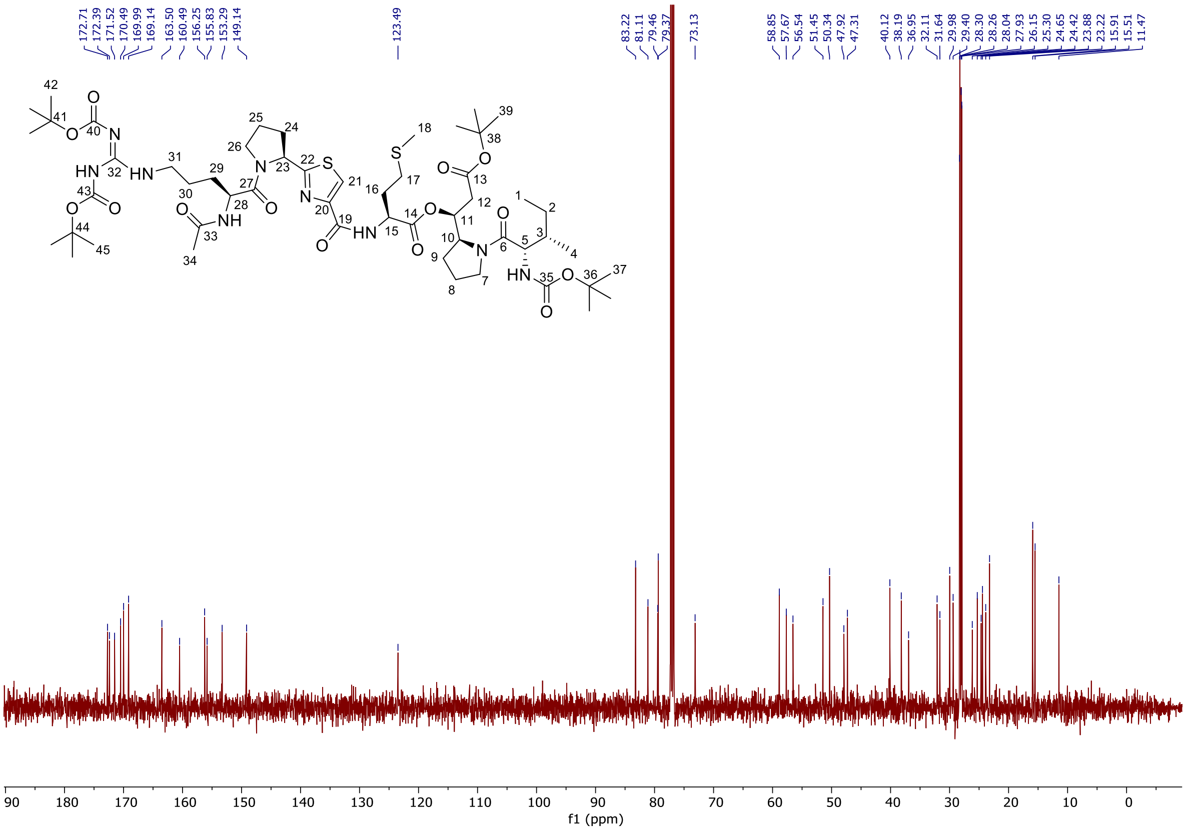

**Pseudotetraivprolid D 5a**

**$^1$H-NMR** (500 MHz, $D_2O$):

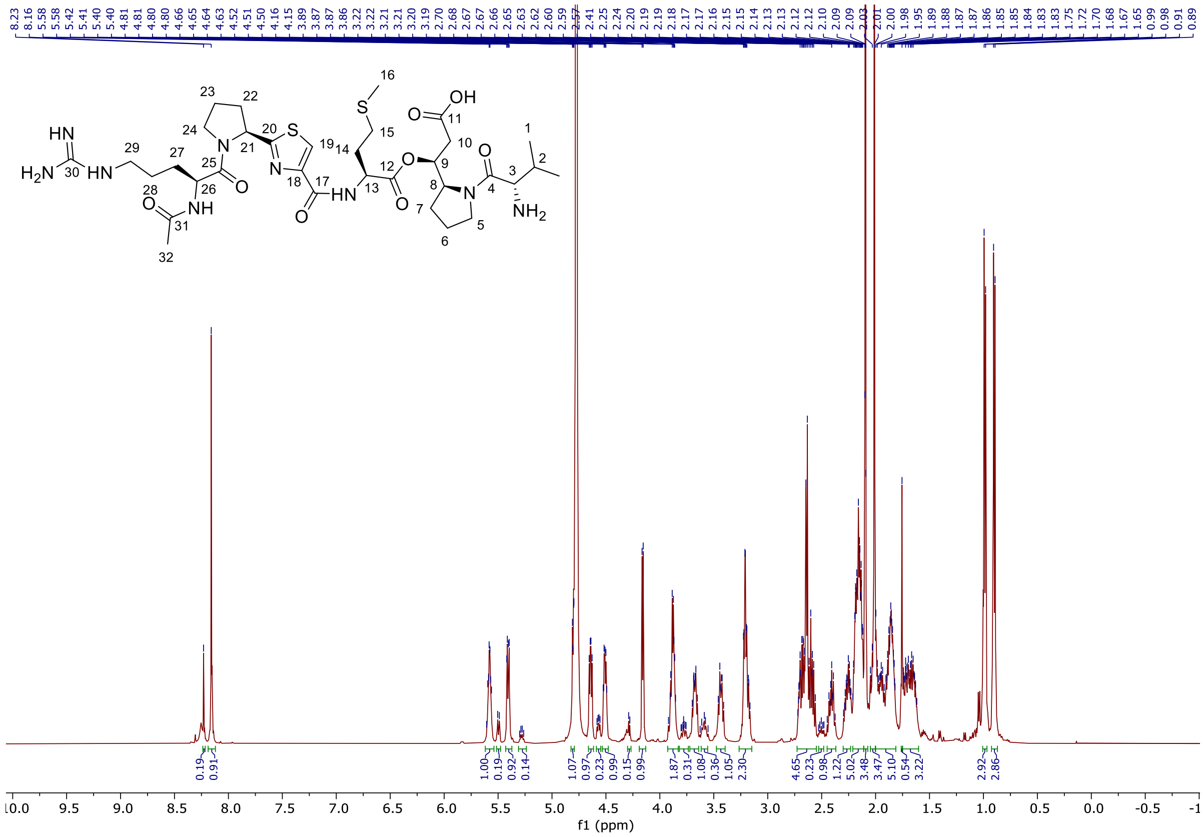


**$^{13}$C-NMR** (125 MHz, $D_2O$):

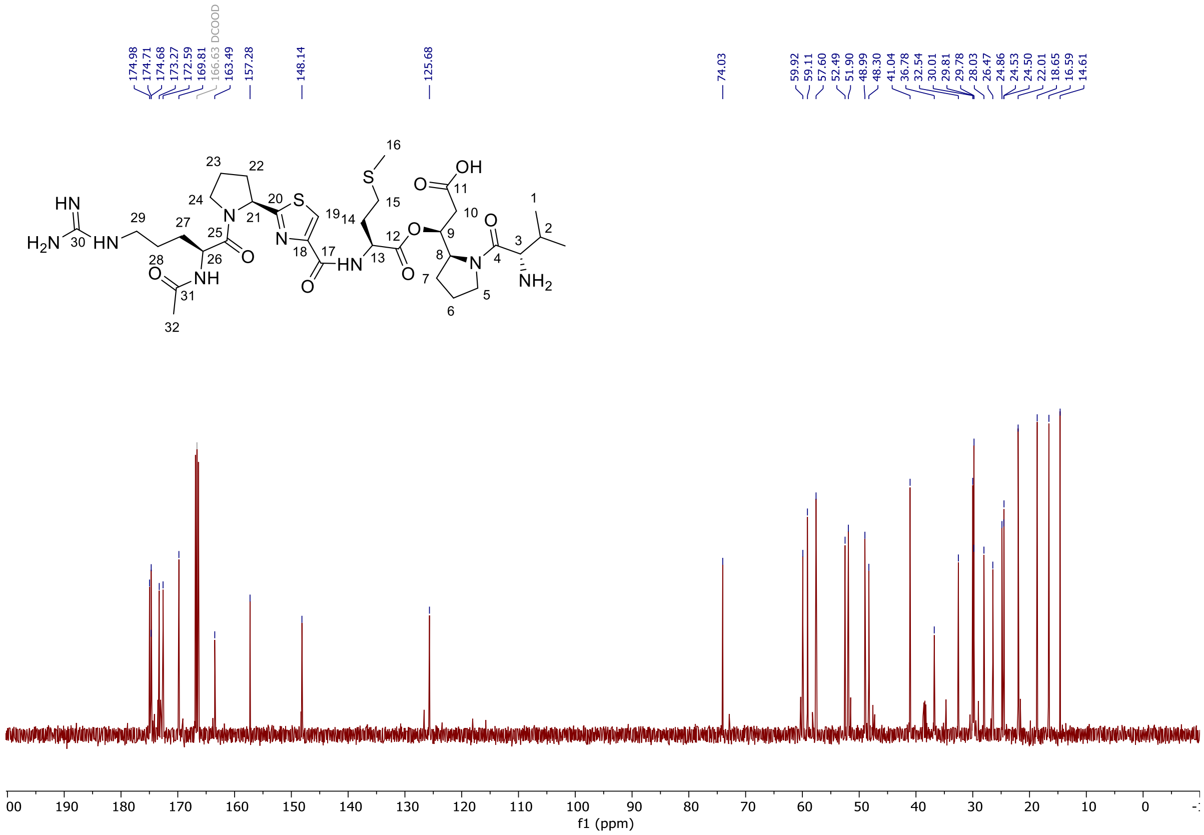

**Pseudotetraivprolid B 4a**

**$^{1}$H-NMR** (500 MHz, $D_2O$):

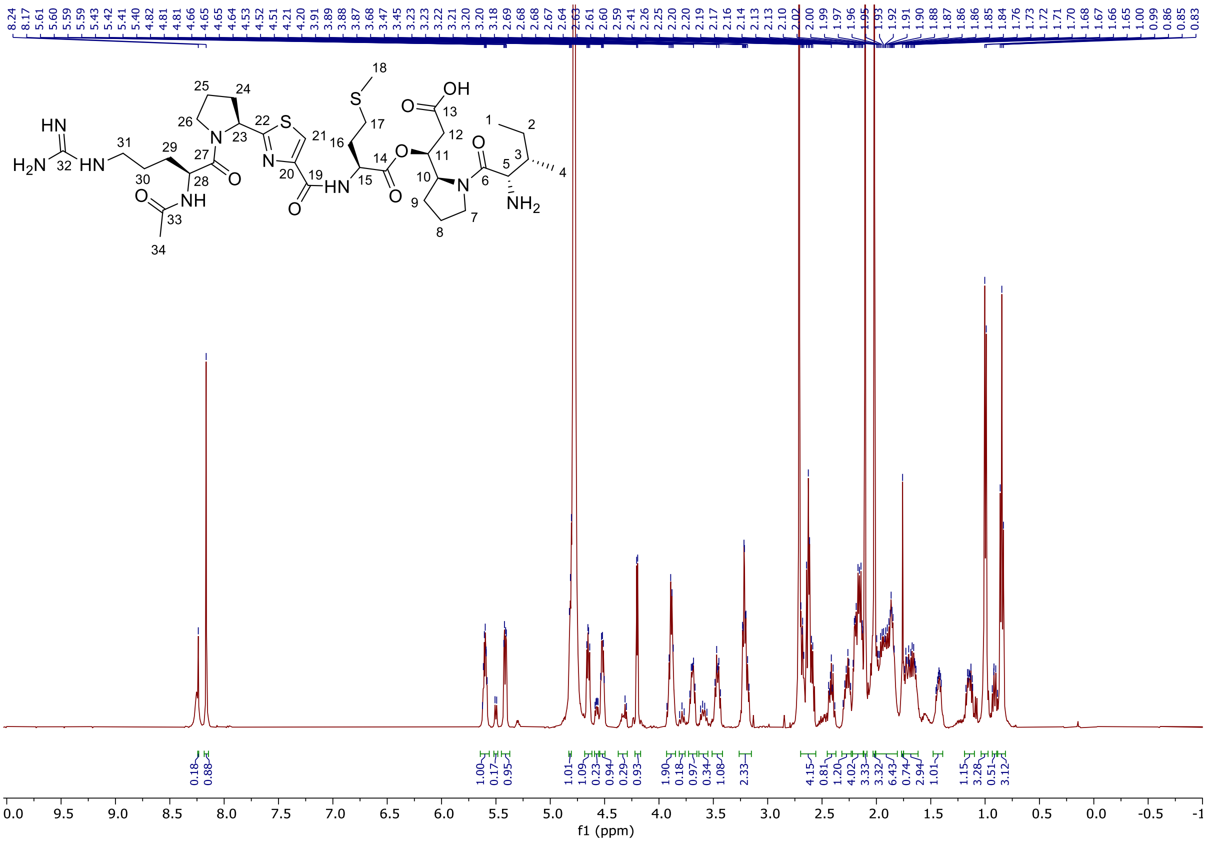

**$^{13}$C-NMR** (125 MHz, $D_2O$):

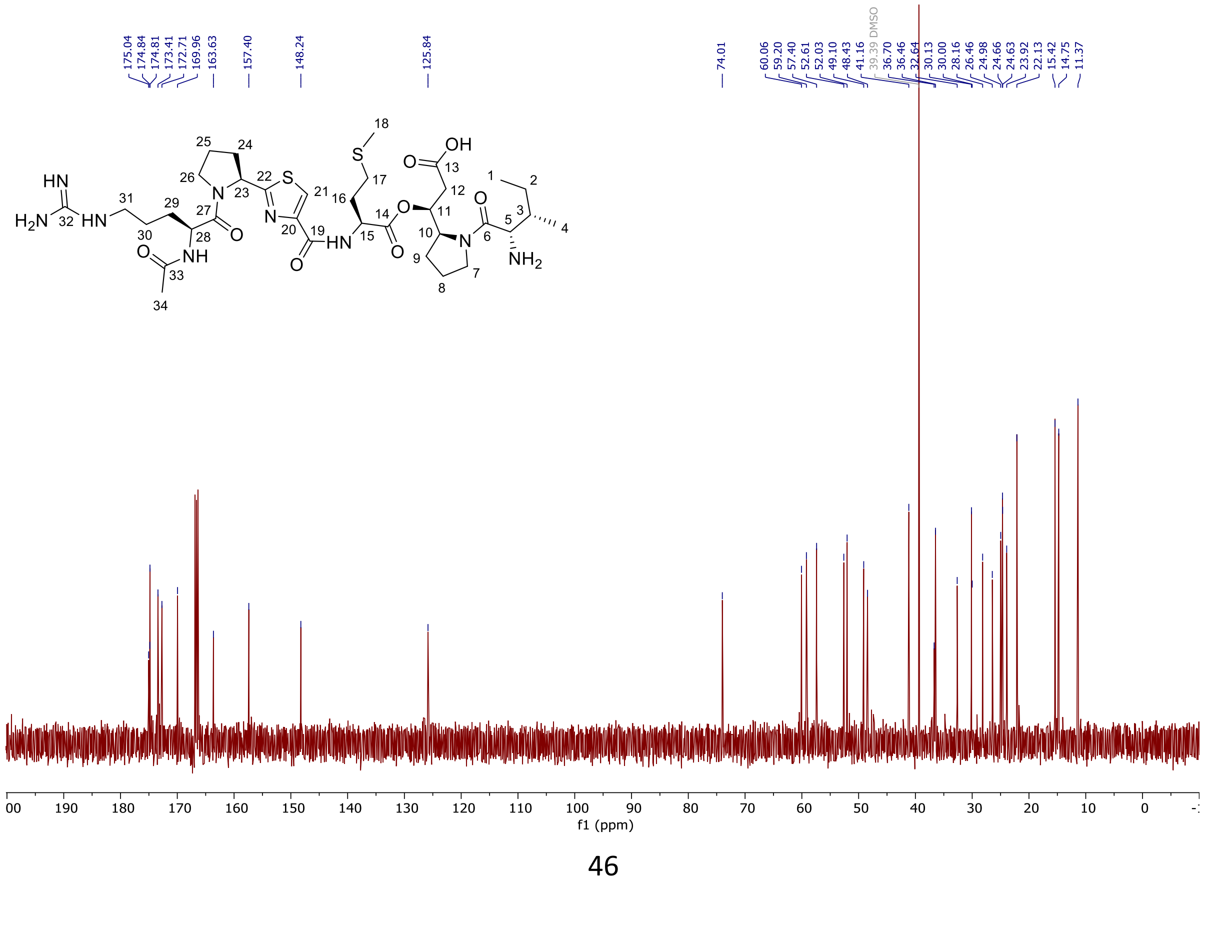

**(*S*)-2-((*tert*-Butoxycarbonyl)amino)-5-(((2,2,2-trichloroethoxy)carbonyl)amino)pentanoic acid SI-19**

**$^{1}$H-NMR** (500 MHz, DMSO-d$_6$):

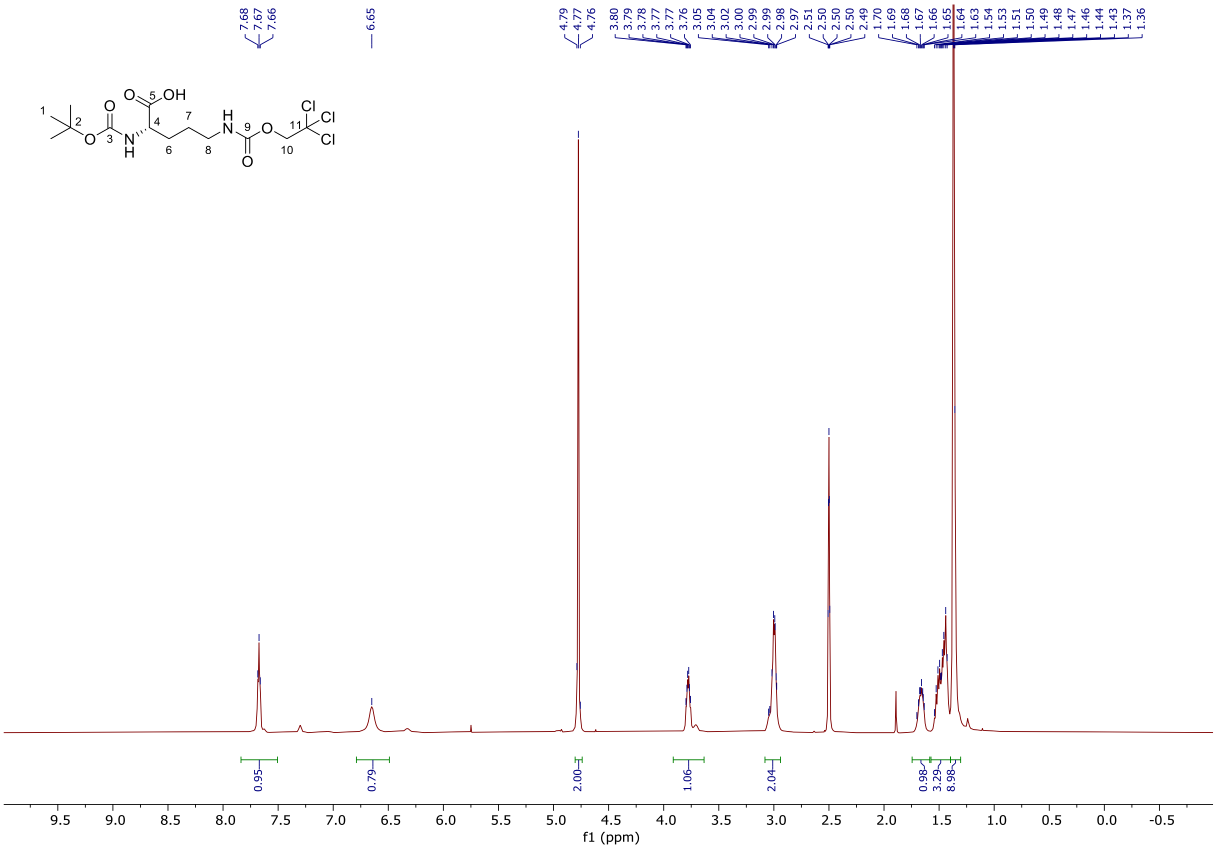


**$^{13}$C-NMR** (125 MHz, DMSO-d$_6$):

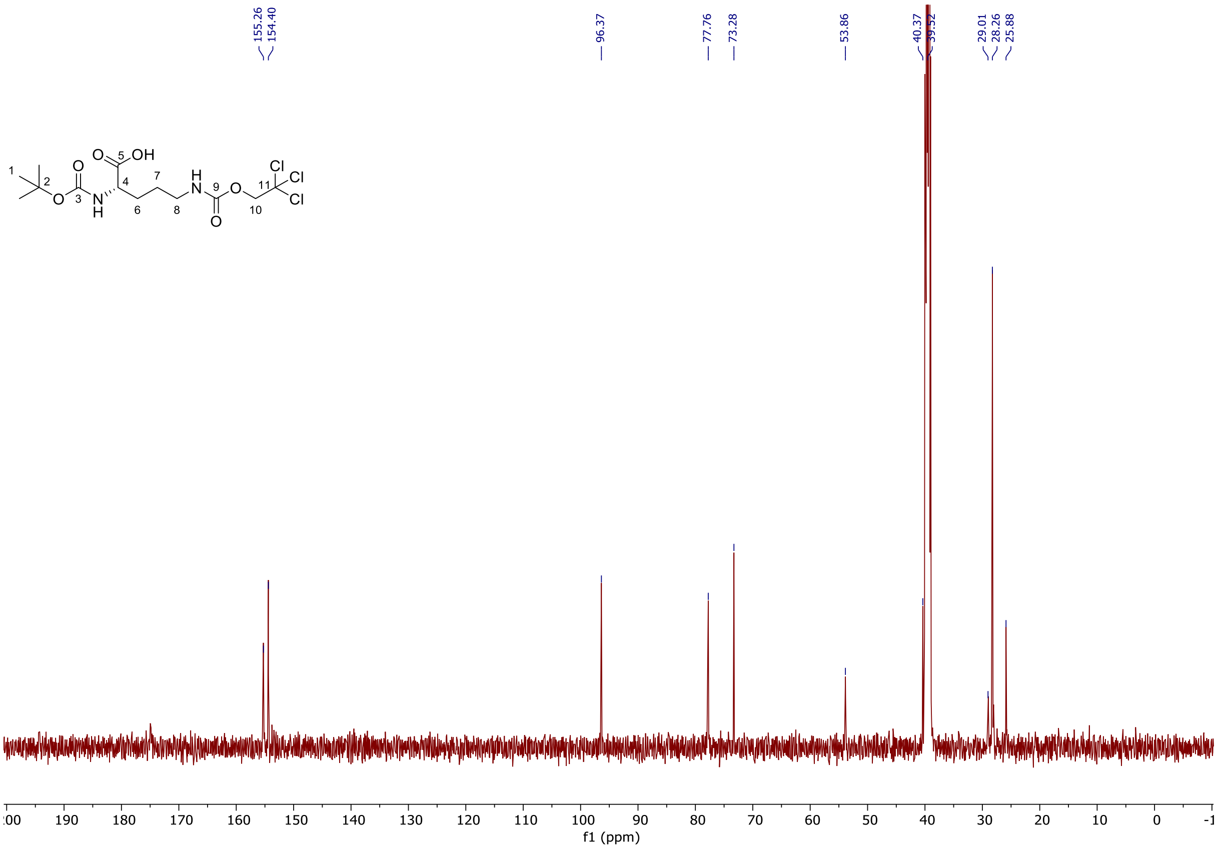
