## Supplementary Information SI-4 for "Identification of pseudotetraivprolide from *Pseudomonas entomophila* give novel insights into the biosynthesis of detoxin/rimosamide-like anti-antibiotics"

### Supplementary Information SI-4: Prediction of structure and function of PipD, PipF and PipG

#### 1. Sequence Information of Proteins:

##### PipD:

**hypothetical protein – annotated by Geneious #BLAST results is Uncharacterized protein (Max similarity: 49.7%)**
MPAWLGPLLRVLETGPLPRQLDCQAYWRLLHRWQAEVVLPLLARALPEHDHAVTTLRRLHQRASLGLRGRVGEWRAALGPVL
LAVFRRAYAFDSAYAQAYDSALDYGLAASNQAMIAEQFGDAEAFARHYAQLSTDANAQAFATANAAASSMLVARAYASEDEQA
CASVVGALARACNWACANAAEQRAAVREQLAAGLQHVASQHPSGRNVWINRH

**hypothetical protein - annotated by antiSMASH #BLAST results is Uncharacterized protein (Max similarity: 49.7%)**
MAGEYAQVPAWLGPLLRVLETGPLPRQLDCQAYWRLLHRWQAEVVLPLLARALPEHDHAVTTLRRLHQRASLGLRGRVGEWR
AALGPVLLAVFRRAYAFDSAYAQAYDSALDYGLAASNQAMIAEQFGDAEAFARHYAQLSTDANAQAFATANAAASSMLVARAYA
SEDEQACASVVGALARACNWACANAAEQRAAVREQLAAGLQHVASQHPSGRNVWINRH

##### PipF:

**hypothetical protein - annotated by Geneious #BLAST results is SGNH hydrolase-type esterase domain-containing protein (Max similarity: 57.8%)**
MDFFTLAPDDVVFRDTELERLYRQALAQLDAPSLGRIPGPLVSSIGLSLHYLATRQNWLCYQGLEGEFDPGFLEGPLFAAIIDTLS
RPALAFYEQALGLGLKVYAVLPPQRVPPMADARVFMAAQTHLIERLTALGVELVDVREAANDDQGRQRAEFCEVDDPLHGNLA
FGELVLGQLLRQVPSKGPALALRSP

**hypothetical protein - annotated by antiSMASH #BLAST results is SGNH hydrolase-type esterase domain-containing protein (Max similarity: 60.9%)**
MKTPFSITTPSFLLLGDSHLGVVGQAARARQLSFSGGPLGAGRDFGVDFFTLAPDDVVFRDTELERLYRQALAQLDAPSLGRIP
GPLVSSIGLSLHYLATRQNWLCYQGLEGEFDPGFLEGPLFAAIIDTLSRPALAFYEQALGLGLKVYAVLPPQRVPPMADARVFMA
AQTHLIERLTALGVELVDVREAANDDQGRQRAEFCEVDDPLHGNLAFGELVLGQLLRQVPSKGPALALRSP

##### PipG:

**GSCFA domain containing protein #BLAST results is GSCFA family protein (Max similarity: 65.9%)**
MNPYQYLPPRAFWRTAIAARPTEQIAELWSPAFTIDAKDAIVTAGSCFAQHIGRALVARGMNWLDAEPAPAEMLEDERKARQYG
VFSFRAGNLYTAAMLRQWLTWALGTQPQSHETWQHEGRFFDPFRPAVETAGFDSEQALFDSREQTLAAIRLAVHRAKVFVFTL
GLTEAWANRESGVVYPVCPGTVRGEFDPRVHEFRNFGFNDTCQAMTEAIALMRTVNPELRLLLTVSPVPLTASATGEHVLSATT
YSKSVLRAVAGQLCQDLPQVDYFPSYEIITGTPFKGAFYQPNRREVTPGGVAFVMRQFFAGLDAEAAPAAPATTPSLACEDLVC
EDAILDYYA

#### 2. Annotation of function based on sequence information

The web server version of CLEAN[1] was used to predict the function of enzymes from sequence. CLEAN is a machine learning model for protein function prediction based on sequence information. The tools only considering sequence information to not seem powerful enough to identify the function of PipD, PipF and PipG.

**Table S4.1.** Function prediction results by CLEAN.

| EC Number Prediction Results | | |
|---|---|---|
| **Identifier** | **Predicted EC Number** | **Function** |
| PipD (Annotation by Geneious) | EC: 3.4.24.83 | Anthrax lethal factor endopeptidase |
| PipD (Annotation by antiSMASH) | EC: 3.4.24.83 | Anthrax lethal factor endopeptidase |
| PipF (Annotation by Geneious) | EC: 1.3.7.2 | 15,16-dihydrobiliverdin:ferredoxin oxidoreductase |
| PipF (Annotation by antiSMASH) | EC: 1.13.11.48 | 3-hydroxy-2-methylquinolin-4-one 2,4-dioxygenase |
| PipG | EC: 3.1.6.6 | Choline-sulfatase |

#### 3. Annotation of function based on structure predictions and structure similarity search

##### 3.1. Search for protein with similar structures

As seen above, the tools only considering sequence information to not seem powerful enough to identify the function of PipD, PipF and PipG. So, structural information based on the predicted structure by AlphaFold3[2] was used to determine their possible functions. Specifically, the isolated single-chain enzyme structured were predicted for further structural search and alignment. The FoldSeek[3] web server was used to search the similar PDB structures in PDB100 database. To filter the search results, the probability of a true positive (TP) match, defined as the probability that two enzymes belong to the same superfamily, was used as the selection criterion.

Considering that there is no significant difference in the predicted structure of AF3 between Annotation by Geneious and Annotation by antiSMASH, in order to minimize the loss of structural information, we always used structures with longer sequences for search in FoldSeek. The screening condition for retrieval results was set to a probability of 1.0 for PipF, as the number of available PDB structures was deemed sufficient for functional and active site analysis. For PipG, because fewer structures with a probability of 1.0 were available compared to PipF, the threshold was relaxed to 0.8 to enable broader structural analysis and comparison. The results are summarized in the following tables. Only PipF and PipG showed significant structural similarities to entries in the PDB100 database. The analysis revealed that many of the structurally similar enzymes are involved in esterification and acetylation (acetyl hydrolysis) reactions. FoldSeek did not retrieve hits for PipD.

**Table S4.2.** FoldSeek results of PipF in PDB100 database.

| Target | Function | Active Sites | Prob. | Score | Query Pos. | Target Pos. |
|---|---|---|---|---|---|---|
| 8GR2 | P-nitrophenyl acetate esterase | Ser138 Asp307 His310 | 1 | 145 | 9-229 (240) | 12-211 (211) |
| 3P94 | Lipase | | 1 | 133 | 6-229 (240) | 18-204 (204) |
| 7TOG | Acetyl xylan esterase | Ser188 Asp363 His366 | 1 | 132 | 9-223 (240) | 171-370 (378) |
| 7DDY | Acetyl xylan esterase | Ser32 Asp200 His203 | 1 | 125 | 11-229 (240) | 2-198 (198) |
| 4HF7 | Lipase | | 1 | 125 | 9-226 (240) | 22-198 (204) |
| 4JGG | Lysophospholipase | Ser9 Asp156 His159 | 1 | 123 | 11-226 (240) | 2-174 (180) |
| 2HSJ | | | 1 | 123 | 4-226 (240) | 29-211 (214) |
| 4RSH | | | 1 | 120 | 11-226 (240) | 3-173 (175) |
| 1YZF | Lipase / Acylhydrolase | | 1 | 120 | 10-232 (240) | 2-194 (195) |
| 7PZG | Sialic acid acetylesterase | Ser52 Asp199 His202 | 1 | 118 | 5-229 (240) | 17-200 (200) |
| 4S1P | Hydrolase | | 1 | 114 | 8-232 (240) | 1-184 (184) |
| 4IYJ | Acylhydrolase | | 1 | 114 | 6-226 (240) | 22-202 (208) |
| 4PPY | Acylhydrolase | | 1 | 114 | 6-227 (240) | 16-201 (208) |
| 4OAO | Acetyl-xylooligosaccharide esterase | | 1 | 111 | 10-226 (240) | 7-209 (219) |
| 4HYQ | Phospholipase | Ser11 Ser 216 His218 | 1 | 110 | 8-228 (240) | 1-236 (236) |
| 4JHL | Acetyl-xylooligosaccharide esterase | Ser15 Asp191 His194 | 1 | 107 | 10-226 (240) | 7-209 (219) |
| 4JKO | Acetyl-xylooligosaccharide esterase (4jhl) | | 1 | 101 | 10-226 (240) | 7-209 (219) |
| 7TJB | Peptidoglycan O-acetyltransferase | | 1 | 95 | 8-234 (240) | 12-220 (222) |
| 4Q9A | Lipase | | 1 | 94 | 9-226 (240) | 8-213 (218) |
| 7TLV | Peptidoglycan O-acetyltransferase | | 1 | 94 | 8-234 (240) | 12-217 (219) |
| 1WAB | Acetylhydrolase | Ser 47 Asp192 His195 | 1 | 89 | 10-224 (240) | 35-204 (212) |

**Table S4.3.** FoldSeek results of PipG in PDB100 database.

| Target | Function | Active Sites | Prob. | Score | Query Pos. | Target Pos. |
|---|---|---|---|---|---|---|
| 4H08 | Hydrolase | | 1 | 117 | 140-316 (344) | 60-197 (200) |
| 5B5S | Acetylesterase | Ser10 Asp179 His182 | 1 | 99 | 39-315 (344) | 2-201 (207) |
| 2HSJ | | | 1 | 99 | 159-313 (344) | 85-213 (214) |
| 1BWR | Acetylhydrolase | Ser47 Asp192 His195 | 1 | 92 | 147-315 (344) | 74-210 (212) |
| 5B5L | Acetylesterase (S10A of 5b5s) | | 1 | 91 | 39-315 (344) | 2-201 (207) |
| 7E16 | Esterase | | 1 | 91 | 36-309 (344) | 1-200 (207) |
| 7BXD | | | 1 | 91 | 160-316 (344) | 75-217 (219) |
| 8IK1 | | | 0.99 | 88 | 36-312 (344) | 1-198 (204) |
| 1BWQ | Acetylhydrolase | Ser47 Asp192 His195 | 0.99 | 88 | 147-315 (344) | 74-210 (212) |
| 1VYH | Acetylhydrolase | Ser48 Asp193 His196 | 0.99 | 85 | 145-317 (344) | 73-215 (218) |
| 4HF7 | Lipase | | 0.99 | 83 | 135-315 (344) | 52-202 (204) |
| 3SKV | Salicylyl-acyltranferase | Ser174 Glu330 His338 | 0.99 | 83 | 152-315 (344) | 205-347 (354) |
| 4XVH | Carbohydrate esterase | | 0.98 | 79 | 160-275 (344) | 205-293 (329) |
| 4Q9A | Lipase | | 0.98 | 78 | 33-314 (344) | 1-216 (218) |
| 7PZG | Sialic acid acetylesterase | Ser52 Asp199 His202 | 0.97 | 76 | 140-317 (344) | 57-199 (200) |
| 3DC7 | | | 0.97 | 76 | 158-316 (344) | 64-206 (208) |
| 4K9S | Peptidoglycan O-acetylesterase | Ser80 Asp366 His369 | 0.97 | 76 | 135-315 (344) | 189-344 (349) |
| 7PZH | Sialic acid acetylesterase | Ser52 Asp199 His202 | 0.96 | 74 | 203-314 (344) | 88-196 (197) |
| 6NKD | Lipase | | 0.93 | 70 | 153-312 (344) | 64-204 (210) |
| 1ES9 | Acetylhydrolase | Ser47 Asp192 His195 | 0.93 | 70 | 67-315 (344) | 6-210 (212) |
| 1BWP | Acetylhydrolase | Ser47 Asp192 His195 | 0.92 | 69 | 67-315 (344) | 6-210 (212) |
| 3DT8 | Acetylhydrolase | Ser47 Asp192 His195 | 0.91 | 68 | 67-315 (344) | 6-209 (211) |
| 3DT9 | Acetylhydrolase | Ser47 Asp192 His196 | 0.91 | 68 | 67-315 (344) | 6-210 (212) |
| 1WAB | Acetylhydrolase | | 0.85 | 64 | 67-315 (344) | 6-210 (212) |
| 6SE1 | Acyltransferase | Ser430 Asp618 His621 | 0.84 | 63 | 35-314 (344) | 39-260 (261) |
| 3R7W | Target of rapamycin | | 0.82 | 62 | 154-277 (344) | 67-156 (294) |

##### 3.2. Identification of PipF and PipG active sites

The identification of the active sites in PipF and PipG started with a structural comparison against similar known structures (Tables S4.2 and S4.3). For PipF, structural alignment revealed a strong overlap with known enzyme active sites (Fig. S4.1a-c), leading to the conclusion that Ser18-His211-Asp208 most likely forms the catalytic triad of PipF.

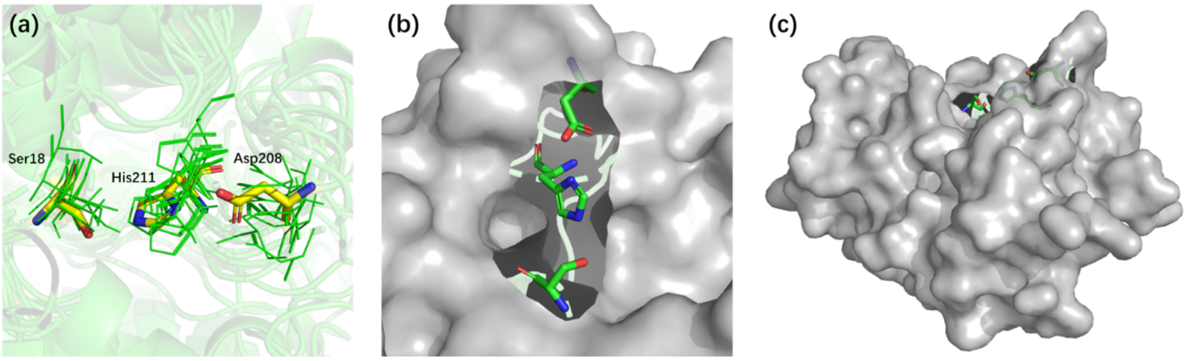


**Figure S4.1. Predicted PipF active sites. (a)** The inferred PipF active site is aligned with the active sites of enzymes reported in the FoldSeek search results in the PDB100 dataset. Green lines: active sites in PDB100; yellow sticks: inferred PipF active sites. **(b) and (c)** The inferred location of the PipF active site on the enzyme surface.

For PipG, the analysis proved more challenging. Structural alignment only provided an approximate location of the active site pocket, and a conserved Ser-His-Asp catalytic triad was not observed in the corresponding region. To infer potential catalytic residues, all surface-exposed residues near the predicted active pocket were considered as candidates (Figure SI-Note 2a). After excluding residues unlikely to participate in catalysis or those presenting steric hindrance, four potential catalytic sites remained (Figure SI-Note 2b and 2c). From these, two possible catalytic triad combinations were identified: Cys47-His51-Glu296 and Thr238-His51-Glu296. Given the closer spatial arrangement of Cys47, His51, and Glu296, we propose that Cys47-His51-Glu296 represents the most likely catalytic triad in PipG.

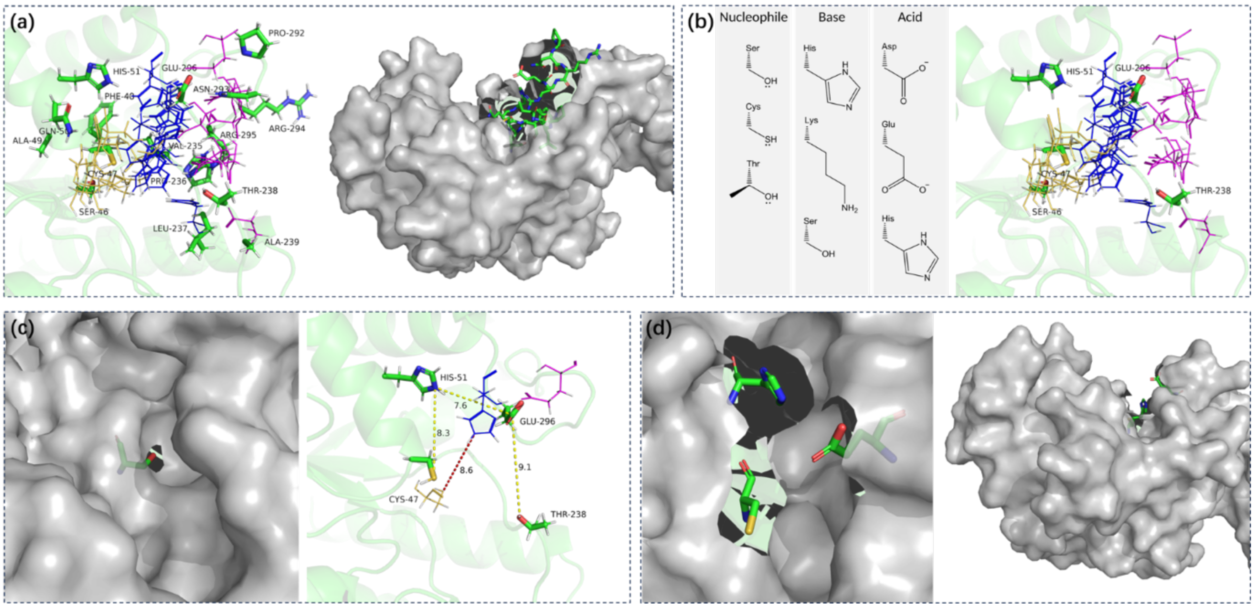


**Figure S4.2. Predicted PipG active sites.** (**a**) The approximate location of the active pocket determined by structural comparison based on the search results of FoldSeek in the PDB100 dataset. All the residues of PipG near the pocket and exposed on the surface were selected as potential active sites set. Yellow lines: Ser; Blue line: His or Glu; Purple line: Asp; Green sticks: residues on the PipG inferred pocket surface. (**b**) Based on the common catalytic triad combination (left), five residues including Ser46, Cys47, His51, Thr238 and Glu296 were screened in the potential active sites set (right). (**c**) Ser46 is very poorly exposed on the surface and is sterically hindered from other residues compared to Cys47, which is also a nucleophile (left). The distance (yellow dashed line) between the putative PipG catalytic residues and the maximum distance (red dashed line) between the catalytic residues in PDB: 3SKV (right). (**d**) Positioning of the Cys47-His51-Glu296 catalytic triad on the surface of PipG.

##### 3.3. Protein interactions among PipD, PipF, and PipG

As PipD, PipF, and PipG are all required for the final step acetylation, we explored a possible interactions between them. Although AlphaFold3 has achieved accurate predictions of protein structures and even complex protein structures, it cannot determine whether a protein is a monomer or a polymer in actual conditions based solely on sequence information. Users are still required to provide the number of each chain in the complex protein. The oligomeric properties of a protein are difficult to predict. Direct experimental evidence would be most relevant, but partly homologous structural information can be helpful, which, however, for PipD, PipF, and PipG is currently not available. Regarding the lack of these details, we modeled the heterooligomeric complex with proteins in monomeric form. Indeed, homodimerization does likely not interfere with heterooligomeric interfaces, because proteins evolve to avoid clashes between their different binding partners, such that the same surface is usually not used for building both homo- and hetero-interfaces.

For PipD and PipF, structures based on different annotation sequence lengths were predicted and compared. This comparison revealed no significant structural differences between the models generated from sequences obtained by different annotation tools. In order to ensure the information is as complete as possible, we chose the longer sequences annotated by antiSMASH for the final dimer and trimer structure predictions. Fig. S4.3a illustrates the comparison of the predicted structures. As shown, PipD and PipF exhibit the same binding mode in both the dimeric and trimeric models (cyan and green). In contrast, the binding mode of PipG with the other two proteins differs noticeably in the trimeric structure (cyan with purple or yellow).

To further assess the stability of protein-protein interactions across different assembly modes, we conducted MD simulations along 100 ns trajectories and calculated the RMSD of the C$\alpha$ atoms (Fig. S4.3b). MD simulations were conducted with GROMACS (version 2022.3) using the Amber99SB-ildn force field and the TIP3P water model. The enzyme was centered in a cubic box with 1 nm between the solute and the box. The system was neutralized with NaCl. Energy minimization was performed applying the steepest descent algorithm until a maximum force of 1000 kJ mol$^{-1}$ nm$^{-1}$ on any atom was reached. The system was equilibrated by a 1 ns NVT run at 310 K, followed by a 1 ns NPT run at 1 atm and 310 K. Pressure and temperature were controlled using the Velocity-rescale and parrinello-Rahman algorithms. All systems were simulated for 100 ns. The RMSD of Cα was calculated.

The data show that, except for the dimer composed of PipD and PipF, there are no substantial differences in RMSD values among the different assembly modes. The PipD-PipF dimer exhibits the most stable and “rigid” conformation, while complexes with PipG generally appear more conformationally flexible, likely due to the inherent pronounced conformational variability of PipG caused by more unstructured regions. Regardless of the assembly mode, the overall structure of the protein complexes remained stable, with no signs of dissociation or secondary structure collapse. This observation confirms that a stable hetero-oligomeric arrangement of PipD, PipF, and PipG is plausible. The alignment of PipF, PipG and PDB structures was performed using Structure Comparison/MatchMaker tool in Chimera [4] software.

It should be noted, that a protein aggregation effect may occur during MD simulations, potentially mimicking complex formation, which should also be evaluated and ruled out in more focused studies. Finally, direct experimental evidence is necessary to confirm a function complex of PipD, PipF and PipG.

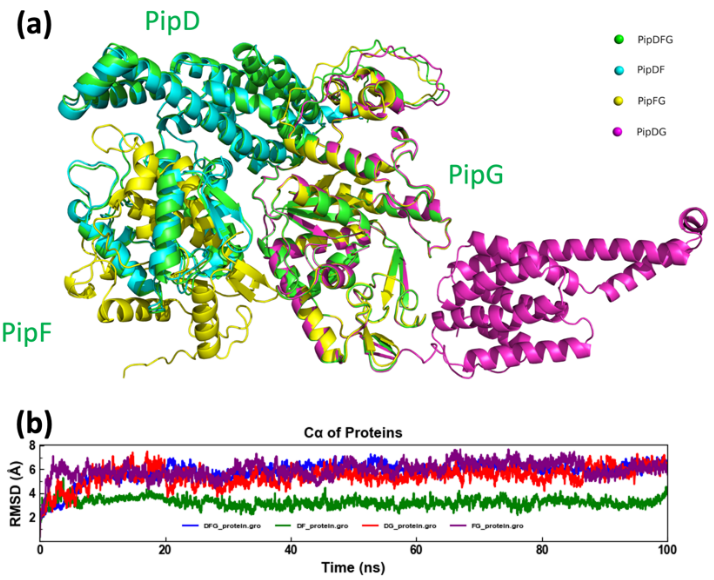


**Figure S4.3. PipDFG structural model. (a)** Comparison between different prediction structures of heterodimeric and heterotrimeric complexes. In the PipDFG, proteins are labelled. (b) RMSD of MD simulation trajectories of four assembly modes
